# Genetic diversity analysis and characterization of Ugandan sorghum

**DOI:** 10.1101/2022.01.31.478463

**Authors:** Subhadra Chakrabarty, Raphael Mufumbo, Steffen Windpassinger, David Jordan, Emma Mace, Rod J. Snowdon, Adrian Hathorn

**Affiliations:** Department of Plant Breeding, Justus Liebig University, Giessen, Germany; Queensland Alliance for Agriculture and Food Innovation, University of Queensland, Warwick, QLD, 4370, Australia

**Keywords:** Sorghum bicolor, genetic diversity, population structure, cold tolerance, temperate climate adaptation, genome wide association study, genebank

## Abstract

The National Genebank of Uganda houses diverse and rich *Sorghum bicolor* germplasm collection. This genetic diversity resource is untapped, under-utilized and has not been systematically incorporated into sorghum breeding programs. In this study, we characterized the germplasm collection using whole genome SNP markers. Discriminant analysis of principal components (DAPC) was implemented to study racial ancestry of the accessions in comparison to a global sorghum diversity set and characterize sub-groups and admixture in the Ugandan germplasm. Genetic structure and phylogenetic analysis was conducted to identify distinct genotypes in the Ugandan collection and relationships among groups. Furthermore, in a case study for identification of potentially useful adaptive trait variation for breeding, we performed genome-wide association studies for juvenile cold tolerance. Genomic regions potentially involved in adaptation of Ugandan sorghum varieties to cooler climatic conditions were identified that could be of interest for expansion of sorghum production into temperate latitudes. The study demonstrates how genebank genomics can potentially facilitate effective and efficient usage of valuable, untapped germplasm collections for agronomic trait evaluation and subsequent allele mining.

## 1. Introduction

*Sorghum bicolor* [L.] Moench (sorghum) is the fifth most important cereal crop globally and shows remarkable diversity, including five different races, their intermediates and several crop forms classified as grain, forage, sweet and broomcorn types (Hariprasanna and Patil 2015). Sorghum has extraordinary untapped variation in grain type, plant type, adaptability, productive capacity and underutilized genetic potential (Lost Crops of Africa 1996). Because of its wide adaptability to drought and heat, sorghum’s importance is expected to increase with the changing global climate and an ongoing increase in the use of marginal lands for agriculture (Paterson et al. 2009).

In Eastern Africa sorghum is traditionally grown as a food-fodder crop by small holder farmers in low-input agricultural systems spanning highland, lowland and semi-arid cropping regions. Uganda is one of only three countries in which all of the five basic races and ten intermediate races of *S. bicolor* are endemic (Reddy et al. 2002). The country’s broad sorghum diversity reflects the variety of environments where the crop is grown, mainly on marginal agricultural lands ranging from extremely arid and semi-arid zones in eastern and northern Uganda to cool highlands in south-western regions. The Uganda National Genebank houses a large collection of *S. bicolor* accessions, including a vast range of landraces whose diversity has yet to be capitalized for use in breeding. This germplasm has not yet been fully characterised and evaluated, limiting its utilisation to date in sorghum improvement programs in Uganda and elsewhere. The considerable geographical and topological diversity of Uganda makes this genetic resource a potentially interesting reservoir for genetic analysis and diversity for adaptive traits of interest for sorghum breeding. For example, cool highland areas in the southwestern Uganda are potential sources of diversity for cold-tolerance traits that could help improve sorghum adaptation in temperate cropping regions.

Effective and efficient management of germplasm from a genebank collection is an essential prerequisite for farmers and breeders to identify, extract and exploit the extensive diversity. Genome-wide characterization of untapped genetic resources using genome sequencing technologies provides new opportunities to sustainable breeding and efficient usage of material.

In this present study, the diverse Uganda National Genebank *S. bicolor* collection, representing different agro-ecological zones of the country, was investigated using whole genome SNP markers and population genetic analysis. The primary objective was to genetically characterize Ugandan sorghum germplasm in the context of global sorghum diversity. As a case study for the value of this resource in adaptive breeding, we furthermore used the available genome data, in association with phenotypic data for juvenile cold tolerance traits, to identify genomic regions enriched with genetic variants associated to low temperature adaptation. The results demonstrate how genebank genomics can help facilitate discovery of economically or biologically important plant diversity and genes as a prerequisite for crop genetic improvement and climatic adaptation.

## 2. Materials and Methods

### Plant materials

A total of 3333 diverse Ugandan *S. bicolor* germplasm accessions (UG set) collected by the Plant Genetic Resources Centre at the Uganda National Genebank were used in this study (Table S1). This germplasm collection represents the entire sorghum diversity from all eco-geographical regions of Uganda. For racial composition analysis, the collection was compared to a global sorghum germplasm collection of 1033 genotypes (global set) which was previously described by (Tao et al. 2020).

### DNA extraction and genotyping

DNA was extracted from seeds by Diversity Arrays Technology Pty Ltd. (www.diversityarrays.com) and genotyped using DArTseq, an efficient genotyping-by-sequencing (GBS) platform which enables discovery of genome-wide markers through genome complexity reduction using restriction enzymes. Subsequently, sorghum reference genome version v3.1.1 (McCormick et al. 2018) was used for sequence alignment and single nucleotide polymorphism (SNP) calling.

### SNP data filtration

A total of 40,290 SNP markers were reported for the global and UG sets. Firstly all nonspecific markers and those belonging to supercontigs were removed. The remaining 34,469 were used for further analysis. For racial ancestry analysis, an extremely high stringency was then applied to remove markers which were duplicated or exhibited greater than 1% missing data and genotypes showing greater than 1% missing data. The global diversity set comprised conversion lines containing introgressed chromosome regions from the Sorghum Conversion Program conducted by Texas Agricultural Experiment Station (Thurber et al. 2013). Hence, to reduce disparity with UG samples in the co-analysis with the global set, all markers from the specific genomic regions impacted by the conversion program were excluded as follows: Sb06: all markers, Sb07: all markers beyond 40Mb, Sb09: all markers beyond 46Mb. Markers with minor allele frequency (MAF) less than 0.01 were also excluded. This set was then imputed using Beagle 5.1 (Browning, Zhou, and Browning 2018) to infer the remaining missing data values. A total of 2,331 common markers between the UG and global set were used for the racial ancestry analysis.

### Population structure and genetic diversity study

In order to understand the racial classification and the population structure of the UG sorghum collection, principal component analysis (PCA) and Discriminant Analysis of Principal Components (DAPC) were implemented using the R package Adegenet (2.1.3) (Jombart, Devillard, and Balloux 2010). To avoid bias caused by the large size of the UG set, we initially used only a representative subset of the UG set to assign racial groupings in comparison to the global collection. Hence, prior to racial group identification of the UG set in comparison to the global set, K-mean clustering was performed to select UG genotypes representative for all identified groups. Ten clusters were identified using the ‘find.clusters’ function. From each of these clusters, ten randomly-selected samples were combined with the global dataset for DAPC co-analysis. To validate the racial assignment of the UG set groups, the DAPC co-analysis was repeated three times, each time a different random selection of 10 genotypes from each identified group. The SNP data was converted to the genlight object bit-level genotype coding scheme using the function ‘vcfR2genlight’ of the vcfR tool (https://github.com/knausb/vcfR). After an initial transformation using the PCA analysis, clusters were subsequently identified using discriminant analysis (DA).

To describe the population structure of the UG germplasm and evaluate the distribution of racial groups in relation to geographical origins of accessions, the filtered marker set (34,466) was pruned to exclude SNPs in strong LD using PLINK software (Purcell et al. 2007) SNPs were pruned with a window of 50 SNPs, step size of 5 makers and r^2^ threshold of 0.5. A total of 12,742 markers for all UG lines were used to analyse population structure using Discriminant analysis (DA). To elaborate the genetic relationship among the accessions a pairwise distance matrix was established using the tool VCF2Dis (https://github.com/BGI-shenzhen/VCF2Dis), which was then converted to a neighbour-joining phylogenetic tree using the R package ape (5.5) (Paradis, Claude, and Strimmer 2004) and visualized with R package ggtree (3.0.4) (Yu 2020). Weir and Cockerham’s Fst (wcFst) (Weir and Cockerham 1984) was calculated among the UG subpopulations using the function ‘--weir-fst-pop’ of vcftools (0.1.17) to identify patterns of genetic differentiation.

### Phenotyping and association mapping

For evaluating juvenile survival under cold stress, two field trials at Gross Gerau (GG), Germany (spring 2019 and 2020) and one climate chamber experiment were conducted on a UG subset of 444 (field trials) and 255 (climate chamber) accessions representing all agro-ecological zones (Table S1) For the field experiments (Table S2), all genotypes were sown in micro-plots consisting of single rows (2.5 × 0.7 m) using an alpha lattice block design with two replications. Being cold stress experiments, sowing times were several weeks earlier than normal for sorghum in that area. Even though the mean soil temperatures were relatively high during the course of the experiments (13.8 and 16.2 °C, respectively) and allowed for a satisfying emergence, in both years several cold nights (up to -1.5 °C) implied strong stress on the seedlings. Around four weeks after emergence, when the last cold event lay 7 days back, the number of surviving plants was scored per plot. For subsequent GWAS, the alpha-lattice adjusted mean value of both years was used. For the climate chamber experiment (CC), 16 seedlings per genotype were established in 12 × 12 × 12 cm pots. Experiments were designed as randomized complete block design with four replications (Table S3) and the number of surviving plants per pots was scored as a measure for juvenile cold stress tolerance.

After eliminating markers and genotypes with more than 25% missing data points, the dataset was imputed with using Beagle 5.1 (Browning, Zhou, and Browning 2018). It was further corrected for MAF lower than 5%. A total of 4,099 markers were used for genome wise association study (GWAS) implemented in R package GenABEL (1.8-0) (Aulchenko, de Koning, and Haley 2007) for juvenile cold tolerance traits.

Population stratification was adjusted by incorporating two frequently used models: principal coordinate and genomic kinship matrix corrections principle component and kinship analysis (Stich et al. 2008). The correlations between each marker and trait can be well identified by these models after variation due to population structure has been controlled.

To reduce the type II error rate and in order to classify a marker–trait association as significant, a threshold of − log10 (p value) ≥ 3.0 was defined (Gabur et al. 2019). Linkage disequilibrium across the entire genome was calculated using the squared allele frequency correlations (r^2^) between each pair of SNPs. Haplotype blocks were calculated using an LD threshold of r^2^ > 0.7 to define blocks, as described by (Gabriel et al. 2002).

Candidate genes were selected based on *Sorghum bicolor* reference genome *v3*.*1*.*1* hosted by Phytozome 12 (https://phytozome.jgi.doe.gov/pz/portal.html#!info?alias=Org_Sbicolor). Orthologous genes within selected haploblocks were identified by homology comparisons of the genomic sequences in maize and rice. Protein sequence alignments were conducted by using blastp option of DIAMOND (Buchfink, Xie, and Huson 2015).

## 3. Results

### Genetic diversity and population structure analysis

Appropriate conservation and effective utilization of novel UG germplasm can be facilitated by understanding the underlying variation and genetic diversity. DAPC between the global and the UG set revealed the presence of all racial groups for the latter (Figure 1). The majority of the accessions could not be directly classified to any particular racial cluster, but rather showed intermediate classifications between two racial groups, indicating the presence of admixture. This was further confirmed by Kmean grouping (Figure 1b). All racial groups were represented in each of the ten identified Kmean clusters, however cluster two showed a strong overrepresentation of caudatum race.

**Figure 1:**
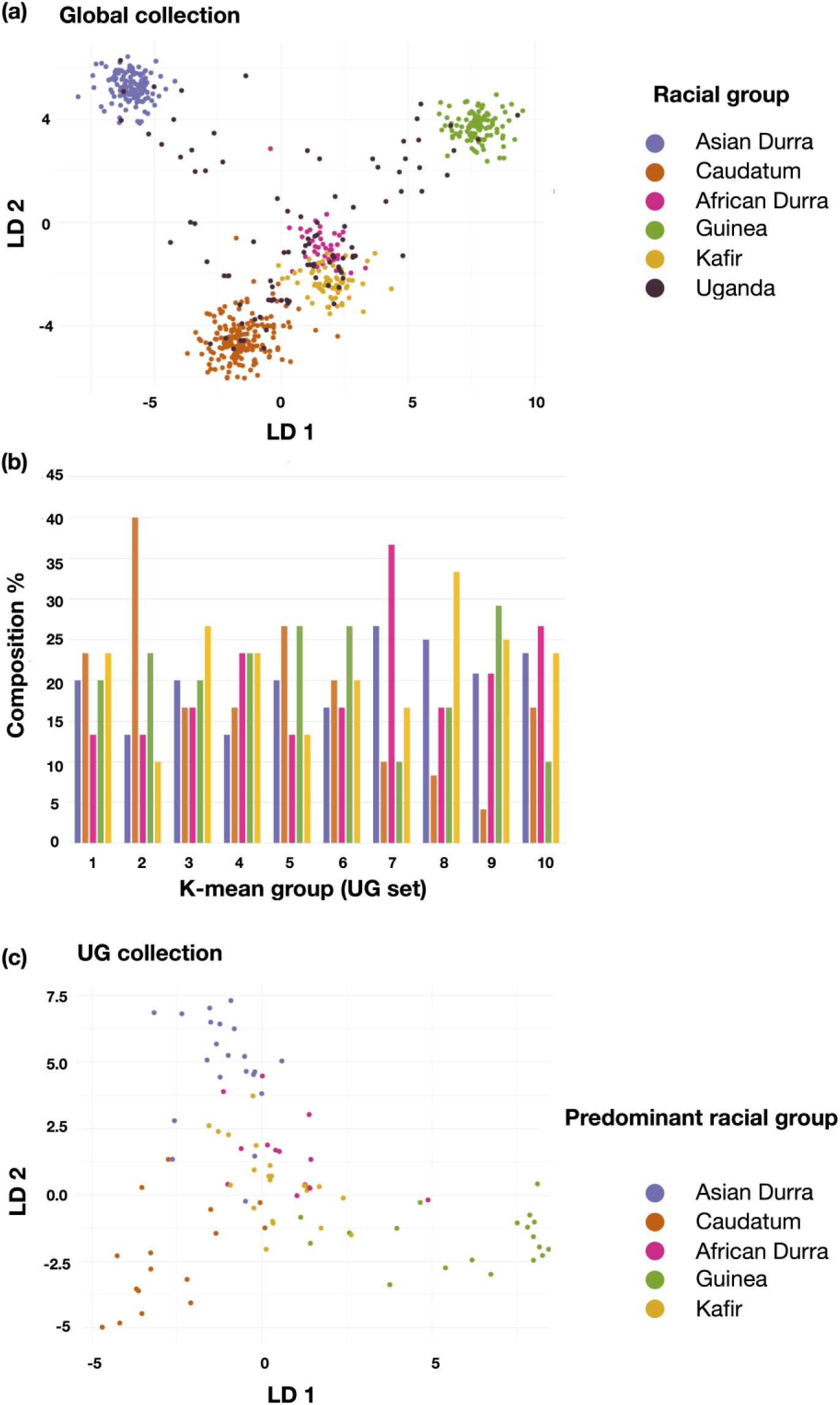
Scatterplot and composition of discriminant coefficients of the DAPC analysis for racial group determination of UG samples based on global diversity set. (a) DA loadings (LD) displaying UG clustering in comparison to global racial groups. (b) Imputed racial composition of ten identified Ugandan Kmean clusters. The x axis represents the clusters whereas the y axis indicates the racial composition in percentage for each cluster. (c) DA loading of Ugandan representative accessions based on races. Each dot represents an individual and the colour code is displayed in the index.

Results from Kmean clustering and DAPC using the three random samples from UG groups co-analysed with the global reference collection confirmed the racial distribution and presence of admixture across the UG diversity set (Figure 2, Figure S1, Table S4).

**Figure 2:**
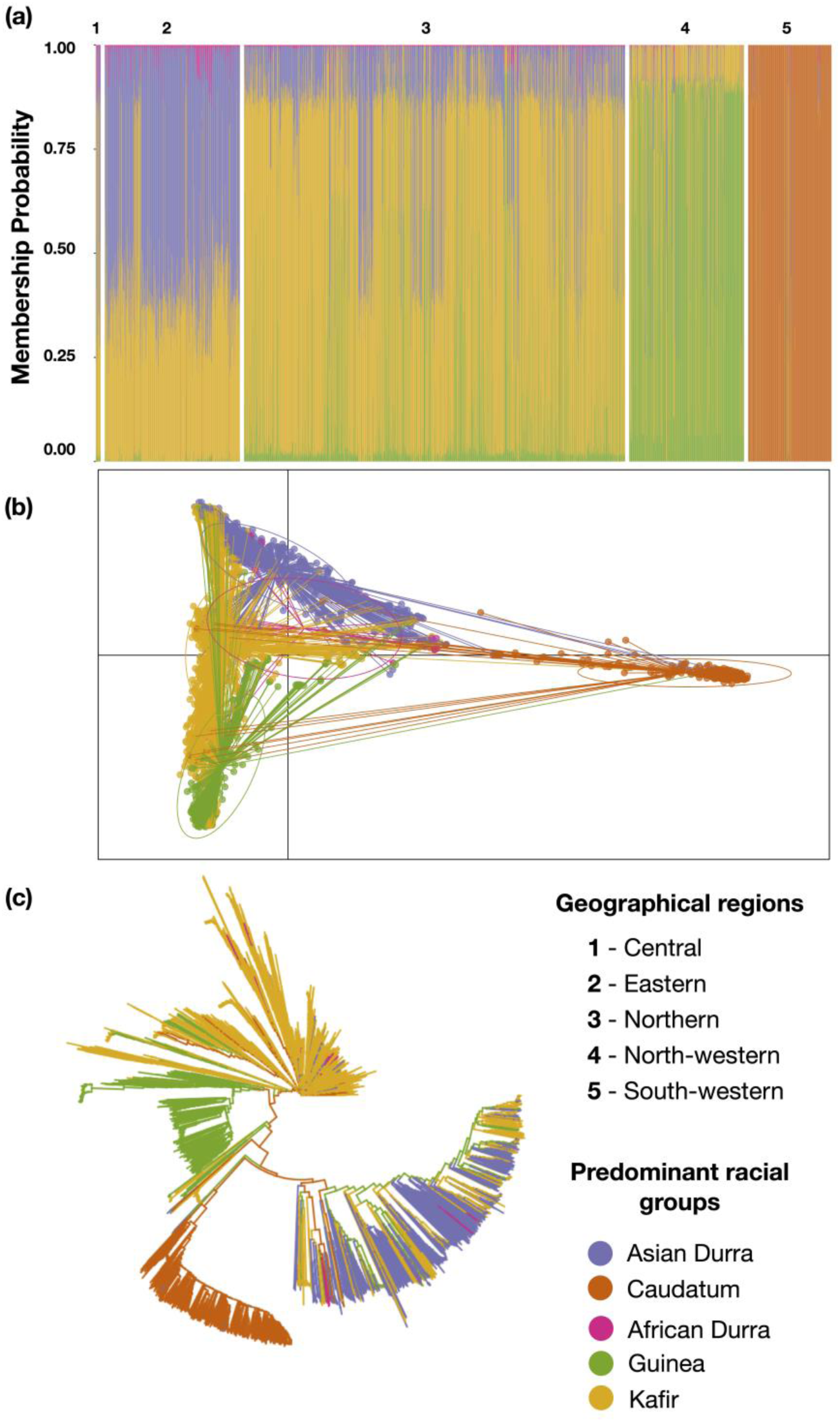
DAPC analysis of the Ugandan genebank sorghum collection. (a) Barplot, (b) Scatter plot and (c) phylogenetic tree analysis of subpopulations. Colours represent predominant racial group assignment of each accession. The first two linear discriminants (LDs) are represented by the axes of the scatterplot. In (a) each vertical bar represents one individual accession, coloured based on the identified racial classification. Circles in (b) represent identified clusters corresponding to racial groups.

To better understand the relationships among the Ugandan accessions based on their imputed racial classification and the geographical origin, a DAPC analysis was performed only for the UG set (Figure 2). Genotypes from the central, eastern, northern and north-western geographical regions were found to be predominantly admixtures, whereas the southwestern region (dominated by genotypes of highland origin) showed a distinct genetic pattern with a predominance of caudatum race (Figure 2a,b). Four hundred PCs (27.4% of variance conserved by first three PCs) and four discriminant eigenvalues were retained in the DAPC analysis in order to capture maximum variance. Overall, diversity in the UG collection is defined more by racial grouping rather than geographical grouping (Figure 2a,b) A phylogenetic tree confirmed the divergence of south-western accessions (Figure 2c).

Due to significant adaptability differences between the lowland and highland races from the Ugandan diversity set, genetic diversity among the sub groups within the population were tested. Fst revealed overall genetic variance of 0.30 (ranging from -0.002 to 0.95), indicating presence of significant genetic structuring or population subdivision within the germplasm.

### Phenotypic variation for juvenile cold stress survival, association mapping and haplotype analysis

A genome-wide association study (GWAS) was conducted for identification of regions of interest associated to juvenile survival under cold stress in two temperate-climate field environments and one controlled-environment climate chamber test. Highly significant differences for cold tolerance among genotypes (p=0.000***, Table S5) were found in all experiments. Comparing lowland- and highland genotypes as groups by a one-way ANOVA, highland genotypes showed a superior cold tolerance in all experiments as expected (p=0.015* for GG19, p=0.000*** for GG20, p=0.000*** for CC).

Significant marker-trait associations to juvenile survival under cold stress, consistent across all test environments, were identified on chromosomes Sb02, Sb06, and Sb09 (Figure 3 a, b). We compared these selected genomic regions with previously curated QTL in sorghum QTL-Atlas (Mace et al. 2019), based on physical position (v3.0) and filtered using category resistance abiotic and subcategory cold tolerance. A sum of nineteen overlapping QTL involved in juvenile cold tolerance were identified (Table S6) in these regions. This result affirms the important role of these genomic regions towards cold adaptation.

**Fig 3:**
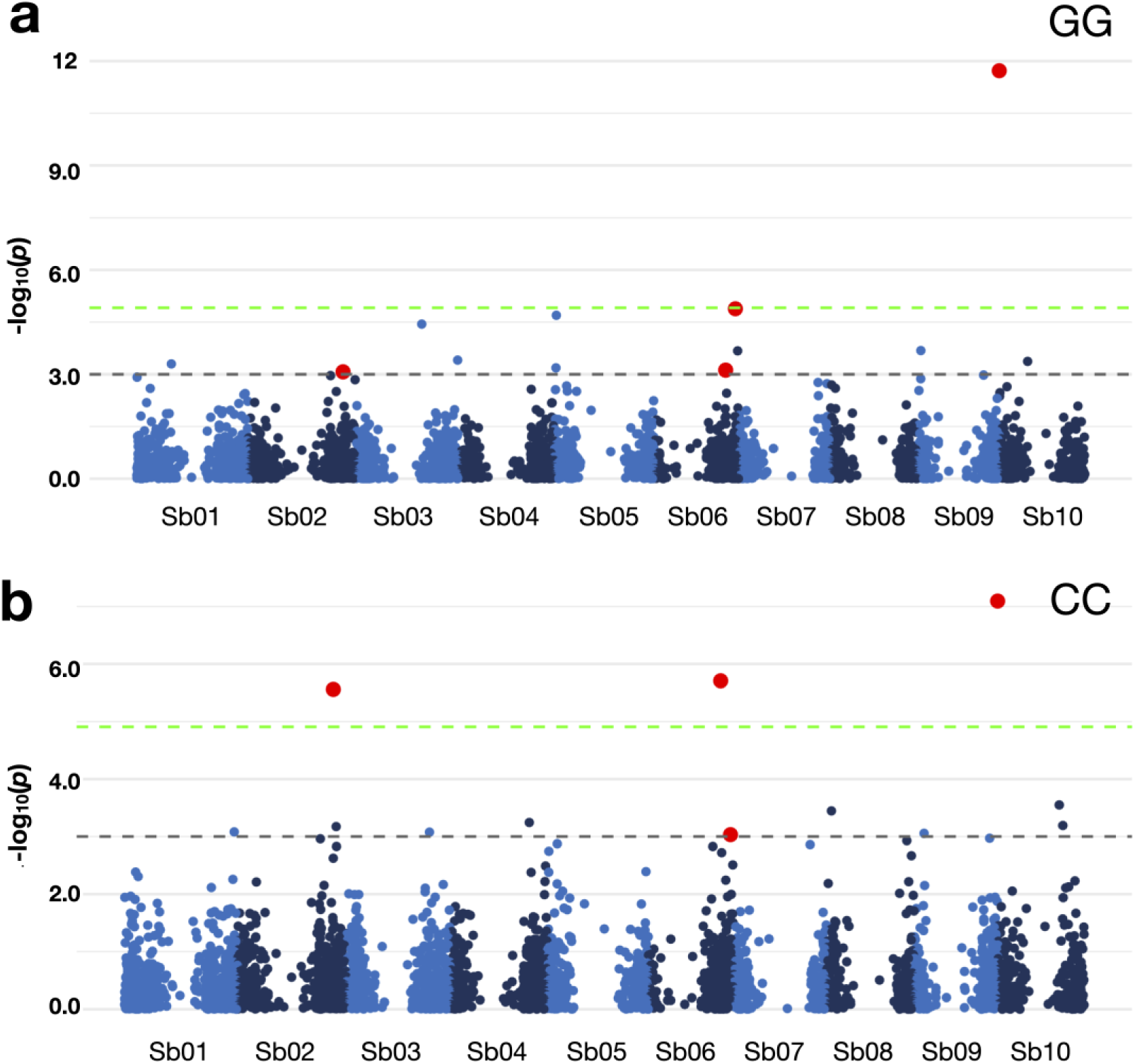
Association mapping of survival under cold stress under (a) field condition (Gross Gerau; GG) and (b) climate chamber (CC). The grey dotted horizontal line indicates a threshold of genome-wide cut-off at -log10(p) > 3.0 while the green line indicates the Bonferroni threshold at -log10(p) > 4.9. The selected associated markers which overlapped under both conditions are marked in red.

Comparing orthologous gene segments of sorghum, maize and rice, six genes involved in various cold acclimatization and tolerance were identified (Table S7). For example, a F-box protein-encoding gene, *Sobic*.*006G245200* was identified at a distance of 9kb from the associated marker Sb06-58472351. This gene family has been shown to be upregulated under cold stress in rice, *Brassica rapa* and pepper (Jain et al. 2007; Rameneni et al. 2018; Venkatesh et al. 2020).

## Discussion

Plant genetic resources like the sorghum collection of the National Genebank of Uganda are extremely important public germplasm resources for local breeders in crop centres of origin and the agricultural and crop research community worldwide. The substantial variation we identified in the Ugandan *S. bicolor* germplasm reflects the highly diverse environments where sorghum grows in Uganda. Accessions with adaptation to drought and heat stress from the arid regions of Northern and Eastern Uganda, and germplasm from the cold highlands of the South-Western Kigezi region, carry potentially useful genetic diversity for heat and cold stress adaptation traits. The latter are of major interest for breeding programs aiming to expand sorghum production into temperate climates in North America, Asia and Europe, whereas heat and drought stress tolerance are becoming increasingly important for global sorghum production in the face of climate change. The rich genetic resource of the Ugandan genebank sorghum collection has not yet been fully characterised and evaluated, limiting its utilisation to date in sorghum improvement programs in Uganda and elsewhere.

Genetic structure has been previously documented in sorghum in smaller germplasm collections (Westengen et al. 2014; Mofokeng et al. 2014; Cuevas and Prom 2020). This study reports genetic diversity in a substantial population of 3333 accessions from a key sorghum centre of origin, comparing it to previously described sorghum diversity by co-analysis with a global diversity set (Tao et al. 2020).

The DAPC method is a good alternative to other population structure analysis software such as STRUCTURE because of its ability to deal with large datasets. Clustering of genotypes presented in this study provide interesting leads for increasing diversity in breeding programs and germplasm utilization. Overall, results from DAPC, Kmean and phylogenetic tree analysis were in agreement and provided evidence that the global racial diversity was well-covered in the UG germplasm. However, considerable admixture between racial groups was identified, particularly between genotypes from lowland areas in Central, Eastern and Northern Uganda, indicating frequent gene flow between these regions. In contrast, the south-western region appeared genetically distinct and was comprised predominantly of Caudatum germplasm. Because this region includes cool-temperature highland areas it may contain interesting adaptive diversity for juvenile or reproductive cold tolerance. Population structure and phylogenetic analysis revealed the distinctness of this group compared to the other eco-geographical regions, presumably due to the distinctness of the highland cultivation environment and a relatively low exchange of germplasm between highly and lowland farmers. Similarly, (Mekbib 2008) reported the existence of caudatum and its intermediate races in the highlands of Ethiopia. This confers with the theory that the spread and diversification of crops to different locations can lead to new variants, a process influenced by genotype by environment interactions and geographical isolation (Pressoir and Berthaud 2004). Based on the principles of artificial selection (Yamasaki et al. 2007), farmers select cultivars with preferred traits and thus increase their proportion compared to other cultivars. According to (Akatwijuka et al. 2016), sorghum from south-western Uganda tend to have semi-compact elliptic panicles, a well-known characteristic of caudatum and its intermediate races.

Sorghum has a high potential for adaptation to a wide range of environmental conditions. Besides yield and other agronomic traits, the improvement of cold tolerance at juvenile and reproductive stages (Schaffasz2019(a,b); S. Chakrabarty et al. 2021) is a major breeding objective for sorghum temperate cropping regions. Early seedling vigour is critical for crop establishment in any environment (Xie et al. 2014) and vigorous germination and growth under low temperatures is essential for early establishment and weed competition in temperate climates. Improving cold tolerance in the early juvenile stage allows higher yield potential and better maturity. Sharifi (2008) observed that survival percentage, germination, and chlorophyll content were good indicators of early cold tolerance in rice. The yield of sorghum is highly temperature dependent, especially between sowing and flowering time (Craufurd et al. 1999; Fiedler et al. 2014). Hence, breeding for juvenile cold tolerance is of utmost importance, especially for temperate European climates. In this study, juvenile survival under low temperature was studied for multi environments. In contrast to emergence and juvenile biomass under cold conditions which have been extensively studied in several publications (e. g. Burow et al. 2011, Fiedler et al. 2014, Schaffasz et al. 2019), the trait juvenile survival has received much less attention so far. Though, it is of utmost importance, because a satisfying emergence is worthless if the seedlings later succumb to cold stress. Promising hotspots for cold tolerance during juvenile development were identified. Since population stratification was accounted for before performing GWAS, we have reduced the likelihood that of genetic background effects are generating spurious associations. We also learned that multiple QTL identified in previous studies (Mace et al. 2019) and genes known to be involved in cold stress endurance were physically co-located with our QTL, suggesting the importance of our selected genomic regions in this regard.

The results from this study therefore pinpoint interesting plant materials and genome regions containing alleles which can be mined for improvement of cold temperature adaptation. Given the complex genetics underlying juvenile cold tolerance (Burow et al. 2011; Bekele et al. 2014; Fiedler et al. 2014) the implementation of genomic prediction appears best suited to improve selection. The data generated in this study represent a promising first step towards the implementation of genomic prediction to enhance cold adaptation traits in sorghum breeding.

## Conclusions

To our knowledge, this is the first extensive study of the unique and large sorghum germplasm collection conserved at the National Genebank of Uganda. The results indicate immense genetic and racial diversity of predominantly admixed accessions, and a unique, relatively genetically isolated resource of caudatum germplasm from the south-western Ugandan highlands. Despite its highly complex nature, juvenile chilling stress is critical for temperate climate adaptation. The discovery of important genomic regions for these traits and high phenotypic variability associated with these regions can potentially enable breeders to enhance early stage chilling tolerance in sorghum. The genomic and phenotypic data collected in this study provide an objective criterion for the selection of accessions for genetic diversity preservation and management, utilization in breeding programs and genetic relationship analysis with other germplasm collections. The results provide important new insight for adaptive crops breeding to support the expansion and stability of sorghum production in the face of increasing abiotic stress constraints and climatic change.

## Acknowledgement

We want to thank Mario Tolksdorf/JLU Experimental Station Gross-Gerau, Germany, Annette Plank and Birgit Keiner (JLU Giessen) for the help in greenhouse.

## Author contributions

SC generated the data, conducted the data analysis wrote the manuscript. RM curated the material and assisted in data collection. SW planned and oversaw field trials and data collection and assisted in data analysis. DJ and EM provided material and assisted in data analysis. RJS conceived the study and edited the manuscript. AH provided ideas and assisted in data analysis.

