## Supplementary figures and images for "Genetic diversity analysis and characterization of Ugandan sorghum"

### Figure S1

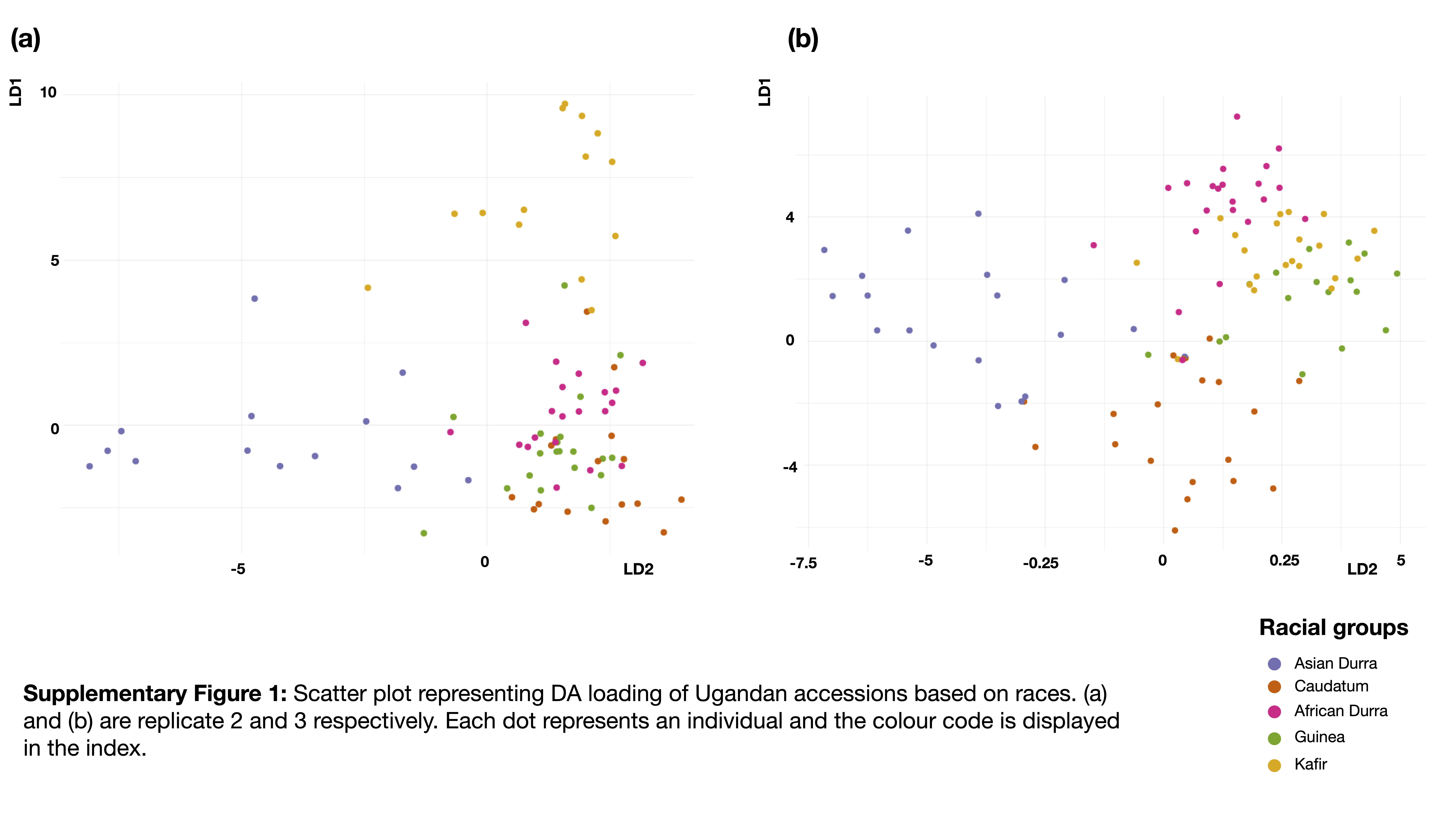
