## Supplementary material for "Genetic diversity analysis and characterization of Ugandan sorghum": Table_S1

Table S1: DAPC analysis of Ugandan lines based on races and geographical origin

| Sample ID | African Durra | Asian Durra | Kafir | Guinea | Caudatum | Geographical_origin |
|---|---|---|---|---|---|---|
| UG_01 | 0.004232767 | 0.607042138 | 0.388408126 | 0.000316969 | 8.73E-15 | Eastern |
| UG_73 | 0.003113858 | 0.569461268 | 0.427131499 | 0.000293374 | 7.42E-16 | Eastern |
| UG_81 | 0.012343179 | 0.671070376 | 0.315956392 | 0.000630053 | 3.67E-11 | Eastern |
| UG_89 | 0.00902292 | 0.179951084 | 0.791308952 | 0.019717043 | 1.47E-12 | Eastern |
| UG_09 | 0.005883148 | 0.475279375 | 0.517715605 | 0.001121872 | 4.87E-14 | Eastern |
| UG_17 | 0.003376067 | 0.616949911 | 0.379443739 | 0.000230283 | 1.82E-15 | Eastern |
| UG_25 | 0.084753258 | 0.675370738 | 0.236389629 | 0.003359345 | 0.00012703 | Eastern |
| UG_33 | 0.005988347 | 0.510006671 | 0.48310107 | 0.000903912 | 6.32E-14 | Eastern |
| UG_41 | 0.010977112 | 0.533984344 | 0.453554963 | 0.001483581 | 5.85E-12 | Eastern |
| UG_49 | 0.014545429 | 0.664302258 | 0.320364013 | 0.0007883 | 1.16E-10 | Eastern |
| UG_57 | 0.003139695 | 0.612693316 | 0.383947928 | 0.00021906 | 1.04E-15 | Eastern |
| UG_65 | 0.072951552 | 0.710358134 | 0.214414222 | 0.002218398 | 5.77E-05 | Eastern |
| UG_02 | 0.005591117 | 0.112594998 | 0.852273194 | 0.029540692 | 9.34E-14 | Eastern |
| UG_74 | 0.003312135 | 0.627804025 | 0.368675209 | 0.000208631 | 1.72E-15 | Eastern |
| UG_82 | 0.014234296 | 0.686037522 | 0.29907784 | 0.000650343 | 1.22E-10 | Eastern |
| UG_90 | 0.07367385 | 0.700951918 | 0.222874695 | 0.002445138 | 5.44E-05 | Eastern |
| UG_10 | 0.000648606 | 0.002786921 | 0.432877625 | 0.563686848 | 6.90E-16 | Eastern |
| UG_18 | 0.095681208 | 0.703895459 | 0.197173865 | 0.002664408 | 0.000585061 | Eastern |
| UG_26 | 0.002658601 | 0.624041783 | 0.373131519 | 0.000168097 | 3.40E-16 | Eastern |
| UG_34 | 0.002594329 | 0.622051419 | 0.375188304 | 0.000165948 | 2.80E-16 | Eastern |
| UG_42 | 0.011548566 | 0.548560449 | 0.43847309 | 0.001417894 | 9.14E-12 | Eastern |
| UG_50 | 0.002945776 | 0.634260229 | 0.362619047 | 0.000174948 | 7.73E-16 | Eastern |
| UG_58 | 0.005908358 | 0.474671501 | 0.518288326 | 0.001131814 | 5.01E-14 | Eastern |
| UG_66 | 0.074842811 | 0.713930577 | 0.208980022 | 0.002170353 | 7.62E-05 | Eastern |
| UG_03 | 0.051753396 | 0.337119404 | 0.582836821 | 0.028289981 | 3.97E-07 | Eastern |
| UG_75 | 0.003397778 | 0.612929238 | 0.383434347 | 0.000238637 | 1.85E-15 | Eastern |

| | | | | | | |
|---|---|---|---|---|---|---|
| UG_83 | 0.002849135 | 0.630339987 | 0.366637423 | 0.000173454 | 5.88E-16 | Eastern |
| UG_91 | 0.091497343 | 0.704994839 | 0.200515855 | 0.002597508 | 0.000394454 | Eastern |
| UG_11 | 0.002398195 | 0.636652874 | 0.360811994 | 0.000136937 | 1.78E-16 | Eastern |
| UG_19 | 0.006423327 | 0.492489024 | 0.499988555 | 0.001099094 | 9.80E-14 | Eastern |
| UG_27 | 0.002566119 | 0.621021353 | 0.376247368 | 0.00016516 | 2.57E-16 | Eastern |
| UG_35 | 0.006238524 | 0.52000406 | 0.472873991 | 0.000883425 | 8.88E-14 | Eastern |
| UG_43 | 0.003231003 | 0.672479912 | 0.324143152 | 0.000145932 | 2.10E-15 | Eastern |
| UG_51 | 0.002593508 | 0.621507719 | 0.375732237 | 0.000166535 | 2.79E-16 | Eastern |
| UG_59 | 0.035487777 | 0.300799576 | 0.637526011 | 0.026186613 | 2.31E-08 | Eastern |
| UG_67 | 0.009541774 | 0.192346299 | 0.779865381 | 0.018246546 | 2.03E-12 | Eastern |
| UG_04 | 0.039622182 | 0.402419703 | 0.54418942 | 0.013768642 | 5.27E-08 | Eastern |
| UG_76 | 0.00252989 | 0.601432798 | 0.395850581 | 0.000186732 | 2.02E-16 | Eastern |
| UG_84 | 0.005388372 | 0.519108322 | 0.474746409 | 0.000756897 | 3.05E-14 | Eastern |
| UG_92 | 0.080552485 | 0.698209699 | 0.218490494 | 0.002634124 | 0.000113198 | Eastern |
| UG_12 | 0.002606852 | 0.622981836 | 0.374245582 | 0.00016573 | 2.92E-16 | Eastern |
| UG_20 | 0.041618483 | 0.662753233 | 0.29340215 | 0.002225809 | 3.25E-07 | Eastern |
| UG_28 | 0.003382058 | 0.525873614 | 0.470311711 | 0.000432617 | 1.08E-15 | Eastern |
| UG_36 | 0.021909577 | 0.689036903 | 0.288069453 | 0.000984063 | 3.16E-09 | Eastern |
| UG_44 | 0.003213943 | 0.640732438 | 0.355869845 | 0.000183775 | 1.53E-15 | Eastern |
| UG_52 | 0.005317622 | 0.642982854 | 0.351385112 | 0.000314411 | 6.02E-14 | Eastern |
| UG_60 | 0.019561669 | 0.492415122 | 0.484390399 | 0.003632809 | 3.41E-10 | Eastern |
| UG_68 | 0.002784976 | 0.623081565 | 0.373955275 | 0.000178184 | 4.72E-16 | Eastern |
| UG_05 | 0.047424941 | 0.669478546 | 0.280739701 | 0.002355835 | 9.76E-07 | Eastern |
| UG_77 | 0.005402967 | 0.153394292 | 0.825479481 | 0.01572326 | 4.32E-14 | Eastern |
| UG_85 | 0.025460958 | 0.057833686 | 0.01157576 | 0.000472165 | 0.90465743 | Eastern |
| UG_93 | 0.008885728 | 0.631426458 | 0.359090445 | 0.00059737 | 2.32E-12 | Eastern |
| UG_13 | 0.00311706 | 0.612253454 | 0.384411493 | 0.000217994 | 9.86E-16 | Eastern |
| UG_21 | 0.006236289 | 0.501383906 | 0.491377957 | 0.001001848 | 8.18E-14 | Eastern |
| UG_29 | 0.086340235 | 0.661296336 | 0.248382462 | 0.003856922 | 0.000124045 | Eastern |

| | | | | | | |
|---|---|---|---|---|---|---|
| UG_37 | 0.089064513 | 0.660932524 | 0.245908458 | 0.003932667 | 0.000161838 | Eastern |
| UG_45 | 0.003430878 | 0.623223991 | 0.373120982 | 0.000224149 | 2.14E-15 | Eastern |
| UG_53 | 0.006482831 | 0.54598898 | 0.446756078 | 0.000772111 | 1.33E-13 | Eastern |
| UG_61 | 0.005638359 | 0.510401039 | 0.483116723 | 0.000843878 | 4.08E-14 | Eastern |
| UG_69 | 0.084031417 | 0.67197402 | 0.24043123 | 0.00345069 | 0.000112643 | Eastern |
| UG_06 | 0.003799397 | 0.620152714 | 0.375791478 | 0.000256411 | 4.38E-15 | Eastern |
| UG_78 | 0.003221974 | 0.642911767 | 0.353684888 | 0.000181372 | 1.59E-15 | Eastern |
| UG_86 | 0.002510585 | 0.578198104 | 0.419073693 | 0.000217617 | 1.64E-16 | Eastern |
| UG_94 | 0.005017007 | 0.509601796 | 0.484634897 | 0.0007463 | 1.74E-14 | Eastern |
| UG_14 | 0.087874619 | 0.668105879 | 0.24020252 | 0.003658972 | 0.00015801 | Eastern |
| UG_22 | 0.002562207 | 0.051259149 | 0.893509529 | 0.052669115 | 1.48E-15 | Eastern |
| UG_30 | 0.005422473 | 0.478653707 | 0.514921283 | 0.001002537 | 2.72E-14 | Eastern |
| UG_38 | 0.049248801 | 0.368053431 | 0.561117833 | 0.021579666 | 2.69E-07 | Eastern |
| UG_46 | 0.005509983 | 0.521305903 | 0.472419908 | 0.000764206 | 3.63E-14 | Eastern |
| UG_54 | 0.060114536 | 0.683206188 | 0.254149835 | 0.002521685 | 7.76E-06 | Eastern |
| UG_62 | 0.017401342 | 0.706892892 | 0.275031474 | 0.000674291 | 6.88E-10 | Eastern |
| UG_70 | 0.003715825 | 0.131400837 | 0.850653621 | 0.014229717 | 3.57E-15 | Eastern |
| UG_07 | 0.00258124 | 0.626770469 | 0.370488752 | 0.000159539 | 2.80E-16 | Eastern |
| UG_79 | 0.090000407 | 0.334359992 | 0.529934898 | 0.045672523 | 3.22E-05 | Eastern |
| UG_87 | 0.009283437 | 0.18345063 | 0.78772882 | 0.019537113 | 1.77E-12 | Eastern |
| UG_15 | 0.002666037 | 0.621810422 | 0.375352212 | 0.000171329 | 3.41E-16 | Eastern |
| UG_23 | 0.083770054 | 0.679439518 | 0.233454025 | 0.003215059 | 0.000121344 | Eastern |
| UG_31 | 0.018692381 | 0.655451636 | 0.324763335 | 0.001092648 | 6.86E-10 | Eastern |
| UG_39 | 0.002671358 | 0.60048273 | 0.39664625 | 0.000199662 | 2.97E-16 | Eastern |
| UG_47 | 0.002839906 | 0.604709407 | 0.392243276 | 0.000207411 | 4.76E-16 | Eastern |
| UG_55 | 0.009340136 | 0.193077031 | 0.779884397 | 0.017698437 | 1.73E-12 | Eastern |
| UG_63 | 0.005891931 | 0.637135655 | 0.356605366 | 0.000367049 | 1.21E-13 | Eastern |
| UG_71 | 0.006159754 | 0.589204772 | 0.404093392 | 0.000542082 | 1.18E-13 | Eastern |
| UG_08 | 0.006249399 | 0.517085568 | 0.475762231 | 0.000902802 | 8.88E-14 | Eastern |

| | | | | | | |
|---|---|---|---|---|---|---|
| UG_80 | 0.061733207 | 0.526019815 | 0.403537627 | 0.008706877 | 2.47E-06 | Eastern |
| UG_88 | 0.006084397 | 0.502514922 | 0.490432979 | 0.000967702 | 6.87E-14 | Eastern |
| UG_16 | 0.093597108 | 0.706160006 | 0.197162242 | 0.002584718 | 0.000495925 | Eastern |
| UG_24 | 0.00586324 | 0.490971667 | 0.502160296 | 0.001004797 | 5.02E-14 | Eastern |
| UG_32 | 0.002605364 | 0.622729196 | 0.374499516 | 0.000165924 | 2.91E-16 | Eastern |
| UG_40 | 0.010843703 | 0.528269687 | 0.459364215 | 0.001522395 | 5.20E-12 | Eastern |
| UG_48 | 0.066314242 | 0.728839786 | 0.203066942 | 0.001744725 | 3.43E-05 | Eastern |
| UG_56 | 0.00318982 | 0.624807372 | 0.371798316 | 0.000204492 | 1.28E-15 | Eastern |
| UG_64 | 0.016677225 | 0.676996248 | 0.305503803 | 0.000822723 | 3.62E-10 | Eastern |
| UG_72 | 0.00272029 | 0.611059824 | 0.386030785 | 0.000189101 | 3.65E-16 | Eastern |
| UG_95 | 0.003018229 | 0.617111135 | 0.3796674 | 0.000203236 | 8.09E-16 | Eastern |
| UG_167 | 0.005779915 | 0.474220428 | 0.518891423 | 0.001108234 | 4.26E-14 | Eastern |
| UG_175 | 0.004592609 | 0.56401564 | 0.430924112 | 0.000467639 | 1.20E-14 | Eastern |
| UG_183 | 0.005599818 | 0.531351824 | 0.462321726 | 0.000726632 | 4.27E-14 | Eastern |
| UG_103 | 0.004650304 | 0.59895137 | 0.396026214 | 0.000372112 | 1.63E-14 | Eastern |
| UG_111 | 0.005122022 | 0.511970756 | 0.482155874 | 0.000751348 | 2.05E-14 | Eastern |
| UG_119 | 0.082795087 | 0.665285035 | 0.248192018 | 0.003637043 | 9.08E-05 | Eastern |
| UG_127 | 0.00644938 | 0.468881839 | 0.52337282 | 0.001295961 | 9.29E-14 | Eastern |
| UG_135 | 0.093348688 | 0.367270543 | 0.502325504 | 0.037012575 | 4.27E-05 | Eastern |
| UG_143 | 0.004164397 | 0.138397619 | 0.842891274 | 0.014546709 | 7.55E-15 | Eastern |
| UG_151 | 0.003503552 | 0.589840551 | 0.4063656 | 0.000290297 | 1.97E-15 | Eastern |
| UG_159 | 0.006328648 | 0.609935054 | 0.383253743 | 0.000482555 | 1.66E-13 | Eastern |
| UG_96 | 0.003086059 | 0.613227231 | 0.383472595 | 0.000214116 | 9.23E-16 | Eastern |
| UG_168 | 0.004899826 | 0.578467637 | 0.416177987 | 0.00045455 | 2.09E-14 | Eastern |
| UG_176 | 0.022527585 | 0.671162841 | 0.305142235 | 0.001167336 | 3.23E-09 | Eastern |
| UG_184 | 0.088400804 | 0.691354526 | 0.217070137 | 0.002942526 | 0.000232008 | Eastern |
| UG_104 | 0.004096469 | 0.574319776 | 0.421199653 | 0.000384103 | 5.57E-15 | Eastern |
| UG_112 | 0.014155633 | 0.594822931 | 0.389745295 | 0.001276141 | 5.40E-11 | Eastern |
| UG_120 | 0.021639135 | 0.58178939 | 0.394396493 | 0.00217498 | 1.15E-09 | Eastern |

| | | | | | | |
|---|---|---|---|---|---|---|
| UG_128 | 0.008622575 | 0.324411537 | 0.661961158 | 0.005004731 | 6.30E-13 | Eastern |
| UG_136 | 0.006318798 | 0.517021947 | 0.475745078 | 0.000914177 | 9.62E-14 | Eastern |
| UG_144 | 0.007692114 | 0.494357632 | 0.496628822 | 0.001321432 | 3.66E-13 | Eastern |
| UG_152 | 0.002600657 | 0.588175953 | 0.409012202 | 0.000211188 | 2.26E-16 | Eastern |
| UG_160 | 0.004755289 | 0.135427271 | 0.84230857 | 0.017508869 | 2.06E-14 | Eastern |
| UG_97 | 0.129612419 | 0.591024673 | 0.268820262 | 0.008412626 | 0.002130021 | Eastern |
| UG_169 | 0.009491888 | 0.522708292 | 0.466430175 | 0.001369645 | 1.91E-12 | Eastern |
| UG_177 | 0.073052695 | 0.615844645 | 0.305996925 | 0.005087283 | 1.85E-05 | Eastern |
| UG_185 | 0.003195657 | 0.604346166 | 0.392221239 | 0.000236938 | 1.12E-15 | Eastern |
| UG_105 | 0.002472081 | 0.645052794 | 0.352341915 | 0.000133209 | 2.37E-16 | Eastern |
| UG_113 | 0.022182497 | 0.566615105 | 0.40872047 | 0.002481926 | 1.26E-09 | Eastern |
| UG_121 | 0.00409108 | 0.574602969 | 0.420923153 | 0.000382797 | 5.53E-15 | Eastern |
| UG_129 | 0.003995279 | 0.566793761 | 0.42881744 | 0.00039352 | 4.45E-15 | Eastern |
| UG_137 | 0.005435716 | 0.475387824 | 0.518148683 | 0.001027777 | 2.74E-14 | Eastern |
| UG_145 | 0.039955747 | 0.609816967 | 0.346995436 | 0.003231704 | 1.46E-07 | Eastern |
| UG_153 | 0.019163172 | 0.684011751 | 0.295928717 | 0.000896358 | 1.10E-09 | Eastern |
| UG_161 | 0.003438916 | 0.605875353 | 0.390431542 | 0.000254189 | 1.92E-15 | Eastern |
| UG_98 | 0.07087405 | 0.689334826 | 0.237092 | 0.002666009 | 3.31E-05 | Eastern |
| UG_170 | 0.006710746 | 0.529643068 | 0.46274997 | 0.000896215 | 1.58E-13 | Eastern |
| UG_178 | 0.006422653 | 0.605110089 | 0.387959834 | 0.000507424 | 1.78E-13 | Eastern |
| UG_186 | 0.005543515 | 0.502926853 | 0.490658315 | 0.000871316 | 3.50E-14 | Eastern |
| UG_106 | 0.021637939 | 0.698231912 | 0.27922886 | 0.000901286 | 3.19E-09 | Eastern |
| UG_114 | 0.00301083 | 0.58675311 | 0.409985219 | 0.000250841 | 6.46E-16 | Eastern |
| UG_122 | 0.004541211 | 0.620259898 | 0.374887129 | 0.000311763 | 1.60E-14 | Eastern |
| UG_130 | 0.034245823 | 0.461989673 | 0.495866987 | 0.007897497 | 2.01E-08 | Eastern |
| UG_138 | 0.058220809 | 0.683466032 | 0.255849353 | 0.002457817 | 5.99E-06 | Eastern |
| UG_146 | 0.005455028 | 0.508336579 | 0.485383132 | 0.000825261 | 3.19E-14 | Eastern |
| UG_154 | 0.005783077 | 0.506776222 | 0.48655146 | 0.000889241 | 4.84E-14 | Eastern |
| UG_162 | 0.003332412 | 0.629028813 | 0.367430577 | 0.000208198 | 1.81E-15 | Eastern |

| | | | | | | |
|---|---|---|---|---|---|---|
| UG_99 | 0.004593635 | 0.562772578 | 0.432162027 | 0.00047176 | 1.20E-14 | Eastern |
| UG_171 | 0.008713212 | 0.653086764 | 0.337701482 | 0.000498541 | 2.41E-12 | Eastern |
| UG_179 | 0.004927163 | 0.46631881 | 0.527772815 | 0.000981212 | 1.30E-14 | Eastern |
| UG_187 | 0.014786892 | 0.680008661 | 0.304494788 | 0.000709658 | 1.53E-10 | Eastern |
| UG_107 | 0.005830635 | 0.532809579 | 0.460607729 | 0.000752056 | 5.77E-14 | Eastern |
| UG_115 | 0.003598822 | 0.609480368 | 0.386660259 | 0.000260551 | 2.74E-15 | Eastern |
| UG_123 | 0.037044617 | 0.692719514 | 0.268668701 | 0.001566985 | 1.83E-07 | Eastern |
| UG_131 | 0.028683215 | 0.657263629 | 0.312404346 | 0.001648793 | 1.75E-08 | Eastern |
| UG_139 | 0.003231166 | 0.557094597 | 0.439341585 | 0.000332652 | 9.04E-16 | Eastern |
| UG_147 | 0.008583979 | 0.514600562 | 0.475517609 | 0.001297849 | 8.86E-13 | Eastern |
| UG_155 | 0.005709154 | 0.485999396 | 0.507282133 | 0.001009317 | 4.06E-14 | Eastern |
| UG_163 | 0.010755619 | 0.644257693 | 0.344319655 | 0.000667033 | 1.04E-11 | Eastern |
| UG_100 | 0.00597974 | 0.545729174 | 0.447583026 | 0.000708059 | 7.40E-14 | Eastern |
| UG_172 | 0.005739298 | 0.515459214 | 0.477970006 | 0.000831482 | 4.75E-14 | Eastern |
| UG_180 | 0.006310964 | 0.489195024 | 0.503391607 | 0.001102406 | 8.51E-14 | Eastern |
| UG_188 | 0.004361883 | 0.540105917 | 0.455012001 | 0.000520199 | 7.28E-15 | Eastern |
| UG_108 | 0.005956118 | 0.526373962 | 0.466865717 | 0.000804203 | 6.53E-14 | Eastern |
| UG_116 | 0.006055958 | 0.492667328 | 0.500247538 | 0.001029176 | 6.39E-14 | Eastern |
| UG_124 | 0.018139763 | 0.585152269 | 0.394936729 | 0.001771239 | 3.16E-10 | Eastern |
| UG_132 | 0.006169054 | 0.519615241 | 0.473340742 | 0.000874963 | 8.18E-14 | Eastern |
| UG_140 | 0.003997636 | 0.112237735 | 0.862943483 | 0.020821147 | 7.97E-15 | Eastern |
| UG_148 | 0.005923742 | 0.529300525 | 0.463992045 | 0.000783688 | 6.37E-14 | Eastern |
| UG_156 | 0.00637457 | 0.542152736 | 0.450694574 | 0.00077812 | 1.16E-13 | Eastern |
| UG_164 | 0.010188012 | 0.443728194 | 0.543551568 | 0.002532225 | 2.41E-12 | Eastern |
| UG_101 | 0.016010909 | 0.563123599 | 0.419051671 | 0.001813821 | 1.10E-10 | Eastern |
| UG_173 | 0.003547375 | 0.130620148 | 0.852142697 | 0.01368978 | 2.57E-15 | Eastern |
| UG_181 | 0.005867922 | 0.519182034 | 0.474119332 | 0.000830712 | 5.67E-14 | Eastern |
| UG_109 | 0.0861689 | 0.666602777 | 0.243425304 | 0.003672474 | 0.000130546 | Eastern |
| UG_117 | 0.00544158 | 0.642742548 | 0.351492872 | 0.000323001 | 7.11E-14 | Eastern |

| | | | | | | |
|---|---|---|---|---|---|---|
| UG_125 | 0.005614175 | 0.584422456 | 0.40945699 | 0.000506379 | 5.84E-14 | Eastern |
| UG_133 | 0.004857903 | 0.094381657 | 0.865532746 | 0.035227693 | 4.63E-14 | Eastern |
| UG_141 | 0.007121205 | 0.522486611 | 0.469388328 | 0.001003856 | 2.35E-13 | Eastern |
| UG_149 | 0.003908128 | 0.581780201 | 0.413965317 | 0.000346354 | 4.14E-15 | Eastern |
| UG_157 | 0.003310458 | 0.553057966 | 0.443280334 | 0.000351242 | 1.05E-15 | Eastern |
| UG_165 | 0.002825792 | 0.60822827 | 0.388744718 | 0.00020122 | 4.71E-16 | Eastern |
| UG_102 | 0.007805319 | 0.547860438 | 0.443400935 | 0.000933307 | 5.20E-13 | Eastern |
| UG_174 | 0.010035564 | 0.664784255 | 0.324649212 | 0.000530968 | 7.54E-12 | Eastern |
| UG_182 | 0.098592467 | 0.553786843 | 0.337439632 | 0.010046535 | 0.000134524 | Eastern |
| UG_110 | 0.003109117 | 0.579466914 | 0.417150601 | 0.000273368 | 7.79E-16 | Eastern |
| UG_118 | 0.002912715 | 0.603011755 | 0.393859666 | 0.000215864 | 5.66E-16 | Eastern |
| UG_126 | 0.087913524 | 0.680666822 | 0.227976343 | 0.003254691 | 0.00018862 | Eastern |
| UG_134 | 0.079651419 | 0.645618466 | 0.270457418 | 0.004221145 | 5.16E-05 | Eastern |
| UG_142 | 0.004867136 | 0.131687123 | 0.844494094 | 0.018951647 | 2.56E-14 | Eastern |
| UG_150 | 0.013093018 | 0.576513758 | 0.409057034 | 0.00133619 | 2.70E-11 | Eastern |
| UG_158 | 0.129868974 | 0.605138899 | 0.255017865 | 0.007425444 | 0.002548817 | Eastern |
| UG_166 | 0.005515182 | 0.430706738 | 0.562359868 | 0.001418212 | 2.69E-14 | Eastern |
| UG_189 | 0.004378625 | 0.115181924 | 0.858585044 | 0.021854408 | 1.48E-14 | Eastern |
| UG_261 | 0.005086372 | 0.156705468 | 0.824089816 | 0.014118344 | 2.69E-14 | Northern |
| UG_269 | 0.004085728 | 0.118691594 | 0.858051168 | 0.019171511 | 8.48E-15 | Northern |
| UG_277 | 0.003499429 | 0.112882663 | 0.865763624 | 0.017854283 | 2.98E-15 | Northern |
| UG_197 | 0.009713431 | 0.182631068 | 0.786973339 | 0.020682162 | 2.48E-12 | Eastern |
| UG_205 | 0.004620594 | 0.161197171 | 0.822160113 | 0.012022122 | 1.28E-14 | Eastern |
| UG_213 | 0.003284328 | 0.079947381 | 0.885259443 | 0.031508847 | 3.58E-15 | Eastern |
| UG_221 | 0.004783221 | 0.137882168 | 0.840320756 | 0.017013856 | 2.09E-14 | Eastern |
| UG_229 | 0.009505832 | 0.173865493 | 0.79429485 | 0.022333826 | 2.27E-12 | Northern |
| UG_237 | 0.004424178 | 0.14673975 | 0.834996582 | 0.013839491 | 1.07E-14 | Northern |
| UG_245 | 0.006740689 | 0.100811837 | 0.848578718 | 0.043868756 | 4.63E-13 | Northern |
| UG_253 | 0.006521893 | 0.098734932 | 0.850719252 | 0.044023923 | 3.77E-13 | Northern |

| | | | | | | |
|---|---|---|---|---|---|---|
| UG_190 | 0.006734862 | 0.468050398 | 0.52384814 | 0.0013666 | 1.27E-13 | Eastern |
| UG_262 | 0.008178127 | 0.143313376 | 0.8206148 | 0.027893696 | 1.01E-12 | Northern |
| UG_270 | 0.005624161 | 0.173675527 | 0.80791778 | 0.012782533 | 4.84E-14 | Northern |
| UG_278 | 0.004515196 | 0.11918092 | 0.855133637 | 0.021170248 | 1.75E-14 | Northern |
| UG_198 | 0.004253702 | 0.56205658 | 0.433254117 | 0.0004356 | 6.82E-15 | Eastern |
| UG_206 | 0.003359885 | 0.626620501 | 0.369805849 | 0.000213765 | 1.89E-15 | Eastern |
| UG_214 | 0.004876369 | 0.117697572 | 0.853895948 | 0.023530111 | 3.15E-14 | Eastern |
| UG_222 | 0.003887504 | 0.095556887 | 0.873300093 | 0.027255516 | 8.73E-15 | Eastern |
| UG_230 | 0.009936316 | 0.182434995 | 0.786399813 | 0.021228876 | 2.94E-12 | Northern |
| UG_238 | 0.00495244 | 0.138491543 | 0.839045945 | 0.017510072 | 2.68E-14 | Northern |
| UG_246 | 0.005305694 | 0.148479207 | 0.829768135 | 0.016446964 | 3.97E-14 | Northern |
| UG_254 | 0.009420053 | 0.183732842 | 0.787068112 | 0.019778993 | 1.96E-12 | Northern |
| UG_191 | 0.019361586 | 0.641596395 | 0.33778429 | 0.001257728 | 7.88E-10 | Eastern |
| UG_263 | 0.003550777 | 0.118316886 | 0.86154813 | 0.016584207 | 3.05E-15 | Northern |
| UG_271 | 0.00509579 | 0.130529685 | 0.844127445 | 0.02024708 | 3.64E-14 | Northern |
| UG_279 | 0.00575521 | 0.101761226 | 0.855823141 | 0.036660423 | 1.40E-13 | Northern |
| UG_199 | 0.004887322 | 0.138160805 | 0.839608239 | 0.017343635 | 2.44E-14 | Eastern |
| UG_207 | 0.010103652 | 0.684488029 | 0.304949187 | 0.000459132 | 9.61E-12 | Eastern |
| UG_215 | 0.003864351 | 0.095704019 | 0.87342423 | 0.0270074 | 8.33E-15 | Eastern |
| UG_223 | 0.015137323 | 0.572129164 | 0.411127397 | 0.001606116 | 7.64E-11 | Eastern |
| UG_231 | 0.009330826 | 0.181452278 | 0.789130379 | 0.020086518 | 1.86E-12 | Northern |
| UG_239 | 0.005269648 | 0.156496319 | 0.823526942 | 0.014707091 | 3.49E-14 | Northern |
| UG_247 | 0.008005086 | 0.139676585 | 0.823661204 | 0.028657125 | 9.03E-13 | Northern |
| UG_255 | 0.007830022 | 0.129665243 | 0.830252829 | 0.032251905 | 8.73E-13 | Northern |
| UG_192 | 0.062743825 | 0.524656132 | 0.403685024 | 0.008912226 | 2.79E-06 | Eastern |
| UG_264 | 0.006598519 | 0.101783248 | 0.84942728 | 0.042190952 | 3.87E-13 | Northern |
| UG_272 | 0.008006869 | 0.134585533 | 0.826641617 | 0.030765981 | 9.65E-13 | Northern |
| UG_280 | 0.003178547 | 0.604286606 | 0.392299212 | 0.000235635 | 1.07E-15 | Northern |
| UG_200 | 0.004201955 | 0.141103264 | 0.840553951 | 0.01414083 | 7.82E-15 | Eastern |

| | | | | | | |
|---|---|---|---|---|---|---|
| UG_208 | 0.003348649 | 0.128693947 | 0.854720887 | 0.013236516 | 1.73E-15 | Eastern |
| UG_216 | 0.003586001 | 0.133009322 | 0.850033927 | 0.01337075 | 2.70E-15 | Eastern |
| UG_224 | 0.000822261 | 0.004150697 | 0.512524968 | 0.482502074 | 8.21E-16 | Eastern |
| UG_232 | 0.005133298 | 0.151493441 | 0.828119847 | 0.015253414 | 3.02E-14 | Northern |
| UG_240 | 0.004276085 | 0.135887341 | 0.844322563 | 0.015514011 | 9.43E-15 | Northern |
| UG_248 | 0.003016796 | 0.582332771 | 0.41439122 | 0.000259212 | 6.37E-16 | Northern |
| UG_256 | 0.003154715 | 0.573589005 | 0.422966977 | 0.000289303 | 8.35E-16 | Northern |
| UG_193 | 6.19E-13 | 1.93E-13 | 1.94E-15 | 1.12E-16 | 1 | Eastern |
| UG_265 | 0.004192344 | 0.129696311 | 0.849484779 | 0.016626565 | 8.81E-15 | Northern |
| UG_273 | 0.009743942 | 0.184020581 | 0.785803183 | 0.020432294 | 2.51E-12 | Northern |
| UG_281 | 0.004919028 | 0.158046517 | 0.823647826 | 0.013386629 | 2.08E-14 | Northern |
| UG_201 | 0.004955024 | 0.118253071 | 0.85307013 | 0.023721775 | 3.51E-14 | Eastern |
| UG_209 | 0.00497296 | 0.163646122 | 0.818752813 | 0.012628105 | 2.14E-14 | Eastern |
| UG_217 | 0.004341868 | 0.137596942 | 0.842669915 | 0.015391276 | 1.03E-14 | Eastern |
| UG_225 | 0.036986898 | 0.66534261 | 0.295708371 | 0.001961987 | 1.33E-07 | Eastern |
| UG_233 | 0.003579878 | 0.104391389 | 0.870832174 | 0.021196559 | 4.04E-15 | Northern |
| UG_241 | 0.003553794 | 0.090212666 | 0.878701018 | 0.027532522 | 5.04E-15 | Northern |
| UG_249 | 0.004899488 | 0.129640066 | 0.84578881 | 0.019671635 | 2.76E-14 | Northern |
| UG_257 | 0.007614443 | 0.136926138 | 0.827208291 | 0.028251129 | 6.46E-13 | Northern |
| UG_194 | 0.010217596 | 0.657132142 | 0.332076692 | 0.000573569 | 8.02E-12 | Eastern |
| UG_266 | 0.00513759 | 0.155639328 | 0.824755143 | 0.014467939 | 2.92E-14 | Northern |
| UG_274 | 0.00938234 | 0.184153736 | 0.786860553 | 0.019603371 | 1.90E-12 | Northern |
| UG_282 | 0.007362474 | 0.125975333 | 0.834723135 | 0.031939058 | 5.83E-13 | Northern |
| UG_202 | 0.005135893 | 0.12214183 | 0.849543139 | 0.023179138 | 4.32E-14 | Eastern |
| UG_210 | 0.004581069 | 0.175783279 | 0.809653325 | 0.009982327 | 1.07E-14 | Eastern |
| UG_218 | 0.089425572 | 0.687957808 | 0.219315072 | 0.003057283 | 0.000244265 | Eastern |
| UG_226 | 0.005675855 | 0.629279604 | 0.364671477 | 0.000373064 | 8.67E-14 | Eastern |
| UG_234 | 0.005277649 | 0.155609702 | 0.824213379 | 0.014899269 | 3.56E-14 | Northern |
| UG_242 | 0.007175821 | 0.126957587 | 0.835221311 | 0.030645281 | 4.75E-13 | Northern |

| | | | | | | |
|---|---|---|---|---|---|---|
| UG_250 | 0.002124737 | 0.036489074 | 0.885205476 | 0.076180713 | 8.19E-16 | Northern |
| UG_258 | 0.006797409 | 0.114263126 | 0.843694091 | 0.035245374 | 3.86E-13 | Northern |
| UG_195 | 0.003553742 | 0.106112784 | 0.869936604 | 0.02039687 | 3.72E-15 | Eastern |
| UG_267 | 0.005165175 | 0.164235318 | 0.817537763 | 0.013061744 | 2.81E-14 | Northern |
| UG_275 | 0.009388843 | 0.18518158 | 0.786034078 | 0.0193955 | 1.90E-12 | Northern |
| UG_203 | 0.002848082 | 0.603037133 | 0.393904253 | 0.000210532 | 4.81E-16 | Eastern |
| UG_211 | 0.003374036 | 0.128534049 | 0.85471414 | 0.013377775 | 1.83E-15 | Eastern |
| UG_219 | 0.004065636 | 0.117872478 | 0.858738614 | 0.019323273 | 8.27E-15 | Eastern |
| UG_227 | 0.006386765 | 0.484701791 | 0.507759961 | 0.001151483 | 9.13E-14 | Eastern |
| UG_235 | 0.007436372 | 0.130321046 | 0.831966282 | 0.0302763 | 5.91E-13 | Northern |
| UG_243 | 0.005500566 | 0.164775967 | 0.815835948 | 0.013887519 | 4.43E-14 | Northern |
| UG_251 | 0.009459705 | 0.184520025 | 0.786326192 | 0.019694077 | 2.01E-12 | Northern |
| UG_259 | 0.004709673 | 0.116298182 | 0.855798895 | 0.02319325 | 2.49E-14 | Northern |
| UG_196 | 0.003857221 | 0.076643218 | 0.87933041 | 0.04016915 | 1.28E-14 | Eastern |
| UG_268 | 9.71E-05 | 0.00013605 | 0.084182143 | 0.915584686 | 3.83E-16 | Northern |
| UG_276 | 0.005409225 | 0.134471085 | 0.83974315 | 0.020376541 | 5.37E-14 | Northern |
| UG_204 | 0.00359882 | 0.130819974 | 0.851717337 | 0.013863869 | 2.85E-15 | Eastern |
| UG_212 | 0.002925611 | 0.608413771 | 0.38845179 | 0.000208827 | 6.06E-16 | Eastern |
| UG_220 | 0.08416858 | 0.657522578 | 0.254280324 | 0.00393357 | 9.49E-05 | Eastern |
| UG_228 | 0.005862553 | 0.629307419 | 0.364443622 | 0.000386407 | 1.10E-13 | Eastern |
| UG_236 | 0.004372858 | 0.108865673 | 0.862500226 | 0.024261243 | 1.63E-14 | Northern |
| UG_244 | 0.00453998 | 0.147237752 | 0.834085792 | 0.014136477 | 1.29E-14 | Northern |
| UG_252 | 0.00413485 | 0.142644337 | 0.839618382 | 0.013602431 | 6.83E-15 | Northern |
| UG_260 | 0.004528876 | 0.140356153 | 0.839621446 | 0.015493525 | 1.36E-14 | Northern |
| UG_283 | 0.005878075 | 0.116208043 | 0.84855767 | 0.029356213 | 1.28E-13 | Northern |
| UG_355 | 0.000132116 | 0.000173615 | 0.088258643 | 0.911435626 | 1.73E-15 | Northern |
| UG_363 | 0.003952374 | 0.128129488 | 0.851944316 | 0.015973822 | 5.84E-15 | Northern |
| UG_371 | 0.004702297 | 0.042196851 | 0.824885542 | 0.12821531 | 2.46E-13 | Northern |
| UG_291 | 0.004257639 | 0.152100468 | 0.831280114 | 0.012361779 | 7.67E-15 | Northern |

| | | | | | | |
|---|---|---|---|---|---|---|
| UG_299 | 0.000234612 | 0.000398079 | 0.13978222 | 0.859585089 | 2.85E-15 | Northern |
| UG_307 | 0.003333062 | 0.069321607 | 0.886178131 | 0.0411672 | 5.36E-15 | Northern |
| UG_315 | 0.009442031 | 0.181930532 | 0.788396759 | 0.020230678 | 2.03E-12 | Northern |
| UG_323 | 0.003389039 | 0.113717975 | 0.865884696 | 0.01700829 | 2.32E-15 | Northern |
| UG_331 | 0.007628138 | 0.126799161 | 0.832837635 | 0.032735066 | 7.49E-13 | Northern |
| UG_339 | 0.000225863 | 0.000343116 | 0.123733741 | 0.87569728 | 4.89E-15 | Northern |
| UG_347 | 0.001394713 | 0.007054227 | 0.575176201 | 0.41637486 | 7.50E-15 | Northern |
| UG_284 | 0.000123766 | 0.000196442 | 0.104259939 | 0.895419853 | 4.21E-16 | North_western |
| UG_356 | 0.004916841 | 0.159816501 | 0.822182527 | 0.013084131 | 2.04E-14 | Northern |
| UG_364 | 0.004358721 | 0.123138125 | 0.853346929 | 0.019156225 | 1.28E-14 | Northern |
| UG_372 | 0.003321752 | 0.115977705 | 0.864668676 | 0.016031868 | 1.94E-15 | Northern |
| UG_292 | 0.002519206 | 0.022554188 | 0.795404776 | 0.179521829 | 1.20E-14 | Northern |
| UG_300 | 0.005175459 | 0.139285828 | 0.837385011 | 0.018153703 | 3.66E-14 | Northern |
| UG_308 | 0.002960094 | 0.030595997 | 0.830923467 | 0.135520442 | 1.68E-14 | Northern |
| UG_316 | 0.002025917 | 0.013537774 | 0.703459752 | 0.280976556 | 1.22E-14 | Northern |
| UG_324 | 0.004061324 | 0.143950593 | 0.838885489 | 0.013102595 | 5.91E-15 | Northern |
| UG_332 | 0.004823394 | 0.1423886 | 0.836667309 | 0.016120698 | 2.11E-14 | Northern |
| UG_340 | 0.002366638 | 0.019579902 | 0.773332367 | 0.204721093 | 1.16E-14 | Northern |
| UG_348 | 0.001888632 | 0.014949261 | 0.743926223 | 0.239235884 | 4.74E-15 | Northern |
| UG_285 | 2.15E-12 | 6.80E-13 | 9.07E-15 | 6.14E-16 | 1 | South_western |
| UG_357 | 0.005183513 | 0.147751382 | 0.830868528 | 0.016196578 | 3.38E-14 | Northern |
| UG_365 | 0.004635222 | 0.066086015 | 0.866822867 | 0.062455896 | 6.99E-14 | Northern |
| UG_373 | 0.00144222 | 0.010699169 | 0.700263746 | 0.287594865 | 1.77E-15 | Northern |
| UG_293 | 0.010301123 | 0.196127555 | 0.774568301 | 0.01900302 | 3.48E-12 | Northern |
| UG_301 | 0.020647006 | 0.247201009 | 0.708272016 | 0.023879969 | 4.60E-10 | Northern |
| UG_309 | 0.002816832 | 0.041791049 | 0.874583155 | 0.080808964 | 4.92E-15 | Northern |
| UG_317 | 0.005441748 | 0.153025139 | 0.825612396 | 0.015920716 | 4.57E-14 | Northern |
| UG_325 | 0.009430313 | 0.18494306 | 0.786089019 | 0.019537608 | 1.96E-12 | Northern |
| UG_333 | 0.00317642 | 0.123111136 | 0.860092261 | 0.013620183 | 1.27E-15 | Northern |

| | | | | | | |
|---|---|---|---|---|---|---|
| UG_341 | 0.003019808 | 0.636005224 | 0.360797413 | 0.000177555 | 9.38E-16 | Northern |
| UG_349 | 0.003470506 | 0.600241018 | 0.396021333 | 0.000267142 | 1.97E-15 | Northern |
| UG_286 | 0.003742083 | 0.100838931 | 0.871715962 | 0.023703025 | 5.96E-15 | Northern |
| UG_358 | 0.003152996 | 0.582404962 | 0.414169988 | 0.000272054 | 8.78E-16 | Northern |
| UG_366 | 0.00552859 | 0.162573048 | 0.817550293 | 0.014348069 | 4.68E-14 | Northern |
| UG_374 | 0.005354167 | 0.356640375 | 0.635676493 | 0.002328964 | 1.96E-14 | Northern |
| UG_294 | 0.00950511 | 0.188514809 | 0.783037156 | 0.018942924 | 2.03E-12 | Northern |
| UG_302 | 0.039588983 | 0.273786613 | 0.650568319 | 0.036056028 | 5.72E-08 | Northern |
| UG_310 | 8.92E-05 | 0.000100029 | 0.064991173 | 0.934819616 | 1.14E-15 | Northern |
| UG_318 | 0.003732207 | 0.121330722 | 0.858259447 | 0.016677623 | 4.21E-15 | Northern |
| UG_326 | 0.009738132 | 0.19219058 | 0.779395706 | 0.018675582 | 2.36E-12 | Northern |
| UG_334 | 0.00629777 | 0.152341827 | 0.82257622 | 0.018784182 | 1.34E-13 | Northern |
| UG_342 | 0.003876344 | 0.121737212 | 0.857124659 | 0.017261785 | 5.52E-15 | Northern |
| UG_350 | 0.001050687 | 0.006604461 | 0.613982055 | 0.378362797 | 8.58E-16 | Northern |
| UG_287 | 0.005947053 | 0.13962254 | 0.833470337 | 0.020960069 | 1.01E-13 | Northern |
| UG_359 | 0.001809171 | 0.013523205 | 0.724482159 | 0.260185465 | 4.79E-15 | Northern |
| UG_367 | 0.005828618 | 0.153793967 | 0.823408092 | 0.016969323 | 7.49E-14 | Northern |
| UG_375 | 0.003543957 | 0.092071222 | 0.877935952 | 0.026448869 | 4.75E-15 | Northern |
| UG_295 | 0.009294765 | 0.184174969 | 0.787124946 | 0.01940532 | 1.77E-12 | Northern |
| UG_303 | 0.006194797 | 0.160895197 | 0.816354048 | 0.016555959 | 1.09E-13 | Northern |
| UG_311 | 0.002168491 | 0.030542229 | 0.864879897 | 0.102409383 | 1.52E-15 | Northern |
| UG_319 | 0.004917507 | 0.156902417 | 0.824601555 | 0.01357852 | 2.10E-14 | Northern |
| UG_327 | 0.000332928 | 0.000941507 | 0.253577109 | 0.745148455 | 5.01E-16 | Northern |
| UG_335 | 0.000130656 | 0.000170331 | 0.087193832 | 0.91250518 | 1.75E-15 | Northern |
| UG_343 | 0.000129623 | 0.00016141 | 0.082986238 | 0.916722729 | 2.26E-15 | Northern |
| UG_351 | 0.003809065 | 0.122374659 | 0.857045854 | 0.016770422 | 4.82E-15 | Northern |
| UG_288 | 0.006100745 | 0.146823882 | 0.82755016 | 0.019525214 | 1.13E-13 | Northern |
| UG_360 | 0.005245854 | 0.156078325 | 0.823961877 | 0.014713944 | 3.39E-14 | Northern |
| UG_368 | 0.006271843 | 0.157240493 | 0.818923699 | 0.017563966 | 1.24E-13 | Northern |

| | | | | | | |
|---|---|---|---|---|---|---|
| UG_376 | 0.009232324 | 0.182197082 | 0.788873351 | 0.019697244 | 1.71E-12 | Northern |
| UG_296 | 0.001938816 | 0.02052246 | 0.813676322 | 0.163862402 | 2.03E-15 | Northern |
| UG_304 | 0.004025319 | 0.154470865 | 0.830227517 | 0.011276299 | 4.98E-15 | Northern |
| UG_312 | 0.00937988 | 0.184659438 | 0.786472407 | 0.019488275 | 1.89E-12 | Northern |
| UG_320 | 0.004495 | 0.128502341 | 0.848759056 | 0.018243604 | 1.49E-14 | Northern |
| UG_328 | 0.004484883 | 0.122575089 | 0.853015526 | 0.019924502 | 1.59E-14 | Northern |
| UG_336 | 0.002893922 | 0.599978492 | 0.396908653 | 0.000218933 | 5.29E-16 | Northern |
| UG_344 | 0.002498714 | 0.577207684 | 0.420075641 | 0.00021796 | 1.58E-16 | Northern |
| UG_352 | 0.004492184 | 0.127905058 | 0.849207137 | 0.018395621 | 1.49E-14 | Northern |
| UG_289 | 0.005562696 | 0.149075118 | 0.82819477 | 0.017167416 | 5.58E-14 | Northern |
| UG_361 | 0.003963884 | 0.069757994 | 0.877614255 | 0.048663867 | 1.92E-14 | Northern |
| UG_369 | 0.003707298 | 0.121485376 | 0.858290122 | 0.016517204 | 4.00E-15 | Northern |
| UG_297 | 0.006939491 | 0.101861421 | 0.846840364 | 0.044358724 | 5.63E-13 | Northern |
| UG_305 | 0.004446399 | 0.122501302 | 0.853288066 | 0.019764233 | 1.49E-14 | Northern |
| UG_313 | 0.003712133 | 0.142432935 | 0.841719095 | 0.012135838 | 3.12E-15 | Northern |
| UG_321 | 0.004092603 | 0.14159142 | 0.840666163 | 0.013649815 | 6.41E-15 | Northern |
| UG_329 | 0.000829106 | 0.00421628 | 0.515949021 | 0.479005593 | 8.21E-16 | Northern |
| UG_337 | 0.009184841 | 0.180361439 | 0.790455166 | 0.019998555 | 1.67E-12 | Northern |
| UG_345 | 0.030980134 | 0.434206104 | 0.526149509 | 0.008664244 | 8.74E-09 | Northern |
| UG_353 | 0.002482134 | 0.568412837 | 0.428875177 | 0.000229852 | 1.43E-16 | Northern |
| UG_290 | 0.004119898 | 0.103754311 | 0.867220592 | 0.0249052 | 1.15E-14 | Northern |
| UG_362 | 0.002476235 | 0.60054451 | 0.396795769 | 0.000183486 | 1.72E-16 | Northern |
| UG_370 | 0.004971482 | 0.16242012 | 0.819791108 | 0.01281729 | 2.16E-14 | Northern |
| UG_298 | 0.00410286 | 0.116289919 | 0.859587037 | 0.020020184 | 9.06E-15 | Northern |
| UG_306 | 0.004077537 | 0.108137952 | 0.864979418 | 0.022805093 | 9.85E-15 | Northern |
| UG_314 | 0.002092467 | 0.014454745 | 0.716040377 | 0.26741241 | 1.24E-14 | Northern |
| UG_322 | 0.00929029 | 0.189044228 | 0.783282138 | 0.018383344 | 1.71E-12 | Northern |
| UG_330 | 0.003906963 | 0.135337405 | 0.846572068 | 0.014183564 | 4.91E-15 | Northern |
| UG_338 | 0.001838992 | 0.0082395 | 0.56713069 | 0.422790819 | 4.19E-14 | Northern |

| | | | | | | |
|---|---|---|---|---|---|---|
| UG_346 | 0.005157511 | 0.146693478 | 0.831809786 | 0.016339225 | 3.29E-14 | Northern |
| UG_354 | 0.004127001 | 0.113635057 | 0.861192159 | 0.021045783 | 9.85E-15 | Northern |
| UG_377 | 0.009332472 | 0.181895343 | 0.788781526 | 0.019990659 | 1.86E-12 | Northern |
| UG_449 | 2.86E-12 | 9.86E-13 | 1.24E-14 | 7.29E-16 | 1 | South_western |
| UG_457 | 3.95E-07 | 3.43E-07 | 1.92E-08 | 1.03E-09 | 0.999999243 | South_western |
| UG_465 | 1.99E-12 | 6.41E-13 | 8.20E-15 | 5.31E-16 | 1 | South_western |
| UG_385 | 0.002475714 | 0.577541096 | 0.419767949 | 0.000215241 | 1.48E-16 | Northern |
| UG_393 | 1.28E-12 | 4.00E-13 | 4.82E-15 | 3.11E-16 | 1 | South_western |
| UG_401 | 1.59E-12 | 5.06E-13 | 6.19E-15 | 3.93E-16 | 1 | South_western |
| UG_409 | 4.32E-13 | 1.34E-13 | 1.25E-15 | 6.90E-17 | 1 | South_western |
| UG_417 | 2.45E-12 | 8.46E-13 | 1.02E-14 | 5.91E-16 | 1 | South_western |
| UG_425 | 4.47E-10 | 2.06E-10 | 5.73E-12 | 4.03E-13 | 0.999999999 | South_western |
| UG_433 | 1.52E-12 | 4.57E-13 | 6.08E-15 | 4.36E-16 | 1 | South_western |
| UG_441 | 7.53E-12 | 2.56E-12 | 4.17E-14 | 2.97E-15 | 1 | South_western |
| UG_378 | 0.004025187 | 0.149837073 | 0.834153827 | 0.011983914 | 5.21E-15 | Northern |
| UG_450 | 1.36E-12 | 4.46E-13 | 5.02E-15 | 2.93E-16 | 1 | South_western |
| UG_458 | 0.050874027 | 0.121096106 | 0.027565372 | 0.001156691 | 0.799307803 | South_western |
| UG_466 | 3.03E-12 | 1.04E-12 | 1.34E-14 | 8.10E-16 | 1 | South_western |
| UG_386 | 0.005219283 | 0.167211882 | 0.814831059 | 0.012737776 | 2.95E-14 | Northern |
| UG_394 | 2.75E-12 | 9.44E-13 | 1.18E-14 | 6.99E-16 | 1 | South_western |
| UG_402 | 6.89E-12 | 2.58E-12 | 3.55E-14 | 2.03E-15 | 1 | South_western |
| UG_410 | 4.39E-12 | 1.55E-12 | 2.09E-14 | 1.25E-15 | 1 | South_western |
| UG_418 | 6.04E-12 | 2.19E-12 | 3.07E-14 | 1.84E-15 | 1 | South_western |
| UG_426 | 3.73E-12 | 1.26E-12 | 1.74E-14 | 1.13E-15 | 1 | South_western |
| UG_434 | 1.02E-12 | 2.96E-13 | 3.75E-15 | 2.72E-16 | 1 | South_western |
| UG_442 | 4.36E-12 | 1.53E-12 | 2.07E-14 | 1.25E-15 | 1 | South_western |
| UG_379 | 0.002674375 | 0.040237373 | 0.875485579 | 0.081602673 | 3.66E-15 | Northern |
| UG_451 | 0.043946227 | 0.105903068 | 0.022639966 | 0.000900783 | 0.826609956 | South_western |
| UG_459 | 2.35E-12 | 8.07E-13 | 9.70E-15 | 5.60E-16 | 1 | South_western |

| | | | | | | |
|---|---|---|---|---|---|---|
| UG_467 | 3.40E-12 | 1.08E-12 | 1.60E-14 | 1.15E-15 | 1 | South_western |
| UG_387 | 0.004227868 | 0.065429997 | 0.872405341 | 0.057936795 | 3.59E-14 | Northern |
| UG_395 | 1.97E-12 | 6.78E-13 | 7.79E-15 | 4.37E-16 | 1 | South_western |
| UG_403 | 4.99E-12 | 1.75E-12 | 2.46E-14 | 1.53E-15 | 1 | South_western |
| UG_411 | 2.75E-12 | 9.52E-13 | 1.18E-14 | 6.88E-16 | 1 | South_western |
| UG_419 | 1.35E-12 | 4.29E-13 | 5.06E-15 | 3.15E-16 | 1 | South_western |
| UG_427 | 1.45E-12 | 4.58E-13 | 5.54E-15 | 3.53E-16 | 1 | South_western |
| UG_435 | 1.68E-11 | 6.07E-12 | 1.09E-13 | 7.68E-15 | 1 | South_western |
| UG_443 | 0.006250318 | 0.01322646 | 0.002050741 | 7.71E-05 | 0.97839534 | South_western |
| UG_380 | 0.002511703 | 0.587580751 | 0.40970351 | 0.000204037 | 1.75E-16 | Northern |
| UG_452 | 0.042000582 | 0.100926335 | 0.021378848 | 0.000848421 | 0.834845814 | South_western |
| UG_460 | 4.68E-06 | 5.04E-06 | 3.72E-07 | 1.87E-08 | 0.99998989 | South_western |
| UG_468 | 1.74E-12 | 5.89E-13 | 6.73E-15 | 3.83E-16 | 1 | South_western |
| UG_388 | 0.002830442 | 0.081405819 | 0.889705997 | 0.026057742 | 1.15E-15 | Northern |
| UG_396 | 2.09E-12 | 6.53E-13 | 8.84E-15 | 6.13E-16 | 1 | South_western |
| UG_404 | 6.68E-10 | 3.34E-10 | 9.06E-12 | 5.74E-13 | 0.999999999 | South_western |
| UG_412 | 1.36E-12 | 4.36E-13 | 5.12E-15 | 3.16E-16 | 1 | South_western |
| UG_420 | 1.88E-12 | 5.88E-13 | 7.74E-15 | 5.27E-16 | 1 | South_western |
| UG_428 | 2.70E-12 | 9.00E-13 | 1.18E-14 | 7.42E-16 | 1 | South_western |
| UG_436 | 2.75E-12 | 8.69E-13 | 1.23E-14 | 8.65E-16 | 1 | South_western |
| UG_444 | 1.39E-09 | 7.62E-10 | 2.14E-11 | 1.25E-12 | 0.999999998 | South_western |
| UG_381 | 0.003761615 | 0.546541903 | 0.449273601 | 0.00042288 | 2.57E-15 | Northern |
| UG_453 | 0.037022154 | 0.086949047 | 0.018372607 | 0.000746895 | 0.856909297 | South_western |
| UG_461 | 0.044345978 | 0.106324118 | 0.02296515 | 0.000924897 | 0.825439857 | South_western |
| UG_469 | 5.23E-12 | 1.81E-12 | 2.62E-14 | 1.68E-15 | 1 | South_western |
| UG_389 | 0.000123004 | 0.000155376 | 0.082606749 | 0.917114872 | 1.71E-15 | Northern |
| UG_397 | 1.16E-12 | 3.44E-13 | 4.37E-15 | 3.10E-16 | 1 | South_western |
| UG_405 | 3.69E-12 | 1.26E-12 | 1.71E-14 | 1.08E-15 | 1 | South_western |
| UG_413 | 9.87E-13 | 2.86E-13 | 3.61E-15 | 2.60E-16 | 1 | South_western |

| | | | | | | |
|---|---|---|---|---|---|---|
| UG_421 | 5.65E-12 | 2.10E-12 | 2.78E-14 | 1.56E-15 | 1 | South_western |
| UG_429 | 2.96E-12 | 1.07E-12 | 1.26E-14 | 6.77E-16 | 1 | South_western |
| UG_437 | 1.82E-12 | 5.74E-13 | 7.43E-15 | 4.98E-16 | 1 | South_western |
| UG_445 | 3.70E-13 | 1.03E-13 | 1.08E-15 | 7.26E-17 | 1 | South_western |
| UG_382 | 0.001511358 | 0.009518296 | 0.655271427 | 0.333698918 | 4.16E-15 | Northern |
| UG_454 | 0.038641954 | 0.091675132 | 0.019317577 | 0.000775123 | 0.849590214 | South_western |
| UG_462 | 6.94E-13 | 2.09E-13 | 2.28E-15 | 1.43E-16 | 1 | South_western |
| UG_470 | 2.57E-07 | 2.16E-07 | 1.14E-08 | 6.12E-10 | 0.999999515 | South_western |
| UG_390 | 0.004363371 | 0.147644515 | 0.834523424 | 0.013468691 | 9.60E-15 | Northern |
| UG_398 | 7.39E-12 | 2.73E-12 | 3.90E-14 | 2.32E-15 | 1 | South_western |
| UG_406 | 7.36E-12 | 2.80E-12 | 3.82E-14 | 2.14E-15 | 1 | South_western |
| UG_414 | 3.24E-10 | 1.50E-10 | 3.85E-12 | 2.58E-13 | 1 | South_western |
| UG_422 | 3.08E-09 | 1.74E-09 | 5.69E-11 | 3.52E-12 | 0.999999995 | South_western |
| UG_430 | 1.83E-12 | 6.20E-13 | 7.19E-15 | 4.14E-16 | 1 | South_western |
| UG_438 | 4.36E-12 | 1.52E-12 | 2.08E-14 | 1.29E-15 | 1 | South_western |
| UG_446 | 4.10E-12 | 1.41E-12 | 1.94E-14 | 1.21E-15 | 1 | South_western |
| UG_383 | 0.003289568 | 0.629001139 | 0.367504007 | 0.000205286 | 1.65E-15 | Northern |
| UG_455 | 2.63E-06 | 2.91E-06 | 1.80E-07 | 7.85E-09 | 0.999994266 | South_western |
| UG_463 | 2.56E-12 | 8.54E-13 | 1.10E-14 | 6.80E-16 | 1 | South_western |
| UG_391 | 3.42E-12 | 1.22E-12 | 1.52E-14 | 8.60E-16 | 1 | South_western |
| UG_399 | 3.29E-13 | 9.23E-14 | 9.35E-16 | 6.11E-17 | 1 | South_western |
| UG_407 | 3.06E-12 | 1.12E-12 | 1.31E-14 | 7.01E-16 | 1 | South_western |
| UG_415 | 3.03E-13 | 8.67E-14 | 8.34E-16 | 5.14E-17 | 1 | South_western |
| UG_423 | 3.28E-12 | 1.06E-12 | 1.52E-14 | 1.05E-15 | 1 | South_western |
| UG_431 | 3.06E-12 | 1.06E-12 | 1.35E-14 | 8.03E-16 | 1 | South_western |
| UG_439 | 2.85E-12 | 9.82E-13 | 1.23E-14 | 7.31E-16 | 1 | South_western |
| UG_447 | 4.58E-13 | 1.34E-13 | 1.38E-15 | 8.69E-17 | 1 | South_western |
| UG_384 | 0.001653849 | 0.007883493 | 0.575326758 | 0.415135899 | 2.02E-14 | Northern |
| UG_456 | 4.14E-08 | 3.00E-08 | 1.27E-09 | 7.06E-11 | 0.999999927 | South_western |

| | | | | | | |
|---|---|---|---|---|---|---|
| UG_464 | 4.13E-12 | 1.41E-12 | 1.97E-14 | 1.27E-15 | 1 | South_western |
| UG_392 | 1.78E-12 | 5.89E-13 | 7.03E-15 | 4.23E-16 | 1 | South_western |
| UG_400 | 2.27E-12 | 7.70E-13 | 9.36E-15 | 5.52E-16 | 1 | South_western |
| UG_408 | 1.40E-11 | 5.59E-12 | 8.28E-14 | 4.61E-15 | 1 | South_western |
| UG_416 | 3.54E-13 | 1.02E-13 | 1.01E-15 | 6.33E-17 | 1 | South_western |
| UG_424 | 2.91E-12 | 1.00E-12 | 1.27E-14 | 7.57E-16 | 1 | South_western |
| UG_432 | 1.96E-11 | 8.00E-12 | 1.24E-13 | 6.96E-15 | 1 | South_western |
| UG_440 | 2.04E-12 | 6.85E-13 | 8.27E-15 | 4.92E-16 | 1 | South_western |
| UG_448 | 1.60E-12 | 5.07E-13 | 6.31E-15 | 4.10E-16 | 1 | South_western |
| UG_471 | 0.00058096 | 0.000903607 | 0.00012442 | 6.13E-06 | 0.99838488 | South_western |
| UG_543 | 0.00077835 | 0.00388602 | 0.502434889 | 0.49290074 | 6.90E-16 | Northern |
| UG_551 | 0.025751147 | 0.501870489 | 0.467860467 | 0.004517894 | 2.74E-09 | Northern |
| UG_559 | 0.000161719 | 0.000215395 | 0.096275066 | 0.90334782 | 3.29E-15 | North_western |
| UG_479 | 6.77E-12 | 2.40E-12 | 3.57E-14 | 2.28E-15 | 1 | South_western |
| UG_487 | 5.00E-11 | 2.14E-11 | 3.91E-13 | 2.30E-14 | 1 | South_western |
| UG_495 | 3.27E-12 | 1.14E-12 | 1.46E-14 | 8.67E-16 | 1 | South_western |
| UG_503 | 4.66E-12 | 1.70E-12 | 2.21E-14 | 1.26E-15 | 1 | South_western |
| UG_511 | 0.004224436 | 0.146334603 | 0.836204369 | 0.013236593 | 7.68E-15 | Northern |
| UG_519 | 0.111301324 | 0.536624509 | 0.339461113 | 0.012276765 | 0.000336288 | Northern |
| UG_527 | 0.001062696 | 0.005845975 | 0.57130971 | 0.421781619 | 1.62E-15 | Northern |
| UG_535 | 0.009262608 | 0.186887953 | 0.785086303 | 0.018763136 | 1.70E-12 | Northern |
| UG_472 | 4.92E-08 | 3.34E-08 | 1.63E-09 | 1.06E-10 | 0.999999916 | South_western |
| UG_544 | 0.001407674 | 0.009740431 | 0.676834519 | 0.312017376 | 2.08E-15 | Northern |
| UG_552 | 0.007788029 | 0.605040007 | 0.386545921 | 0.000626043 | 7.26E-13 | Northern |
| UG_560 | 0.000121561 | 0.000210302 | 0.11289735 | 0.886770788 | 2.31E-16 | North_western |
| UG_480 | 3.82E-12 | 1.27E-12 | 1.81E-14 | 1.20E-15 | 1 | South_western |
| UG_488 | 3.68E-13 | 1.04E-13 | 1.07E-15 | 7.02E-17 | 1 | South_western |
| UG_496 | 1.16E-11 | 4.46E-12 | 6.64E-14 | 3.85E-15 | 1 | South_western |
| UG_504 | 1.31E-12 | 3.96E-13 | 5.00E-15 | 3.43E-16 | 1 | South_western |

| | | | | | | |
|---|---|---|---|---|---|---|
| UG_512 | 0.003063143 | 0.654654745 | 0.3421248 | 0.000157312 | 1.21E-15 | Northern |
| UG_520 | 0.003958791 | 0.122175475 | 0.856329506 | 0.017536228 | 6.40E-15 | Northern |
| UG_528 | 0.055966884 | 0.389634834 | 0.533625986 | 0.020771559 | 7.37E-07 | Northern |
| UG_536 | 0.009175009 | 0.185099778 | 0.786779795 | 0.018945417 | 1.60E-12 | Northern |
| UG_473 | 3.00E-09 | 1.83E-09 | 5.30E-11 | 2.80E-12 | 0.999999995 | South_western |
| UG_545 | 0.003815546 | 0.128054904 | 0.85273473 | 0.01539482 | 4.52E-15 | Northern |
| UG_553 | 0.000138104 | 0.000235255 | 0.116355558 | 0.883271083 | 4.04E-16 | North_western |
| UG_561 | 0.000135698 | 0.000227355 | 0.113738503 | 0.885898443 | 4.20E-16 | North_western |
| UG_481 | 2.79E-12 | 9.14E-13 | 1.24E-14 | 8.15E-16 | 1 | South_western |
| UG_489 | 3.08E-12 | 1.04E-12 | 1.38E-14 | 8.68E-16 | 1 | South_western |
| UG_497 | 4.69E-06 | 5.31E-06 | 3.64E-07 | 1.65E-08 | 0.999989611 | South_western |
| UG_505 | 2.07E-12 | 7.12E-13 | 8.33E-15 | 4.74E-16 | 1 | South_western |
| UG_513 | 0.049607036 | 0.345657808 | 0.579172709 | 0.025562162 | 2.85E-07 | Northern |
| UG_521 | 0.003808871 | 0.09323693 | 0.875055189 | 0.027899011 | 7.87E-15 | Northern |
| UG_529 | 0.003912377 | 0.122353653 | 0.856467268 | 0.017266702 | 5.86E-15 | Northern |
| UG_537 | 0.003967338 | 0.130403962 | 0.850126841 | 0.015501859 | 5.83E-15 | Northern |
| UG_474 | 9.73E-13 | 3.00E-13 | 3.44E-15 | 2.18E-16 | 1 | South_western |
| UG_546 | 0.003730371 | 0.116894092 | 0.861477333 | 0.017898204 | 4.47E-15 | Northern |
| UG_554 | 0.000155929 | 0.000214839 | 0.09831018 | 0.901319053 | 2.34E-15 | North_western |
| UG_562 | 0.000133937 | 0.000228254 | 0.115140445 | 0.884497364 | 3.60E-16 | North_western |
| UG_482 | 1.43E-12 | 4.76E-13 | 5.33E-15 | 3.06E-16 | 1 | South_western |
| UG_490 | 6.33E-13 | 1.83E-13 | 2.08E-15 | 1.40E-16 | 1 | South_western |
| UG_498 | 0.035389799 | 0.08387898 | 0.017258504 | 0.000682953 | 0.862789764 | South_western |
| UG_506 | 2.19E-12 | 7.35E-13 | 9.05E-15 | 5.48E-16 | 1 | South_western |
| UG_514 | 0.000841629 | 0.005101085 | 0.576149506 | 0.417907779 | 3.89E-16 | Northern |
| UG_522 | 0.003752969 | 0.122984929 | 0.856914775 | 0.016347327 | 4.28E-15 | Northern |
| UG_530 | 0.000349832 | 0.001273142 | 0.319886368 | 0.678490659 | 1.46E-16 | Northern |
| UG_538 | 0.005887867 | 0.370773331 | 0.621008724 | 0.002330078 | 3.94E-14 | Northern |
| UG_475 | 1.07E-12 | 3.21E-13 | 3.93E-15 | 2.69E-16 | 1 | South_western |

| | | | | | | |
|---|---|---|---|---|---|---|
| UG_547 | 0.001156788 | 0.005758555 | 0.548051526 | 0.445033131 | 3.65E-15 | Northern |
| UG_555 | 0.000128096 | 0.000219702 | 0.114045199 | 0.885607004 | 2.95E-16 | North_western |
| UG_563 | 0.000135304 | 0.000232459 | 0.116491554 | 0.883140682 | 3.56E-16 | North_western |
| UG_483 | 1.55E-12 | 5.10E-13 | 5.90E-15 | 3.49E-16 | 1 | South_western |
| UG_491 | 2.09E-12 | 7.20E-13 | 8.38E-15 | 4.73E-16 | 1 | South_western |
| UG_499 | 5.54E-13 | 1.66E-13 | 1.73E-15 | 1.06E-16 | 1 | South_western |
| UG_507 | 3.88E-12 | 1.36E-12 | 1.79E-14 | 1.07E-15 | 1 | South_western |
| UG_515 | 0.05837665 | 0.394820765 | 0.526015193 | 0.020786363 | 1.03E-06 | Northern |
| UG_523 | 0.003118515 | 0.090814715 | 0.882394942 | 0.023671828 | 1.90E-15 | Northern |
| UG_531 | 0.00373503 | 0.607505875 | 0.388483833 | 0.000275261 | 3.54E-15 | Northern |
| UG_539 | 0.003992285 | 0.147650198 | 0.836127262 | 0.012230255 | 5.02E-15 | Northern |
| UG_476 | 2.63E-12 | 8.58E-13 | 1.15E-14 | 7.56E-16 | 1 | South_western |
| UG_548 | 0.003519317 | 0.583382006 | 0.412793554 | 0.000305123 | 1.96E-15 | Northern |
| UG_556 | 0.00015781 | 0.000212615 | 0.09653628 | 0.903093295 | 2.81E-15 | North_western |
| UG_564 | 0.000124335 | 0.000165753 | 0.087595471 | 0.912114441 | 1.27E-15 | North_western |
| UG_484 | 1.22E-12 | 3.91E-13 | 4.43E-15 | 2.66E-16 | 1 | South_western |
| UG_492 | 1.41E-11 | 5.28E-12 | 8.60E-14 | 5.44E-15 | 1 | South_western |
| UG_500 | 9.52E-13 | 2.78E-13 | 3.44E-15 | 2.43E-16 | 1 | South_western |
| UG_508 | 0.115164013 | 0.551271408 | 0.321899452 | 0.011160558 | 0.000504569 | Northern |
| UG_516 | 0.000417788 | 0.001638645 | 0.358086072 | 0.639857494 | 1.94E-16 | Northern |
| UG_524 | 0.118333324 | 0.582047927 | 0.289954461 | 0.008826264 | 0.000838023 | Northern |
| UG_532 | 0.00630657 | 0.142936274 | 0.829441073 | 0.021316083 | 1.50E-13 | Northern |
| UG_540 | 0.026805586 | 0.570494848 | 0.399775297 | 0.002924263 | 5.30E-09 | Northern |
| UG_477 | 8.67E-12 | 3.18E-12 | 4.77E-14 | 2.95E-15 | 1 | South_western |
| UG_549 | 0.004925831 | 0.151966024 | 0.828608403 | 0.014499741 | 2.23E-14 | Northern |
| UG_557 | 0.000125836 | 0.000208753 | 0.109621388 | 0.890044023 | 3.40E-16 | North_western |
| UG_485 | 0.000295832 | 0.000473768 | 5.29E-05 | 2.21E-06 | 0.999175299 | South_western |
| UG_493 | 5.82E-13 | 1.80E-13 | 1.80E-15 | 1.04E-16 | 1 | South_western |
| UG_501 | 2.55E-12 | 8.73E-13 | 1.08E-14 | 6.33E-16 | 1 | South_western |

| | | | | | | |
|---|---|---|---|---|---|---|
| UG_509 | 0.00305052 | 0.591645712 | 0.405057778 | 0.00024599 | 7.33E-16 | Northern |
| UG_517 | 0.141488115 | 0.615561723 | 0.229235455 | 0.006669345 | 0.007045362 | Northern |
| UG_525 | 0.00358541 | 0.600579028 | 0.395559296 | 0.000276266 | 2.51E-15 | Northern |
| UG_533 | 0.109257725 | 0.545493631 | 0.333585736 | 0.011357713 | 0.000305195 | Northern |
| UG_541 | 0.008694635 | 0.222527723 | 0.756643873 | 0.012133769 | 8.63E-13 | Northern |
| UG_478 | 8.19E-12 | 3.01E-12 | 4.44E-14 | 2.71E-15 | 1 | South_western |
| UG_550 | 0.000922005 | 0.004645735 | 0.525398793 | 0.469033467 | 1.33E-15 | Northern |
| UG_558 | 0.000105804 | 0.000133803 | 0.078269636 | 0.921490757 | 9.90E-16 | North_western |
| UG_486 | 2.19E-10 | 1.05E-10 | 2.30E-12 | 1.34E-13 | 1 | South_western |
| UG_494 | 4.18E-12 | 1.45E-12 | 1.98E-14 | 1.23E-15 | 1 | South_western |
| UG_502 | 1.03E-12 | 3.08E-13 | 3.76E-15 | 2.57E-16 | 1 | South_western |
| UG_510 | 0.126846777 | 0.598659004 | 0.264772468 | 0.007833915 | 0.001887836 | Northern |
| UG_518 | 0.004049334 | 0.145378761 | 0.837763879 | 0.012808025 | 5.70E-15 | Northern |
| UG_526 | 0.004274304 | 0.10081302 | 0.867603053 | 0.027309624 | 1.59E-14 | Northern |
| UG_534 | 0.057813279 | 0.392820547 | 0.528462683 | 0.020902538 | 9.52E-07 | Northern |
| UG_542 | 0.009342377 | 0.181327731 | 0.789188953 | 0.020140939 | 1.88E-12 | Northern |
| UG_565 | 0.000125133 | 0.000215309 | 0.113457102 | 0.886202456 | 2.65E-16 | North_western |
| UG_637 | 0.000114775 | 0.000174875 | 0.097382564 | 0.902327785 | 4.13E-16 | North_western |
| UG_645 | 0.009265025 | 0.599001677 | 0.390945062 | 0.000788236 | 2.48E-12 | Eastern |
| UG_653 | 0.084866905 | 0.600738657 | 0.307988155 | 0.006350486 | 5.58E-05 | Eastern |
| UG_573 | 0.000184918 | 0.000285275 | 0.117014094 | 0.882515713 | 2.15E-15 | North_western |
| UG_581 | 0.000182957 | 0.000283618 | 0.117131431 | 0.882401994 | 2.01E-15 | North_western |
| UG_589 | 0.000146383 | 0.000237936 | 0.113388875 | 0.886226806 | 6.67E-16 | North_western |
| UG_597 | 0.000159443 | 0.000212532 | 0.095858875 | 0.90376915 | 3.11E-15 | North_western |
| UG_605 | 0.000111637 | 0.000149745 | 0.084752063 | 0.914986555 | 8.30E-16 | North_western |
| UG_613 | 9.33E-05 | 0.000143803 | 0.091392596 | 0.908370291 | 1.82E-16 | North_western |
| UG_621 | 0.000122262 | 0.000203482 | 0.108846468 | 0.890827788 | 3.00E-16 | North_western |
| UG_629 | 0.000102962 | 0.000149304 | 0.08906048 | 0.910687255 | 3.81E-16 | North_western |
| UG_566 | 0.000105889 | 0.000153299 | 0.089824826 | 0.909915986 | 4.25E-16 | North_western |

| | | | | | | |
|---|---|---|---|---|---|---|
| UG_638 | 0.000131325 | 0.000228783 | 0.116861254 | 0.882778638 | 2.93E-16 | North_western |
| UG_646 | 0.004512458 | 0.648642479 | 0.346593117 | 0.000251946 | 1.92E-14 | Eastern |
| UG_654 | 0.076627914 | 0.47431992 | 0.434077535 | 0.01496377 | 1.09E-05 | Eastern |
| UG_574 | 0.000132341 | 0.000223929 | 0.113836486 | 0.885807244 | 3.61E-16 | North_western |
| UG_582 | 0.002970553 | 0.60159353 | 0.395213098 | 0.00022282 | 6.46E-16 | North_western |
| UG_590 | 0.000147925 | 0.000217785 | 0.10310239 | 0.896531901 | 1.28E-15 | North_western |
| UG_598 | 0.000140441 | 0.000204194 | 0.099927419 | 0.899727946 | 1.15E-15 | North_western |
| UG_606 | 0.000147745 | 0.000236504 | 0.112045322 | 0.887570429 | 7.58E-16 | North_western |
| UG_614 | 8.46E-05 | 0.000103206 | 0.069499424 | 0.930312793 | 5.56E-16 | North_western |
| UG_622 | 0.000122012 | 0.000202493 | 0.108460147 | 0.891215348 | 3.03E-16 | North_western |
| UG_630 | 0.124255318 | 0.596650292 | 0.269641554 | 0.007931851 | 0.001520985 | North_western |
| UG_567 | 0.000116827 | 0.000198866 | 0.109521659 | 0.890162648 | 2.21E-16 | North_western |
| UG_639 | 0.00012187 | 0.000203325 | 0.108986228 | 0.890688577 | 2.92E-16 | North_western |
| UG_647 | 0.040497319 | 0.555171283 | 0.399469739 | 0.004861549 | 1.10E-07 | Eastern |
| UG_655 | 0.061214164 | 0.655759525 | 0.27978715 | 0.003232651 | 6.51E-06 | Eastern |
| UG_575 | 0.000131135 | 0.000181211 | 0.092612062 | 0.907075592 | 1.23E-15 | North_western |
| UG_583 | 0.00012537 | 0.000213353 | 0.11229488 | 0.887366397 | 2.86E-16 | North_western |
| UG_591 | 0.000168258 | 0.000232275 | 0.101243445 | 0.898356022 | 3.04E-15 | North_western |
| UG_599 | 0.001825821 | 0.007307693 | 0.529006639 | 0.461859847 | 6.82E-14 | North_western |
| UG_607 | 0.007101694 | 0.08409483 | 0.845354719 | 0.063448756 | 1.01E-12 | North_western |
| UG_615 | 6.59E-05 | 8.34E-05 | 0.065865808 | 0.933984898 | 1.79E-16 | North_western |
| UG_623 | 0.006762017 | 0.08159056 | 0.848005126 | 0.063642297 | 7.44E-13 | North_western |
| UG_631 | 0.000122523 | 0.00020309 | 0.108489233 | 0.891185154 | 3.10E-16 | North_western |
| UG_568 | 0.000149452 | 0.000238135 | 0.111990743 | 0.887621671 | 8.13E-16 | North_western |
| UG_640 | 0.000118909 | 0.000202632 | 0.110337787 | 0.889340673 | 2.34E-16 | North_western |
| UG_648 | 0.047148085 | 0.653984422 | 0.296206115 | 0.002660588 | 7.90E-07 | Eastern |
| UG_656 | 0.010295205 | 0.628051943 | 0.360935844 | 0.000717008 | 6.63E-12 | Eastern |
| UG_576 | 0.000125824 | 0.000212322 | 0.1114973 | 0.888164555 | 3.05E-16 | North_western |
| UG_584 | 0.000126591 | 0.000206565 | 0.108059718 | 0.891607126 | 3.84E-16 | North_western |

| | | | | | | |
|---|---|---|---|---|---|---|
| UG_592 | 0.000146816 | 0.000237408 | 0.112925504 | 0.886690272 | 6.96E-16 | North_western |
| UG_608 | 0.000131819 | 0.000225073 | 0.114703888 | 0.884939221 | 3.36E-16 | North_western |
| UG_616 | 9.15E-05 | 0.000143923 | 0.092626519 | 0.907138043 | 1.50E-16 | North_western |
| UG_624 | 9.78E-05 | 0.000138095 | 0.085108736 | 0.914655417 | 3.72E-16 | North_western |
| UG_632 | 0.000146946 | 0.000192719 | 0.091552317 | 0.908108018 | 2.57E-15 | North_western |
| UG_569 | 9.85E-05 | 0.000143524 | 0.08804816 | 0.911709785 | 3.16E-16 | North_western |
| UG_641 | 0.000145357 | 0.000193066 | 0.09236715 | 0.907294427 | 2.28E-15 | North_western |
| UG_649 | 0.058172563 | 0.697059039 | 0.242585746 | 0.002175556 | 7.10E-06 | Eastern |
| UG_657 | 0.084475154 | 0.59950019 | 0.309577729 | 0.00639385 | 5.31E-05 | Eastern |
| UG_577 | 0.000135067 | 0.000201374 | 0.101048259 | 0.8986153 | 8.51E-16 | North_western |
| UG_585 | 0.000116219 | 0.000163381 | 0.09019232 | 0.90952808 | 7.15E-16 | North_western |
| UG_593 | 0.000124141 | 0.00021169 | 0.112126567 | 0.887537602 | 2.73E-16 | North_western |
| UG_601 | 0.000107396 | 0.000145048 | 0.084147751 | 0.915599805 | 6.91E-16 | North_western |
| UG_609 | 0.000120739 | 0.000161444 | 0.086929643 | 0.912788174 | 1.12E-15 | North_western |
| UG_617 | 7.87E-05 | 9.20E-05 | 0.064799372 | 0.935029938 | 5.60E-16 | North_western |
| UG_625 | 0.000152756 | 0.000207632 | 0.096259001 | 0.903380612 | 2.36E-15 | North_western |
| UG_633 | 0.000156763 | 0.000214959 | 0.098027649 | 0.901600628 | 2.46E-15 | North_western |
| UG_570 | 0.000126283 | 0.00018243 | 0.095542015 | 0.904149272 | 8.13E-16 | North_western |
| UG_642 | 0.000201124 | 0.000337598 | 0.131027405 | 0.868433873 | 1.73E-15 | North_western |
| UG_650 | 0.084298274 | 0.657841776 | 0.253837494 | 0.003925862 | 9.66E-05 | Eastern |
| UG_658 | 0.087035085 | 0.576632148 | 0.328483794 | 0.007792867 | 5.61E-05 | Eastern |
| UG_578 | 0.000129526 | 0.000208015 | 0.107232087 | 0.892430371 | 4.61E-16 | North_western |
| UG_586 | 0.000143163 | 0.000227663 | 0.110062877 | 0.889566297 | 7.04E-16 | North_western |
| UG_594 | 0.000125433 | 0.000206316 | 0.10856638 | 0.89110187 | 3.54E-16 | North_western |
| UG_602 | 0.00012245 | 0.000208713 | 0.111527778 | 0.888141058 | 2.60E-16 | North_western |
| UG_610 | 0.00012044 | 0.000201518 | 0.108838196 | 0.890839846 | 2.75E-16 | North_western |
| UG_618 | 7.50E-05 | 0.000100843 | 0.07347349 | 0.926350714 | 1.95E-16 | North_western |
| UG_626 | 0.007005727 | 0.08279587 | 0.84591628 | 0.064282123 | 9.40E-13 | North_western |
| UG_634 | 0.000138106 | 0.000237676 | 0.117543131 | 0.882081087 | 3.79E-16 | North_western |

| | | | | | | |
|---|---|---|---|---|---|---|
| UG_571 | 0.000124887 | 0.000211749 | 0.111729447 | 0.887933917 | 2.89E-16 | North_western |
| UG_643 | 0.006436575 | 0.608531508 | 0.384535465 | 0.000496452 | 1.86E-13 | Eastern |
| UG_651 | 0.073242512 | 0.540624596 | 0.377120255 | 0.009002083 | 1.06E-05 | Eastern |
| UG_579 | 0.000106044 | 0.000135667 | 0.079264075 | 0.920494213 | 9.28E-16 | North_western |
| UG_587 | 0.000125906 | 0.000205453 | 0.107852921 | 0.891815721 | 3.77E-16 | North_western |
| UG_595 | 0.000131813 | 0.00022592 | 0.115136138 | 0.884506129 | 3.28E-16 | North_western |
| UG_603 | 0.000147776 | 0.000190621 | 0.090214874 | 0.909446729 | 2.91E-15 | North_western |
| UG_611 | 0.000102115 | 0.000152944 | 0.091747044 | 0.907997898 | 3.02E-16 | North_western |
| UG_619 | 0.000110356 | 0.000150662 | 0.085922029 | 0.913816953 | 7.13E-16 | North_western |
| UG_627 | 0.000126232 | 0.000215919 | 0.113144092 | 0.886513757 | 2.84E-16 | North_western |
| UG_635 | 0.007137096 | 0.084432454 | 0.84510328 | 0.06332717 | 1.03E-12 | North_western |
| UG_572 | 0.0001185 | 0.00015458 | 0.084197835 | 0.915529085 | 1.22E-15 | North_western |
| UG_644 | 0.084148685 | 0.656894533 | 0.25490782 | 0.003954935 | 9.40E-05 | Eastern |
| UG_652 | 0.079308395 | 0.576764713 | 0.336596837 | 0.007304082 | 2.60E-05 | Eastern |
| UG_580 | 9.05E-05 | 0.000124682 | 0.080677875 | 0.919106972 | 3.29E-16 | North_western |
| UG_588 | 0.000135791 | 0.000195975 | 0.097986834 | 0.9016814 | 1.06E-15 | North_western |
| UG_596 | 0.000136248 | 0.000191933 | 0.095744462 | 0.903927356 | 1.25E-15 | North_western |
| UG_604 | 0.000159925 | 0.000267042 | 0.120191944 | 0.879381089 | 7.78E-16 | North_western |
| UG_612 | 0.000115643 | 0.000191716 | 0.106278083 | 0.893414558 | 2.51E-16 | North_western |
| UG_620 | 9.65E-05 | 0.000151989 | 0.094577559 | 0.905173969 | 1.79E-16 | North_western |
| UG_628 | 0.000161291 | 0.00029562 | 0.132150516 | 0.867392573 | 4.53E-16 | North_western |
| UG_636 | 0.006608415 | 0.081365193 | 0.849518775 | 0.062507616 | 6.29E-13 | North_western |
| UG_659 | 0.084641135 | 0.600818479 | 0.308150255 | 0.006335528 | 5.46E-05 | Eastern |
| UG_731 | 0.004097146 | 0.579814524 | 0.415718466 | 0.000369864 | 5.77E-15 | Eastern |
| UG_739 | 0.004875252 | 0.510963818 | 0.483444436 | 0.000716494 | 1.42E-14 | Eastern |
| UG_747 | 0.016880429 | 0.456577387 | 0.522565411 | 0.003976774 | 1.02E-10 | Eastern |
| UG_667 | 0.090158425 | 0.565819592 | 0.335289073 | 0.008663658 | 6.93E-05 | Eastern |
| UG_675 | 0.086617047 | 0.694734462 | 0.215623894 | 0.002820863 | 0.000203735 | Eastern |
| UG_683 | 0.035480511 | 0.529602807 | 0.429788475 | 0.005128172 | 3.48E-08 | Eastern |

| | | | | | | |
|---|---|---|---|---|---|---|
| UG_691 | 0.006811667 | 0.472301966 | 0.519542184 | 0.001344183 | 1.40E-13 | Eastern |
| UG_699 | 0.119376287 | 0.591961325 | 0.27950929 | 0.008152523 | 0.001000574 | Eastern |
| UG_707 | 0.141245561 | 0.599088635 | 0.246324557 | 0.007846234 | 0.005495013 | Eastern |
| UG_715 | 0.151045102 | 0.612002742 | 0.215749305 | 0.006567552 | 0.014635299 | Eastern |
| UG_723 | 0.003667374 | 0.092539863 | 0.876619694 | 0.02717307 | 6.05E-15 | Northern |
| UG_660 | 0.081255972 | 0.665424608 | 0.249647821 | 0.003594178 | 7.74E-05 | Eastern |
| UG_732 | 0.008041845 | 0.399605583 | 0.589688466 | 0.002664106 | 3.93E-13 | Eastern |
| UG_740 | 0.012760916 | 0.505272815 | 0.47984643 | 0.002119839 | 1.55E-11 | Eastern |
| UG_748 | 0.004325995 | 0.53395817 | 0.461178363 | 0.000537472 | 6.65E-15 | Eastern |
| UG_668 | 0.009292839 | 0.502118038 | 0.487049622 | 0.001539501 | 1.50E-12 | Eastern |
| UG_676 | 0.043990862 | 0.377203192 | 0.560613995 | 0.018191837 | 1.14E-07 | Eastern |
| UG_684 | 0.008157339 | 0.587825819 | 0.403273849 | 0.000742993 | 9.07E-13 | Eastern |
| UG_692 | 0.062305655 | 0.667374526 | 0.267341874 | 0.002969364 | 8.58E-06 | Eastern |
| UG_700 | 0.11791144 | 0.598743013 | 0.274723073 | 0.007663625 | 0.000958848 | Eastern |
| UG_708 | 0.14447484 | 0.606519118 | 0.233990516 | 0.007244105 | 0.007771421 | Eastern |
| UG_716 | 0.151399554 | 0.608929765 | 0.218632198 | 0.006788075 | 0.014250409 | Eastern |
| UG_724 | 0.003279291 | 0.06386669 | 0.886203254 | 0.046650765 | 5.67E-15 | Northern |
| UG_661 | 0.087061741 | 0.578144036 | 0.32703304 | 0.007704244 | 5.69E-05 | Eastern |
| UG_733 | 0.005121143 | 0.604008722 | 0.390470923 | 0.000399211 | 3.41E-14 | Eastern |
| UG_741 | 0.021258263 | 0.398661546 | 0.572509798 | 0.007570391 | 4.92E-10 | Eastern |
| UG_749 | 0.003400367 | 0.105523514 | 0.871418074 | 0.019658044 | 2.72E-15 | Eastern |
| UG_669 | 0.082593489 | 0.603037134 | 0.308197852 | 0.006126141 | 4.54E-05 | Eastern |
| UG_677 | 0.003497111 | 0.088142309 | 0.880125336 | 0.028235244 | 4.68E-15 | Eastern |
| UG_685 | 0.025857712 | 0.432963254 | 0.533895289 | 0.007283743 | 2.25E-09 | Eastern |
| UG_693 | 0.017102643 | 0.654418769 | 0.327473202 | 0.001005385 | 3.51E-10 | Eastern |
| UG_701 | 2.19E-12 | 7.37E-13 | 9.01E-15 | 5.39E-16 | 1 | South_western |
| UG_709 | 0.006558223 | 0.501333254 | 0.49104956 | 0.001058963 | 1.18E-13 | Eastern |
| UG_717 | 0.005407686 | 0.169828062 | 0.811939697 | 0.012824555 | 3.74E-14 | Northern |
| UG_725 | 0.002381599 | 0.038578589 | 0.881094594 | 0.077945218 | 1.69E-15 | Northern |

| | | | | | | |
|---|---|---|---|---|---|---|
| UG_662 | 0.007106486 | 0.34481761 | 0.644606306 | 0.003469598 | 1.53E-13 | Eastern |
| UG_734 | 0.004896819 | 0.583131668 | 0.411531716 | 0.000439797 | 2.15E-14 | Eastern |
| UG_742 | 0.004614721 | 0.657926518 | 0.337217752 | 0.000241009 | 2.44E-14 | Eastern |
| UG_750 | 0.007077126 | 0.469002945 | 0.522486545 | 0.001433383 | 1.83E-13 | Eastern |
| UG_670 | 0.046164646 | 0.109545449 | 0.024320292 | 0.001008129 | 0.818961483 | Eastern |
| UG_678 | 0.062970615 | 0.657768102 | 0.276006687 | 0.00324621 | 8.39E-06 | Eastern |
| UG_686 | 0.083152736 | 0.659742605 | 0.25318607 | 0.003830681 | 8.79E-05 | Eastern |
| UG_694 | 0.085904656 | 0.661396568 | 0.248736867 | 0.003843037 | 0.000118872 | Eastern |
| UG_702 | 4.89E-12 | 1.75E-12 | 2.37E-14 | 1.42E-15 | 1 | South_western |
| UG_710 | 0.125615094 | 0.5710134 | 0.29219052 | 0.009880284 | 0.001300703 | Eastern |
| UG_718 | 0.059067678 | 0.397137009 | 0.523131998 | 0.020662183 | 1.13E-06 | Northern |
| UG_726 | 0.006580415 | 0.110209144 | 0.846791259 | 0.036419183 | 3.25E-13 | Northern |
| UG_663 | 0.003142167 | 0.583400142 | 0.413188532 | 0.00026916 | 8.62E-16 | Eastern |
| UG_735 | 0.003963353 | 0.56228652 | 0.433347823 | 0.000402303 | 4.09E-15 | Eastern |
| UG_743 | 0.018459301 | 0.46152214 | 0.515795614 | 0.004222945 | 1.99E-10 | Eastern |
| UG_751 | 0.007109681 | 0.480438492 | 0.511119034 | 0.001332793 | 1.96E-13 | Eastern |
| UG_671 | 0.02303653 | 0.608285318 | 0.366760668 | 0.001917482 | 2.21E-09 | Eastern |
| UG_679 | 0.003578063 | 0.534627877 | 0.461360166 | 0.000433894 | 1.69E-15 | Eastern |
| UG_687 | 0.081578378 | 0.663231976 | 0.251438219 | 0.003673511 | 7.79E-05 | Eastern |
| UG_695 | 0.085547094 | 0.655301013 | 0.254997565 | 0.004048047 | 0.000106281 | Eastern |
| UG_703 | 6.15E-12 | 2.27E-12 | 3.10E-14 | 1.80E-15 | 1 | South_western |
| UG_711 | 0.093891014 | 0.565232957 | 0.331847639 | 0.008931415 | 9.70E-05 | Eastern |
| UG_719 | 0.004984905 | 0.472834045 | 0.521230268 | 0.000950782 | 1.45E-14 | Northern |
| UG_727 | 0.032745398 | 0.606618877 | 0.357890901 | 0.002744794 | 3.09E-08 | Eastern |
| UG_664 | 0.008874818 | 0.530730269 | 0.459189033 | 0.00120588 | 1.22E-12 | Eastern |
| UG_736 | 0.005262584 | 0.598495894 | 0.395813944 | 0.000427578 | 4.00E-14 | Eastern |
| UG_744 | 0.019796263 | 0.458054149 | 0.517497283 | 0.004652304 | 3.31E-10 | Eastern |
| UG_752 | 0.003536846 | 0.575576624 | 0.420562717 | 0.000323813 | 1.94E-15 | Eastern |
| UG_672 | 0.010009354 | 0.525190993 | 0.46337347 | 0.001426183 | 2.85E-12 | Eastern |

| | | | | | | |
|---|---|---|---|---|---|---|
| UG_680 | 0.063835477 | 0.663800968 | 0.269238005 | 0.003115508 | 1.00E-05 | Eastern |
| UG_688 | 0.070010035 | 0.674267092 | 0.252666957 | 0.003031419 | 2.45E-05 | Eastern |
| UG_696 | 0.056799309 | 0.450878781 | 0.478613775 | 0.013707196 | 9.38E-07 | Eastern |
| UG_704 | 1.13E-12 | 3.74E-13 | 4.01E-15 | 2.26E-16 | 1 | South_western |
| UG_712 | 0.003536373 | 0.112954347 | 0.865473694 | 0.018035586 | 3.21E-15 | Eastern |
| UG_720 | 0.002994158 | 0.06472639 | 0.890772877 | 0.041506576 | 2.80E-15 | Northern |
| UG_728 | 0.043613397 | 0.387256918 | 0.552333587 | 0.01679599 | 1.07E-07 | Eastern |
| UG_665 | 0.010209041 | 0.506869219 | 0.48127161 | 0.001650129 | 3.04E-12 | Eastern |
| UG_737 | 0.012805954 | 0.488904994 | 0.495910656 | 0.002378396 | 1.49E-11 | Eastern |
| UG_745 | 0.012440452 | 0.480045945 | 0.505064479 | 0.002449124 | 1.16E-11 | Eastern |
| UG_673 | 0.005958572 | 0.600058469 | 0.393498469 | 0.00048449 | 9.98E-14 | Eastern |
| UG_681 | 0.00315682 | 0.57431699 | 0.422238121 | 0.000288069 | 8.43E-16 | Eastern |
| UG_689 | 0.035887735 | 0.517877692 | 0.440610084 | 0.005624453 | 3.58E-08 | Eastern |
| UG_697 | 0.004826022 | 0.583823961 | 0.410919274 | 0.000430743 | 1.94E-14 | Eastern |
| UG_705 | 2.18E-12 | 6.96E-13 | 9.17E-15 | 6.09E-16 | 1 | South_western |
| UG_713 | 0.116582154 | 0.595015283 | 0.279693835 | 0.007876572 | 0.000832156 | Eastern |
| UG_721 | 0.006697431 | 0.243145522 | 0.742629789 | 0.007527258 | 1.18E-13 | Northern |
| UG_729 | 0.042838844 | 0.372834631 | 0.566020249 | 0.018306183 | 9.26E-08 | Eastern |
| UG_666 | 0.118453433 | 0.599939776 | 0.272996297 | 0.007597865 | 0.001012629 | Eastern |
| UG_738 | 0.00696442 | 0.477938648 | 0.513771543 | 0.001325389 | 1.67E-13 | Eastern |
| UG_746 | 0.011513291 | 0.504202374 | 0.48237186 | 0.001912475 | 7.24E-12 | Eastern |
| UG_674 | 0.078233059 | 0.666817488 | 0.251420471 | 0.003472002 | 5.70E-05 | Eastern |
| UG_682 | 0.028159753 | 0.479967699 | 0.486126739 | 0.005745805 | 4.91E-09 | Eastern |
| UG_690 | 0.048967691 | 0.670381495 | 0.27824857 | 0.002400972 | 1.27E-06 | Eastern |
| UG_698 | 0.019443222 | 0.612226013 | 0.366762656 | 0.001568109 | 6.41E-10 | Eastern |
| UG_706 | 0.092247017 | 0.405529355 | 0.47421505 | 0.027968397 | 4.02E-05 | Central |
| UG_714 | 0.094483937 | 0.586927641 | 0.310892769 | 0.007573365 | 0.000122287 | Eastern |
| UG_722 | 0.046306687 | 0.321613024 | 0.603386252 | 0.028693866 | 1.71E-07 | Northern |
| UG_730 | 0.006350167 | 0.379309447 | 0.611960937 | 0.002379449 | 6.88E-14 | Eastern |

| | | | | | | |
|---|---|---|---|---|---|---|
| UG_753 | 0.022951152 | 0.448508115 | 0.522750532 | 0.0057902 | 9.64E-10 | Eastern |
| UG_825 | 0.004559568 | 0.04858418 | 0.845970336 | 0.100885916 | 1.32E-13 | Northern |
| UG_833 | 0.003249974 | 0.05986593 | 0.885262355 | 0.051621741 | 6.13E-15 | Northern |
| UG_841 | 0.004498012 | 0.040434213 | 0.82388774 | 0.131180035 | 1.97E-13 | Northern |
| UG_761 | 0.003316169 | 0.581196745 | 0.415197007 | 0.000290079 | 1.26E-15 | Eastern |
| UG_769 | 0.014648345 | 0.345128584 | 0.632670538 | 0.007552533 | 3.03E-11 | Eastern |
| UG_777 | 0.00509471 | 0.616403126 | 0.378138597 | 0.000363567 | 3.59E-14 | Eastern |
| UG_785 | 0.00533013 | 0.480409197 | 0.51328854 | 0.000972132 | 2.42E-14 | Eastern |
| UG_793 | 0.027157939 | 0.448433516 | 0.517529012 | 0.00687953 | 3.38E-09 | Eastern |
| UG_801 | 0.002804423 | 0.04000782 | 0.870990219 | 0.086197538 | 5.32E-15 | Northern |
| UG_809 | 0.002350671 | 0.022830827 | 0.807982995 | 0.166835506 | 6.66E-15 | Northern |
| UG_817 | 0.004569223 | 0.122942072 | 0.852278666 | 0.020210038 | 1.81E-14 | Northern |
| UG_754 | 0.003813595 | 0.581751657 | 0.414097557 | 0.000337192 | 3.47E-15 | Eastern |
| UG_826 | 0.003602536 | 0.109648306 | 0.867288088 | 0.019461069 | 3.87E-15 | Northern |
| UG_834 | 0.002086564 | 0.020188696 | 0.799728756 | 0.177995983 | 3.82E-15 | Northern |
| UG_842 | 0.004698965 | 0.145530567 | 0.834756478 | 0.015013989 | 1.69E-14 | Northern |
| UG_762 | 0.004242575 | 0.540097913 | 0.455154947 | 0.000504565 | 5.95E-15 | Eastern |
| UG_770 | 0.005171038 | 0.47142201 | 0.522407481 | 0.000999472 | 1.88E-14 | Eastern |
| UG_778 | 0.003865388 | 0.573597537 | 0.422174996 | 0.00036208 | 3.64E-15 | Eastern |
| UG_786 | 0.003912188 | 0.574333374 | 0.421389376 | 0.000365062 | 3.99E-15 | Eastern |
| UG_794 | 0.001868191 | 0.016502114 | 0.770903078 | 0.210726618 | 3.07E-15 | Northern |
| UG_802 | 0.003911367 | 0.112369831 | 0.863423425 | 0.020295377 | 6.78E-15 | Northern |
| UG_810 | 0.003593831 | 0.123381129 | 0.85752125 | 0.015503789 | 3.10E-15 | Northern |
| UG_818 | 0.005432232 | 0.09972598 | 0.859025095 | 0.035816693 | 9.51E-14 | Northern |
| UG_755 | 0.003807302 | 0.577568932 | 0.41827731 | 0.000346457 | 3.34E-15 | Eastern |
| UG_827 | 0.00435931 | 0.164025933 | 0.820719539 | 0.010895218 | 8.16E-15 | Northern |
| UG_835 | 0.003442666 | 0.084161797 | 0.882203163 | 0.030192374 | 4.57E-15 | Northern |
| UG_843 | 0.003335766 | 0.111722211 | 0.867650503 | 0.01729152 | 2.14E-15 | Northern |
| UG_763 | 0.006786759 | 0.491427932 | 0.500609571 | 0.001175738 | 1.46E-13 | Eastern |

| | | | | | | |
|---|---|---|---|---|---|---|
| UG_771 | 0.005989212 | 0.474202218 | 0.518656083 | 0.001152487 | 5.52E-14 | Eastern |
| UG_779 | 0.036828832 | 0.482875661 | 0.472952914 | 0.007342556 | 3.75E-08 | Eastern |
| UG_787 | 0.036722107 | 0.489685297 | 0.466606079 | 0.00698648 | 3.77E-08 | Eastern |
| UG_795 | 0.003727031 | 0.071123504 | 0.880968678 | 0.044180787 | 1.16E-14 | Northern |
| UG_803 | 0.004943194 | 0.157876358 | 0.823693585 | 0.013486863 | 2.16E-14 | Northern |
| UG_811 | 0.001316637 | 0.009462901 | 0.681375284 | 0.307845178 | 1.33E-15 | Northern |
| UG_819 | 0.003616334 | 0.126413872 | 0.855075285 | 0.014894508 | 3.12E-15 | Northern |
| UG_756 | 0.006918875 | 0.490025188 | 0.501843678 | 0.001212258 | 1.67E-13 | Eastern |
| UG_828 | 0.002520906 | 0.013914555 | 0.668625312 | 0.314939226 | 6.83E-14 | Northern |
| UG_836 | 0.005125342 | 0.138481967 | 0.838223583 | 0.018169108 | 3.44E-14 | Northern |
| UG_844 | 0.004582961 | 0.14388863 | 0.83658336 | 0.01494505 | 1.43E-14 | Northern |
| UG_764 | 0.007626943 | 0.468501332 | 0.522311034 | 0.001560692 | 3.14E-13 | Eastern |
| UG_772 | 0.012840834 | 0.519511704 | 0.465710545 | 0.001936916 | 1.72E-11 | Eastern |
| UG_780 | 0.003790403 | 0.564652942 | 0.431179842 | 0.000376813 | 3.00E-15 | Eastern |
| UG_788 | 0.006800459 | 0.489748171 | 0.502259537 | 0.001191834 | 1.47E-13 | Eastern |
| UG_796 | 0.003294701 | 0.117305182 | 0.863851105 | 0.015549012 | 1.79E-15 | Northern |
| UG_804 | 0.000517198 | 0.001784365 | 0.343457737 | 0.654240701 | 8.94E-16 | Northern |
| UG_812 | 0.004186821 | 0.127127747 | 0.85142876 | 0.017256673 | 9.02E-15 | Northern |
| UG_820 | 0.00490973 | 0.130345947 | 0.845233707 | 0.019510616 | 2.78E-14 | Northern |
| UG_757 | 0.004732322 | 0.511864119 | 0.482714368 | 0.000689191 | 1.15E-14 | Eastern |
| UG_829 | 0.003874503 | 0.104802098 | 0.868426 | 0.022897398 | 7.17E-15 | Northern |
| UG_837 | 0.003434174 | 0.084652914 | 0.882113414 | 0.029799498 | 4.43E-15 | Northern |
| UG_845 | 0.004009516 | 0.132758977 | 0.848087334 | 0.015144173 | 6.12E-15 | Northern |
| UG_765 | 0.009161553 | 0.537191766 | 0.452452322 | 0.001194358 | 1.58E-12 | Eastern |
| UG_773 | 0.004707322 | 0.594998807 | 0.399906153 | 0.000387718 | 1.74E-14 | Eastern |
| UG_781 | 0.008843444 | 0.326972237 | 0.659141251 | 0.005043067 | 7.57E-13 | Eastern |
| UG_789 | 0.005020936 | 0.513532007 | 0.48071973 | 0.000727327 | 1.78E-14 | Eastern |
| UG_797 | 0.005201099 | 0.078775046 | 0.864023252 | 0.052000603 | 1.12E-13 | Northern |
| UG_805 | 0.003588795 | 0.083022895 | 0.881063493 | 0.032324817 | 6.38E-15 | Northern |

| | | | | | | |
|---|---|---|---|---|---|---|
| UG_813 | 0.004942454 | 0.115365185 | 0.854907005 | 0.024785356 | 3.60E-14 | Northern |
| UG_821 | 0.004574041 | 0.040981987 | 0.823885645 | 0.130558327 | 2.16E-13 | Northern |
| UG_758 | 0.03534878 | 0.479254805 | 0.47816271 | 0.007233678 | 2.71E-08 | Eastern |
| UG_830 | 0.004541307 | 0.153855704 | 0.82864632 | 0.012956668 | 1.21E-14 | Northern |
| UG_838 | 0.004619662 | 0.14860268 | 0.83263382 | 0.014143837 | 1.44E-14 | Northern |
| UG_846 | 0.001422129 | 0.010639005 | 0.701320004 | 0.286618861 | 1.61E-15 | Northern |
| UG_766 | 0.022109695 | 0.5359274 | 0.43890228 | 0.003060623 | 1.03E-09 | Eastern |
| UG_774 | 0.005085763 | 0.525791867 | 0.468443516 | 0.000678854 | 2.07E-14 | Eastern |
| UG_782 | 0.102585997 | 0.556725008 | 0.33043864 | 0.010057426 | 0.00019293 | Eastern |
| UG_790 | 0.027303069 | 0.460910777 | 0.505438502 | 0.006347649 | 3.65E-09 | Eastern |
| UG_798 | 0.006325018 | 0.131360617 | 0.837151667 | 0.025162698 | 1.76E-13 | Northern |
| UG_806 | 0.004433425 | 0.131486717 | 0.846884234 | 0.017195624 | 1.30E-14 | Northern |
| UG_814 | 0.003547331 | 0.116997266 | 0.862529993 | 0.01692541 | 3.09E-15 | Northern |
| UG_822 | 0.004511324 | 0.039794178 | 0.821102536 | 0.134591961 | 2.11E-13 | Northern |
| UG_759 | 0.012889285 | 0.47175031 | 0.512668639 | 0.002691765 | 1.47E-11 | Eastern |
| UG_831 | 8.79E-05 | 0.000101176 | 0.066370219 | 0.933440668 | 9.27E-16 | Northern |
| UG_839 | 0.000855828 | 0.003716806 | 0.467856919 | 0.527570447 | 2.01E-15 | Northern |
| UG_767 | 0.023323101 | 0.458029825 | 0.513133384 | 0.005513688 | 1.12E-09 | Eastern |
| UG_775 | 0.005498279 | 0.436959776 | 0.556188285 | 0.00135366 | 2.67E-14 | Eastern |
| UG_783 | 0.036341753 | 0.470527481 | 0.485239198 | 0.007891536 | 3.24E-08 | Eastern |
| UG_791 | 0.025663787 | 0.448975998 | 0.518889073 | 0.00647114 | 2.22E-09 | Eastern |
| UG_799 | 0.004154944 | 0.110459897 | 0.863025982 | 0.022359177 | 1.09E-14 | Northern |
| UG_807 | 0.004519581 | 0.124313139 | 0.85161104 | 0.01955624 | 1.64E-14 | Northern |
| UG_815 | 0.004068827 | 0.113930914 | 0.86137319 | 0.02062707 | 8.83E-15 | Northern |
| UG_823 | 0.004121053 | 0.128066498 | 0.851086487 | 0.016725962 | 7.93E-15 | Northern |
| UG_760 | 0.011172828 | 0.517503272 | 0.469632215 | 0.001691685 | 6.15E-12 | Eastern |
| UG_832 | 0.004856923 | 0.129929647 | 0.845808119 | 0.019405311 | 2.58E-14 | Northern |
| UG_840 | 0.005590179 | 0.106060109 | 0.855367869 | 0.032981843 | 1.04E-13 | Northern |
| UG_768 | 0.026201002 | 0.398524731 | 0.565881426 | 0.009392838 | 2.33E-09 | Eastern |

| | | | | | | |
|---|---|---|---|---|---|---|
| UG_776 | 0.036104792 | 0.485105148 | 0.471697228 | 0.007092799 | 3.25E-08 | Eastern |
| UG_784 | 0.004836698 | 0.529870906 | 0.464667546 | 0.000624849 | 1.46E-14 | Eastern |
| UG_792 | 0.004999887 | 0.485570036 | 0.50855516 | 0.000874917 | 1.55E-14 | Eastern |
| UG_800 | 0.005027084 | 0.161998885 | 0.81993343 | 0.013040602 | 2.35E-14 | Northern |
| UG_808 | 0.057480315 | 0.393757868 | 0.52810236 | 0.020658546 | 9.11E-07 | Northern |
| UG_816 | 0.004371557 | 0.089369124 | 0.871440606 | 0.034818713 | 2.37E-14 | Northern |
| UG_824 | 0.004649963 | 0.039740574 | 0.817143451 | 0.138466011 | 2.68E-13 | Northern |
| UG_847 | 0.003571779 | 0.116895167 | 0.862453415 | 0.017079639 | 3.25E-15 | Northern |
| UG_919 | 0.004339418 | 0.123846418 | 0.852957008 | 0.018857156 | 1.22E-14 | Northern |
| UG_927 | 9.58E-05 | 0.000111902 | 0.069575936 | 0.930216367 | 1.14E-15 | Northern |
| UG_935 | 0.006347547 | 0.095286129 | 0.852744223 | 0.045622101 | 3.32E-13 | Northern |
| UG_855 | 0.002691643 | 0.051614749 | 0.890960104 | 0.054733505 | 2.10E-15 | Northern |
| UG_863 | 0.003589064 | 0.117804625 | 0.861689505 | 0.016916806 | 3.33E-15 | Northern |
| UG_871 | 0.004540683 | 0.136191083 | 0.842788948 | 0.016479285 | 1.46E-14 | Northern |
| UG_879 | 0.000814359 | 0.00292501 | 0.40252035 | 0.593740281 | 4.32E-15 | Northern |
| UG_887 | 0.004188392 | 0.036255814 | 0.815448595 | 0.1441072 | 1.55E-13 | Northern |
| UG_895 | 0.005023685 | 0.130263325 | 0.844693323 | 0.020019666 | 3.29E-14 | Northern |
| UG_903 | 0.008682046 | 0.141429661 | 0.819436989 | 0.030451304 | 1.61E-12 | Northern |
| UG_911 | 0.003486456 | 0.094008816 | 0.877484838 | 0.025019891 | 4.04E-15 | Northern |
| UG_848 | 0.005910623 | 0.148911354 | 0.826818436 | 0.018359587 | 8.72E-14 | Northern |
| UG_920 | 0.003573363 | 0.134834092 | 0.848623007 | 0.012969539 | 2.58E-15 | Northern |
| UG_928 | 0.000671402 | 0.003369508 | 0.486836375 | 0.509122715 | 3.66E-16 | Northern |
| UG_936 | 0.003981192 | 0.130792002 | 0.849755907 | 0.015470898 | 5.95E-15 | Northern |
| UG_856 | 0.004303002 | 0.132385906 | 0.846877908 | 0.016433184 | 1.03E-14 | Northern |
| UG_864 | 0.000333299 | 0.000936049 | 0.252111602 | 0.746619049 | 5.24E-16 | Northern |
| UG_872 | 0.004586441 | 0.13569121 | 0.842944554 | 0.016777794 | 1.58E-14 | Northern |
| UG_880 | 0.003148867 | 0.032379298 | 0.832515916 | 0.131955919 | 2.31E-14 | Northern |
| UG_888 | 0.003758617 | 0.151538139 | 0.833828019 | 0.010875225 | 3.11E-15 | Northern |
| UG_896 | 0.000868811 | 0.002464175 | 0.340548902 | 0.656118112 | 1.96E-14 | Northern |

| | | | | | | |
|---|---|---|---|---|---|---|
| UG_904 | 0.004000799 | 0.124581197 | 0.85433277 | 0.017085235 | 6.69E-15 | Northern |
| UG_912 | 0.004199093 | 0.120942845 | 0.855809757 | 0.019048306 | 1.00E-14 | Northern |
| UG_849 | 0.00339628 | 0.104582231 | 0.872056338 | 0.019965151 | 2.74E-15 | Northern |
| UG_921 | 0.003022404 | 0.607782872 | 0.388977275 | 0.000217449 | 7.64E-16 | Northern |
| UG_929 | 0.002756476 | 0.620679683 | 0.376384625 | 0.000179216 | 4.30E-16 | Northern |
| UG_937 | 0.000595212 | 0.002224239 | 0.380821477 | 0.616359073 | 1.01E-15 | Northern |
| UG_857 | 0.004944358 | 0.13476633 | 0.841858217 | 0.018431095 | 2.77E-14 | Northern |
| UG_865 | 0.004455302 | 0.039730196 | 0.822381228 | 0.133433274 | 1.92E-13 | Northern |
| UG_873 | 0.001179376 | 0.004169429 | 0.438501162 | 0.556150033 | 2.07E-14 | Northern |
| UG_881 | 0.004451636 | 0.119533351 | 0.855279613 | 0.020735401 | 1.57E-14 | Northern |
| UG_889 | 0.003783316 | 0.119168722 | 0.859531798 | 0.017516164 | 4.79E-15 | Northern |
| UG_897 | 0.00404532 | 0.125738348 | 0.853231014 | 0.016985319 | 7.14E-15 | Northern |
| UG_905 | 0.004367967 | 0.129646242 | 0.848595403 | 0.017390388 | 1.19E-14 | Northern |
| UG_913 | 0.003641067 | 0.1174754 | 0.861610366 | 0.017273167 | 3.71E-15 | Northern |
| UG_850 | 0.00088694 | 0.004041295 | 0.487629243 | 0.507442523 | 1.84E-15 | Northern |
| UG_922 | 0.004647273 | 0.160717647 | 0.822464725 | 0.012170356 | 1.34E-14 | Northern |
| UG_930 | 0.003991384 | 0.151452777 | 0.832932704 | 0.011623135 | 4.82E-15 | Northern |
| UG_938 | 0.004488658 | 0.156442479 | 0.826695043 | 0.012373821 | 1.08E-14 | Northern |
| UG_858 | 0.004234627 | 0.14966043 | 0.833412946 | 0.012691996 | 7.56E-15 | Northern |
| UG_866 | 0.003545471 | 0.107244348 | 0.869265135 | 0.019945047 | 3.59E-15 | Northern |
| UG_874 | 0.007778001 | 0.126138905 | 0.832352549 | 0.033730545 | 8.73E-13 | Northern |
| UG_882 | 0.004014768 | 0.11657579 | 0.859940529 | 0.019468913 | 7.69E-15 | Northern |
| UG_890 | 0.039847979 | 0.253112924 | 0.66397187 | 0.043067162 | 6.52E-08 | Northern |
| UG_898 | 0.003669754 | 0.114930746 | 0.863238442 | 0.018161057 | 4.08E-15 | Northern |
| UG_906 | 0.005943977 | 0.197384289 | 0.786267741 | 0.010403993 | 6.13E-14 | Northern |
| UG_914 | 0.003049501 | 0.582178515 | 0.414509381 | 0.000262604 | 6.89E-16 | Northern |
| UG_851 | 0.00263215 | 0.041378712 | 0.879132913 | 0.076856226 | 3.02E-15 | Northern |
| UG_923 | 0.000597757 | 0.002779715 | 0.448715398 | 0.54790713 | 3.33E-16 | Northern |
| UG_931 | 0.003074466 | 0.596103808 | 0.400581198 | 0.000240528 | 7.99E-16 | Northern |

| | | | | | | |
|---|---|---|---|---|---|---|
| UG_939 | 0.000574989 | 0.0021648 | 0.379381365 | 0.617878846 | 8.48E-16 | Northern |
| UG_859 | 0.003824026 | 0.135488977 | 0.846859521 | 0.013827476 | 4.19E-15 | Northern |
| UG_867 | 0.004454028 | 0.039934191 | 0.823196252 | 0.132415529 | 1.89E-13 | Northern |
| UG_875 | 0.004676594 | 0.041960279 | 0.824716994 | 0.128646132 | 2.40E-13 | Northern |
| UG_883 | 0.000702299 | 0.003040856 | 0.444281863 | 0.551974981 | 9.15E-16 | Northern |
| UG_891 | 0.003184378 | 0.117660592 | 0.864255348 | 0.014899683 | 1.39E-15 | Northern |
| UG_899 | 0.004894246 | 0.16721874 | 0.816007009 | 0.011880005 | 1.85E-14 | Northern |
| UG_907 | 0.000116787 | 0.000142075 | 0.07800561 | 0.921735528 | 1.80E-15 | Northern |
| UG_915 | 0.003575149 | 0.11950052 | 0.860532295 | 0.016392036 | 3.15E-15 | Northern |
| UG_852 | 0.058384896 | 0.394903356 | 0.525933892 | 0.020776825 | 1.03E-06 | Northern |
| UG_924 | 0.003543311 | 0.109794319 | 0.867592631 | 0.019069739 | 3.42E-15 | Northern |
| UG_932 | 0.002796882 | 0.60988696 | 0.387119531 | 0.000196627 | 4.42E-16 | Northern |
| UG_940 | 8.71E-05 | 9.96E-05 | 0.065713015 | 0.934100377 | 9.29E-16 | Northern |
| UG_860 | 0.004688254 | 0.116785607 | 0.855626486 | 0.022899653 | 2.39E-14 | Northern |
| UG_868 | 0.002964299 | 0.022478055 | 0.769331539 | 0.205226108 | 4.48E-14 | Northern |
| UG_876 | 0.000793971 | 0.002238851 | 0.330141698 | 0.66682548 | 1.43E-14 | Northern |
| UG_884 | 0.007691879 | 0.126758323 | 0.832510831 | 0.033038967 | 7.97E-13 | Northern |
| UG_892 | 0.003821898 | 0.103822411 | 0.869391467 | 0.022964223 | 6.60E-15 | Northern |
| UG_900 | 0.004338237 | 0.131203578 | 0.847589219 | 0.016868966 | 1.11E-14 | Northern |
| UG_908 | 0.000934126 | 0.005111179 | 0.554370673 | 0.439584023 | 9.61E-16 | Northern |
| UG_916 | 0.004374353 | 0.142165402 | 0.838907361 | 0.014552884 | 1.04E-14 | Northern |
| UG_853 | 0.004583554 | 0.161997128 | 0.821620279 | 0.011799039 | 1.20E-14 | Northern |
| UG_925 | 0.000570969 | 0.002028655 | 0.361778771 | 0.635621606 | 1.12E-15 | Northern |
| UG_933 | 0.003519434 | 0.117070462 | 0.862648233 | 0.01676187 | 2.91E-15 | Northern |
| UG_861 | 0.058352729 | 0.398912148 | 0.522546752 | 0.02018734 | 1.03E-06 | Northern |
| UG_869 | 0.007526492 | 0.126625968 | 0.833482593 | 0.032364947 | 6.80E-13 | Northern |
| UG_877 | 0.004416307 | 0.039816758 | 0.823777979 | 0.131988957 | 1.78E-13 | Northern |
| UG_885 | 0.004367321 | 0.146076508 | 0.835786089 | 0.013770082 | 9.82E-15 | Northern |
| UG_893 | 0.003949388 | 0.097440713 | 0.871869969 | 0.026739931 | 9.45E-15 | Northern |

| | | | | | | |
|---|---|---|---|---|---|---|
| UG_901 | 0.004292538 | 0.118348358 | 0.857034328 | 0.020324776 | 1.22E-14 | Northern |
| UG_909 | 0.00312559 | 0.582025357 | 0.414578904 | 0.000270149 | 8.22E-16 | Northern |
| UG_917 | 0.003092565 | 0.588827406 | 0.40782534 | 0.000254689 | 7.95E-16 | Northern |
| UG_854 | 0.002460648 | 0.021645131 | 0.790057112 | 0.185837109 | 1.14E-14 | Northern |
| UG_926 | 0.00344238 | 0.121710585 | 0.8596554 | 0.015191634 | 2.32E-15 | Northern |
| UG_934 | 0.005050372 | 0.147977979 | 0.831269861 | 0.015701788 | 2.78E-14 | Northern |
| UG_862 | 0.006361285 | 0.095376759 | 0.852616349 | 0.045645608 | 3.36E-13 | Northern |
| UG_870 | 0.003554373 | 0.116918788 | 0.862543474 | 0.016983365 | 3.14E-15 | Northern |
| UG_878 | 0.004488149 | 0.039784904 | 0.821696656 | 0.13403029 | 2.02E-13 | Northern |
| UG_886 | 0.003585704 | 0.04176405 | 0.852896389 | 0.101753857 | 3.11E-14 | Northern |
| UG_894 | 0.001048783 | 0.003510525 | 0.408512059 | 0.586928633 | 1.71E-14 | Northern |
| UG_902 | 0.001269391 | 0.009170622 | 0.679099257 | 0.31046073 | 1.11E-15 | Northern |
| UG_910 | 0.000106915 | 0.000131337 | 0.076274282 | 0.923487467 | 1.23E-15 | Northern |
| UG_918 | 0.000100565 | 0.000121346 | 0.073226503 | 0.926551585 | 1.11E-15 | Northern |
| UG_941 | 0.004438582 | 0.12973443 | 0.848157404 | 0.017669584 | 1.34E-14 | Northern |
| UG_1013 | 0.072965941 | 0.581266567 | 0.339110419 | 0.006643482 | 1.36E-05 | Eastern |
| UG_1021 | 0.004490904 | 0.03905834 | 0.818746957 | 0.137703799 | 2.15E-13 | Northern |
| UG_1029 | 0.004232118 | 0.136704772 | 0.84389983 | 0.015163281 | 8.66E-15 | Northern |
| UG_949 | 0.002983837 | 0.616927283 | 0.379887941 | 0.00020094 | 7.43E-16 | Northern |
| UG_957 | 0.004985407 | 0.148274106 | 0.831317202 | 0.015423285 | 2.52E-14 | Northern |
| UG_965 | 0.003822385 | 0.573752424 | 0.422067935 | 0.000357257 | 3.36E-15 | Eastern |
| UG_973 | 0.067330239 | 0.699997147 | 0.230310154 | 0.002337524 | 2.49E-05 | Eastern |
| UG_981 | 0.002863315 | 0.583680165 | 0.413214137 | 0.000242383 | 4.40E-16 | Eastern |
| UG_989 | 0.005961595 | 0.474031487 | 0.518858927 | 0.001147991 | 5.34E-14 | Eastern |
| UG_997 | 0.018720777 | 0.613181907 | 0.366599473 | 0.001497843 | 4.87E-10 | Eastern |
| UG_1005 | 0.103269143 | 0.605768544 | 0.283821802 | 0.006825611 | 0.0003149 | Eastern |
| UG_942 | 0.002689267 | 0.545152005 | 0.451864259 | 0.000294469 | 2.25E-16 | Northern |
| UG_1014 | 0.004836263 | 0.638606722 | 0.356264402 | 0.000292612 | 2.92E-14 | Eastern |
| UG_1022 | 0.000473437 | 0.001733833 | 0.351319497 | 0.646473233 | 4.58E-16 | Northern |

| | | | | | | |
|---|---|---|---|---|---|---|
| UG_1030 | 0.003478797 | 0.106839199 | 0.869999809 | 0.019682195 | 3.14E-15 | Northern |
| UG_950 | 0.003664763 | 0.151106119 | 0.834588823 | 0.010640295 | 2.60E-15 | Northern |
| UG_958 | 0.004533163 | 0.136841471 | 0.842327532 | 0.016297834 | 1.43E-14 | Northern |
| UG_966 | 0.041779465 | 0.68414782 | 0.272200606 | 0.001871684 | 4.24E-07 | Eastern |
| UG_974 | 0.003498834 | 0.583650492 | 0.412548075 | 0.000302599 | 1.88E-15 | Eastern |
| UG_982 | 0.003064768 | 0.603628193 | 0.393079661 | 0.000227377 | 8.21E-16 | Eastern |
| UG_990 | 0.005511761 | 0.527228497 | 0.466525367 | 0.000734376 | 3.73E-14 | Eastern |
| UG_998 | 0.003202835 | 0.615057286 | 0.381519669 | 0.000220211 | 1.22E-15 | Eastern |
| UG_1006 | 0.102532852 | 0.606570707 | 0.283842003 | 0.006756291 | 0.000298147 | Eastern |
| UG_943 | 0.001028623 | 0.005895425 | 0.581104593 | 0.411971359 | 1.17E-15 | Northern |
| UG_1015 | 0.006628094 | 0.606324949 | 0.386526312 | 0.000520645 | 2.26E-13 | Eastern |
| UG_1023 | 0.004868488 | 0.168554572 | 0.814954424 | 0.011622516 | 1.76E-14 | Northern |
| UG_1031 | 0.003541743 | 0.136191807 | 0.847671215 | 0.012595235 | 2.38E-15 | Northern |
| UG_951 | 0.011800295 | 0.151762243 | 0.799938212 | 0.03649925 | 1.40E-11 | Northern |
| UG_959 | 0.004420615 | 0.138853433 | 0.841309858 | 0.015416093 | 1.16E-14 | Northern |
| UG_967 | 0.003205429 | 0.613858497 | 0.382713783 | 0.000222291 | 1.22E-15 | Eastern |
| UG_975 | 0.002810831 | 0.598107986 | 0.398866402 | 0.00021478 | 4.23E-16 | Eastern |
| UG_983 | 0.003301369 | 0.625492744 | 0.370994525 | 0.000211363 | 1.65E-15 | Eastern |
| UG_991 | 0.008943406 | 0.613568798 | 0.376803572 | 0.000684224 | 2.12E-12 | Eastern |
| UG_999 | 0.004218081 | 0.132849314 | 0.846957477 | 0.015975127 | 8.86E-15 | Eastern |
| UG_1007 | 0.088016569 | 0.702452625 | 0.206636565 | 0.002628422 | 0.000265818 | Eastern |
| UG_944 | 0.004532402 | 0.139851409 | 0.839999785 | 0.015616404 | 1.38E-14 | Northern |
| UG_1016 | 0.005357873 | 0.540333818 | 0.453657064 | 0.000651244 | 3.24E-14 | Eastern |
| UG_1024 | 0.003435562 | 0.071825149 | 0.884812374 | 0.039926915 | 6.22E-15 | Northern |
| UG_1032 | 0.005343788 | 0.10075945 | 0.85933955 | 0.034557211 | 8.25E-14 | Northern |
| UG_952 | 0.000618793 | 0.002254458 | 0.377436786 | 0.619689963 | 1.35E-15 | Northern |
| UG_960 | 0.009629994 | 0.193615639 | 0.778578437 | 0.01817593 | 2.15E-12 | Northern |
| UG_968 | 0.004861744 | 0.6101869 | 0.38459038 | 0.000360975 | 2.44E-14 | Eastern |
| UG_976 | 0.010362119 | 0.684215265 | 0.304949982 | 0.000472634 | 1.15E-11 | Eastern |

| | | | | | | |
|---|---|---|---|---|---|---|
| UG_984 | 0.003389762 | 0.105980149 | 0.871195992 | 0.019434098 | 2.64E-15 | Eastern |
| UG_992 | 0.003282158 | 0.614928751 | 0.381562642 | 0.00022645 | 1.46E-15 | Eastern |
| UG_1000 | 0.003066866 | 0.634186888 | 0.362563236 | 0.00018301 | 1.03E-15 | Eastern |
| UG_1008 | 0.007191492 | 0.658688407 | 0.333731477 | 0.000388624 | 6.22E-13 | Eastern |
| UG_945 | 0.009172765 | 0.187350324 | 0.785000665 | 0.018476247 | 1.57E-12 | Northern |
| UG_1017 | 0.011548753 | 0.572814392 | 0.414437449 | 0.001199407 | 1.05E-11 | Eastern |
| UG_1025 | 0.005060124 | 0.151835744 | 0.828152085 | 0.014952047 | 2.71E-14 | Northern |
| UG_1033 | 0.001710727 | 0.017259283 | 0.7952988 | 0.18573119 | 1.30E-15 | Northern |
| UG_953 | 0.003080958 | 0.5913998 | 0.405270109 | 0.000249133 | 7.86E-16 | Northern |
| UG_961 | 0.009216404 | 0.187835783 | 0.784476998 | 0.018470814 | 1.62E-12 | Northern |
| UG_969 | 0.003457689 | 0.621889657 | 0.374424403 | 0.000228251 | 2.24E-15 | Eastern |
| UG_977 | 0.003270619 | 0.596715259 | 0.39975767 | 0.000256452 | 1.25E-15 | Eastern |
| UG_985 | 0.007665238 | 0.576590686 | 0.414993103 | 0.000750973 | 5.37E-13 | Eastern |
| UG_993 | 0.092172873 | 0.580053849 | 0.319806286 | 0.007873542 | 9.34E-05 | Eastern |
| UG_1001 | 0.087918264 | 0.706250952 | 0.203023708 | 0.002526913 | 0.000280163 | Eastern |
| UG_1009 | 0.003211754 | 0.584073521 | 0.412440249 | 0.000274476 | 1.01E-15 | Eastern |
| UG_946 | 0.003266796 | 0.122594299 | 0.859983333 | 0.014155572 | 1.56E-15 | Northern |
| UG_1018 | 0.042279143 | 0.661413317 | 0.294025874 | 0.002281303 | 3.63E-07 | Eastern |
| UG_1026 | 0.005402254 | 0.101728751 | 0.858521795 | 0.0343472 | 8.78E-14 | Northern |
| UG_1034 | 0.005093539 | 0.109417929 | 0.857244184 | 0.028244348 | 4.95E-14 | Northern |
| UG_954 | 0.00163201 | 0.01960891 | 0.828180752 | 0.150578327 | 6.00E-16 | Northern |
| UG_962 | 0.003591682 | 0.103294945 | 0.87141862 | 0.021694753 | 4.22E-15 | Northern |
| UG_970 | 0.010852225 | 0.643998944 | 0.34447412 | 0.00067471 | 1.11E-11 | Eastern |
| UG_978 | 0.003119271 | 0.60050472 | 0.396139016 | 0.000236993 | 9.13E-16 | Eastern |
| UG_986 | 0.003151833 | 0.601692316 | 0.39491811 | 0.000237741 | 9.93E-16 | Eastern |
| UG_994 | 0.002351858 | 0.597105462 | 0.400365165 | 0.000177514 | 1.16E-16 | Eastern |
| UG_1002 | 0.011702993 | 0.70769374 | 0.280158482 | 0.000444785 | 3.65E-11 | Eastern |
| UG_1010 | 0.004103794 | 0.585321869 | 0.410217689 | 0.000356648 | 6.04E-15 | Eastern |
| UG_947 | 0.004328441 | 0.122463217 | 0.853993946 | 0.019214396 | 1.23E-14 | Northern |

| | | | | | | |
|---|---|---|---|---|---|---|
| UG_1019 | 0.003434113 | 0.617718195 | 0.378614314 | 0.000233378 | 2.07E-15 | Eastern |
| UG_1027 | 0.004165416 | 0.093068606 | 0.872004849 | 0.030761129 | 1.53E-14 | Northern |
| UG_955 | 0.005031882 | 0.159701746 | 0.821831492 | 0.01343488 | 2.42E-14 | Northern |
| UG_963 | 0.003362801 | 0.022182641 | 0.74493371 | 0.229520848 | 1.30E-13 | Northern |
| UG_971 | 0.003083729 | 0.607262779 | 0.389430342 | 0.00022315 | 8.80E-16 | Eastern |
| UG_979 | 0.008163831 | 0.530277914 | 0.460453435 | 0.00110482 | 6.60E-13 | Eastern |
| UG_987 | 0.002670778 | 0.632163073 | 0.365006779 | 0.00015937 | 3.74E-16 | Eastern |
| UG_995 | 0.020287223 | 0.710411123 | 0.268538166 | 0.000763485 | 2.27E-09 | Eastern |
| UG_1003 | 0.003808705 | 0.619460149 | 0.376472769 | 0.000258377 | 4.44E-15 | Eastern |
| UG_1011 | 0.101766108 | 0.604025274 | 0.287062234 | 0.006874241 | 0.000272143 | Eastern |
| UG_948 | 0.004569573 | 0.128292318 | 0.848510977 | 0.018627132 | 1.69E-14 | Northern |
| UG_1020 | 0.002799845 | 0.587554523 | 0.40941546 | 0.000230172 | 3.84E-16 | Eastern |
| UG_1028 | 0.000455487 | 0.001713216 | 0.354961194 | 0.642870103 | 3.41E-16 | Northern |
| UG_956 | 0.003125642 | 0.600655586 | 0.395981494 | 0.000237278 | 9.28E-16 | Northern |
| UG_964 | 0.045072653 | 0.652024016 | 0.300302181 | 0.002600608 | 5.44E-07 | Eastern |
| UG_972 | 0.013073995 | 0.587864218 | 0.397829558 | 0.001232229 | 2.87E-11 | Eastern |
| UG_980 | 0.003780334 | 0.653363434 | 0.342655947 | 0.000200285 | 5.52E-15 | Eastern |
| UG_988 | 0.015643506 | 0.570924814 | 0.411755008 | 0.001676672 | 9.67E-11 | Eastern |
| UG_996 | 0.003143345 | 0.608509383 | 0.388121343 | 0.000225929 | 1.02E-15 | Eastern |
| UG_1004 | 0.099645212 | 0.603353867 | 0.289938737 | 0.006837168 | 0.000225015 | Eastern |
| UG_1012 | 0.021588582 | 0.698833464 | 0.278683173 | 0.000894779 | 3.16E-09 | Eastern |
| UG_1035 | 0.000779725 | 0.004516857 | 0.552063713 | 0.442639706 | 3.46E-16 | Northern |
| UG_1107 | 0.004029862 | 0.126074964 | 0.853066609 | 0.016828564 | 6.92E-15 | Northern |
| UG_1115 | 0.003497172 | 0.07060552 | 0.883993776 | 0.041903533 | 7.36E-15 | Northern |
| UG_1123 | 0.003605282 | 0.131181782 | 0.851396547 | 0.013816389 | 2.87E-15 | Northern |
| UG_1043 | 0.007624974 | 0.123716093 | 0.834396884 | 0.034262049 | 7.81E-13 | Northern |
| UG_1051 | 0.006436353 | 0.093992238 | 0.852166116 | 0.047405292 | 3.78E-13 | Northern |
| UG_1059 | 0.001076637 | 0.007987207 | 0.669660355 | 0.321275802 | 4.79E-16 | Northern |
| UG_1067 | 0.005182423 | 0.146482179 | 0.831864592 | 0.016470807 | 3.42E-14 | Northern |

| | | | | | | |
|---|---|---|---|---|---|---|
| UG_1075 | 0.004621651 | 0.049190025 | 0.845959447 | 0.100228877 | 1.42E-13 | Northern |
| UG_1083 | 0.005024344 | 0.128031901 | 0.846245649 | 0.020698106 | 3.39E-14 | Northern |
| UG_1091 | 0.004073852 | 0.134032222 | 0.846770711 | 0.015123215 | 6.77E-15 | Northern |
| UG_1099 | 0.004553674 | 0.122189378 | 0.852883191 | 0.020373757 | 1.78E-14 | Northern |
| UG_1036 | 0.003955314 | 0.123478064 | 0.855399676 | 0.017166945 | 6.25E-15 | Northern |
| UG_1108 | 0.004357284 | 0.11997954 | 0.855540576 | 0.0201226 | 1.33E-14 | Northern |
| UG_1116 | 0.00179919 | 0.022173682 | 0.837932539 | 0.138094589 | 8.90E-16 | Northern |
| UG_1124 | 0.004090789 | 0.12187669 | 0.855781381 | 0.018251141 | 8.17E-15 | Northern |
| UG_1044 | 0.004543488 | 0.040218047 | 0.821860185 | 0.13337828 | 2.16E-13 | Northern |
| UG_1052 | 0.005051336 | 0.084957355 | 0.865745269 | 0.044246039 | 7.66E-14 | Northern |
| UG_1060 | 0.003497188 | 0.103835628 | 0.871787309 | 0.020879875 | 3.44E-15 | Northern |
| UG_1068 | 0.000745125 | 0.003785621 | 0.503075775 | 0.49239348 | 5.30E-16 | Northern |
| UG_1076 | 0.000843825 | 0.0034059 | 0.442755505 | 0.552994771 | 2.74E-15 | Northern |
| UG_1084 | 0.004354933 | 0.134809929 | 0.844765005 | 0.016070134 | 1.09E-14 | Northern |
| UG_1092 | 0.005450963 | 0.162075746 | 0.818254866 | 0.014218426 | 4.24E-14 | Northern |
| UG_1100 | 0.004263995 | 0.121066477 | 0.85534302 | 0.019326508 | 1.12E-14 | Northern |
| UG_1037 | 0.003822273 | 0.118890625 | 0.859497545 | 0.017789556 | 5.19E-15 | Northern |
| UG_1109 | 0.000612745 | 0.002369602 | 0.394147775 | 0.602869878 | 9.47E-16 | Northern |
| UG_1117 | 0.004645379 | 0.132835545 | 0.844792194 | 0.017726882 | 1.79E-14 | Northern |
| UG_1125 | 0.005512318 | 0.136919434 | 0.837490323 | 0.020077925 | 5.98E-14 | Northern |
| UG_1045 | 0.004209278 | 0.139712645 | 0.841632205 | 0.014445872 | 8.04E-15 | Northern |
| UG_1053 | 0.003876794 | 0.080110203 | 0.878666819 | 0.037346184 | 1.21E-14 | Northern |
| UG_1061 | 0.004074342 | 0.134743873 | 0.846211937 | 0.014969848 | 6.72E-15 | Northern |
| UG_1069 | 0.003237026 | 0.103548082 | 0.873897324 | 0.019317568 | 1.96E-15 | Northern |
| UG_1077 | 0.003168348 | 0.102054101 | 0.875379773 | 0.019397777 | 1.72E-15 | Northern |
| UG_1085 | 0.003612466 | 0.123804263 | 0.857094995 | 0.015488276 | 3.21E-15 | Northern |
| UG_1093 | 0.000678189 | 0.002631334 | 0.40614371 | 0.590546768 | 1.39E-15 | Northern |
| UG_1101 | 0.003458887 | 0.10131778 | 0.87362167 | 0.021601663 | 3.32E-15 | Northern |
| UG_1038 | 0.004202757 | 0.127526165 | 0.851047893 | 0.017223185 | 9.22E-15 | Northern |

| | | | | | | |
|---|---|---|---|---|---|---|
| UG_1110 | 0.003901503 | 0.119524693 | 0.858570916 | 0.018002889 | 5.97E-15 | Northern |
| UG_1118 | 0.003020062 | 0.103223946 | 0.875722807 | 0.018033184 | 1.19E-15 | Northern |
| UG_1126 | 0.000536107 | 0.001920358 | 0.357659181 | 0.639884355 | 8.40E-16 | Northern |
| UG_1046 | 0.003615429 | 0.093130861 | 0.876798251 | 0.026455459 | 5.38E-15 | Northern |
| UG_1054 | 0.002941109 | 0.03911576 | 0.864516112 | 0.093427019 | 8.11E-15 | Northern |
| UG_1062 | 0.004580516 | 0.137358906 | 0.841700249 | 0.01636033 | 1.53E-14 | Northern |
| UG_1070 | 0.004905192 | 0.127713088 | 0.847110741 | 0.020270979 | 2.85E-14 | Northern |
| UG_1086 | 0.00084731 | 0.005136353 | 0.576994478 | 0.417021859 | 4.00E-16 | Northern |
| UG_1094 | 0.000653299 | 0.00349566 | 0.504744311 | 0.491106731 | 2.40E-16 | Northern |
| UG_1102 | 0.004364854 | 0.135884387 | 0.843888403 | 0.015862356 | 1.10E-14 | Northern |
| UG_1039 | 0.00372166 | 0.114698729 | 0.863071288 | 0.018508323 | 4.54E-15 | Northern |
| UG_1111 | 0.000963803 | 0.004729337 | 0.52181743 | 0.47248943 | 1.82E-15 | Northern |
| UG_1119 | 0.003171497 | 0.060068454 | 0.88669575 | 0.050064299 | 5.07E-15 | Northern |
| UG_1127 | 0.003454838 | 0.072333774 | 0.88454541 | 0.039665978 | 6.39E-15 | Northern |
| UG_1047 | 0.005780287 | 0.1190972 | 0.847574691 | 0.027547822 | 1.08E-13 | Northern |
| UG_1055 | 0.004178915 | 0.118644903 | 0.857520709 | 0.019655472 | 1.00E-14 | Northern |
| UG_1063 | 0.004830383 | 0.137911877 | 0.840070751 | 0.017186989 | 2.25E-14 | Northern |
| UG_1071 | 0.004915192 | 0.062445188 | 0.859950913 | 0.072688708 | 1.24E-13 | Northern |
| UG_1079 | 0.004866147 | 0.136268763 | 0.841132381 | 0.017732708 | 2.42E-14 | Northern |
| UG_1087 | 0.003654098 | 0.100801131 | 0.872419376 | 0.023125395 | 5.01E-15 | Northern |
| UG_1095 | 0.003764692 | 0.113069191 | 0.863912841 | 0.019253276 | 5.07E-15 | Northern |
| UG_1103 | 0.00475066 | 0.042111106 | 0.823372251 | 0.129765983 | 2.68E-13 | Northern |
| UG_1040 | 0.00367674 | 0.112430577 | 0.864919868 | 0.018972815 | 4.30E-15 | Northern |
| UG_1112 | 0.003149167 | 0.074564203 | 0.888151363 | 0.034135267 | 3.02E-15 | Northern |
| UG_1120 | 0.004344835 | 0.113477921 | 0.859885077 | 0.022292167 | 1.44E-14 | Northern |
| UG_1128 | 0.003910854 | 0.12214549 | 0.85662777 | 0.017315886 | 5.86E-15 | Northern |
| UG_1048 | 0.004436075 | 0.045042022 | 0.840110618 | 0.110411285 | 1.31E-13 | Northern |
| UG_1056 | 0.000475169 | 0.001707327 | 0.346330379 | 0.651487125 | 5.15E-16 | Northern |
| UG_1064 | 0.003457385 | 0.105768564 | 0.870848795 | 0.019925256 | 3.06E-15 | Northern |

| | | | | | | |
|---|---|---|---|---|---|---|
| UG_1072 | 0.00672197 | 0.100450483 | 0.848801121 | 0.044026426 | 4.57E-13 | Northern |
| UG_1080 | 0.001344207 | 0.010015175 | 0.694381129 | 0.294259489 | 1.27E-15 | Northern |
| UG_1088 | 0.006422126 | 0.14690427 | 0.826072981 | 0.020600624 | 1.64E-13 | Northern |
| UG_1096 | 0.002910117 | 0.039555882 | 0.866634653 | 0.090899348 | 7.26E-15 | Northern |
| UG_1104 | 0.00409819 | 0.120623569 | 0.856626902 | 0.018651339 | 8.43E-15 | Northern |
| UG_1041 | 0.003951804 | 0.107645238 | 0.866158058 | 0.0222449 | 7.89E-15 | Northern |
| UG_1113 | 0.003188685 | 0.094981803 | 0.879509703 | 0.022319809 | 2.06E-15 | Northern |
| UG_1121 | 0.00425614 | 0.07093031 | 0.873972934 | 0.050840616 | 3.15E-14 | Northern |
| UG_1049 | 0.001818945 | 0.015315962 | 0.756782332 | 0.22608276 | 3.20E-15 | Northern |
| UG_1057 | 0.005556681 | 0.086930249 | 0.8606686 | 0.04684447 | 1.49E-13 | Northern |
| UG_1065 | 0.001383579 | 0.009074155 | 0.658392592 | 0.331149674 | 2.39E-15 | Northern |
| UG_1073 | 0.00390555 | 0.140008295 | 0.842820732 | 0.013265424 | 4.64E-15 | Northern |
| UG_1081 | 0.003639778 | 0.045964047 | 0.861443232 | 0.088952943 | 2.71E-14 | Northern |
| UG_1089 | 0.003436671 | 0.050821416 | 0.874078217 | 0.071663696 | 1.37E-14 | Northern |
| UG_1097 | 0.004493327 | 0.039856812 | 0.821833064 | 0.133816797 | 2.03E-13 | Northern |
| UG_1105 | 0.003967529 | 0.115739314 | 0.8608054 | 0.019487758 | 7.14E-15 | Northern |
| UG_1042 | 0.001096543 | 0.006721206 | 0.61064803 | 0.381534221 | 1.15E-15 | Northern |
| UG_1114 | 0.000505791 | 0.00212853 | 0.398979343 | 0.598386335 | 2.84E-16 | Northern |
| UG_1122 | 0.003787763 | 0.085507227 | 0.878260159 | 0.032444851 | 8.95E-15 | Northern |
| UG_1050 | 0.004904031 | 0.155944269 | 0.825446347 | 0.013705353 | 2.07E-14 | Northern |
| UG_1058 | 0.003544565 | 0.049205263 | 0.869431088 | 0.077819085 | 1.87E-14 | Northern |
| UG_1066 | 0.003721948 | 0.113561825 | 0.863853498 | 0.01886273 | 4.62E-15 | Northern |
| UG_1074 | 0.000703498 | 0.003089104 | 0.448950135 | 0.547257262 | 8.58E-16 | Northern |
| UG_1082 | 0.002126345 | 0.022338126 | 0.817610918 | 0.157924611 | 3.23E-15 | Northern |
| UG_1090 | 0.003861944 | 0.128446181 | 0.852186513 | 0.015505362 | 4.91E-15 | Northern |
| UG_1098 | 0.004372264 | 0.139178695 | 0.841284485 | 0.015164556 | 1.07E-14 | Northern |
| UG_1106 | 0.005101914 | 0.11460943 | 0.854337778 | 0.025950879 | 4.61E-14 | Northern |
| UG_1129 | 0.004850254 | 0.137140054 | 0.840556943 | 0.017452749 | 2.34E-14 | Northern |
| UG_1201 | 0.002182988 | 0.011243258 | 0.631795398 | 0.354778356 | 4.80E-14 | North_western |

| | | | | | | |
|---|---|---|---|---|---|---|
| UG_1209 | 0.008149201 | 0.092777157 | 0.837655956 | 0.061417686 | 2.28E-12 | North_western |
| UG_1217 | 0.008502828 | 0.094763199 | 0.834986539 | 0.061747434 | 3.01E-12 | North_western |
| UG_1137 | 0.003998677 | 0.134934809 | 0.846437462 | 0.014629053 | 5.84E-15 | Northern |
| UG_1145 | 0.042082223 | 0.259968463 | 0.655194429 | 0.042754789 | 9.63E-08 | Northern |
| UG_1153 | 0.005000415 | 0.059645978 | 0.855712658 | 0.079640949 | 1.58E-13 | Northern |
| UG_1161 | 0.003299587 | 0.140013462 | 0.84564423 | 0.01104272 | 1.36E-15 | Northern |
| UG_1169 | 0.007798492 | 0.130370185 | 0.830040753 | 0.03179057 | 8.39E-13 | Northern |
| UG_1177 | 0.000592085 | 0.001626357 | 0.295824398 | 0.701957161 | 5.31E-15 | Northern |
| UG_1185 | 8.66E-05 | 0.000103996 | 0.06897827 | 0.93083116 | 6.67E-16 | North_western |
| UG_1193 | 0.00036648 | 0.000866089 | 0.222538287 | 0.776229144 | 2.04E-15 | North_western |
| UG_1130 | 0.003762513 | 0.120424863 | 0.85874575 | 0.017066874 | 4.52E-15 | Northern |
| UG_1202 | 0.000472796 | 0.001669176 | 0.340932406 | 0.656925622 | 5.56E-16 | North_western |
| UG_1210 | 0.000638382 | 0.002530133 | 0.406044288 | 0.590787196 | 9.81E-16 | North_western |
| UG_1218 | 8.78E-05 | 0.000108278 | 0.071206008 | 0.928597865 | 5.98E-16 | North_western |
| UG_1138 | 0.008106195 | 0.131463058 | 0.827853465 | 0.032577281 | 1.10E-12 | Northern |
| UG_1146 | 0.040808309 | 0.256394849 | 0.659973532 | 0.042823233 | 7.72E-08 | Northern |
| UG_1154 | 0.004197623 | 0.130670001 | 0.848722551 | 0.016409825 | 8.78E-15 | Northern |
| UG_1162 | 0.003532358 | 0.148103841 | 0.837723825 | 0.010639977 | 2.05E-15 | Northern |
| UG_1170 | 0.004375342 | 0.124431284 | 0.852339621 | 0.018853753 | 1.29E-14 | Northern |
| UG_1178 | 0.003653991 | 0.125099828 | 0.855876843 | 0.015369337 | 3.42E-15 | Northern |
| UG_1186 | 0.006830812 | 0.080325909 | 0.846842043 | 0.066001235 | 8.32E-13 | North_western |
| UG_1194 | 0.006375046 | 0.076853878 | 0.850333214 | 0.066437862 | 5.46E-13 | North_western |
| UG_1131 | 0.004074312 | 0.116948984 | 0.859317646 | 0.019659058 | 8.52E-15 | Northern |
| UG_1203 | 8.22E-05 | 0.000103996 | 0.071368186 | 0.928445615 | 4.00E-16 | North_western |
| UG_1211 | 0.000701285 | 0.002708806 | 0.408433777 | 0.588156133 | 1.63E-15 | North_western |
| UG_1219 | 0.000111837 | 0.000177142 | 0.100314193 | 0.899396828 | 2.95E-16 | North_western |
| UG_1139 | 0.00411924 | 0.025938719 | 0.748882712 | 0.221059328 | 3.95E-13 | Northern |
| UG_1147 | 0.040720828 | 0.258572297 | 0.658740869 | 0.041965932 | 7.52E-08 | Northern |
| UG_1155 | 0.001544272 | 0.011930752 | 0.718752259 | 0.267772717 | 2.06E-15 | Northern |

| | | | | | | |
|---|---|---|---|---|---|---|
| UG_1163 | 0.004182918 | 0.123141654 | 0.85434812 | 0.018327308 | 9.45E-15 | Northern |
| UG_1171 | 0.003980389 | 0.109957186 | 0.864521137 | 0.021541288 | 8.01E-15 | Northern |
| UG_1179 | 0.004053026 | 0.12421242 | 0.85431013 | 0.017424423 | 7.39E-15 | Northern |
| UG_1187 | 0.006704053 | 0.078674092 | 0.847514086 | 0.067107769 | 7.57E-13 | North_western |
| UG_1195 | 0.000772624 | 0.002094689 | 0.317013981 | 0.680118706 | 1.59E-14 | North_western |
| UG_1132 | 0.007536015 | 0.161027488 | 0.811064605 | 0.020371891 | 4.60E-13 | Northern |
| UG_1204 | 0.001418873 | 0.005463791 | 0.487076701 | 0.506040635 | 2.88E-14 | North_western |
| UG_1212 | 0.000646222 | 0.002481766 | 0.397786488 | 0.599085524 | 1.21E-15 | North_western |
| UG_1220 | 0.010280495 | 0.122603804 | 0.819851302 | 0.0472644 | 7.37E-12 | North_western |
| UG_1140 | 0.00743313 | 0.1249562 | 0.834858149 | 0.032752521 | 6.35E-13 | Northern |
| UG_1148 | 0.003991115 | 0.132580441 | 0.848319855 | 0.015108589 | 5.93E-15 | Northern |
| UG_1156 | 0.000918784 | 0.004383217 | 0.506872038 | 0.487825961 | 1.70E-15 | Northern |
| UG_1164 | 0.003669609 | 0.11149664 | 0.86559959 | 0.019234161 | 4.31E-15 | Northern |
| UG_1172 | 0.005362801 | 0.160384286 | 0.81998446 | 0.014268454 | 3.82E-14 | Northern |
| UG_1180 | 0.005110155 | 0.123476754 | 0.848828584 | 0.022584508 | 4.08E-14 | Northern |
| UG_1188 | 0.005553803 | 0.055038916 | 0.839489037 | 0.099918244 | 4.33E-13 | North_western |
| UG_1196 | 0.000116059 | 0.000147719 | 0.081497388 | 0.918238835 | 1.33E-15 | North_western |
| UG_1133 | 0.005539906 | 0.103741453 | 0.856697089 | 0.034021552 | 1.02E-13 | Northern |
| UG_1205 | 0.000155961 | 0.000265279 | 0.121322812 | 0.878255948 | 6.34E-16 | North_western |
| UG_1213 | 0.00030729 | 0.000637072 | 0.185889009 | 0.813166628 | 2.29E-15 | North_western |
| UG_1221 | 0.010132315 | 0.118169049 | 0.821902144 | 0.049796493 | 7.10E-12 | North_western |
| UG_1141 | 0.005144659 | 0.152396018 | 0.827348926 | 0.015110397 | 3.05E-14 | Northern |
| UG_1149 | 0.00160055 | 0.015133958 | 0.775219321 | 0.208046172 | 1.18E-15 | Northern |
| UG_1157 | 0.002405814 | 0.017167155 | 0.73752658 | 0.24290045 | 2.09E-14 | Northern |
| UG_1165 | 0.007613281 | 0.128161696 | 0.832205317 | 0.032019707 | 7.24E-13 | Northern |
| UG_1173 | 0.005467101 | 0.075020987 | 0.860072323 | 0.059439588 | 1.81E-13 | Northern |
| UG_1181 | 0.004627994 | 0.041304729 | 0.823646109 | 0.130421168 | 2.31E-13 | Northern |
| UG_1189 | 0.00997027 | 0.115404023 | 0.8234935 | 0.051132207 | 6.60E-12 | North_western |
| UG_1197 | 9.59E-05 | 0.000128315 | 0.079998859 | 0.919776975 | 4.86E-16 | North_western |

| | | | | | | |
|---|---|---|---|---|---|---|
| UG_1134 | 0.004262269 | 0.064292044 | 0.871286503 | 0.060159185 | 3.97E-14 | Northern |
| UG_1206 | 0.00361673 | 0.027782928 | 0.785443691 | 0.183156651 | 1.10E-13 | North_western |
| UG_1214 | 0.001176102 | 0.005481874 | 0.52813683 | 0.465205193 | 5.31E-15 | North_western |
| UG_1222 | 0.0008404 | 0.002327699 | 0.330790557 | 0.666041344 | 1.96E-14 | North_western |
| UG_1142 | 0.007695855 | 0.130676821 | 0.830410527 | 0.031216797 | 7.58E-13 | Northern |
| UG_1150 | 0.007455341 | 0.126295716 | 0.834044587 | 0.032204356 | 6.36E-13 | Northern |
| UG_1158 | 0.001512027 | 0.009134853 | 0.642196542 | 0.347156579 | 4.94E-15 | Northern |
| UG_1166 | 0.005247353 | 0.136104265 | 0.839378064 | 0.019270318 | 4.21E-14 | Northern |
| UG_1174 | 9.82E-05 | 0.000114985 | 0.070368063 | 0.929418713 | 1.24E-15 | Northern |
| UG_1182 | 0.000479691 | 0.001683431 | 0.34071298 | 0.657123899 | 6.08E-16 | North_western |
| UG_1190 | 0.000734846 | 0.002915768 | 0.421887283 | 0.574462103 | 1.71E-15 | North_western |
| UG_1198 | 0.000105751 | 0.000136418 | 0.079855544 | 0.919902287 | 8.73E-16 | North_western |
| UG_1135 | 0.003238638 | 0.060432366 | 0.885706682 | 0.050622314 | 5.84E-15 | Northern |
| UG_1207 | 9.48E-05 | 0.000117484 | 0.073620413 | 0.926167268 | 7.62E-16 | North_western |
| UG_1215 | 0.000621277 | 0.001827558 | 0.317973148 | 0.679578017 | 4.37E-15 | North_western |
| UG_1143 | 0.040812352 | 0.251163986 | 0.663264042 | 0.044759541 | 7.90E-08 | Northern |
| UG_1151 | 0.004596154 | 0.166197512 | 0.81797042 | 0.011235914 | 1.18E-14 | Northern |
| UG_1159 | 0.004760448 | 0.127454333 | 0.848076643 | 0.019708576 | 2.30E-14 | Northern |
| UG_1167 | 0.007489955 | 0.125536252 | 0.834246136 | 0.032727657 | 6.66E-13 | Northern |
| UG_1175 | 0.001549658 | 0.009336651 | 0.644034992 | 0.345078699 | 5.57E-15 | Northern |
| UG_1183 | 0.003840268 | 0.024317106 | 0.744999681 | 0.226842945 | 2.79E-13 | North_western |
| UG_1191 | 0.010338427 | 0.122690514 | 0.819500187 | 0.047470872 | 7.67E-12 | North_western |
| UG_1199 | 0.000163663 | 0.00026096 | 0.115763477 | 0.8838119 | 1.13E-15 | North_western |
| UG_1136 | 0.004542826 | 0.040286925 | 0.822138551 | 0.133031699 | 2.15E-13 | Northern |
| UG_1208 | 7.66E-05 | 9.18E-05 | 0.065821023 | 0.93401055 | 4.34E-16 | North_western |
| UG_1216 | 0.003653999 | 0.028249947 | 0.787409889 | 0.180686165 | 1.13E-13 | North_western |
| UG_1144 | 0.040776642 | 0.253311841 | 0.661993443 | 0.043917997 | 7.77E-08 | Northern |
| UG_1152 | 0.005295954 | 0.146080198 | 0.831673638 | 0.01695021 | 4.02E-14 | Northern |
| UG_1160 | 0.00353031 | 0.145054626 | 0.840332417 | 0.011082648 | 2.10E-15 | Northern |

| | | | | | | |
|---|---|---|---|---|---|---|
| UG_1168 | 0.003723142 | 0.101507244 | 0.871481547 | 0.023288067 | 5.67E-15 | Northern |
| UG_1176 | 0.002788174 | 0.013993954 | 0.649913299 | 0.333304572 | 1.57E-13 | Northern |
| UG_1184 | 8.93E-05 | 0.000131929 | 0.086186306 | 0.913592482 | 2.03E-16 | North_western |
| UG_1192 | 0.007718073 | 0.095660943 | 0.841436357 | 0.055184627 | 1.42E-12 | North_western |
| UG_1200 | 0.000126689 | 0.000186189 | 0.097323025 | 0.902364097 | 7.39E-16 | North_western |
| UG_1223 | 0.000116692 | 0.000180158 | 0.099276034 | 0.900427116 | 4.04E-16 | North_western |
| UG_1295 | 0.030673082 | 0.220913267 | 0.70362538 | 0.044788261 | 1.04E-08 | North_western |
| UG_1303 | 0.000488421 | 0.001207736 | 0.254282323 | 0.74402152 | 4.63E-15 | North_western |
| UG_1311 | 0.006693017 | 0.079604579 | 0.84801704 | 0.065685364 | 7.28E-13 | North_western |
| UG_1231 | 0.000249599 | 0.000595989 | 0.197863434 | 0.801290977 | 4.57E-16 | North_western |
| UG_1239 | 0.000160848 | 0.000237652 | 0.10663212 | 0.89296938 | 1.69E-15 | North_western |
| UG_1247 | 0.000159363 | 0.000244642 | 0.110417601 | 0.889178394 | 1.29E-15 | North_western |
| UG_1255 | 0.003080526 | 0.016811889 | 0.685282285 | 0.2948253 | 1.75E-13 | North_western |
| UG_1263 | 0.000118559 | 0.00017496 | 0.095410958 | 0.904295522 | 5.67E-16 | North_western |
| UG_1271 | 0.000152294 | 0.000274098 | 0.127204497 | 0.872369112 | 4.11E-16 | North_western |
| UG_1279 | 0.087299569 | 0.662484887 | 0.246238741 | 0.003838163 | 0.00013864 | North_western |
| UG_1287 | 0.007826489 | 0.089939167 | 0.839978595 | 0.062255749 | 1.80E-12 | North_western |
| UG_1224 | 0.000110748 | 0.000150999 | 0.085918925 | 0.913819328 | 7.28E-16 | North_western |
| UG_1296 | 0.030989237 | 0.221930225 | 0.702277298 | 0.044803229 | 1.12E-08 | North_western |
| UG_1304 | 0.006433858 | 0.122863253 | 0.841624588 | 0.029078301 | 2.25E-13 | North_western |
| UG_1312 | 0.031836111 | 0.230386785 | 0.695226391 | 0.042550699 | 1.31E-08 | North_western |
| UG_1232 | 0.000242995 | 0.000581355 | 0.196454696 | 0.802720954 | 4.09E-16 | North_western |
| UG_1240 | 0.006811775 | 0.08060782 | 0.847148154 | 0.065432251 | 8.08E-13 | North_western |
| UG_1248 | 0.006710847 | 0.079618326 | 0.847832701 | 0.065838126 | 7.42E-13 | North_western |
| UG_1256 | 0.002243413 | 0.011437678 | 0.631548698 | 0.35477021 | 5.64E-14 | North_western |
| UG_1264 | 0.000111513 | 0.000167984 | 0.095274815 | 0.904445688 | 4.00E-16 | North_western |
| UG_1272 | 0.000982041 | 0.003117442 | 0.385449317 | 0.6104512 | 1.74E-14 | North_western |
| UG_1280 | 0.000408994 | 0.001006155 | 0.239149122 | 0.759435729 | 2.44E-15 | North_western |
| UG_1288 | 0.08724834 | 0.66660941 | 0.242300733 | 0.003695993 | 0.000145524 | North_western |

| | | | | | | |
|---|---|---|---|---|---|---|
| UG_1225 | 0.000129088 | 0.000232339 | 0.119964562 | 0.87967401 | 2.25E-16 | North_western |
| UG_1297 | 0.030649309 | 0.216876744 | 0.706004477 | 0.046469459 | 1.06E-08 | North_western |
| UG_1305 | 0.000105871 | 0.000147481 | 0.086381357 | 0.913365291 | 5.41E-16 | North_western |
| UG_1313 | 0.000115944 | 0.000149344 | 0.082463026 | 0.917271687 | 1.23E-15 | North_western |
| UG_1233 | 0.000122971 | 0.000161266 | 0.085804265 | 0.913911497 | 1.35E-15 | North_western |
| UG_1241 | 0.000165401 | 0.000321187 | 0.141076839 | 0.858436573 | 3.48E-16 | North_western |
| UG_1249 | 0.006620984 | 0.078142881 | 0.848184352 | 0.067051782 | 7.00E-13 | North_western |
| UG_1257 | 0.017149222 | 0.103582583 | 0.77630107 | 0.102967124 | 5.26E-10 | North_western |
| UG_1265 | 0.000127861 | 0.000195342 | 0.101528819 | 0.898147978 | 6.00E-16 | North_western |
| UG_1273 | 0.000225943 | 0.000512188 | 0.182243093 | 0.817018776 | 4.31E-16 | North_western |
| UG_1281 | 0.000349669 | 0.000958045 | 0.250429432 | 0.748262854 | 7.24E-16 | North_western |
| UG_1289 | 0.000164275 | 0.000285923 | 0.126416585 | 0.873133217 | 6.65E-16 | North_western |
| UG_1226 | 0.000127067 | 0.000173694 | 0.090559605 | 0.909139635 | 1.17E-15 | North_western |
| UG_1298 | 0.001105883 | 0.004502603 | 0.476466696 | 0.517924818 | 7.88E-15 | North_western |
| UG_1306 | 0.00677202 | 0.080316316 | 0.847455378 | 0.065456285 | 7.79E-13 | North_western |
| UG_1314 | 0.001029852 | 0.00435231 | 0.48037441 | 0.514243427 | 4.90E-15 | North_western |
| UG_1234 | 0.000116084 | 0.000181475 | 0.100342717 | 0.899359724 | 3.67E-16 | North_western |
| UG_1242 | 0.001916116 | 0.00918564 | 0.594065883 | 0.394832361 | 3.70E-14 | North_western |
| UG_1250 | 0.000128088 | 0.000180233 | 0.093519682 | 0.906171996 | 1.01E-15 | North_western |
| UG_1258 | 0.087136753 | 0.663552993 | 0.245374378 | 0.003797583 | 0.000138293 | North_western |
| UG_1266 | 0.000159208 | 0.000237628 | 0.107323381 | 0.892279784 | 1.53E-15 | North_western |
| UG_1274 | 9.67E-05 | 0.000128322 | 0.079551894 | 0.920223098 | 5.29E-16 | North_western |
| UG_1282 | 0.000122393 | 0.000192394 | 0.102841306 | 0.896843908 | 4.29E-16 | North_western |
| UG_1290 | 0.000113708 | 0.000169714 | 0.095059063 | 0.904657516 | 4.54E-16 | North_western |
| UG_1227 | 0.000113983 | 0.000154171 | 0.08612245 | 0.913609397 | 8.49E-16 | North_western |
| UG_1299 | 0.001625008 | 0.006551122 | 0.517863436 | 0.473960434 | 4.05E-14 | North_western |
| UG_1307 | 0.002158504 | 0.011575055 | 0.643339244 | 0.342927197 | 3.86E-14 | North_western |
| UG_1315 | 0.000127888 | 0.000207478 | 0.107831056 | 0.891833578 | 4.13E-16 | North_western |
| UG_1235 | 0.001422799 | 0.004598797 | 0.431265023 | 0.562713381 | 6.96E-14 | North_western |

| | | | | | | |
|---|---|---|---|---|---|---|
| UG_1243 | 0.006634711 | 0.080704251 | 0.849032924 | 0.063628114 | 6.60E-13 | North_western |
| UG_1251 | 0.000103807 | 0.000170197 | 0.101098775 | 0.898627221 | 1.82E-16 | North_western |
| UG_1259 | 7.94E-05 | 0.000110875 | 0.077943993 | 0.92186572 | 1.90E-16 | North_western |
| UG_1267 | 9.80E-05 | 0.00013476 | 0.082905806 | 0.916861474 | 4.43E-16 | North_western |
| UG_1275 | 0.000699087 | 0.002704042 | 0.408512966 | 0.588083905 | 1.60E-15 | North_western |
| UG_1283 | 8.97E-05 | 0.000130469 | 0.084959908 | 0.914819923 | 2.27E-16 | North_western |
| UG_1291 | 0.000102846 | 0.000136546 | 0.081398394 | 0.918362214 | 6.59E-16 | North_western |
| UG_1228 | 0.000108836 | 0.000173564 | 0.100015087 | 0.899702512 | 2.57E-16 | North_western |
| UG_1300 | 0.006761748 | 0.079144784 | 0.847095216 | 0.066998251 | 7.97E-13 | North_western |
| UG_1308 | 0.000470273 | 0.001351935 | 0.286620193 | 0.711557599 | 1.70E-15 | North_western |
| UG_1316 | 0.001045558 | 0.004435882 | 0.483395035 | 0.511123525 | 5.12E-15 | North_western |
| UG_1236 | 8.07E-05 | 0.000112388 | 0.0782199 | 0.921587039 | 2.04E-16 | North_western |
| UG_1244 | 0.00659994 | 0.079034564 | 0.84878798 | 0.065577516 | 6.66E-13 | North_western |
| UG_1252 | 0.000191514 | 0.000307554 | 0.123286413 | 0.876214519 | 1.91E-15 | North_western |
| UG_1260 | 0.001353462 | 0.004262196 | 0.417640504 | 0.576743839 | 6.50E-14 | North_western |
| UG_1268 | 0.000165177 | 0.000246364 | 0.108673634 | 0.890914825 | 1.76E-15 | North_western |
| UG_1276 | 0.001240478 | 0.004155526 | 0.427225836 | 0.56737816 | 3.34E-14 | North_western |
| UG_1284 | 0.000136354 | 0.000200208 | 0.099848865 | 0.899814573 | 9.69E-16 | North_western |
| UG_1292 | 0.000243962 | 0.000472296 | 0.160985975 | 0.838297768 | 1.48E-15 | North_western |
| UG_1229 | 0.00264566 | 0.017007023 | 0.717657487 | 0.26268983 | 4.74E-14 | North_western |
| UG_1301 | 0.000230869 | 0.000437431 | 0.154710181 | 0.84462152 | 1.37E-15 | North_western |
| UG_1309 | 0.000104212 | 0.000149741 | 0.088627842 | 0.911118205 | 4.21E-16 | North_western |
| UG_1237 | 9.82E-05 | 0.000132244 | 0.081195811 | 0.918573735 | 5.11E-16 | North_western |
| UG_1245 | 0.000456678 | 0.001116812 | 0.246454123 | 0.751972388 | 3.83E-15 | North_western |
| UG_1253 | 0.084122582 | 0.655654634 | 0.256132708 | 0.003997708 | 9.24E-05 | North_western |
| UG_1261 | 0.006697299 | 0.07830708 | 0.847425152 | 0.067570469 | 7.59E-13 | North_western |
| UG_1269 | 8.47E-05 | 0.00011569 | 0.07803167 | 0.921767926 | 2.75E-16 | North_western |
| UG_1277 | 0.000157689 | 0.000235931 | 0.107213748 | 0.892392632 | 1.46E-15 | North_western |
| UG_1285 | 0.001396898 | 0.005399726 | 0.486519488 | 0.506683888 | 2.65E-14 | North_western |

| | | | | | | |
|---|---|---|---|---|---|---|
| UG_1293 | 0.000160421 | 0.000234975 | 0.105610159 | 0.893994445 | 1.77E-15 | North_western |
| UG_1230 | 6.73E-05 | 8.29E-05 | 0.064450129 | 0.935399702 | 2.33E-16 | North_western |
| UG_1302 | 0.000107822 | 0.000149293 | 0.086425137 | 0.913317748 | 6.00E-16 | North_western |
| UG_1310 | 0.002353714 | 0.010954069 | 0.607601458 | 0.379090759 | 1.02E-13 | North_western |
| UG_1238 | 0.006765354 | 0.078105881 | 0.846591114 | 0.068537651 | 8.24E-13 | North_western |
| UG_1246 | 9.11E-05 | 0.000126085 | 0.081238697 | 0.918544127 | 3.28E-16 | North_western |
| UG_1254 | 0.000105075 | 0.000144064 | 0.084764932 | 0.914985928 | 5.82E-16 | North_western |
| UG_1262 | 0.006794531 | 0.082693182 | 0.848010043 | 0.062502244 | 7.48E-13 | North_western |
| UG_1270 | 0.00357369 | 0.574936779 | 0.421160541 | 0.000328989 | 2.08E-15 | North_western |
| UG_1278 | 0.00018775 | 0.000281678 | 0.114435514 | 0.885095059 | 2.70E-15 | North_western |
| UG_1286 | 0.000502977 | 0.001116879 | 0.233125244 | 0.765254901 | 9.63E-15 | North_western |
| UG_1294 | 0.009941055 | 0.119002702 | 0.822818653 | 0.04823759 | 6.07E-12 | North_western |
| UG_1317 | 0.000101671 | 0.00013519 | 0.08118206 | 0.918581078 | 6.26E-16 | North_western |
| UG_1389 | 0.000917495 | 0.002124623 | 0.292291786 | 0.704666096 | 7.43E-14 | North_western |
| UG_1397 | 0.004225656 | 0.03172479 | 0.789640917 | 0.174408637 | 2.49E-13 | North_western |
| UG_1405 | 0.004111851 | 0.111142499 | 0.862888252 | 0.021857399 | 9.97E-15 | Northern |
| UG_1325 | 0.00013176 | 0.000181385 | 0.092416962 | 0.907269894 | 1.28E-15 | North_western |
| UG_1333 | 0.010032055 | 0.123281385 | 0.821032967 | 0.045653593 | 6.07E-12 | North_western |
| UG_1341 | 0.000196296 | 0.000311703 | 0.122992999 | 0.876499002 | 2.23E-15 | North_western |
| UG_1349 | 0.000943764 | 0.00296808 | 0.378493101 | 0.617595055 | 1.56E-14 | North_western |
| UG_1365 | 0.003827304 | 0.126123049 | 0.854143211 | 0.015906436 | 4.74E-15 | Northern |
| UG_1373 | 0.003039615 | 0.655177946 | 0.341627065 | 0.000155374 | 1.15E-15 | Northern |
| UG_1381 | 0.00341884 | 0.637317956 | 0.359061505 | 0.000201699 | 2.33E-15 | North_western |
| UG_1318 | 0.000720649 | 0.00288504 | 0.42249616 | 0.57389815 | 1.51E-15 | North_western |
| UG_1390 | 9.33E-05 | 0.000122292 | 0.077536052 | 0.922248373 | 5.02E-16 | North_western |
| UG_1398 | 0.000303033 | 0.000747509 | 0.217446898 | 0.78150256 | 7.78E-16 | North_western |
| UG_1406 | 0.003630317 | 0.087488941 | 0.879109398 | 0.029771344 | 6.26E-15 | Northern |
| UG_1326 | 0.000116559 | 0.00016558 | 0.091247566 | 0.908470295 | 6.76E-16 | North_western |
| UG_1334 | 9.37E-05 | 0.000125166 | 0.07915953 | 0.920621597 | 4.54E-16 | North_western |

| | | | | | | |
|---|---|---|---|---|---|---|
| UG_1342 | 0.001334447 | 0.003998463 | 0.400884005 | 0.593783085 | 7.95E-14 | North_western |
| UG_1350 | 2.00E-12 | 6.98E-13 | 7.87E-15 | 4.28E-16 | 1 | North_western |
| UG_1366 | 0.001226336 | 0.008445701 | 0.660583802 | 0.329744161 | 1.15E-15 | Northern |
| UG_1374 | 0.005089149 | 0.095980997 | 0.863060565 | 0.035869289 | 6.32E-14 | Northern |
| UG_1382 | 0.00015509 | 0.000257765 | 0.118338972 | 0.881248173 | 7.16E-16 | North_western |
| UG_1319 | 0.006721949 | 0.082476321 | 0.848682449 | 0.062119282 | 6.94E-13 | North_western |
| UG_1391 | 0.000181115 | 0.000261497 | 0.108735507 | 0.89082188 | 3.00E-15 | North_western |
| UG_1399 | 0.000101595 | 0.000155362 | 0.093520387 | 0.906222656 | 2.60E-16 | North_western |
| UG_1407 | 0.002555946 | 0.077263614 | 0.894490851 | 0.02568959 | 6.05E-16 | Northern |
| UG_1327 | 0.001668006 | 0.009222894 | 0.624756473 | 0.364352627 | 1.10E-14 | North_western |
| UG_1335 | 0.001163342 | 0.004734866 | 0.482201922 | 0.511899871 | 9.72E-15 | North_western |
| UG_1343 | 0.000388071 | 0.000936446 | 0.231066692 | 0.76760879 | 2.24E-15 | North_western |
| UG_1351 | 0.006472087 | 0.392999635 | 0.598325919 | 0.002202358 | 8.02E-14 | North_western |
| UG_1367 | 0.003965294 | 0.155975685 | 0.829179463 | 0.010879558 | 4.40E-15 | Northern |
| UG_1375 | 0.011735353 | 0.18773054 | 0.776691837 | 0.02384227 | 9.66E-12 | Northern |
| UG_1383 | 0.000390369 | 0.000792117 | 0.197436208 | 0.801381306 | 6.33E-15 | North_western |
| UG_1320 | 0.000123788 | 0.000172261 | 0.091338774 | 0.908365177 | 9.56E-16 | North_western |
| UG_1392 | 0.001850733 | 0.005280397 | 0.421434062 | 0.571434808 | 3.80E-13 | North_western |
| UG_1400 | 0.003007223 | 0.096559918 | 0.880101279 | 0.02033158 | 1.30E-15 | Northern |
| UG_1408 | 0.0045129 | 0.136373633 | 0.842785409 | 0.016328059 | 1.39E-14 | Northern |
| UG_1328 | 0.007022755 | 0.084694734 | 0.846285391 | 0.06199712 | 9.10E-13 | North_western |
| UG_1336 | 9.44E-05 | 0.00012617 | 0.079410309 | 0.920369098 | 4.66E-16 | North_western |
| UG_1344 | 0.000440547 | 0.001078468 | 0.243854425 | 0.754626559 | 3.32E-15 | North_western |
| UG_1352 | 0.004805475 | 0.49777996 | 0.496643516 | 0.000771049 | 1.22E-14 | North_western |
| UG_1368 | 0.018647561 | 0.224894001 | 0.730053355 | 0.026405083 | 2.39E-10 | Northern |
| UG_1376 | 0.009210473 | 0.152606825 | 0.810217662 | 0.027965041 | 2.20E-12 | Northern |
| UG_1384 | 0.009487826 | 0.075482506 | 0.815136601 | 0.099893067 | 1.18E-11 | North_western |
| UG_1321 | 0.000175566 | 0.000242949 | 0.103055367 | 0.896526118 | 3.49E-15 | North_western |
| UG_1393 | 0.003323884 | 0.053818442 | 0.879739342 | 0.063118331 | 9.26E-15 | North_western |

| | | | | | | |
|---|---|---|---|---|---|---|
| UG_1401 | 0.00301556 | 0.094352507 | 0.881347267 | 0.021284665 | 1.39E-15 | Northern |
| UG_1409 | 0.003134926 | 0.107307196 | 0.872105415 | 0.017452463 | 1.46E-15 | Northern |
| UG_1329 | 0.008795605 | 0.09832521 | 0.83298537 | 0.059893815 | 3.58E-12 | North_western |
| UG_1337 | 0.002351991 | 0.01506604 | 0.705691408 | 0.276890561 | 2.81E-14 | North_western |
| UG_1345 | 0.000344346 | 0.000618881 | 0.168767761 | 0.830269012 | 8.22E-15 | North_western |
| UG_1353 | 0.007046956 | 0.416992753 | 0.573919238 | 0.002041054 | 1.55E-13 | North_western |
| UG_1369 | 0.013187335 | 0.194850727 | 0.767018343 | 0.024943595 | 2.18E-11 | Northern |
| UG_1385 | 0.001103354 | 0.004544669 | 0.479974613 | 0.514377364 | 7.38E-15 | North_western |
| UG_1322 | 0.000152495 | 0.00023797 | 0.110481753 | 0.889127782 | 9.95E-16 | North_western |
| UG_1394 | 0.000104842 | 0.000150045 | 0.088462992 | 0.91128212 | 4.41E-16 | North_western |
| UG_1402 | 0.004138634 | 0.144226249 | 0.838312441 | 0.013322677 | 6.76E-15 | Northern |
| UG_1410 | 0.003967245 | 0.128025414 | 0.85194345 | 0.016063891 | 6.01E-15 | Northern |
| UG_1330 | 0.032468765 | 0.234171156 | 0.691455095 | 0.04190497 | 1.49E-08 | North_western |
| UG_1338 | 0.000426121 | 0.000917723 | 0.214772292 | 0.783883864 | 6.18E-15 | North_western |
| UG_1346 | 0.000326918 | 0.000765066 | 0.212426184 | 0.786481832 | 1.41E-15 | North_western |
| UG_1354 | 0.005109488 | 0.471443868 | 0.5224604 | 0.000986244 | 1.73E-14 | North_western |
| UG_1370 | 0.011150952 | 0.191027284 | 0.77601061 | 0.021811154 | 6.46E-12 | Northern |
| UG_1378 | 0.000862492 | 0.003454893 | 0.442816633 | 0.552865983 | 3.11E-15 | South_western |
| UG_1386 | 0.001077823 | 0.004500681 | 0.481728448 | 0.512693047 | 6.27E-15 | North_western |
| UG_1323 | 0.00047001 | 0.001285768 | 0.274534861 | 0.723709361 | 2.25E-15 | North_western |
| UG_1395 | 0.009290291 | 0.073942353 | 0.815539812 | 0.101227544 | 1.06E-11 | North_western |
| UG_1403 | 0.003651007 | 0.104077191 | 0.870502149 | 0.021769653 | 4.69E-15 | Northern |
| UG_1331 | 0.031111823 | 0.2261895 | 0.699451954 | 0.043246712 | 1.12E-08 | North_western |
| UG_1339 | 0.000482585 | 0.001070883 | 0.229878708 | 0.768567824 | 8.27E-15 | North_western |
| UG_1347 | 0.003539471 | 0.039723512 | 0.848246911 | 0.108490106 | 3.22E-14 | North_western |
| UG_1363 | 0.006782636 | 0.342409616 | 0.647450143 | 0.003357605 | 1.09E-13 | Northern |
| UG_1371 | 1.69E-12 | 5.32E-13 | 6.77E-15 | 4.48E-16 | 1 | South_western |
| UG_1379 | 0.000281619 | 0.000665782 | 0.204090404 | 0.794962196 | 7.60E-16 | North_western |
| UG_1387 | 0.00022339 | 0.000442957 | 0.159806784 | 0.839526868 | 9.23E-16 | North_western |

| | | | | | | |
|---|---|---|---|---|---|---|
| UG_1324 | 0.000358768 | 0.000819401 | 0.214188541 | 0.78463329 | 2.30E-15 | North_western |
| UG_1396 | 0.000163927 | 0.000269993 | 0.119615304 | 0.879950776 | 9.26E-16 | North_western |
| UG_1404 | 0.003662029 | 0.119719263 | 0.859855247 | 0.016763461 | 3.75E-15 | Northern |
| UG_1332 | 0.000222606 | 0.000408244 | 0.148022766 | 0.851346383 | 1.46E-15 | North_western |
| UG_1340 | 9.04E-05 | 0.000113888 | 0.07358796 | 0.926207747 | 5.77E-16 | North_western |
| UG_1348 | 0.001363361 | 0.004246862 | 0.415078765 | 0.579311012 | 7.08E-14 | North_western |
| UG_1364 | 0.020805053 | 0.204586752 | 0.738710064 | 0.035898131 | 6.12E-10 | Northern |
| UG_1380 | 0.000490165 | 0.001245486 | 0.260765879 | 0.73749847 | 4.01E-15 | North_western |
| UG_1388 | 0.005271364 | 0.111523758 | 0.854939895 | 0.028264982 | 6.16E-14 | North_western |
| UG_1411 | 0.002862578 | 0.101075835 | 0.878355899 | 0.017705688 | 8.34E-16 | Northern |
| UG_1483 | 0.00050725 | 0.001573286 | 0.312810492 | 0.685108971 | 1.49E-15 | Northern |
| UG_1491 | 0.007043737 | 0.130103098 | 0.83415499 | 0.028698176 | 3.97E-13 | Northern |
| UG_1499 | 0.003340838 | 0.111075287 | 0.868072404 | 0.01751147 | 2.18E-15 | Northern |
| UG_1419 | 0.00309129 | 0.093578832 | 0.881137341 | 0.022192537 | 1.69E-15 | Northern |
| UG_1427 | 0.003868767 | 0.142036021 | 0.841332046 | 0.012763166 | 4.23E-15 | Northern |
| UG_1435 | 0.004312954 | 0.122703051 | 0.853914744 | 0.01906925 | 1.19E-14 | Northern |
| UG_1443 | 0.002763701 | 0.077501733 | 0.891973932 | 0.027760634 | 1.07E-15 | Northern |
| UG_1451 | 0.000109791 | 0.00013367 | 0.076320916 | 0.923435623 | 1.44E-15 | Northern |
| UG_1459 | 0.008915006 | 0.244587715 | 0.736375843 | 0.010121436 | 9.45E-13 | Northern |
| UG_1467 | 0.00969615 | 0.121277863 | 0.823575834 | 0.045450153 | 4.85E-12 | Northern |
| UG_1475 | 0.008306065 | 0.210008799 | 0.768599723 | 0.013085413 | 6.58E-13 | Northern |
| UG_1484 | 0.010544966 | 0.202819683 | 0.768473804 | 0.018161547 | 3.96E-12 | Northern |
| UG_1492 | 0.00237802 | 0.070034469 | 0.899212117 | 0.028375394 | 4.34E-16 | Northern |
| UG_1500 | 0.003608439 | 0.118711009 | 0.860912517 | 0.016768035 | 3.41E-15 | Northern |
| UG_1420 | 0.003535108 | 0.09013188 | 0.878909077 | 0.027423935 | 4.85E-15 | Northern |
| UG_1428 | 0.003647899 | 0.137433434 | 0.846142428 | 0.012776239 | 2.90E-15 | Northern |
| UG_1436 | 0.004390793 | 0.139444456 | 0.840987646 | 0.015177104 | 1.10E-14 | Northern |
| UG_1444 | 0.004345624 | 0.12567944 | 0.851613474 | 0.018361462 | 1.21E-14 | Northern |
| UG_1452 | 0.009225503 | 0.117376846 | 0.827526945 | 0.045870706 | 3.56E-12 | Northern |

| | | | | | | |
|---|---|---|---|---|---|---|
| UG_1460 | 0.008118211 | 0.180270283 | 0.794049846 | 0.01756166 | 6.76E-13 | Northern |
| UG_1468 | 0.006620973 | 0.098115547 | 0.850052739 | 0.045210742 | 4.28E-13 | Northern |
| UG_1476 | 0.005621289 | 0.360483234 | 0.631507385 | 0.002388093 | 2.80E-14 | Northern |
| UG_1413 | 0.002822472 | 0.083096605 | 0.889051802 | 0.025029121 | 1.09E-15 | Northern |
| UG_1485 | 0.006289273 | 0.405331415 | 0.5864236 | 0.001955713 | 6.63E-14 | Northern |
| UG_1493 | 0.004539966 | 0.149282824 | 0.832422212 | 0.013754998 | 1.26E-14 | Northern |
| UG_1501 | 0.004145812 | 0.10900728 | 0.86397973 | 0.022867178 | 1.10E-14 | Northern |
| UG_1421 | 0.00303757 | 0.086438313 | 0.885331492 | 0.025192624 | 1.73E-15 | Northern |
| UG_1429 | 0.003455738 | 0.116330407 | 0.863579098 | 0.016634757 | 2.58E-15 | Northern |
| UG_1437 | 0.003403119 | 0.106970058 | 0.870448332 | 0.019178491 | 2.67E-15 | Northern |
| UG_1445 | 0.003061655 | 0.099831172 | 0.877625802 | 0.019481371 | 1.39E-15 | Northern |
| UG_1453 | 0.009570415 | 0.117496183 | 0.825420653 | 0.04751275 | 4.68E-12 | Northern |
| UG_1461 | 0.008362304 | 0.233932661 | 0.747274465 | 0.01043057 | 6.17E-13 | Northern |
| UG_1469 | 0.004555427 | 0.143418697 | 0.837081744 | 0.014944132 | 1.37E-14 | Northern |
| UG_1477 | 0.008209094 | 0.210195101 | 0.768698607 | 0.012897197 | 6.03E-13 | Northern |
| UG_1414 | 0.003551837 | 0.128954048 | 0.853439327 | 0.014054789 | 2.65E-15 | Northern |
| UG_1486 | 0.00451102 | 0.161700923 | 0.822148577 | 0.01163948 | 1.07E-14 | Northern |
| UG_1494 | 0.009286413 | 0.11617797 | 0.827492449 | 0.047043169 | 3.82E-12 | Northern |
| UG_1502 | 0.005129293 | 0.183725608 | 0.800843027 | 0.010302072 | 2.29E-14 | Northern |
| UG_1422 | 0.002802351 | 0.090192071 | 0.885623224 | 0.021382354 | 8.81E-16 | Northern |
| UG_1430 | 0.003125015 | 0.098683964 | 0.877842856 | 0.020348165 | 1.66E-15 | Northern |
| UG_1438 | 0.003421763 | 0.110053639 | 0.868239036 | 0.018285563 | 2.64E-15 | Northern |
| UG_1446 | 0.006398499 | 0.14129964 | 0.830165783 | 0.022136078 | 1.70E-13 | Northern |
| UG_1454 | 0.009334345 | 0.116744549 | 0.827047556 | 0.04687355 | 3.93E-12 | Northern |
| UG_1462 | 0.006792233 | 0.294618376 | 0.693696479 | 0.004892912 | 1.14E-13 | Northern |
| UG_1470 | 0.006334075 | 0.156803585 | 0.819013399 | 0.017848941 | 1.34E-13 | Northern |
| UG_1478 | 0.001650953 | 0.013393928 | 0.738543091 | 0.246412029 | 2.34E-15 | Northern |
| UG_1415 | 0.004722738 | 0.154792756 | 0.82712967 | 0.013354836 | 1.59E-14 | Northern |
| UG_1487 | 0.003400936 | 0.106889736 | 0.870516975 | 0.019192352 | 2.66E-15 | Northern |

| | | | | | | |
|---|---|---|---|---|---|---|
| UG_1495 | 0.003739788 | 0.122179577 | 0.857588488 | 0.016492147 | 4.22E-15 | Northern |
| UG_1503 | 0.004310643 | 0.153480476 | 0.829904159 | 0.012304721 | 8.28E-15 | Northern |
| UG_1423 | 0.003368031 | 0.111539368 | 0.867566384 | 0.017526217 | 2.30E-15 | Northern |
| UG_1431 | 0.0034149 | 0.108939992 | 0.869045384 | 0.018599724 | 2.65E-15 | Northern |
| UG_1439 | 0.003260976 | 0.11638031 | 0.864747386 | 0.015611328 | 1.69E-15 | Northern |
| UG_1447 | 0.003840178 | 0.101367107 | 0.870662203 | 0.024130511 | 7.14E-15 | Northern |
| UG_1455 | 0.009010771 | 0.117434226 | 0.828803973 | 0.04475103 | 2.99E-12 | Northern |
| UG_1463 | 0.007964573 | 0.168069442 | 0.804133497 | 0.019832488 | 6.49E-13 | Northern |
| UG_1471 | 0.010455526 | 0.210648808 | 0.762278874 | 0.016616792 | 3.55E-12 | Northern |
| UG_1479 | 0.006560564 | 0.408834334 | 0.582606651 | 0.001998451 | 9.07E-14 | Northern |
| UG_1416 | 0.003977219 | 0.114471863 | 0.861600287 | 0.019950631 | 7.41E-15 | Northern |
| UG_1488 | 0.003441385 | 0.10069129 | 0.874133245 | 0.021734081 | 3.23E-15 | Northern |
| UG_1496 | 0.004649213 | 0.154688944 | 0.827514237 | 0.013147606 | 1.42E-14 | Northern |
| UG_1504 | 0.00108884 | 0.006725348 | 0.612345751 | 0.37984006 | 1.08E-15 | Northern |
| UG_1424 | 0.002740487 | 0.097671007 | 0.881578601 | 0.018009905 | 6.45E-16 | Northern |
| UG_1432 | 0.005341435 | 0.126649288 | 0.845453092 | 0.022556185 | 5.41E-14 | Northern |
| UG_1440 | 0.00324755 | 0.129884072 | 0.854292808 | 0.01257557 | 1.36E-15 | Northern |
| UG_1448 | 0.003503648 | 0.093432904 | 0.877627081 | 0.025436367 | 4.24E-15 | Northern |
| UG_1456 | 0.005756846 | 0.35891847 | 0.632844782 | 0.002479902 | 3.32E-14 | Northern |
| UG_1464 | 0.004163921 | 0.08604387 | 0.874368279 | 0.03542393 | 1.78E-14 | Northern |
| UG_1472 | 0.008986589 | 0.207200844 | 0.769163458 | 0.014649109 | 1.19E-12 | Northern |
| UG_1480 | 0.008077486 | 0.352827548 | 0.635337555 | 0.003757412 | 3.90E-13 | Northern |
| UG_1417 | 0.003542637 | 0.124080791 | 0.857276852 | 0.015099719 | 2.77E-15 | Northern |
| UG_1489 | 0.003003307 | 0.131293076 | 0.854395082 | 0.011308536 | 7.58E-16 | Northern |
| UG_1497 | 0.002723565 | 0.042733289 | 0.879071145 | 0.075472001 | 3.61E-15 | Northern |
| UG_1425 | 0.003595505 | 0.086374801 | 0.879868455 | 0.030161239 | 5.98E-15 | Northern |
| UG_1433 | 0.002958445 | 0.096380975 | 0.880614072 | 0.020046509 | 1.16E-15 | Northern |
| UG_1441 | 0.002649983 | 0.080654246 | 0.891992392 | 0.024703379 | 7.25E-16 | Northern |
| UG_1449 | 0.003957024 | 0.140002681 | 0.842584178 | 0.013456118 | 5.11E-15 | Northern |

| | | | | | | |
|---|---|---|---|---|---|---|
| UG_1457 | 0.007174206 | 0.28304354 | 0.704066469 | 0.005715785 | 1.74E-13 | Northern |
| UG_1465 | 0.015445954 | 0.273782551 | 0.696718057 | 0.014053438 | 4.90E-11 | Northern |
| UG_1473 | 0.007057193 | 0.353266231 | 0.636444237 | 0.003232339 | 1.46E-13 | Northern |
| UG_1481 | 0.000591675 | 0.002558562 | 0.424595737 | 0.572254026 | 4.62E-16 | Northern |
| UG_1418 | 0.004402185 | 0.146323455 | 0.83543165 | 0.013842711 | 1.04E-14 | Northern |
| UG_1490 | 0.004007586 | 0.144204303 | 0.838918304 | 0.012869806 | 5.35E-15 | Northern |
| UG_1498 | 0.003960103 | 0.136631498 | 0.84528127 | 0.014127129 | 5.34E-15 | Northern |
| UG_1426 | 0.004120365 | 0.132772354 | 0.84751351 | 0.015593771 | 7.47E-15 | Northern |
| UG_1434 | 0.003202498 | 0.099423158 | 0.876772379 | 0.020601965 | 1.95E-15 | Northern |
| UG_1442 | 0.004491315 | 0.138095275 | 0.841562419 | 0.015850991 | 1.32E-14 | Northern |
| UG_1450 | 0.009415796 | 0.119690611 | 0.825699018 | 0.045194575 | 4.00E-12 | Northern |
| UG_1458 | 0.007040311 | 0.184506932 | 0.794075019 | 0.014377738 | 2.30E-13 | Northern |
| UG_1466 | 0.011778849 | 0.229443625 | 0.743087867 | 0.015689659 | 7.76E-12 | Northern |
| UG_1474 | 0.008933331 | 0.191680114 | 0.782249631 | 0.017136923 | 1.26E-12 | Northern |
| UG_1482 | 0.003505666 | 0.093303211 | 0.877674262 | 0.025516861 | 4.27E-15 | Northern |
| UG_1505 | 0.002943595 | 0.063966951 | 0.891465673 | 0.041623781 | 2.53E-15 | Northern |
| UG_1577 | 0.003394227 | 0.116052393 | 0.86416344 | 0.01638994 | 2.27E-15 | Northern |
| UG_1585 | 0.003639236 | 0.142464343 | 0.84202465 | 0.01187177 | 2.70E-15 | Northern |
| UG_1593 | 0.004543822 | 0.149157078 | 0.832508407 | 0.013790693 | 1.27E-14 | Northern |
| UG_1513 | 0.001534709 | 0.013347861 | 0.75058312 | 0.234534309 | 1.30E-15 | Northern |
| UG_1521 | 0.001147528 | 0.006823682 | 0.605944119 | 0.386084672 | 1.59E-15 | Northern |
| UG_1529 | 0.003848485 | 0.127226569 | 0.853190314 | 0.015734632 | 4.86E-15 | Northern |
| UG_1537 | 0.004258191 | 0.124337325 | 0.85306547 | 0.018339015 | 1.06E-14 | Northern |
| UG_1545 | 0.003012209 | 0.09403695 | 0.881559465 | 0.021391375 | 1.38E-15 | Northern |
| UG_1553 | 0.004345299 | 0.165306744 | 0.819660964 | 0.010686993 | 7.89E-15 | Northern |
| UG_1561 | 0.002932979 | 0.126626924 | 0.858614364 | 0.011825733 | 6.76E-16 | Northern |
| UG_1569 | 0.002890636 | 0.094403344 | 0.882386267 | 0.020319753 | 1.02E-15 | Northern |
| UG_1506 | 0.003651556 | 0.101826058 | 0.871844236 | 0.02267815 | 4.89E-15 | Northern |
| UG_1578 | 0.003087383 | 0.095705994 | 0.879945874 | 0.021260749 | 1.60E-15 | Northern |

| | | | | | | |
|---|---|---|---|---|---|---|
| UG_1586 | 0.003624748 | 0.114092 | 0.864110811 | 0.018172441 | 3.78E-15 | Northern |
| UG_1594 | 0.004688773 | 0.151892955 | 0.829657676 | 0.013760596 | 1.55E-14 | Northern |
| UG_1514 | 0.003260173 | 0.100072647 | 0.87592001 | 0.02074717 | 2.20E-15 | Northern |
| UG_1522 | 0.004161479 | 0.144616626 | 0.837890935 | 0.013330959 | 7.01E-15 | Northern |
| UG_1530 | 0.00396966 | 0.095004853 | 0.872863306 | 0.028162181 | 1.03E-14 | Northern |
| UG_1538 | 0.002212409 | 0.0365121 | 0.882107644 | 0.079167846 | 1.11E-15 | Northern |
| UG_1546 | 0.003507254 | 0.12306458 | 0.858253011 | 0.015175154 | 2.61E-15 | Northern |
| UG_1554 | 0.002763285 | 0.062728518 | 0.894181561 | 0.040326636 | 1.65E-15 | Northern |
| UG_1562 | 0.00323246 | 0.115478012 | 0.865594302 | 0.015695226 | 1.60E-15 | Northern |
| UG_1570 | 0.006019499 | 0.14270509 | 0.830926213 | 0.020349199 | 1.07E-13 | Northern |
| UG_1507 | 0.004091942 | 0.13712502 | 0.84424926 | 0.014533778 | 6.74E-15 | Northern |
| UG_1579 | 0.004439781 | 0.155171329 | 0.827959446 | 0.012429444 | 1.01E-14 | Northern |
| UG_1587 | 0.003543905 | 0.10571338 | 0.870262151 | 0.020480564 | 3.67E-15 | Northern |
| UG_1595 | 0.003899017 | 0.10595494 | 0.867559733 | 0.022586309 | 7.36E-15 | Northern |
| UG_1515 | 0.002825604 | 0.072728413 | 0.89260821 | 0.031837773 | 1.43E-15 | Northern |
| UG_1523 | 0.003441652 | 0.135851859 | 0.84843768 | 0.012268808 | 1.94E-15 | Northern |
| UG_1531 | 0.003704123 | 0.124482805 | 0.85606637 | 0.015746703 | 3.81E-15 | Northern |
| UG_1539 | 0.004094035 | 0.143727525 | 0.838920848 | 0.013257593 | 6.28E-15 | Northern |
| UG_1547 | 0.003966146 | 0.149796715 | 0.834437771 | 0.011799368 | 4.68E-15 | Northern |
| UG_1555 | 0.004292804 | 0.149064282 | 0.833659102 | 0.012983812 | 8.40E-15 | Northern |
| UG_1563 | 0.002972401 | 0.088306224 | 0.885047117 | 0.023674259 | 1.41E-15 | Northern |
| UG_1571 | 0.002643902 | 0.080541335 | 0.892109198 | 0.024705565 | 7.15E-16 | Northern |
| UG_1508 | 0.00382014 | 0.130818988 | 0.850571226 | 0.014789647 | 4.40E-15 | Northern |
| UG_1580 | 0.003327126 | 0.123569819 | 0.858882863 | 0.014220192 | 1.76E-15 | Northern |
| UG_1588 | 0.003160748 | 0.072466226 | 0.888332562 | 0.036040464 | 3.30E-15 | Northern |
| UG_1596 | 0.003881888 | 0.119286506 | 0.858857745 | 0.01797386 | 5.78E-15 | Northern |
| UG_1516 | 0.003525116 | 0.103224281 | 0.871958642 | 0.021291962 | 3.68E-15 | Northern |
| UG_1524 | 0.003021675 | 0.132643018 | 0.85317696 | 0.011158347 | 7.79E-16 | Northern |
| UG_1532 | 0.003337589 | 0.081394986 | 0.884234842 | 0.031032583 | 3.89E-15 | Northern |

| | | | | | | |
|---|---|---|---|---|---|---|
| UG_1540 | 0.000558103 | 0.002075093 | 0.372577487 | 0.624789316 | 8.06E-16 | Northern |
| UG_1548 | 0.004019272 | 0.144512744 | 0.838611917 | 0.012856067 | 5.45E-15 | Northern |
| UG_1556 | 0.00360602 | 0.13571287 | 0.847748404 | 0.012932706 | 2.72E-15 | Northern |
| UG_1564 | 0.004622788 | 0.151038414 | 0.830634434 | 0.013704364 | 1.41E-14 | Northern |
| UG_1572 | 0.002742955 | 0.077476173 | 0.892224966 | 0.027555906 | 1.01E-15 | Northern |
| UG_1509 | 0.00373846 | 0.112122452 | 0.864723587 | 0.019415501 | 4.89E-15 | Northern |
| UG_1581 | 0.004355981 | 0.129353874 | 0.8488753 | 0.017414845 | 1.17E-14 | Northern |
| UG_1589 | 0.00312092 | 0.099775709 | 0.87719585 | 0.019907521 | 1.61E-15 | Northern |
| UG_1597 | 0.007177102 | 0.165653018 | 0.80889239 | 0.01827749 | 3.08E-13 | Northern |
| UG_1517 | 0.004063929 | 0.173827153 | 0.813144501 | 0.008964417 | 4.52E-15 | Northern |
| UG_1525 | 0.001875465 | 0.016932243 | 0.776468686 | 0.204723605 | 2.90E-15 | Northern |
| UG_1533 | 0.003743095 | 0.123124191 | 0.856867205 | 0.016265509 | 4.19E-15 | Northern |
| UG_1541 | 0.000559902 | 0.002286752 | 0.400917992 | 0.596235354 | 4.98E-16 | Northern |
| UG_1549 | 0.00410186 | 0.146511255 | 0.836596759 | 0.012790125 | 6.19E-15 | Northern |
| UG_1557 | 0.00389925 | 0.124238314 | 0.855155692 | 0.016706744 | 5.57E-15 | Northern |
| UG_1565 | 0.004724668 | 0.146024425 | 0.834248955 | 0.015001952 | 1.75E-14 | Northern |
| UG_1573 | 0.002689022 | 0.074095206 | 0.89398679 | 0.029228981 | 9.56E-16 | Northern |
| UG_1510 | 0.0040898 | 0.140438772 | 0.841610562 | 0.013860866 | 6.46E-15 | Northern |
| UG_1582 | 0.004731896 | 0.126605573 | 0.8488278 | 0.019834732 | 2.23E-14 | Northern |
| UG_1590 | 0.003056135 | 0.077334108 | 0.888613395 | 0.030996361 | 2.25E-15 | Northern |
| UG_1598 | 0.003322247 | 0.067640021 | 0.886218521 | 0.04281921 | 5.52E-15 | Northern |
| UG_1518 | 0.003482173 | 0.110509134 | 0.867520344 | 0.018488349 | 2.98E-15 | Northern |
| UG_1526 | 0.004305196 | 0.126331183 | 0.851365327 | 0.017998294 | 1.12E-14 | Northern |
| UG_1534 | 0.003344576 | 0.102273674 | 0.873905538 | 0.020476212 | 2.55E-15 | Northern |
| UG_1542 | 0.001073347 | 0.005932588 | 0.574048744 | 0.418945321 | 1.65E-15 | Northern |
| UG_1550 | 0.003256201 | 0.115358374 | 0.865534184 | 0.015851241 | 1.69E-15 | Northern |
| UG_1558 | 0.003515095 | 0.139434138 | 0.845123222 | 0.011927545 | 2.17E-15 | Northern |
| UG_1566 | 0.003586863 | 0.11382911 | 0.864537084 | 0.018046943 | 3.51E-15 | Northern |
| UG_1574 | 0.003625704 | 0.11882409 | 0.860726267 | 0.016823939 | 3.53E-15 | Northern |

| | | | | | | |
|---|---|---|---|---|---|---|
| UG_1511 | 0.002427832 | 0.032772968 | 0.862509107 | 0.102290093 | 2.99E-15 | Northern |
| UG_1583 | 0.002725578 | 0.086404864 | 0.888413882 | 0.022455676 | 7.80E-16 | Northern |
| UG_1591 | 0.003203548 | 0.10289697 | 0.874569658 | 0.019329824 | 1.84E-15 | Northern |
| UG_1519 | 0.004632602 | 0.157074703 | 0.825591426 | 0.012701269 | 1.35E-14 | Northern |
| UG_1527 | 0.001068109 | 0.006018709 | 0.579879341 | 0.413033841 | 1.49E-15 | Northern |
| UG_1535 | 0.003768141 | 0.125071643 | 0.855264575 | 0.01589564 | 4.29E-15 | Northern |
| UG_1543 | 0.004362805 | 0.1723532 | 0.813427189 | 0.009856806 | 7.67E-15 | Northern |
| UG_1551 | 0.003924406 | 0.116427022 | 0.860603337 | 0.019045234 | 6.52E-15 | Northern |
| UG_1559 | 0.003472382 | 0.130237719 | 0.852837246 | 0.013452652 | 2.21E-15 | Northern |
| UG_1567 | 0.00325452 | 0.117488613 | 0.863958725 | 0.015298142 | 1.63E-15 | Northern |
| UG_1575 | 0.003730703 | 0.117642104 | 0.860943458 | 0.017683735 | 4.42E-15 | Northern |
| UG_1512 | 0.003521943 | 0.130040358 | 0.852736265 | 0.013701434 | 2.46E-15 | Northern |
| UG_1584 | 0.003892397 | 0.100829807 | 0.870555958 | 0.024721838 | 7.96E-15 | Northern |
| UG_1592 | 0.001070852 | 0.006055831 | 0.581366448 | 0.411506869 | 1.48E-15 | Northern |
| UG_1520 | 0.004050521 | 0.134243212 | 0.846722616 | 0.014983651 | 6.47E-15 | Northern |
| UG_1528 | 0.003023006 | 0.069555168 | 0.890445946 | 0.036975879 | 2.58E-15 | Northern |
| UG_1536 | 0.002804814 | 0.084232694 | 0.888706741 | 0.024255751 | 1.01E-15 | Northern |
| UG_1544 | 0.000688018 | 0.0033592 | 0.480679365 | 0.515273418 | 4.63E-16 | Northern |
| UG_1552 | 0.001094767 | 0.006312015 | 0.590338681 | 0.402254537 | 1.49E-15 | Northern |
| UG_1560 | 0.004228607 | 0.163027049 | 0.822072632 | 0.010671712 | 6.60E-15 | Northern |
| UG_1568 | 0.004158042 | 0.118333173 | 0.857860478 | 0.019648307 | 9.69E-15 | Northern |
| UG_1576 | 0.00375698 | 0.144408331 | 0.839869984 | 0.011964705 | 3.33E-15 | Northern |
| UG_1599 | 0.004100557 | 0.140981034 | 0.841123036 | 0.013795373 | 6.55E-15 | Northern |
| UG_1671 | 0.009133046 | 0.18203005 | 0.789327186 | 0.019509718 | 1.58E-12 | Northern |
| UG_1679 | 0.000514949 | 0.001911033 | 0.363679574 | 0.633894444 | 5.93E-16 | Northern |
| UG_1687 | 0.003751603 | 0.128857899 | 0.852456291 | 0.014934206 | 3.95E-15 | Northern |
| UG_1607 | 0.005506716 | 0.148852511 | 0.828608449 | 0.017032324 | 5.20E-14 | Northern |
| UG_1615 | 0.000622755 | 0.002619633 | 0.421686687 | 0.575070925 | 6.53E-16 | Northern |
| UG_1623 | 0.090515575 | 0.688082355 | 0.218060049 | 0.00306925 | 0.00027277 | Northern |

| | | | | | | |
|---|---|---|---|---|---|---|
| UG_1631 | 0.00172964 | 0.018406657 | 0.80758664 | 0.172277063 | 1.16E-15 | Northern |
| UG_1639 | 0.004631293 | 0.157065644 | 0.825604209 | 0.012698853 | 1.35E-14 | Northern |
| UG_1647 | 0.006421666 | 0.1262607 | 0.839751719 | 0.027565915 | 2.11E-13 | Northern |
| UG_1655 | 0.003568191 | 0.070142354 | 0.883014454 | 0.043275001 | 8.67E-15 | Northern |
| UG_1663 | 0.012607299 | 0.12803439 | 0.805821836 | 0.053536475 | 3.14E-11 | Northern |
| UG_1600 | 0.00063806 | 0.002589452 | 0.413259477 | 0.583513011 | 8.66E-16 | Northern |
| UG_1672 | 0.009403345 | 0.193101735 | 0.779674137 | 0.017820783 | 1.81E-12 | Northern |
| UG_1680 | 0.003424311 | 0.083341758 | 0.882677132 | 0.030556799 | 4.48E-15 | Northern |
| UG_1688 | 0.001881028 | 0.019943067 | 0.812151336 | 0.166024569 | 1.74E-15 | Northern |
| UG_1608 | 0.00421497 | 0.127818207 | 0.850765803 | 0.017201021 | 9.39E-15 | Northern |
| UG_1616 | 0.001239492 | 0.007274368 | 0.61049187 | 0.38099427 | 2.35E-15 | Northern |
| UG_1624 | 0.003357637 | 0.596979076 | 0.399399772 | 0.000263515 | 1.52E-15 | Northern |
| UG_1632 | 0.000523641 | 0.001983616 | 0.371369742 | 0.626123001 | 5.68E-16 | Northern |
| UG_1640 | 0.003931942 | 0.135543702 | 0.846285054 | 0.014239302 | 5.13E-15 | Northern |
| UG_1648 | 0.002577515 | 0.039392285 | 0.87660188 | 0.081428321 | 2.92E-15 | Northern |
| UG_1656 | 0.000935403 | 0.004504172 | 0.512048855 | 0.48251157 | 1.76E-15 | Northern |
| UG_1664 | 0.003126838 | 0.108672207 | 0.871207561 | 0.016993394 | 1.40E-15 | Northern |
| UG_1601 | 0.000234804 | 0.000403801 | 0.141668625 | 0.857692771 | 2.63E-15 | Northern |
| UG_1673 | 0.009619943 | 0.194212271 | 0.778127459 | 0.018040328 | 2.13E-12 | Northern |
| UG_1681 | 0.00068859 | 0.003128639 | 0.457445195 | 0.538737576 | 6.63E-16 | Northern |
| UG_1689 | 0.003847784 | 0.122155769 | 0.856983903 | 0.017012543 | 5.20E-15 | Northern |
| UG_1609 | 0.001750288 | 0.011103634 | 0.673183637 | 0.313962441 | 7.74E-15 | Northern |
| UG_1617 | 0.002251024 | 0.03360688 | 0.872491787 | 0.091650309 | 1.57E-15 | Northern |
| UG_1625 | 0.003758313 | 0.116000425 | 0.861933465 | 0.018307798 | 4.78E-15 | Northern |
| UG_1633 | 0.004777817 | 0.138333041 | 0.840003835 | 0.016885307 | 2.06E-14 | Northern |
| UG_1641 | 0.006291749 | 0.123463669 | 0.842099167 | 0.028145415 | 1.89E-13 | Northern |
| UG_1649 | 0.007758457 | 0.126702243 | 0.832176036 | 0.033363264 | 8.50E-13 | Northern |
| UG_1657 | 0.000512939 | 0.001928639 | 0.367055171 | 0.630503252 | 5.45E-16 | Northern |
| UG_1665 | 0.000487216 | 0.001797651 | 0.356344459 | 0.641370674 | 4.93E-16 | Northern |

| | | | | | | |
|---|---|---|---|---|---|---|
| UG_1602 | 0.004507754 | 0.03960021 | 0.820443465 | 0.135448571 | 2.12E-13 | Northern |
| UG_1674 | 0.003239946 | 0.118740427 | 0.863101586 | 0.014918041 | 1.55E-15 | Northern |
| UG_1682 | 0.004357306 | 0.11261267 | 0.860345347 | 0.022684677 | 1.49E-14 | Northern |
| UG_1690 | 0.002801739 | 0.587550462 | 0.409417448 | 0.000230351 | 3.86E-16 | Northern |
| UG_1610 | 0.004480448 | 0.039724388 | 0.821672476 | 0.134122688 | 2.01E-13 | Northern |
| UG_1618 | 0.000902556 | 0.005254696 | 0.570989263 | 0.422853486 | 6.27E-16 | Northern |
| UG_1626 | 0.000334228 | 0.001038788 | 0.275840508 | 0.722786476 | 2.97E-16 | Northern |
| UG_1634 | 0.010108291 | 0.082636534 | 0.815623934 | 0.091631241 | 1.54E-11 | Northern |
| UG_1642 | 0.003912748 | 0.148343146 | 0.835889255 | 0.011854851 | 4.30E-15 | Northern |
| UG_1650 | 0.004166548 | 0.048105139 | 0.853573044 | 0.094155269 | 6.78E-14 | Northern |
| UG_1658 | 0.001662614 | 0.011943057 | 0.705033211 | 0.281361118 | 3.79E-15 | Northern |
| UG_1666 | 0.032747961 | 0.261087409 | 0.672711641 | 0.033452974 | 1.41E-08 | Northern |
| UG_1603 | 0.004459219 | 0.039166763 | 0.820062056 | 0.136311963 | 2.01E-13 | Northern |
| UG_1675 | 0.002538748 | 0.036376439 | 0.870168444 | 0.090916369 | 3.19E-15 | Northern |
| UG_1683 | 0.003968506 | 0.049056716 | 0.859791694 | 0.087183084 | 4.44E-14 | Northern |
| UG_1691 | 0.004638289 | 0.159631956 | 0.823417861 | 0.012311894 | 1.33E-14 | Northern |
| UG_1611 | 0.00376035 | 0.119849719 | 0.859176713 | 0.017213217 | 4.54E-15 | Northern |
| UG_1619 | 0.000563044 | 0.002153457 | 0.381816252 | 0.615467246 | 7.18E-16 | Northern |
| UG_1627 | 0.004183631 | 0.129588464 | 0.849611853 | 0.016616052 | 8.69E-15 | Northern |
| UG_1635 | 0.004419663 | 0.117259745 | 0.856981023 | 0.021339568 | 1.54E-14 | Northern |
| UG_1643 | 0.004071377 | 0.123597665 | 0.854654115 | 0.017676843 | 7.71E-15 | Northern |
| UG_1651 | 0.115399183 | 0.560878929 | 0.312808094 | 0.010358897 | 0.000554896 | Northern |
| UG_1659 | 0.000377602 | 0.001292948 | 0.311037567 | 0.687291883 | 2.75E-16 | Northern |
| UG_1667 | 0.0112705 | 0.21547322 | 0.756112069 | 0.017144211 | 6.01E-12 | Northern |
| UG_1604 | 0.003091378 | 0.128064595 | 0.856593062 | 0.012250964 | 9.73E-16 | Northern |
| UG_1676 | 0.000654274 | 0.002839472 | 0.437004949 | 0.559501306 | 6.79E-16 | Northern |
| UG_1684 | 0.004587783 | 0.133704675 | 0.844436476 | 0.017271067 | 1.62E-14 | Northern |
| UG_1692 | 0.00040949 | 0.001286032 | 0.296307673 | 0.701996805 | 6.09E-16 | Northern |
| UG_1612 | 0.001741537 | 0.01595206 | 0.774186245 | 0.208120158 | 1.95E-15 | Northern |

| | | | | | | |
|---|---|---|---|---|---|---|
| UG_1620 | 0.00442997 | 0.04083151 | 0.827145786 | 0.127592735 | 1.70E-13 | Northern |
| UG_1628 | 0.000112288 | 0.000133821 | 0.075291987 | 0.924461904 | 1.78E-15 | Northern |
| UG_1636 | 0.00052414 | 0.001837953 | 0.349341585 | 0.648296322 | 8.63E-16 | Northern |
| UG_1644 | 0.001081322 | 0.006500274 | 0.602661494 | 0.389756911 | 1.18E-15 | Northern |
| UG_1652 | 0.00051573 | 0.001830609 | 0.351140406 | 0.646513254 | 7.58E-16 | Northern |
| UG_1660 | 0.002516332 | 0.038203991 | 0.875781584 | 0.083498093 | 2.63E-15 | Northern |
| UG_1668 | 0.000514278 | 0.001906956 | 0.36330572 | 0.634273047 | 5.93E-16 | Northern |
| UG_1605 | 0.004223339 | 0.131225004 | 0.848168569 | 0.016383088 | 9.12E-15 | Northern |
| UG_1677 | 0.000838062 | 0.004276541 | 0.518357824 | 0.476527572 | 8.44E-16 | Northern |
| UG_1685 | 0.000513508 | 0.001868532 | 0.357745953 | 0.639872007 | 6.52E-16 | Northern |
| UG_1613 | 0.001380312 | 0.010069363 | 0.690793551 | 0.297756775 | 1.55E-15 | Northern |
| UG_1621 | 0.048230035 | 0.323327274 | 0.599036937 | 0.02940552 | 2.34E-07 | Northern |
| UG_1629 | 0.004458188 | 0.569309991 | 0.425795393 | 0.000436427 | 9.99E-15 | Northern |
| UG_1637 | 0.000481367 | 0.001776955 | 0.355256907 | 0.642484771 | 4.69E-16 | Northern |
| UG_1645 | 0.11875223 | 0.554550193 | 0.314983859 | 0.011034482 | 0.000679236 | Northern |
| UG_1653 | 0.033097923 | 0.262866905 | 0.670740514 | 0.033294643 | 1.51E-08 | Northern |
| UG_1661 | 0.000930889 | 0.005521297 | 0.5807908 | 0.412757014 | 6.58E-16 | Northern |
| UG_1669 | 0.000718897 | 0.003539261 | 0.488519623 | 0.507222219 | 5.32E-16 | Northern |
| UG_1606 | 0.007473883 | 0.102148012 | 0.842786795 | 0.04759131 | 9.74E-13 | Northern |
| UG_1678 | 0.003259386 | 0.066822625 | 0.88704026 | 0.042877728 | 4.91E-15 | Northern |
| UG_1686 | 0.003261055 | 0.10544945 | 0.872470961 | 0.018818534 | 2.00E-15 | Northern |
| UG_1614 | 0.004377413 | 0.03962513 | 0.824115099 | 0.131882358 | 1.68E-13 | Northern |
| UG_1622 | 0.134396744 | 0.573485028 | 0.279759021 | 0.009854036 | 0.002505171 | Northern |
| UG_1630 | 0.005130551 | 0.156186984 | 0.824336958 | 0.014345508 | 2.88E-14 | Northern |
| UG_1638 | 0.009394206 | 0.120572039 | 0.825543117 | 0.044490638 | 3.87E-12 | Northern |
| UG_1646 | 0.000467105 | 0.001724425 | 0.352225095 | 0.645583375 | 4.16E-16 | Northern |
| UG_1654 | 0.009444176 | 0.191338157 | 0.780971512 | 0.018246156 | 1.90E-12 | Northern |
| UG_1662 | 0.00139599 | 0.009743934 | 0.678612855 | 0.310247222 | 1.94E-15 | Northern |
| UG_1670 | 0.000491251 | 0.001794534 | 0.354345346 | 0.643368869 | 5.37E-16 | Northern |

| | | | | | | |
|---|---|---|---|---|---|---|
| UG_1693 | 0.003452542 | 0.546046355 | 0.450115146 | 0.000385957 | 1.38E-15 | Northern |
| UG_1765 | 0.00835714 | 0.522274321 | 0.468171898 | 0.001196641 | 7.54E-13 | Northern |
| UG_1773 | 0.003217703 | 0.638954555 | 0.35764133 | 0.000186412 | 1.52E-15 | Northern |
| UG_1781 | 0.003186979 | 0.651506995 | 0.345137801 | 0.000168224 | 1.57E-15 | Northern |
| UG_1701 | 0.004591764 | 0.130823371 | 0.846552915 | 0.01803195 | 1.69E-14 | Northern |
| UG_1709 | 0.004090054 | 0.13409811 | 0.846638171 | 0.015173665 | 6.96E-15 | Northern |
| UG_1717 | 0.007964725 | 0.149824551 | 0.817317249 | 0.024893476 | 7.75E-13 | Northern |
| UG_1725 | 0.004543483 | 0.151768521 | 0.830366372 | 0.013321625 | 1.24E-14 | Northern |
| UG_1733 | 0.003840562 | 0.06234916 | 0.87674821 | 0.057062068 | 1.95E-14 | Northern |
| UG_1741 | 0.000682292 | 0.002898047 | 0.434934795 | 0.561484866 | 8.97E-16 | Northern |
| UG_1749 | 0.00409319 | 0.125611948 | 0.853059706 | 0.017235156 | 7.80E-15 | Northern |
| UG_1757 | 0.002796786 | 0.593550609 | 0.403432105 | 0.0002205 | 3.96E-16 | Northern |
| UG_1694 | 0.000521503 | 0.00204828 | 0.381570443 | 0.615859774 | 4.61E-16 | Northern |
| UG_1766 | 0.003366136 | 0.604450313 | 0.391932793 | 0.000250758 | 1.63E-15 | Northern |
| UG_1774 | 0.006291458 | 0.133968034 | 0.835642594 | 0.024097914 | 1.64E-13 | Northern |
| UG_1782 | 0.003383607 | 0.603422541 | 0.392939826 | 0.000254026 | 1.68E-15 | Northern |
| UG_1702 | 0.000493325 | 0.00182644 | 0.358594017 | 0.639086218 | 5.08E-16 | Northern |
| UG_1710 | 0.032928638 | 0.264352059 | 0.669998545 | 0.032720744 | 1.45E-08 | Northern |
| UG_1718 | 0.004018945 | 0.116828187 | 0.859742089 | 0.019410779 | 7.72E-15 | Northern |
| UG_1726 | 0.004472215 | 0.127946972 | 0.849284483 | 0.01829633 | 1.45E-14 | Northern |
| UG_1734 | 0.00231397 | 0.037111366 | 0.879979149 | 0.080595515 | 1.50E-15 | Northern |
| UG_1742 | 0.000627975 | 0.002645176 | 0.42303723 | 0.573689619 | 6.71E-16 | Northern |
| UG_1750 | 0.004942173 | 0.154953073 | 0.826106889 | 0.013997865 | 2.22E-14 | Northern |
| UG_1758 | 0.003252729 | 0.603284004 | 0.393219836 | 0.00024343 | 1.26E-15 | Northern |
| UG_1695 | 0.000477785 | 0.001841212 | 0.366826551 | 0.630854453 | 3.61E-16 | Northern |
| UG_1767 | 0.003483477 | 0.606752282 | 0.389508001 | 0.000256239 | 2.12E-15 | Northern |
| UG_1775 | 0.006880013 | 0.114026729 | 0.843267945 | 0.035825313 | 4.24E-13 | Northern |
| UG_1783 | 0.002431119 | 0.596574426 | 0.400809609 | 0.000184846 | 1.46E-16 | Northern |
| UG_1703 | 0.000683591 | 0.003319929 | 0.478196739 | 0.517799742 | 4.63E-16 | Northern |

| | | | | | | |
|---|---|---|---|---|---|---|
| UG_1711 | 0.003086659 | 0.604532587 | 0.392153032 | 0.000227721 | 8.70E-16 | Northern |
| UG_1719 | 0.000465608 | 0.001672294 | 0.344185083 | 0.653677014 | 4.77E-16 | Northern |
| UG_1727 | 0.004201053 | 0.14968709 | 0.833533428 | 0.012578429 | 7.13E-15 | Northern |
| UG_1735 | 0.003788374 | 0.131679113 | 0.850061442 | 0.014471071 | 4.10E-15 | Northern |
| UG_1743 | 0.003685946 | 0.139927199 | 0.843914793 | 0.012472062 | 3.05E-15 | Northern |
| UG_1751 | 0.003431354 | 0.082304865 | 0.882943826 | 0.031319955 | 4.66E-15 | Northern |
| UG_1759 | 0.019430374 | 0.672161511 | 0.307409532 | 0.000998582 | 1.08E-09 | Northern |
| UG_1696 | 0.032254596 | 0.254980316 | 0.678043759 | 0.034721316 | 1.28E-08 | Northern |
| UG_1768 | 0.003192132 | 0.648641019 | 0.347994723 | 0.000172126 | 1.55E-15 | Northern |
| UG_1776 | 0.003177857 | 0.564039927 | 0.43247079 | 0.000311426 | 8.33E-16 | Northern |
| UG_1784 | 0.002972541 | 0.620780551 | 0.376052227 | 0.00019468 | 7.44E-16 | Northern |
| UG_1704 | 0.007931002 | 0.130711392 | 0.829164644 | 0.032192962 | 9.46E-13 | Northern |
| UG_1712 | 0.002846002 | 0.61436475 | 0.382595048 | 0.000194199 | 5.18E-16 | Northern |
| UG_1720 | 0.003597022 | 0.106855927 | 0.869152413 | 0.020394638 | 4.01E-15 | Northern |
| UG_1728 | 0.00588343 | 0.14382259 | 0.830734938 | 0.019559042 | 8.91E-14 | Northern |
| UG_1736 | 0.004514738 | 0.121008841 | 0.853912051 | 0.02056437 | 1.70E-14 | Northern |
| UG_1744 | 0.004511672 | 0.122683651 | 0.852786625 | 0.020018052 | 1.66E-14 | Northern |
| UG_1752 | 0.002797192 | 0.037844317 | 0.865594697 | 0.093763795 | 6.03E-15 | Northern |
| UG_1760 | 0.049878198 | 0.655832248 | 0.291540503 | 0.002747793 | 1.26E-06 | Northern |
| UG_1697 | 0.032692404 | 0.264997216 | 0.669990886 | 0.032319481 | 1.37E-08 | Northern |
| UG_1769 | 0.001905341 | 0.010252197 | 0.630798768 | 0.357043694 | 2.20E-14 | Northern |
| UG_1777 | 0.005246003 | 0.404829658 | 0.588316182 | 0.001608157 | 1.78E-14 | Northern |
| UG_1785 | 0.015896341 | 0.587024619 | 0.395555804 | 0.001523237 | 1.21E-10 | Northern |
| UG_1705 | 0.003736778 | 0.086745572 | 0.878349106 | 0.031168544 | 7.87E-15 | Northern |
| UG_1713 | 0.006235172 | 0.122425301 | 0.843013113 | 0.028326415 | 1.79E-13 | Northern |
| UG_1721 | 0.00376595 | 0.098262469 | 0.872937789 | 0.025033792 | 6.56E-15 | Northern |
| UG_1729 | 0.003775174 | 0.130170718 | 0.85131114 | 0.014742968 | 4.07E-15 | Northern |
| UG_1737 | 0.000102813 | 0.000119847 | 0.071257651 | 0.928519689 | 1.49E-15 | Northern |
| UG_1745 | 0.003701081 | 0.138170223 | 0.845285676 | 0.012843019 | 3.20E-15 | Northern |

| | | | | | | |
|---|---|---|---|---|---|---|
| UG_1753 | 0.005001577 | 0.1662605 | 0.816432561 | 0.012305362 | 2.18E-14 | Northern |
| UG_1761 | 0.018749516 | 0.569536417 | 0.409668707 | 0.002045359 | 3.66E-10 | Northern |
| UG_1698 | 0.031791496 | 0.254674723 | 0.679203122 | 0.034330647 | 1.15E-08 | Northern |
| UG_1770 | 0.003116919 | 0.606589499 | 0.3900667 | 0.000226881 | 9.47E-16 | Northern |
| UG_1778 | 0.004886246 | 0.456873696 | 0.537203112 | 0.001036946 | 1.19E-14 | Northern |
| UG_1786 | 0.005930554 | 0.401324409 | 0.59085872 | 0.001886317 | 4.30E-14 | Northern |
| UG_1706 | 0.003829522 | 0.113098579 | 0.863472222 | 0.019599677 | 5.74E-15 | Northern |
| UG_1714 | 0.003337667 | 0.099332341 | 0.875758219 | 0.021571772 | 2.65E-15 | Northern |
| UG_1722 | 0.003850263 | 0.111176307 | 0.864611614 | 0.020361815 | 6.15E-15 | Northern |
| UG_1730 | 0.003518453 | 0.117915031 | 0.862038082 | 0.016528434 | 2.87E-15 | Northern |
| UG_1738 | 9.59E-05 | 0.000110623 | 0.068728244 | 0.931065271 | 1.24E-15 | Northern |
| UG_1746 | 0.003272612 | 0.120234018 | 0.861769833 | 0.014723537 | 1.64E-15 | Northern |
| UG_1754 | 0.002954623 | 0.114378406 | 0.868165077 | 0.014501894 | 8.45E-16 | Northern |
| UG_1762 | 0.003191154 | 0.644482458 | 0.352148982 | 0.000177406 | 1.50E-15 | Northern |
| UG_1699 | 0.001597355 | 0.014023065 | 0.756479712 | 0.227899868 | 1.52E-15 | Northern |
| UG_1771 | 0.00280239 | 0.031962754 | 0.844083561 | 0.121151294 | 9.69E-15 | Northern |
| UG_1779 | 0.004756035 | 0.441011041 | 0.553110729 | 0.001122195 | 9.40E-15 | Northern |
| UG_1707 | 0.003877155 | 0.115360897 | 0.861631666 | 0.019130282 | 6.07E-15 | Northern |
| UG_1715 | 0.002727114 | 0.044854782 | 0.882526688 | 0.069891416 | 3.23E-15 | Northern |
| UG_1723 | 0.004724931 | 0.145650202 | 0.83454583 | 0.015079037 | 1.75E-14 | Northern |
| UG_1731 | 0.004620333 | 0.134441904 | 0.843719718 | 0.017218045 | 1.69E-14 | Northern |
| UG_1739 | 0.003123956 | 0.099911053 | 0.877086952 | 0.019878039 | 1.61E-15 | Northern |
| UG_1747 | 0.003820479 | 0.151192691 | 0.833865925 | 0.011120905 | 3.51E-15 | Northern |
| UG_1755 | 0.003203498 | 0.093179233 | 0.880380084 | 0.023237186 | 2.21E-15 | Northern |
| UG_1763 | 0.003006567 | 0.621571689 | 0.375225708 | 0.000196035 | 8.13E-16 | Northern |
| UG_1700 | 0.00756205 | 0.12560906 | 0.833809945 | 0.033018945 | 7.14E-13 | Northern |
| UG_1772 | 0.003792683 | 0.513710644 | 0.481963485 | 0.000533188 | 2.34E-15 | Northern |
| UG_1780 | 0.003409128 | 0.07709552 | 0.884536743 | 0.034958609 | 5.07E-15 | Northern |
| UG_1708 | 0.000480127 | 0.001842428 | 0.366108325 | 0.63156912 | 3.77E-16 | Northern |

| | | | | | | |
|---|---|---|---|---|---|---|
| UG_1716 | 0.003621065 | 0.115469589 | 0.863165916 | 0.01774343 | 3.67E-15 | Northern |
| UG_1724 | 0.004694697 | 0.114334219 | 0.857103187 | 0.023867898 | 2.51E-14 | Northern |
| UG_1732 | 0.003809604 | 0.13046595 | 0.850901315 | 0.014823131 | 4.33E-15 | Northern |
| UG_1740 | 0.004283548 | 0.118156193 | 0.85721837 | 0.020341889 | 1.21E-14 | Northern |
| UG_1748 | 0.005708495 | 0.164076547 | 0.815636173 | 0.014578785 | 5.84E-14 | Northern |
| UG_1756 | 0.003343652 | 0.069265981 | 0.886029959 | 0.041360408 | 5.50E-15 | Northern |
| UG_1764 | 0.005531445 | 0.452739107 | 0.540506818 | 0.001222631 | 2.90E-14 | Northern |
| UG_1787 | 0.044788736 | 0.66521369 | 0.287671946 | 0.002325034 | 5.93E-07 | Northern |
| UG_1859 | 0.003776537 | 0.115322096 | 0.862291948 | 0.018609419 | 5.01E-15 | Northern |
| UG_1867 | 0.004254667 | 0.18490732 | 0.802546057 | 0.008291955 | 5.82E-15 | Northern |
| UG_1875 | 0.004175888 | 0.126851837 | 0.85169191 | 0.017280365 | 8.88E-15 | Northern |
| UG_1795 | 0.002786479 | 0.61580396 | 0.38122178 | 0.000187781 | 4.49E-16 | Northern |
| UG_1803 | 0.002559491 | 0.613576079 | 0.38369079 | 0.00017364 | 2.39E-16 | Northern |
| UG_1811 | 0.004657099 | 0.146100954 | 0.834486669 | 0.014755278 | 1.57E-14 | Northern |
| UG_1819 | 0.002436617 | 0.596731892 | 0.400646385 | 0.000185106 | 1.49E-16 | Northern |
| UG_1827 | 0.003289623 | 0.108524472 | 0.870190409 | 0.017995496 | 2.03E-15 | Northern |
| UG_1835 | 0.004351485 | 0.11237406 | 0.860531587 | 0.022742868 | 1.48E-14 | Northern |
| UG_1843 | 0.004258431 | 0.145679095 | 0.83659124 | 0.013471234 | 8.20E-15 | Northern |
| UG_1851 | 0.003755823 | 0.04166354 | 0.847965396 | 0.106615241 | 4.47E-14 | Northern |
| UG_1788 | 0.004152758 | 0.115131365 | 0.860046748 | 0.020669129 | 1.01E-14 | Northern |
| UG_1860 | 0.0047539 | 0.125267853 | 0.849635234 | 0.020343013 | 2.34E-14 | Northern |
| UG_1868 | 0.006092068 | 0.377590635 | 0.614015241 | 0.002302056 | 5.08E-14 | Northern |
| UG_1876 | 0.004449986 | 0.107715263 | 0.862624441 | 0.02521031 | 1.89E-14 | Northern |
| UG_1796 | 0.00296715 | 0.618444263 | 0.378391034 | 0.000197553 | 7.22E-16 | Northern |
| UG_1804 | 0.000548317 | 0.002146757 | 0.385949516 | 0.61135541 | 5.72E-16 | Northern |
| UG_1812 | 0.006002151 | 0.424835119 | 0.567541726 | 0.001621004 | 4.90E-14 | Northern |
| UG_1820 | 0.002882635 | 0.555436821 | 0.44138404 | 0.000296504 | 3.92E-16 | Northern |
| UG_1828 | 0.003861501 | 0.119477422 | 0.858843222 | 0.017817855 | 5.54E-15 | Northern |
| UG_1836 | 0.005039912 | 0.125468475 | 0.847905797 | 0.021585816 | 3.59E-14 | Northern |

| | | | | | | |
|---|---|---|---|---|---|---|
| UG_1844 | 0.004107484 | 0.12122514 | 0.856147397 | 0.018519979 | 8.50E-15 | Northern |
| UG_1852 | 0.002456171 | 0.038886532 | 0.879344683 | 0.079312614 | 2.09E-15 | Northern |
| UG_1789 | 0.029005229 | 0.366906848 | 0.591034906 | 0.013053012 | 4.83E-09 | Northern |
| UG_1861 | 0.006016194 | 0.111801403 | 0.849866302 | 0.032316102 | 1.63E-13 | Northern |
| UG_1869 | 0.004421872 | 0.161768078 | 0.822429726 | 0.011380325 | 9.24E-15 | Northern |
| UG_1877 | 0.005218089 | 0.113975494 | 0.853953517 | 0.0268529 | 5.49E-14 | Northern |
| UG_1797 | 0.002215982 | 0.023643789 | 0.822816746 | 0.151323483 | 3.75E-15 | Northern |
| UG_1805 | 0.00497203 | 0.093429208 | 0.864844312 | 0.03675445 | 5.61E-14 | Northern |
| UG_1813 | 0.003218616 | 0.641259138 | 0.355338881 | 0.000183364 | 1.55E-15 | Northern |
| UG_1821 | 0.010806814 | 0.517444184 | 0.470116181 | 0.001632821 | 4.82E-12 | Northern |
| UG_1829 | 0.005724423 | 0.652668337 | 0.341289999 | 0.000317241 | 1.12E-13 | Northern |
| UG_1837 | 0.00383451 | 0.117306426 | 0.860546006 | 0.018313058 | 5.44E-15 | Northern |
| UG_1845 | 0.001552786 | 0.012355828 | 0.727571375 | 0.258520011 | 1.90E-15 | Northern |
| UG_1853 | 0.005704298 | 0.10967258 | 0.852950533 | 0.03167259 | 1.14E-13 | Northern |
| UG_1790 | 0.002390686 | 0.594748045 | 0.402677501 | 0.000183768 | 1.28E-16 | Northern |
| UG_1862 | 0.003592405 | 0.128269961 | 0.853761251 | 0.014376383 | 2.90E-15 | Northern |
| UG_1870 | 0.005794171 | 0.442896146 | 0.549933229 | 0.001376453 | 3.96E-14 | Northern |
| UG_1878 | 0.000125666 | 0.000166679 | 0.087484719 | 0.912222936 | 1.36E-15 | Northern |
| UG_1798 | 0.003171441 | 0.568073291 | 0.428453018 | 0.000302251 | 8.40E-16 | Northern |
| UG_1806 | 0.01390059 | 0.520496228 | 0.463509624 | 0.002093558 | 3.10E-11 | Northern |
| UG_1814 | 0.029960187 | 0.371959551 | 0.585080804 | 0.012999452 | 6.18E-09 | Northern |
| UG_1822 | 0.004365712 | 0.530797369 | 0.464282234 | 0.000554685 | 7.00E-15 | Northern |
| UG_1830 | 0.004164504 | 0.126412404 | 0.852077891 | 0.017345201 | 8.76E-15 | Northern |
| UG_1838 | 0.005782746 | 0.105010219 | 0.854413166 | 0.034793869 | 1.37E-13 | Northern |
| UG_1846 | 0.002640378 | 0.073298235 | 0.894832515 | 0.029228873 | 8.54E-16 | Northern |
| UG_1854 | 0.000561592 | 0.001811189 | 0.332992886 | 0.664634333 | 1.78E-15 | Northern |
| UG_1791 | 0.002390015 | 0.60226377 | 0.395171913 | 0.000174302 | 1.34E-16 | Northern |
| UG_1863 | 0.003156028 | 0.098540602 | 0.877682768 | 0.020620601 | 1.78E-15 | Northern |
| UG_1871 | 9.25E-05 | 0.000109191 | 0.069448156 | 0.930350196 | 9.40E-16 | Northern |

| | | | | | | |
|---|---|---|---|---|---|---|
| UG_1879 | 8.79E-05 | 0.000104176 | 0.068407722 | 0.931400191 | 7.68E-16 | Northern |
| UG_1799 | 0.007988089 | 0.128325981 | 0.830109805 | 0.033576125 | 1.03E-12 | Northern |
| UG_1807 | 0.003615083 | 0.11865523 | 0.860913254 | 0.016816433 | 3.46E-15 | Northern |
| UG_1815 | 0.003282924 | 0.648305751 | 0.348233349 | 0.000177976 | 1.90E-15 | Northern |
| UG_1823 | 0.057573166 | 0.389048231 | 0.531988287 | 0.021389398 | 9.18E-07 | Northern |
| UG_1831 | 0.004196157 | 0.107955082 | 0.864261743 | 0.023587018 | 1.22E-14 | Northern |
| UG_1839 | 0.002444776 | 0.044847231 | 0.889993794 | 0.062714198 | 1.42E-15 | Northern |
| UG_1847 | 0.003709923 | 0.120428309 | 0.859052819 | 0.01680895 | 4.08E-15 | Northern |
| UG_1855 | 0.003374596 | 0.101778336 | 0.873985654 | 0.020861413 | 2.75E-15 | Northern |
| UG_1792 | 0.003241562 | 0.635353706 | 0.361211812 | 0.00019292 | 1.56E-15 | Northern |
| UG_1864 | 0.006170798 | 0.374815971 | 0.616631121 | 0.00238211 | 5.56E-14 | Northern |
| UG_1872 | 0.004781652 | 0.115104253 | 0.856081483 | 0.024032613 | 2.84E-14 | Northern |
| UG_1880 | 0.005285043 | 0.110652127 | 0.855304438 | 0.028758392 | 6.37E-14 | Northern |
| UG_1800 | 0.005596514 | 0.17038983 | 0.810793792 | 0.013219864 | 4.79E-14 | Northern |
| UG_1808 | 0.00306199 | 0.609913718 | 0.386806985 | 0.000217307 | 8.52E-16 | Northern |
| UG_1816 | 0.011260919 | 0.559181625 | 0.428274564 | 0.001282892 | 8.05E-12 | Northern |
| UG_1824 | 0.00407652 | 0.128808699 | 0.850767057 | 0.016347724 | 7.26E-15 | Northern |
| UG_1832 | 0.004084279 | 0.148805844 | 0.834766192 | 0.012343685 | 5.85E-15 | Northern |
| UG_1840 | 0.003107351 | 0.113569612 | 0.867798528 | 0.015524509 | 1.24E-15 | Northern |
| UG_1848 | 0.005172936 | 0.132473844 | 0.842356247 | 0.019996972 | 3.96E-14 | Northern |
| UG_1856 | 0.005931169 | 0.137242032 | 0.835218368 | 0.021608431 | 1.02E-13 | Northern |
| UG_1793 | 0.004737674 | 0.496158183 | 0.498336684 | 0.000767458 | 1.09E-14 | Northern |
| UG_1865 | 0.004763299 | 0.081476721 | 0.868920671 | 0.044839309 | 5.40E-14 | Northern |
| UG_1873 | 0.004037283 | 0.158543236 | 0.826683579 | 0.010735902 | 4.90E-15 | Northern |
| UG_1801 | 0.00829616 | 0.530357329 | 0.460222832 | 0.001123679 | 7.43E-13 | Northern |
| UG_1809 | 0.06818222 | 0.472491996 | 0.44553045 | 0.01379109 | 4.24E-06 | Northern |
| UG_1817 | 0.00556787 | 0.460241466 | 0.533020746 | 0.001169918 | 3.11E-14 | Northern |
| UG_1825 | 0.002599313 | 0.02523815 | 0.813654683 | 0.158507855 | 1.08E-14 | Northern |
| UG_1833 | 0.00317718 | 0.106556807 | 0.87232408 | 0.017941934 | 1.63E-15 | Northern |

| | | | | | | |
|---|---|---|---|---|---|---|
| UG_1841 | 0.000124725 | 0.000157672 | 0.083092554 | 0.916625049 | 1.79E-15 | Northern |
| UG_1849 | 0.002669263 | 0.040492525 | 0.876197911 | 0.0806403 | 3.55E-15 | Northern |
| UG_1857 | 0.001919161 | 0.013287234 | 0.708422376 | 0.276371229 | 8.34E-15 | Northern |
| UG_1794 | 0.004146984 | 0.1305808 | 0.849054234 | 0.016217981 | 8.04E-15 | Northern |
| UG_1866 | 0.000549462 | 0.001257856 | 0.246506683 | 0.751686 | 1.13E-14 | Northern |
| UG_1874 | 0.003704895 | 0.110648684 | 0.865931681 | 0.01971474 | 4.68E-15 | Northern |
| UG_1802 | 0.000750806 | 0.003831706 | 0.505456765 | 0.489960722 | 5.35E-16 | Northern |
| UG_1810 | 0.008868334 | 0.205689191 | 0.770773325 | 0.01466915 | 1.09E-12 | Northern |
| UG_1818 | 0.003250823 | 0.654942528 | 0.341639024 | 0.000167625 | 1.87E-15 | Northern |
| UG_1826 | 0.00273436 | 0.026557862 | 0.816401705 | 0.154306072 | 1.38E-14 | Northern |
| UG_1834 | 0.004085762 | 0.129509432 | 0.850188191 | 0.016216615 | 7.31E-15 | Northern |
| UG_1842 | 0.007423934 | 0.138195677 | 0.827351142 | 0.027029248 | 5.27E-13 | Northern |
| UG_1850 | 0.003485234 | 0.138781902 | 0.845805854 | 0.01192701 | 2.05E-15 | Northern |
| UG_1858 | 0.006851998 | 0.081107305 | 0.846912351 | 0.065128346 | 8.33E-13 | Northern |
| UG_1881 | 9.70E-05 | 0.000112758 | 0.069531052 | 0.930259153 | 1.24E-15 | Northern |
| UG_1953 | 0.000103775 | 0.000144094 | 0.085471966 | 0.914280165 | 5.14E-16 | North_western |
| UG_1961 | 0.000110867 | 0.000176244 | 0.100364794 | 0.899348094 | 2.80E-16 | North_western |
| UG_1969 | 0.000122326 | 0.000170866 | 0.09129103 | 0.908415779 | 8.95E-16 | North_western |
| UG_1889 | 0.000536264 | 0.002291889 | 0.410076992 | 0.587094856 | 3.31E-16 | Northern |
| UG_1897 | 0.000618864 | 0.002320919 | 0.386016622 | 0.611043594 | 1.16E-15 | Northern |
| UG_1905 | 0.00010854 | 0.000130034 | 0.07475863 | 0.925002797 | 1.53E-15 | Northern |
| UG_1913 | 0.004492669 | 0.126737837 | 0.850044539 | 0.018724954 | 1.52E-14 | Northern |
| UG_1921 | 0.004501934 | 0.100706905 | 0.865888003 | 0.028903157 | 2.33E-14 | Northern |
| UG_1929 | 0.004402281 | 0.130812795 | 0.847548629 | 0.017236296 | 1.24E-14 | Northern |
| UG_1937 | 0.000437049 | 0.000919486 | 0.211962688 | 0.786680777 | 7.80E-15 | North_western |
| UG_1945 | 9.89E-05 | 0.0001325 | 0.080979792 | 0.918788798 | 5.41E-16 | North_western |
| UG_1882 | 0.003743288 | 0.119701264 | 0.859385728 | 0.01716972 | 4.40E-15 | Northern |
| UG_1954 | 0.124580765 | 0.58691633 | 0.278468033 | 0.008627771 | 0.001407102 | North_western |
| UG_1962 | 0.000111275 | 0.000173854 | 0.098763257 | 0.900951614 | 3.16E-16 | North_western |

| | | | | | | |
|---|---|---|---|---|---|---|
| UG_1970 | 7.81E-05 | 0.000104664 | 0.074313185 | 0.925504084 | 2.30E-16 | North_western |
| UG_1890 | 0.00483961 | 0.162525658 | 0.820201475 | 0.012433257 | 1.77E-14 | Northern |
| UG_1898 | 0.000564778 | 0.002408032 | 0.415049777 | 0.581977413 | 4.12E-16 | Northern |
| UG_1906 | 0.004180913 | 0.140320681 | 0.841279561 | 0.014218846 | 7.60E-15 | Northern |
| UG_1914 | 0.00447734 | 0.22901337 | 0.760969223 | 0.005540067 | 6.64E-15 | Northern |
| UG_1922 | 0.003647097 | 0.041463911 | 0.850372642 | 0.104516349 | 3.61E-14 | Northern |
| UG_1930 | 0.003939405 | 0.130590986 | 0.850128319 | 0.015341289 | 5.53E-15 | Northern |
| UG_1938 | 0.000139596 | 0.000217621 | 0.106927829 | 0.892714954 | 7.27E-16 | North_western |
| UG_1946 | 0.016348729 | 0.097584399 | 0.777476363 | 0.108590509 | 4.20E-10 | North_western |
| UG_1883 | 0.003748494 | 0.113472139 | 0.863743622 | 0.019035746 | 4.88E-15 | Northern |
| UG_1955 | 9.81E-05 | 0.000129721 | 0.079651319 | 0.920120812 | 5.73E-16 | North_western |
| UG_1963 | 0.000140579 | 0.000238856 | 0.116796132 | 0.882824433 | 4.38E-16 | North_western |
| UG_1971 | 0.087382378 | 0.670013961 | 0.238864151 | 0.003585202 | 0.000154308 | North_western |
| UG_1891 | 0.004447806 | 0.036568261 | 0.809069749 | 0.149914184 | 2.42E-13 | Northern |
| UG_1899 | 0.000337803 | 0.000888053 | 0.238679861 | 0.760094283 | 8.07E-16 | Northern |
| UG_1907 | 0.004253256 | 0.096226707 | 0.86993022 | 0.029589818 | 1.67E-14 | Northern |
| UG_1915 | 0.004482132 | 0.039240458 | 0.819722615 | 0.136554795 | 2.08E-13 | Northern |
| UG_1923 | 0.004359528 | 0.101722507 | 0.866490068 | 0.027427897 | 1.80E-14 | Northern |
| UG_1931 | 0.004577939 | 0.103627312 | 0.863885037 | 0.027909713 | 2.50E-14 | Northern |
| UG_1939 | 8.75E-05 | 0.000108717 | 0.07170928 | 0.92809454 | 5.58E-16 | North_western |
| UG_1947 | 0.016371493 | 0.101220603 | 0.779871136 | 0.102536768 | 3.88E-10 | North_western |
| UG_1884 | 0.001301519 | 0.008971405 | 0.667298001 | 0.322429075 | 1.49E-15 | Northern |
| UG_1956 | 0.00010052 | 0.000153544 | 0.093054588 | 0.906691348 | 2.52E-16 | North_western |
| UG_1964 | 0.083015843 | 0.672210521 | 0.241251104 | 0.003420842 | 0.000101691 | North_western |
| UG_1972 | 7.63E-05 | 9.26E-05 | 0.066538281 | 0.933292815 | 3.98E-16 | North_western |
| UG_1892 | 0.005951649 | 0.092668606 | 0.856494743 | 0.044885002 | 2.17E-13 | Northern |
| UG_1900 | 0.002702194 | 0.620692214 | 0.376430295 | 0.000175297 | 3.73E-16 | Northern |
| UG_1908 | 0.006278446 | 0.074118378 | 0.850058202 | 0.069544974 | 5.29E-13 | Northern |
| UG_1916 | 0.000472136 | 0.001924992 | 0.382105722 | 0.61549715 | 2.55E-16 | Northern |

| | | | | | | |
|---|---|---|---|---|---|---|
| UG_1924 | 0.000549209 | 0.002181846 | 0.390482923 | 0.606786022 | 5.33E-16 | Northern |
| UG_1932 | 0.004313364 | 0.113130706 | 0.860307955 | 0.022247975 | 1.37E-14 | Northern |
| UG_1940 | 0.096057335 | 0.401681708 | 0.472612262 | 0.02959304 | 5.57E-05 | North_western |
| UG_1948 | 0.015748883 | 0.097944023 | 0.781879264 | 0.10442783 | 3.11E-10 | North_western |
| UG_1885 | 0.00820184 | 0.092000698 | 0.83708936 | 0.062708102 | 2.44E-12 | Northern |
| UG_1957 | 0.000123087 | 0.000169438 | 0.090156836 | 0.909550639 | 1.00E-15 | North_western |
| UG_1965 | 0.000100593 | 0.000127642 | 0.07709881 | 0.922672954 | 8.09E-16 | North_western |
| UG_1973 | 9.51E-05 | 0.000121991 | 0.076367333 | 0.923415572 | 6.18E-16 | North_western |
| UG_1893 | 0.00640646 | 0.096616813 | 0.852044049 | 0.044932678 | 3.45E-13 | Northern |
| UG_1901 | 0.007314189 | 0.122459757 | 0.836780267 | 0.033445788 | 5.84E-13 | Northern |
| UG_1909 | 0.002980354 | 0.123894111 | 0.86057143 | 0.012554104 | 7.87E-16 | Northern |
| UG_1917 | 0.006299492 | 0.099936704 | 0.85218658 | 0.041577224 | 2.84E-13 | Northern |
| UG_1925 | 0.004427003 | 0.039205652 | 0.821110712 | 0.135256633 | 1.90E-13 | Northern |
| UG_1933 | 0.004924489 | 0.100320462 | 0.862770807 | 0.031984242 | 4.55E-14 | Northern |
| UG_1941 | 0.000101158 | 0.000131179 | 0.078989574 | 0.920778089 | 7.20E-16 | North_western |
| UG_1949 | 0.016689365 | 0.098906374 | 0.776159022 | 0.108245239 | 4.77E-10 | North_western |
| UG_1886 | 0.003768233 | 0.189044336 | 0.800256195 | 0.006931235 | 2.34E-15 | Northern |
| UG_1958 | 9.70E-05 | 0.000125214 | 0.07743921 | 0.922338609 | 6.35E-16 | North_western |
| UG_1966 | 0.000102997 | 0.000154296 | 0.092052862 | 0.907689845 | 3.11E-16 | North_western |
| UG_1974 | 0.000124982 | 0.000179765 | 0.094766627 | 0.904928626 | 8.05E-16 | North_western |
| UG_1894 | 0.0005537 | 0.002234103 | 0.396030099 | 0.601182098 | 5.07E-16 | Northern |
| UG_1902 | 0.003031965 | 0.02798288 | 0.812199372 | 0.156785783 | 2.65E-14 | Northern |
| UG_1910 | 0.004448261 | 0.107708023 | 0.86264062 | 0.025203097 | 1.88E-14 | Northern |
| UG_1918 | 0.004116802 | 0.11894921 | 0.857685614 | 0.019248374 | 8.93E-15 | Northern |
| UG_1926 | 0.004023628 | 0.09506301 | 0.872378531 | 0.02853483 | 1.14E-14 | Northern |
| UG_1934 | 0.004269805 | 0.105904586 | 0.864924013 | 0.024901597 | 1.44E-14 | Northern |
| UG_1942 | 0.000107255 | 0.000173759 | 0.101076006 | 0.89864298 | 2.21E-16 | North_western |
| UG_1950 | 0.016323893 | 0.097733767 | 0.777761333 | 0.108181007 | 4.14E-10 | North_western |
| UG_1887 | 0.001526499 | 0.012669252 | 0.737581407 | 0.248222843 | 1.50E-15 | Northern |

| | | | | | | |
|---|---|---|---|---|---|---|
| UG_1959 | 0.000171895 | 0.000278516 | 0.119693256 | 0.879856333 | 1.22E-15 | North_western |
| UG_1967 | 0.000135313 | 0.000218691 | 0.109617818 | 0.890028178 | 5.19E-16 | North_western |
| UG_1895 | 0.004367479 | 0.046998221 | 0.846626154 | 0.102008146 | 1.03E-13 | Northern |
| UG_1903 | 0.004508731 | 0.096361292 | 0.867748757 | 0.03138122 | 2.56E-14 | Northern |
| UG_1911 | 0.004384149 | 0.103781591 | 0.865244974 | 0.026589286 | 1.81E-14 | Northern |
| UG_1919 | 0.004843288 | 0.093471438 | 0.865948709 | 0.035736565 | 4.62E-14 | Northern |
| UG_1927 | 0.00010491 | 0.000124293 | 0.072993398 | 0.926777399 | 1.45E-15 | Northern |
| UG_1935 | 0.004420897 | 0.131101939 | 0.84723627 | 0.017240894 | 1.28E-14 | Northern |
| UG_1943 | 0.00012032 | 0.000161258 | 0.087024379 | 0.912694043 | 1.09E-15 | North_western |
| UG_1951 | 0.016258834 | 0.098099797 | 0.778482601 | 0.107158767 | 3.98E-10 | North_western |
| UG_1888 | 0.001194221 | 0.009005971 | 0.685742655 | 0.304057153 | 7.12E-16 | Northern |
| UG_1960 | 0.000100536 | 0.00013415 | 0.081138559 | 0.918626755 | 5.88E-16 | North_western |
| UG_1968 | 0.000117445 | 0.000176939 | 0.097087894 | 0.902617721 | 4.82E-16 | North_western |
| UG_1896 | 0.005120524 | 0.128164767 | 0.845635525 | 0.021079184 | 3.89E-14 | Northern |
| UG_1904 | 0.005063286 | 0.103409762 | 0.860376082 | 0.03115087 | 5.27E-14 | Northern |
| UG_1912 | 0.000444299 | 0.001171771 | 0.261234022 | 0.737149907 | 2.23E-15 | Northern |
| UG_1920 | 0.004434329 | 0.040268475 | 0.824983317 | 0.130313878 | 1.78E-13 | Northern |
| UG_1928 | 0.000103021 | 0.000122086 | 0.072525839 | 0.927249055 | 1.36E-15 | Northern |
| UG_1936 | 0.002241106 | 0.020346358 | 0.790684596 | 0.18672794 | 6.59E-15 | Northern |
| UG_1944 | 0.000120922 | 0.000155671 | 0.083692068 | 0.91603134 | 1.43E-15 | North_western |
| UG_1952 | 8.67E-05 | 0.000114796 | 0.076239782 | 0.923558702 | 3.64E-16 | North_western |
| UG_1975 | 0.015963796 | 0.097078451 | 0.779752201 | 0.107205552 | 3.54E-10 | North_western |
| UG_2047 | 0.004569193 | 0.161405294 | 0.822179312 | 0.011846201 | 1.18E-14 | Northern |
| UG_2055 | 0.000544729 | 0.002133102 | 0.385305645 | 0.612016524 | 5.56E-16 | Northern |
| UG_2063 | 0.056227347 | 0.66936534 | 0.271700033 | 0.002703472 | 3.81E-06 | Northern |
| UG_1983 | 8.52E-05 | 0.000104235 | 0.069884795 | 0.929925793 | 5.60E-16 | North_western |
| UG_1991 | 0.000102177 | 0.000161093 | 0.096642608 | 0.903094122 | 2.20E-16 | North_western |
| UG_1999 | 8.73E-05 | 0.000105763 | 0.069772707 | 0.930034183 | 6.55E-16 | North_western |
| UG_2007 | 0.002365541 | 0.031642277 | 0.860708405 | 0.105283777 | 2.69E-15 | Northern |

| | | | | | | |
|---|---|---|---|---|---|---|
| UG_2015 | 0.00307768 | 0.601592398 | 0.395098195 | 0.000231727 | 8.35E-16 | Northern |
| UG_2023 | 0.003867979 | 0.113756611 | 0.862780706 | 0.019594705 | 6.11E-15 | Northern |
| UG_2031 | 0.000559355 | 0.002315734 | 0.40494655 | 0.592178361 | 4.62E-16 | Northern |
| UG_2039 | 0.006140895 | 0.120297466 | 0.844751512 | 0.028810127 | 1.66E-13 | Northern |
| UG_1976 | 0.000165533 | 0.00025227 | 0.111119059 | 0.888463138 | 1.55E-15 | North_western |
| UG_2048 | 0.00363349 | 0.129863476 | 0.852293072 | 0.014209962 | 3.09E-15 | Northern |
| UG_2056 | 0.003027055 | 0.074925182 | 0.889580539 | 0.032467224 | 2.24E-15 | Northern |
| UG_2064 | 0.003156697 | 0.643978393 | 0.35268897 | 0.000175941 | 1.38E-15 | Northern |
| UG_1984 | 9.60E-05 | 0.000134694 | 0.083934373 | 0.915834885 | 3.66E-16 | North_western |
| UG_1992 | 9.56E-05 | 0.000106579 | 0.066282515 | 0.933515339 | 1.52E-15 | North_western |
| UG_2000 | 0.000526462 | 0.002171113 | 0.397183468 | 0.600118958 | 3.70E-16 | Northern |
| UG_2008 | 0.058129184 | 0.396583425 | 0.524834037 | 0.020452355 | 9.98E-07 | Northern |
| UG_2016 | 0.003707607 | 0.111877558 | 0.865091632 | 0.019323203 | 4.62E-15 | Northern |
| UG_2024 | 0.000639591 | 0.002721226 | 0.428212271 | 0.568426913 | 6.86E-16 | Northern |
| UG_2032 | 0.000566048 | 0.002108538 | 0.374598624 | 0.62272679 | 8.44E-16 | Northern |
| UG_2040 | 0.00483273 | 0.139188687 | 0.839089113 | 0.01688947 | 2.22E-14 | Northern |
| UG_1977 | 0.000192235 | 0.00029652 | 0.118624908 | 0.880886336 | 2.48E-15 | North_western |
| UG_2049 | 8.68E-05 | 9.89E-05 | 0.065430271 | 0.934384015 | 9.36E-16 | Northern |
| UG_2057 | 0.004374275 | 0.143511462 | 0.837829268 | 0.014284994 | 1.02E-14 | Northern |
| UG_2065 | 0.004281335 | 0.134598409 | 0.845294431 | 0.015825824 | 9.67E-15 | Northern |
| UG_1985 | 0.000106723 | 0.000168842 | 0.098516736 | 0.901207698 | 2.51E-16 | North_western |
| UG_1993 | 0.000123506 | 0.000172096 | 0.091385039 | 0.908319359 | 9.40E-16 | North_western |
| UG_2001 | 0.003868794 | 0.124055855 | 0.855462362 | 0.016612989 | 5.27E-15 | Northern |
| UG_2009 | 0.003390189 | 0.60419919 | 0.392157436 | 0.000253185 | 1.71E-15 | Northern |
| UG_2017 | 0.002442263 | 0.092106658 | 0.887713011 | 0.017738068 | 3.10E-16 | Northern |
| UG_2025 | 0.000105691 | 0.000125451 | 0.073331549 | 0.92643731 | 1.47E-15 | Northern |
| UG_2033 | 0.000768789 | 0.003301222 | 0.451918745 | 0.544011244 | 1.38E-15 | Northern |
| UG_2041 | 0.004202962 | 0.133515292 | 0.846522325 | 0.015759421 | 8.56E-15 | Northern |
| UG_1978 | 0.000214536 | 0.000356419 | 0.132692926 | 0.866736118 | 2.34E-15 | North_western |

| | | | | | | |
|---|---|---|---|---|---|---|
| UG_2050 | 0.003330811 | 0.133096085 | 0.851248446 | 0.012324658 | 1.58E-15 | Northern |
| UG_2058 | 0.00378029 | 0.13274653 | 0.849260728 | 0.014212453 | 3.98E-15 | Northern |
| UG_2066 | 0.003525437 | 0.47393284 | 0.521897873 | 0.00064385 | 1.18E-15 | Northern |
| UG_1986 | 0.004122355 | 0.112959457 | 0.861659991 | 0.021258197 | 9.87E-15 | North_western |
| UG_1994 | 0.000100559 | 0.000143405 | 0.086816849 | 0.912939187 | 3.88E-16 | North_western |
| UG_2002 | 0.003433684 | 0.118784008 | 0.861908003 | 0.015874305 | 2.37E-15 | Northern |
| UG_2010 | 0.004142817 | 0.140360167 | 0.841426079 | 0.014070937 | 7.11E-15 | Northern |
| UG_2018 | 0.003119046 | 0.626555099 | 0.370128855 | 0.000197 | 1.10E-15 | Northern |
| UG_2026 | 0.003044933 | 0.09115546 | 0.882881997 | 0.02291761 | 1.59E-15 | Northern |
| UG_2034 | 0.003470458 | 0.08257487 | 0.882444849 | 0.031509824 | 5.03E-15 | Northern |
| UG_2042 | 0.004725809 | 0.118034481 | 0.854604155 | 0.022635556 | 2.49E-14 | Northern |
| UG_1979 | 0.000151221 | 0.000195656 | 0.091251939 | 0.908401184 | 3.10E-15 | North_western |
| UG_2051 | 0.003701262 | 0.119001217 | 0.860144209 | 0.017153312 | 4.09E-15 | Northern |
| UG_2059 | 0.000120047 | 0.000148354 | 0.080068894 | 0.919662706 | 1.80E-15 | Northern |
| UG_2067 | 0.00024715 | 0.000807676 | 0.26218578 | 0.736759394 | 7.08E-17 | Northern |
| UG_1987 | 9.32E-05 | 0.000119547 | 0.075780377 | 0.924006835 | 5.77E-16 | North_western |
| UG_1995 | 0.000119939 | 0.000162848 | 0.088075008 | 0.911642205 | 9.95E-16 | North_western |
| UG_2003 | 0.003499284 | 0.076903055 | 0.883512887 | 0.036084775 | 6.18E-15 | Northern |
| UG_2011 | 0.004526965 | 0.134460821 | 0.844172694 | 0.01683952 | 1.46E-14 | Northern |
| UG_2019 | 0.003353896 | 0.650265626 | 0.346200867 | 0.000179611 | 2.26E-15 | Northern |
| UG_2027 | 9.87E-05 | 0.000118003 | 0.072037592 | 0.927745699 | 1.10E-15 | Northern |
| UG_2035 | 0.004820088 | 0.162954734 | 0.819912025 | 0.012313153 | 1.71E-14 | Northern |
| UG_2043 | 0.004317431 | 0.142069074 | 0.839246035 | 0.014367459 | 9.43E-15 | Northern |
| UG_1980 | 0.000142188 | 0.000186479 | 0.090464542 | 0.909206791 | 2.28E-15 | North_western |
| UG_2052 | 0.003368857 | 0.11557963 | 0.864666808 | 0.016384705 | 2.16E-15 | Northern |
| UG_2060 | 0.004019934 | 0.141197926 | 0.841321124 | 0.013461016 | 5.65E-15 | Northern |
| UG_2068 | 0.003622141 | 0.105758916 | 0.869670491 | 0.020948453 | 4.30E-15 | Northern |
| UG_1988 | 0.000131052 | 0.000166538 | 0.085056626 | 0.914645784 | 2.07E-15 | North_western |
| UG_1996 | 0.0001368 | 0.000215845 | 0.107438935 | 0.892208421 | 6.27E-16 | North_western |

| | | | | | | |
|---|---|---|---|---|---|---|
| UG_2004 | 0.057680895 | 0.395068894 | 0.526718015 | 0.020531259 | 9.38E-07 | Northern |
| UG_2012 | 0.003481316 | 0.082167617 | 0.882455837 | 0.03189523 | 5.20E-15 | Northern |
| UG_2020 | 0.004521804 | 0.159496726 | 0.823984085 | 0.011997385 | 1.11E-14 | Northern |
| UG_2028 | 0.000536027 | 0.002232544 | 0.402139265 | 0.595092164 | 3.78E-16 | Northern |
| UG_2036 | 0.003511745 | 0.103532549 | 0.871867904 | 0.021087802 | 3.56E-15 | Northern |
| UG_2044 | 9.08E-05 | 0.000104284 | 0.067029418 | 0.932775481 | 1.05E-15 | Northern |
| UG_1981 | 0.000121916 | 0.000179274 | 0.096034527 | 0.903664283 | 6.41E-16 | North_western |
| UG_2053 | 0.00420459 | 0.11886236 | 0.857216921 | 0.019716129 | 1.04E-14 | Northern |
| UG_2061 | 0.004852239 | 0.149445636 | 0.830954202 | 0.014747923 | 2.05E-14 | Northern |
| UG_1989 | 6.55E-05 | 7.94E-05 | 0.062801101 | 0.937054059 | 2.32E-16 | North_western |
| UG_1997 | 0.000110833 | 0.000164852 | 0.093853271 | 0.905871044 | 4.23E-16 | North_western |
| UG_2005 | 9.30E-05 | 0.000108828 | 0.068921057 | 0.930877066 | 1.02E-15 | Northern |
| UG_2013 | 0.000509304 | 0.002152582 | 0.401040568 | 0.596297546 | 2.85E-16 | Northern |
| UG_2021 | 0.000549383 | 0.002164534 | 0.388037913 | 0.60924817 | 5.57E-16 | Northern |
| UG_2029 | 0.040879438 | 0.256906383 | 0.659505778 | 0.042708323 | 7.80E-08 | Northern |
| UG_2037 | 0.003591353 | 0.078389294 | 0.882180105 | 0.035839247 | 7.20E-15 | Northern |
| UG_2045 | 0.009456092 | 0.190651101 | 0.78148594 | 0.018406867 | 1.92E-12 | Northern |
| UG_1982 | 9.65E-05 | 0.000121759 | 0.075491117 | 0.924290614 | 7.23E-16 | North_western |
| UG_2054 | 0.004995677 | 0.16914943 | 0.813988245 | 0.011866648 | 2.11E-14 | Northern |
| UG_2062 | 0.004043781 | 0.098424367 | 0.870616157 | 0.026915695 | 1.10E-14 | Northern |
| UG_1990 | 0.000123899 | 0.000187794 | 0.099586655 | 0.900101652 | 5.63E-16 | North_western |
| UG_1998 | 0.000622129 | 0.001388649 | 0.250759358 | 0.747229864 | 2.08E-14 | North_western |
| UG_2006 | 0.004068694 | 0.149294137 | 0.83442451 | 0.012212659 | 5.67E-15 | Northern |
| UG_2014 | 0.00348305 | 0.127203142 | 0.855184103 | 0.014129705 | 2.35E-15 | Northern |
| UG_2022 | 0.001810387 | 0.024324271 | 0.852264828 | 0.121600514 | 7.13E-16 | Northern |
| UG_2030 | 0.003168391 | 0.573047562 | 0.42349227 | 0.000291777 | 8.59E-16 | Northern |
| UG_2038 | 9.64E-05 | 0.000112129 | 0.069438302 | 0.93035318 | 1.20E-15 | Northern |
| UG_2046 | 0.004278552 | 0.127389526 | 0.850738078 | 0.017593844 | 1.05E-14 | Northern |
| UG_2069 | 0.003817175 | 0.116562946 | 0.861174041 | 0.018445838 | 5.32E-15 | Northern |

| | | | | | | |
|---|---|---|---|---|---|---|
| UG_2141 | 0.005624678 | 0.110454807 | 0.853117454 | 0.030803061 | 1.01E-13 | Northern |
| UG_2149 | 0.003495156 | 0.115636551 | 0.863835683 | 0.017032609 | 2.83E-15 | Northern |
| UG_2157 | 0.004470423 | 0.105138895 | 0.863892503 | 0.026498179 | 2.04E-14 | Northern |
| UG_2077 | 0.003087887 | 0.606331394 | 0.390355764 | 0.000224955 | 8.83E-16 | Northern |
| UG_2085 | 0.002907534 | 0.61330821 | 0.383583911 | 0.000200345 | 6.00E-16 | Northern |
| UG_2093 | 0.004194198 | 0.11979857 | 0.856634785 | 0.019372447 | 1.01E-14 | Northern |
| UG_2101 | 0.003509645 | 0.605528599 | 0.390701154 | 0.000260602 | 2.22E-15 | Northern |
| UG_2109 | 0.000693142 | 0.003244985 | 0.467865361 | 0.528196512 | 5.87E-16 | Northern |
| UG_2117 | 0.004479127 | 0.130266609 | 0.847551693 | 0.01770257 | 1.42E-14 | Northern |
| UG_2125 | 0.000832531 | 0.004938765 | 0.567724927 | 0.426503777 | 4.09E-16 | Northern |
| UG_2133 | 0.088536046 | 0.654790227 | 0.252388978 | 0.004142546 | 0.000142203 | Northern |
| UG_2070 | 0.003768072 | 0.110965056 | 0.865299105 | 0.019967766 | 5.27E-15 | Northern |
| UG_2142 | 0.005415324 | 0.090845615 | 0.861530186 | 0.042208874 | 1.12E-13 | Northern |
| UG_2150 | 0.00060057 | 0.002254918 | 0.383173018 | 0.613971493 | 1.02E-15 | Northern |
| UG_2158 | 9.57E-05 | 0.000106573 | 0.066227052 | 0.933570697 | 1.54E-15 | Northern |
| UG_2078 | 0.001821612 | 0.026388869 | 0.863336394 | 0.108453125 | 5.95E-16 | Northern |
| UG_2086 | 8.99E-05 | 0.000102068 | 0.066011624 | 0.933796421 | 1.09E-15 | Northern |
| UG_2094 | 0.003156643 | 0.609397311 | 0.387220479 | 0.000225567 | 1.06E-15 | Northern |
| UG_2102 | 0.00331734 | 0.596614381 | 0.399807592 | 0.000260687 | 1.39E-15 | Northern |
| UG_2110 | 0.003096044 | 0.635882956 | 0.360838324 | 0.000182676 | 1.12E-15 | Northern |
| UG_2118 | 0.003298648 | 0.118237263 | 0.863128947 | 0.015335141 | 1.78E-15 | Northern |
| UG_2126 | 0.000572404 | 0.00214564 | 0.37761438 | 0.619667576 | 8.53E-16 | Northern |
| UG_2134 | 0.003469443 | 0.122910146 | 0.858585894 | 0.015034517 | 2.42E-15 | Northern |
| UG_2071 | 0.006477186 | 0.115041616 | 0.845376205 | 0.033104993 | 2.66E-13 | Northern |
| UG_2143 | 0.003057238 | 0.066794527 | 0.88999083 | 0.040157405 | 3.06E-15 | Northern |
| UG_2151 | 0.000579082 | 0.00210129 | 0.369324591 | 0.627995037 | 1.06E-15 | Northern |
| UG_2159 | 0.00332366 | 0.606845297 | 0.389587914 | 0.00024313 | 1.51E-15 | Northern |
| UG_2079 | 0.004120308 | 0.49923321 | 0.496002079 | 0.000644404 | 4.01E-15 | Northern |
| UG_2087 | 0.001454885 | 0.011855226 | 0.728056234 | 0.258633654 | 1.29E-15 | Northern |

| | | | | | | |
|---|---|---|---|---|---|---|
| UG_2095 | 0.003541321 | 0.611175923 | 0.385029846 | 0.00025291 | 2.47E-15 | Northern |
| UG_2103 | 0.003380607 | 0.621746414 | 0.374650109 | 0.00022287 | 1.90E-15 | Northern |
| UG_2111 | 0.003166312 | 0.623731564 | 0.372897727 | 0.000204397 | 1.20E-15 | Northern |
| UG_2119 | 0.004623267 | 0.131509885 | 0.845884876 | 0.017981973 | 1.76E-14 | Northern |
| UG_2127 | 0.000101727 | 0.000120992 | 0.07245903 | 0.927318252 | 1.27E-15 | Northern |
| UG_2135 | 0.007074077 | 0.109864446 | 0.84359667 | 0.039464807 | 5.59E-13 | Northern |
| UG_2072 | 0.00298145 | 0.602547965 | 0.394248355 | 0.00022223 | 6.68E-16 | Northern |
| UG_2144 | 0.007453208 | 0.127599568 | 0.833367583 | 0.031579641 | 6.23E-13 | Northern |
| UG_2152 | 0.000562774 | 0.002238091 | 0.39341836 | 0.603780774 | 5.84E-16 | Northern |
| UG_2160 | 0.004396397 | 0.154608099 | 0.82860783 | 0.012387674 | 9.46E-15 | Northern |
| UG_2080 | 0.003741743 | 0.09164831 | 0.876358774 | 0.028251173 | 7.14E-15 | Northern |
| UG_2088 | 0.000673616 | 0.002781824 | 0.424673564 | 0.571870995 | 9.84E-16 | Northern |
| UG_2096 | 0.004031582 | 0.162587916 | 0.823193562 | 0.010186939 | 4.68E-15 | Northern |
| UG_2104 | 0.003072594 | 0.626304948 | 0.370428349 | 0.000194108 | 9.86E-16 | Northern |
| UG_2112 | 0.003457681 | 0.622341595 | 0.37397321 | 0.000227514 | 2.25E-15 | Northern |
| UG_2120 | 0.002202504 | 0.034951877 | 0.878407865 | 0.084437754 | 1.20E-15 | Northern |
| UG_2128 | 9.36E-05 | 0.000107274 | 0.067631117 | 0.932167976 | 1.19E-15 | Northern |
| UG_2136 | 0.003417918 | 0.646149152 | 0.350243889 | 0.00018904 | 2.50E-15 | Northern |
| UG_2073 | 0.003253794 | 0.611236875 | 0.385279095 | 0.000230236 | 1.34E-15 | Northern |
| UG_2145 | 0.000600363 | 0.002493889 | 0.41373813 | 0.583167618 | 6.02E-16 | Northern |
| UG_2153 | 0.003908192 | 0.123690217 | 0.855510636 | 0.016890956 | 5.71E-15 | Northern |
| UG_2161 | 0.003341714 | 0.607687425 | 0.388727721 | 0.00024314 | 1.58E-15 | Northern |
| UG_2081 | 0.003170931 | 0.587710506 | 0.408854686 | 0.000263877 | 9.46E-16 | Northern |
| UG_2089 | 0.002979729 | 0.570310125 | 0.426432351 | 0.000277795 | 5.42E-16 | Northern |
| UG_2097 | 0.003110529 | 0.075972659 | 0.888322906 | 0.032593906 | 2.66E-15 | Northern |
| UG_2105 | 0.00352124 | 0.126975116 | 0.855156214 | 0.01434743 | 2.55E-15 | Northern |
| UG_2113 | 0.00354811 | 0.645170999 | 0.351082477 | 0.000198414 | 3.25E-15 | Northern |
| UG_2121 | 0.003989151 | 0.128397336 | 0.851544263 | 0.01606925 | 6.23E-15 | Northern |
| UG_2129 | 0.002597084 | 0.093156583 | 0.885686422 | 0.01855991 | 4.75E-16 | Northern |

| | | | | | | |
|---|---|---|---|---|---|---|
| UG_2137 | 0.004812434 | 0.164349204 | 0.818756517 | 0.012081845 | 1.67E-14 | Northern |
| UG_2074 | 0.003191438 | 0.588725153 | 0.407819512 | 0.000263897 | 9.97E-16 | Northern |
| UG_2146 | 0.003199404 | 0.601901498 | 0.394657741 | 0.000241356 | 1.11E-15 | Northern |
| UG_2154 | 0.002840512 | 0.649639864 | 0.347369427 | 0.000150197 | 6.73E-16 | Northern |
| UG_2162 | 0.003359502 | 0.607454281 | 0.388941244 | 0.000244973 | 1.64E-15 | Northern |
| UG_2082 | 0.004583108 | 0.149312647 | 0.832213343 | 0.013890901 | 1.35E-14 | Northern |
| UG_2090 | 0.003781482 | 0.126551248 | 0.854068778 | 0.015598492 | 4.32E-15 | Northern |
| UG_2098 | 0.003649353 | 0.115651086 | 0.862860205 | 0.017839356 | 3.88E-15 | Northern |
| UG_2106 | 0.003894134 | 0.088115202 | 0.876348054 | 0.03164261 | 1.03E-14 | Northern |
| UG_2114 | 0.005489329 | 0.121131956 | 0.848111019 | 0.025267696 | 7.15E-14 | Northern |
| UG_2122 | 0.000539351 | 0.002174171 | 0.392919845 | 0.604366634 | 4.59E-16 | Northern |
| UG_2130 | 0.004166159 | 0.140170365 | 0.841469045 | 0.014194431 | 7.42E-15 | Northern |
| UG_2138 | 0.002821719 | 0.62915419 | 0.367851011 | 0.00017308 | 5.44E-16 | Northern |
| UG_2075 | 0.003868416 | 0.105918034 | 0.867801812 | 0.022411738 | 6.95E-15 | Northern |
| UG_2147 | 0.000541239 | 0.002191315 | 0.394608958 | 0.602658488 | 4.55E-16 | Northern |
| UG_2155 | 0.003335937 | 0.655306698 | 0.341185359 | 0.000172007 | 2.27E-15 | Northern |
| UG_2083 | 0.003827127 | 0.150278483 | 0.834616597 | 0.011277793 | 3.59E-15 | Northern |
| UG_2091 | 0.003949037 | 0.143897318 | 0.839434216 | 0.012719428 | 4.82E-15 | Northern |
| UG_2099 | 0.004757042 | 0.138578182 | 0.839916393 | 0.016748383 | 1.99E-14 | Northern |
| UG_2107 | 0.003094568 | 0.612247687 | 0.384441481 | 0.000216264 | 9.35E-16 | Northern |
| UG_2115 | 0.004085106 | 0.133775957 | 0.846913932 | 0.015225004 | 6.93E-15 | Northern |
| UG_2123 | 0.000530132 | 0.002161444 | 0.394487846 | 0.602820579 | 4.04E-16 | Northern |
| UG_2131 | 0.003592653 | 0.135213464 | 0.848219872 | 0.012974011 | 2.67E-15 | Northern |
| UG_2139 | 0.000556231 | 0.002217368 | 0.392880745 | 0.604345656 | 5.50E-16 | Northern |
| UG_2076 | 0.003152946 | 0.589246703 | 0.407340914 | 0.000259437 | 9.17E-16 | Northern |
| UG_2148 | 9.63E-05 | 0.000111393 | 0.069022928 | 0.930769402 | 1.24E-15 | Northern |
| UG_2156 | 0.007660511 | 0.125127639 | 0.833505675 | 0.033706176 | 7.91E-13 | Northern |
| UG_2084 | 0.003682872 | 0.635120688 | 0.360973979 | 0.000222461 | 3.92E-15 | Northern |
| UG_2092 | 0.004641045 | 0.12936431 | 0.847355975 | 0.01863867 | 1.86E-14 | Northern |

| | | | | | | |
|---|---|---|---|---|---|---|
| UG_2100 | 0.009342807 | 0.188185755 | 0.783803771 | 0.018667667 | 1.79E-12 | Northern |
| UG_2108 | 0.000552716 | 0.002187576 | 0.390043044 | 0.607216664 | 5.57E-16 | Northern |
| UG_2116 | 0.004445562 | 0.141260368 | 0.839298039 | 0.014996031 | 1.18E-14 | Northern |
| UG_2124 | 0.000586501 | 0.002140216 | 0.372289057 | 0.624984226 | 1.08E-15 | Northern |
| UG_2132 | 0.003667234 | 0.101970857 | 0.871639901 | 0.022722008 | 5.03E-15 | Northern |
| UG_2140 | 0.003607639 | 0.094916088 | 0.875984446 | 0.025491827 | 5.10E-15 | Northern |
| UG_2163 | 0.003129311 | 0.587329558 | 0.409280393 | 0.000260738 | 8.57E-16 | Northern |
| UG_2235 | 9.00E-13 | 2.73E-13 | 3.14E-15 | 2.04E-16 | 1 | South_western |
| UG_2243 | 0.038549802 | 0.089629752 | 0.019452157 | 0.000813281 | 0.851555007 | South_western |
| UG_2251 | 3.49E-13 | 1.05E-13 | 9.71E-16 | 5.55E-17 | 1 | South_western |
| UG_2171 | 0.002991643 | 0.044840525 | 0.87559007 | 0.076577761 | 6.51E-15 | Northern |
| UG_2179 | 0.003187207 | 0.58944172 | 0.407108873 | 0.0002622 | 9.92E-16 | Northern |
| UG_2187 | 0.004178462 | 0.1267588 | 0.851746454 | 0.017316284 | 8.93E-15 | Northern |
| UG_2195 | 0.001758877 | 0.014858503 | 0.754781412 | 0.228601208 | 2.71E-15 | Northern |
| UG_2203 | 0.003533491 | 0.606882493 | 0.389323952 | 0.000260064 | 2.35E-15 | Northern |
| UG_2211 | 0.004338344 | 0.150965145 | 0.831890433 | 0.012806078 | 8.90E-15 | Northern |
| UG_2219 | 0.001194046 | 0.007931509 | 0.646243094 | 0.344631351 | 1.19E-15 | Northern |
| UG_2227 | 6.96E-13 | 1.96E-13 | 2.37E-15 | 1.72E-16 | 1 | South_western |
| UG_2164 | 0.002904494 | 0.580352461 | 0.416491063 | 0.000251981 | 4.78E-16 | Northern |
| UG_2236 | 4.66E-13 | 1.34E-13 | 1.42E-15 | 9.29E-17 | 1 | South_western |
| UG_2244 | 0.029534162 | 0.069212872 | 0.013766921 | 0.000540129 | 0.886945917 | South_western |
| UG_2252 | 1.37E-13 | 3.45E-14 | 3.31E-16 | 2.34E-17 | 1 | South_western |
| UG_2172 | 0.004862533 | 0.169768091 | 0.813930782 | 0.011438594 | 1.73E-14 | Northern |
| UG_2180 | 0.004663674 | 0.17033311 | 0.814145608 | 0.010857608 | 1.27E-14 | Northern |
| UG_2188 | 0.003447875 | 0.102641349 | 0.872897157 | 0.021013619 | 3.16E-15 | Northern |
| UG_2196 | 0.003978768 | 0.139016007 | 0.843279521 | 0.013725704 | 5.37E-15 | Northern |
| UG_2204 | 0.003444818 | 0.621131939 | 0.375194695 | 0.000228548 | 2.17E-15 | Northern |
| UG_2212 | 0.002808824 | 0.055770965 | 0.891251371 | 0.05016884 | 2.42E-15 | Northern |
| UG_2220 | 0.00315709 | 0.110387362 | 0.869786 | 0.016669549 | 1.46E-15 | Northern |

| | | | | | | |
|---|---|---|---|---|---|---|
| UG_2228 | 5.92E-13 | 1.68E-13 | 1.93E-15 | 1.35E-16 | 1 | South_western |
| UG_2165 | 0.003404975 | 0.583806635 | 0.412495064 | 0.000293327 | 1.55E-15 | Northern |
| UG_2237 | 3.27E-13 | 9.32E-14 | 9.21E-16 | 5.81E-17 | 1 | South_western |
| UG_2245 | 1.89E-13 | 5.05E-14 | 4.81E-16 | 3.19E-17 | 1 | South_western |
| UG_2253 | 1.06E-13 | 2.76E-14 | 2.36E-16 | 1.49E-17 | 1 | South_western |
| UG_2173 | 0.005397285 | 0.140140911 | 0.835698792 | 0.018763011 | 4.93E-14 | Northern |
| UG_2181 | 0.003462861 | 0.582093808 | 0.414140923 | 0.000302408 | 1.73E-15 | Northern |
| UG_2189 | 0.003893861 | 0.13301512 | 0.848473534 | 0.014617485 | 4.93E-15 | Northern |
| UG_2197 | 0.003088972 | 0.612896259 | 0.383799926 | 0.000214843 | 9.28E-16 | Northern |
| UG_2205 | 0.00303962 | 0.593154886 | 0.403563039 | 0.000242454 | 7.21E-16 | Northern |
| UG_2213 | 0.003469427 | 0.613670055 | 0.3826176 | 0.000242918 | 2.16E-15 | Northern |
| UG_2221 | 0.006174503 | 0.440141131 | 0.55218021 | 0.001504156 | 6.24E-14 | Northern |
| UG_2229 | 7.90E-13 | 2.30E-13 | 2.73E-15 | 1.89E-16 | 1 | South_western |
| UG_2166 | 0.008900832 | 0.183461212 | 0.788954755 | 0.018683202 | 1.30E-12 | Northern |
| UG_2238 | 1.73E-13 | 4.58E-14 | 4.34E-16 | 2.91E-17 | 1 | South_western |
| UG_2246 | 5.69E-13 | 1.68E-13 | 1.80E-15 | 1.15E-16 | 1 | South_western |
| UG_2254 | 5.08E-13 | 1.31E-13 | 1.68E-15 | 1.39E-16 | 1 | South_western |
| UG_2174 | 0.001423004 | 0.015152185 | 0.794138125 | 0.189286685 | 4.58E-16 | Northern |
| UG_2182 | 0.003270169 | 0.592147792 | 0.404317318 | 0.000264721 | 1.22E-15 | Northern |
| UG_2190 | 0.003075441 | 0.617695788 | 0.379022133 | 0.000206638 | 9.30E-16 | Northern |
| UG_2198 | 0.003083745 | 0.581480059 | 0.41516904 | 0.000267157 | 7.43E-16 | Northern |
| UG_2206 | 0.005246874 | 0.179840877 | 0.803875662 | 0.011036586 | 2.78E-14 | Northern |
| UG_2214 | 0.003579367 | 0.576625033 | 0.41946985 | 0.00032575 | 2.12E-15 | Northern |
| UG_2222 | 1.01E-12 | 3.16E-13 | 3.56E-15 | 2.19E-16 | 1 | South_western |
| UG_2230 | 5.88E-13 | 1.74E-13 | 1.87E-15 | 1.20E-16 | 1 | South_western |
| UG_2167 | 0.002895687 | 0.623198495 | 0.373719936 | 0.000185882 | 6.27E-16 | Northern |
| UG_2239 | 6.42E-13 | 1.89E-13 | 2.10E-15 | 1.37E-16 | 1 | South_western |
| UG_2247 | 6.52E-13 | 1.86E-13 | 2.17E-15 | 1.52E-16 | 1 | South_western |
| UG_2255 | 0.0018048 | 0.008068714 | 0.564245166 | 0.42588132 | 3.91E-14 | South_western |

| | | | | | | |
|---|---|---|---|---|---|---|
| UG_2175 | 0.00366766 | 0.106045368 | 0.869163763 | 0.021123208 | 4.69E-15 | Northern |
| UG_2183 | 0.003401379 | 0.096872238 | 0.876667165 | 0.023059217 | 3.19E-15 | Northern |
| UG_2191 | 0.00315962 | 0.599204875 | 0.39739292 | 0.000242585 | 9.94E-16 | Northern |
| UG_2199 | 0.003280672 | 0.613189123 | 0.383301057 | 0.000229148 | 1.44E-15 | Northern |
| UG_2207 | 0.003233897 | 0.624842063 | 0.371716475 | 0.000207564 | 1.41E-15 | Northern |
| UG_2215 | 0.004019498 | 0.128914591 | 0.85099022 | 0.016075691 | 6.54E-15 | Northern |
| UG_2223 | 0.031724119 | 0.073012144 | 0.015222776 | 0.000627751 | 0.87941321 | South_western |
| UG_2231 | 2.01E-13 | 5.35E-14 | 5.18E-16 | 3.47E-17 | 1 | South_western |
| UG_2168 | 0.053496153 | 0.664186168 | 0.279599679 | 0.002715595 | 2.41E-06 | Northern |
| UG_2240 | 5.38E-13 | 1.56E-13 | 1.69E-15 | 1.11E-16 | 1 | South_western |
| UG_2248 | 2.13E-13 | 6.12E-14 | 5.38E-16 | 3.13E-17 | 1 | South_western |
| UG_2256 | 8.09E-13 | 2.33E-13 | 2.82E-15 | 2.00E-16 | 1 | South_western |
| UG_2176 | 0.001863282 | 0.019400874 | 0.80784408 | 0.170891763 | 1.76E-15 | Northern |
| UG_2184 | 0.00950633 | 0.192162432 | 0.780120477 | 0.01821076 | 1.98E-12 | Northern |
| UG_2192 | 0.003686451 | 0.126400316 | 0.854702956 | 0.015210277 | 3.59E-15 | Northern |
| UG_2200 | 0.003942254 | 0.554389854 | 0.44124575 | 0.000422142 | 3.77E-15 | Northern |
| UG_2208 | 0.004089558 | 0.117297642 | 0.858986112 | 0.019626688 | 8.71E-15 | Northern |
| UG_2216 | 0.089376954 | 0.664942793 | 0.241705925 | 0.003798515 | 0.000175812 | Northern |
| UG_2224 | 4.30E-13 | 1.21E-13 | 1.30E-15 | 8.73E-17 | 1 | South_western |
| UG_2232 | 9.97E-13 | 3.07E-13 | 3.54E-15 | 2.26E-16 | 1 | South_western |
| UG_2169 | 0.089048588 | 0.659108658 | 0.247687352 | 0.003997539 | 0.000157862 | Northern |
| UG_2241 | 8.92E-13 | 2.72E-13 | 3.10E-15 | 1.98E-16 | 1 | South_western |
| UG_2249 | 1.33E-12 | 4.53E-13 | 4.81E-15 | 2.60E-16 | 1 | South_western |
| UG_2177 | 0.004011784 | 0.1373831 | 0.844431377 | 0.014173739 | 5.82E-15 | Northern |
| UG_2185 | 0.004395681 | 0.143581893 | 0.837675735 | 0.014346692 | 1.06E-14 | Northern |
| UG_2193 | 0.003571927 | 0.600109273 | 0.396042772 | 0.000276028 | 2.43E-15 | Northern |
| UG_2201 | 0.004013982 | 0.138136755 | 0.843818355 | 0.014030908 | 5.79E-15 | Northern |
| UG_2209 | 0.005317702 | 0.158958225 | 0.82132924 | 0.014394833 | 3.64E-14 | Northern |
| UG_2217 | 0.003676146 | 0.114587807 | 0.863437444 | 0.018298604 | 4.16E-15 | Northern |

| | | | | | | |
|---|---|---|---|---|---|---|
| UG_2225 | 1.27E-12 | 3.76E-13 | 4.85E-15 | 3.44E-16 | 1 | South_western |
| UG_2233 | 0.004025386 | 0.12405613 | 0.854580056 | 0.017338429 | 7.05E-15 | South_western |
| UG_2170 | 0.003405158 | 0.618550019 | 0.377814982 | 0.000229841 | 1.96E-15 | Northern |
| UG_2242 | 8.09E-13 | 2.49E-13 | 2.73E-15 | 1.68E-16 | 1 | South_western |
| UG_2250 | 4.07E-13 | 1.15E-13 | 1.22E-15 | 8.11E-17 | 1 | South_western |
| UG_2178 | 0.003858344 | 0.616905642 | 0.378969108 | 0.000266906 | 4.79E-15 | Northern |
| UG_2186 | 0.004643489 | 0.132276593 | 0.845215511 | 0.017864406 | 1.80E-14 | Northern |
| UG_2194 | 0.003895716 | 0.093896373 | 0.874001307 | 0.028206604 | 9.17E-15 | Northern |
| UG_2202 | 0.000578867 | 0.002332705 | 0.400458247 | 0.596630181 | 6.09E-16 | Northern |
| UG_2210 | 0.00387528 | 0.128633212 | 0.851972071 | 0.015519437 | 5.02E-15 | Northern |
| UG_2218 | 0.003821659 | 0.12813575 | 0.852639941 | 0.01540265 | 4.57E-15 | Northern |
| UG_2226 | 2.26E-13 | 6.08E-14 | 6.01E-16 | 4.05E-17 | 1 | South_western |
| UG_2234 | 0.006210903 | 0.012656482 | 0.002074617 | 8.43E-05 | 0.9789737 | South_western |
| UG_2257 | 1.46E-12 | 4.40E-13 | 5.77E-15 | 4.08E-16 | 1 | South_western |
| UG_2329 | 8.77E-13 | 2.66E-13 | 3.04E-15 | 1.95E-16 | 1 | South_western |
| UG_2337 | 0.003880853 | 0.09543983 | 0.873411991 | 0.027267326 | 8.64E-15 | Northern |
| UG_2345 | 0.004037609 | 0.13685036 | 0.844730561 | 0.01438147 | 6.13E-15 | Northern |
| UG_2265 | 5.32E-13 | 1.61E-13 | 1.64E-15 | 9.84E-17 | 1 | South_western |
| UG_2273 | 3.22E-13 | 9.13E-14 | 9.03E-16 | 5.72E-17 | 1 | South_western |
| UG_2281 | 8.16E-13 | 2.52E-13 | 2.75E-15 | 1.68E-16 | 1 | South_western |
| UG_2289 | 8.39E-13 | 2.46E-13 | 2.93E-15 | 2.01E-16 | 1 | South_western |
| UG_2297 | 2.45E-13 | 6.82E-14 | 6.51E-16 | 4.13E-17 | 1 | South_western |
| UG_2305 | 2.49E-13 | 7.16E-14 | 6.52E-16 | 3.86E-17 | 1 | South_western |
| UG_2313 | 2.59E-13 | 7.08E-14 | 7.05E-16 | 4.68E-17 | 1 | South_western |
| UG_2321 | 9.89E-13 | 3.13E-13 | 3.45E-15 | 2.08E-16 | 1 | South_western |
| UG_2258 | 1.06E-13 | 2.67E-14 | 2.43E-16 | 1.67E-17 | 1 | South_western |
| UG_2330 | 3.68E-13 | 1.02E-13 | 1.08E-15 | 7.44E-17 | 1 | South_western |
| UG_2338 | 0.00066528 | 0.003640364 | 0.514304879 | 0.481389477 | 2.32E-16 | Northern |
| UG_2346 | 0.002533893 | 0.024515172 | 0.811556928 | 0.161394007 | 9.66E-15 | Northern |

| | | | | | | |
|---|---|---|---|---|---|---|
| UG_2266 | 6.40E-12 | 2.27E-12 | 3.33E-14 | 2.11E-15 | 1 | South_western |
| UG_2274 | 7.65E-13 | 2.22E-13 | 2.62E-15 | 1.80E-16 | 1 | South_western |
| UG_2282 | 5.76E-13 | 1.73E-13 | 1.81E-15 | 1.11E-16 | 1 | South_western |
| UG_2290 | 0.02987138 | 0.370526193 | 0.586506615 | 0.013095805 | 6.04E-09 | South_western |
| UG_2298 | 1.87E-13 | 5.12E-14 | 4.66E-16 | 2.91E-17 | 1 | South_western |
| UG_2306 | 5.05E-10 | 2.42E-10 | 6.54E-12 | 4.34E-13 | 0.999999999 | South_western |
| UG_2314 | 2.20E-13 | 5.74E-14 | 5.88E-16 | 4.18E-17 | 1 | South_western |
| UG_2322 | 6.52E-12 | 2.31E-12 | 3.41E-14 | 2.18E-15 | 1 | South_western |
| UG_2259 | 0.004031719 | 0.127723091 | 0.851824735 | 0.016420455 | 6.79E-15 | South_western |
| UG_2331 | 2.05E-09 | 1.07E-09 | 3.57E-11 | 2.45E-12 | 0.999999997 | South_western |
| UG_2339 | 0.003690716 | 0.087810901 | 0.87840523 | 0.030093152 | 7.01E-15 | Northern |
| UG_2347 | 0.00252521 | 0.02672455 | 0.827913139 | 0.142837101 | 7.24E-15 | Northern |
| UG_2267 | 0.000524202 | 0.001962097 | 0.367996646 | 0.629517054 | 6.08E-16 | South_western |
| UG_2275 | 0.005191083 | 0.115550495 | 0.85322964 | 0.026028782 | 5.16E-14 | South_western |
| UG_2283 | 1.52E-12 | 5.20E-13 | 5.70E-15 | 3.14E-16 | 1 | South_western |
| UG_2291 | 2.68E-10 | 1.38E-10 | 2.86E-12 | 1.49E-13 | 1 | South_western |
| UG_2299 | 1.16E-12 | 3.25E-13 | 4.52E-15 | 3.61E-16 | 1 | South_western |
| UG_2307 | 1.90E-13 | 5.32E-14 | 4.74E-16 | 2.87E-17 | 1 | South_western |
| UG_2315 | 4.09E-11 | 1.65E-11 | 3.12E-13 | 1.99E-14 | 1 | South_western |
| UG_2323 | 2.84E-13 | 7.74E-14 | 7.88E-16 | 5.31E-17 | 1 | South_western |
| UG_2260 | 5.99E-13 | 1.75E-13 | 1.93E-15 | 1.27E-16 | 1 | South_western |
| UG_2332 | 0.003542045 | 0.072897427 | 0.883405705 | 0.040154822 | 7.56E-15 | Northern |
| UG_2340 | 0.003822931 | 0.09199665 | 0.87548013 | 0.028700288 | 8.30E-15 | Northern |
| UG_2348 | 0.002548672 | 0.051299019 | 0.893834701 | 0.052317608 | 1.42E-15 | Northern |
| UG_2268 | 1.81E-12 | 6.31E-13 | 6.98E-15 | 3.78E-16 | 1 | South_western |
| UG_2276 | 3.57E-13 | 9.79E-14 | 1.05E-15 | 7.24E-17 | 1 | South_western |
| UG_2284 | 3.78E-13 | 1.04E-13 | 1.12E-15 | 7.80E-17 | 1 | South_western |
| UG_2292 | 1.14E-12 | 3.50E-13 | 4.21E-15 | 2.77E-16 | 1 | South_western |
| UG_2300 | 1.84E-12 | 6.37E-13 | 7.11E-15 | 3.89E-16 | 1 | South_western |

| | | | | | | |
|---|---|---|---|---|---|---|
| UG_2308 | 1.67E-11 | 6.71E-12 | 1.03E-13 | 5.88E-15 | 1 | South_western |
| UG_2316 | 2.58E-13 | 6.74E-14 | 7.16E-16 | 5.21E-17 | 1 | South_western |
| UG_2324 | 1.69E-13 | 4.41E-14 | 4.24E-16 | 2.90E-17 | 1 | South_western |
| UG_2261 | 1.20E-13 | 3.03E-14 | 2.82E-16 | 1.97E-17 | 1 | South_western |
| UG_2333 | 0.003722903 | 0.123676045 | 0.856568691 | 0.016032361 | 4.00E-15 | Northern |
| UG_2341 | 0.002493878 | 0.04699793 | 0.89128236 | 0.059225831 | 1.48E-15 | Northern |
| UG_2349 | 0.003493581 | 0.072570068 | 0.884037565 | 0.039898785 | 6.90E-15 | Northern |
| UG_2269 | 5.32E-11 | 2.33E-11 | 4.17E-13 | 2.36E-14 | 1 | South_western |
| UG_2277 | 1.44E-11 | 5.74E-12 | 8.62E-14 | 4.86E-15 | 1 | South_western |
| UG_2285 | 2.89E-12 | 1.01E-12 | 1.25E-14 | 7.16E-16 | 1 | South_western |
| UG_2293 | 3.49E-13 | 9.31E-14 | 1.03E-15 | 7.57E-17 | 1 | South_western |
| UG_2301 | 1.98E-12 | 6.37E-13 | 8.14E-15 | 5.27E-16 | 1 | South_western |
| UG_2309 | 3.35E-13 | 9.14E-14 | 9.72E-16 | 6.75E-17 | 1 | South_western |
| UG_2317 | 5.38E-13 | 1.61E-13 | 1.67E-15 | 1.03E-16 | 1 | South_western |
| UG_2325 | 3.20E-12 | 1.18E-12 | 1.38E-14 | 7.24E-16 | 1 | South_western |
| UG_2262 | 1.20E-12 | 4.00E-13 | 4.28E-15 | 2.39E-16 | 1 | South_western |
| UG_2334 | 0.003716545 | 0.121038362 | 0.858565997 | 0.016679096 | 4.10E-15 | Northern |
| UG_2342 | 0.000536128 | 0.002452783 | 0.431227875 | 0.565783214 | 2.33E-16 | Northern |
| UG_2350 | 0.003905397 | 0.102734025 | 0.869396438 | 0.023964141 | 7.88E-15 | Northern |
| UG_2270 | 3.25E-13 | 9.20E-14 | 9.16E-16 | 5.85E-17 | 1 | South_western |
| UG_2278 | 7.81E-13 | 2.36E-13 | 2.64E-15 | 1.69E-16 | 1 | South_western |
| UG_2286 | 1.25E-12 | 3.91E-13 | 4.65E-15 | 2.97E-16 | 1 | South_western |
| UG_2294 | 3.76E-13 | 1.09E-13 | 1.08E-15 | 6.69E-17 | 1 | South_western |
| UG_2302 | 5.65E-13 | 1.65E-13 | 1.79E-15 | 1.17E-16 | 1 | South_western |
| UG_2310 | 1.54E-12 | 5.30E-13 | 5.71E-15 | 3.07E-16 | 1 | South_western |
| UG_2318 | 3.54E-12 | 1.18E-12 | 1.64E-14 | 1.07E-15 | 1 | South_western |
| UG_2326 | 1.44E-12 | 5.08E-13 | 5.19E-15 | 2.62E-16 | 1 | South_western |
| UG_2263 | 9.79E-13 | 3.27E-13 | 3.31E-15 | 1.77E-16 | 1 | South_western |
| UG_2335 | 0.004278394 | 0.088197124 | 0.872662916 | 0.034861565 | 2.07E-14 | Northern |

| | | | | | | |
|---|---|---|---|---|---|---|
| UG_2343 | 0.00409741 | 0.107203956 | 0.865400408 | 0.023298225 | 1.04E-14 | Northern |
| UG_2271 | 8.52E-13 | 2.60E-13 | 2.93E-15 | 1.85E-16 | 1 | South_western |
| UG_2279 | 2.69E-12 | 8.94E-13 | 1.17E-14 | 7.41E-16 | 1 | South_western |
| UG_2287 | 4.89E-13 | 1.45E-13 | 1.49E-15 | 9.13E-17 | 1 | South_western |
| UG_2295 | 2.92E-13 | 8.05E-14 | 8.11E-16 | 5.37E-17 | 1 | South_western |
| UG_2303 | 1.49E-12 | 4.88E-13 | 5.66E-15 | 3.37E-16 | 1 | South_western |
| UG_2311 | 2.35E-13 | 6.19E-14 | 6.34E-16 | 4.47E-17 | 1 | South_western |
| UG_2319 | 2.75E-12 | 9.77E-13 | 1.16E-14 | 6.47E-16 | 1 | South_western |
| UG_2327 | 1.42E-12 | 4.50E-13 | 5.40E-15 | 3.41E-16 | 1 | South_western |
| UG_2264 | 6.38E-13 | 2.06E-13 | 1.98E-15 | 1.06E-16 | 1 | South_western |
| UG_2336 | 0.003120805 | 0.073579798 | 0.888680689 | 0.034618708 | 2.91E-15 | Northern |
| UG_2344 | 0.004624676 | 0.141127607 | 0.83856999 | 0.015677727 | 1.57E-14 | Northern |
| UG_2272 | 2.78E-13 | 7.53E-14 | 7.73E-16 | 5.29E-17 | 1 | South_western |
| UG_2280 | 1.61E-10 | 7.88E-11 | 1.55E-12 | 8.23E-14 | 1 | South_western |
| UG_2288 | 1.99E-12 | 7.03E-13 | 7.79E-15 | 4.14E-16 | 1 | South_western |
| UG_2296 | 2.99E-13 | 7.86E-14 | 8.56E-16 | 6.27E-17 | 1 | South_western |
| UG_2304 | 1.90E-13 | 5.01E-14 | 4.88E-16 | 3.33E-17 | 1 | South_western |
| UG_2312 | 4.19E-13 | 1.17E-13 | 1.27E-15 | 8.65E-17 | 1 | South_western |
| UG_2320 | 2.00E-13 | 5.46E-14 | 5.11E-16 | 3.27E-17 | 1 | South_western |
| UG_2328 | 4.28E-13 | 1.27E-13 | 1.26E-15 | 7.57E-17 | 1 | South_western |
| UG_2351 | 0.005446274 | 0.162341353 | 0.818053887 | 0.014158487 | 4.21E-14 | Northern |
| UG_2423 | 0.00068322 | 0.002469664 | 0.385577657 | 0.611269458 | 2.08E-15 | North_western |
| UG_2431 | 0.006934579 | 0.082542344 | 0.846551809 | 0.063971268 | 8.76E-13 | North_western |
| UG_2439 | 0.006871811 | 0.081335028 | 0.846788698 | 0.065004463 | 8.46E-13 | North_western |
| UG_2359 | 0.003836288 | 0.094997311 | 0.874000178 | 0.027166223 | 8.01E-15 | Northern |
| UG_2367 | 0.000276564 | 0.000564579 | 0.176771833 | 0.822387024 | 1.70E-15 | Northern |
| UG_2375 | 0.003020467 | 0.130700309 | 0.854799145 | 0.011480078 | 7.95E-16 | Northern |
| UG_2383 | 0.00074871 | 0.004063173 | 0.525512264 | 0.469675852 | 3.95E-16 | Northern |
| UG_2391 | 0.000947276 | 0.005308527 | 0.563976321 | 0.429767877 | 9.14E-16 | Northern |

| | | | | | | |
|---|---|---|---|---|---|---|
| UG_2399 | 0.000344491 | 0.00069905 | 0.189335759 | 0.8096207 | 3.98E-15 | Northern |
| UG_2407 | 0.005117393 | 0.128428649 | 0.845471407 | 0.020982551 | 3.86E-14 | Northern |
| UG_2415 | 0.000184068 | 0.000345259 | 0.141590338 | 0.857880335 | 6.36E-16 | North_western |
| UG_2352 | 0.004503059 | 0.161264958 | 0.822551264 | 0.011680719 | 1.06E-14 | Northern |
| UG_2424 | 0.000638065 | 0.002323964 | 0.38057868 | 0.616459291 | 1.53E-15 | North_western |
| UG_2432 | 0.0067502 | 0.082337478 | 0.848355846 | 0.062556476 | 7.19E-13 | North_western |
| UG_2440 | 0.000114517 | 0.000164721 | 0.091814384 | 0.907906378 | 5.87E-16 | North_western |
| UG_2360 | 0.004265003 | 0.123021002 | 0.853965124 | 0.018748871 | 1.09E-14 | Northern |
| UG_2368 | 0.001371942 | 0.010640393 | 0.708275083 | 0.279712583 | 1.19E-15 | Northern |
| UG_2376 | 0.002794906 | 0.602883267 | 0.39411542 | 0.000206408 | 4.19E-16 | Northern |
| UG_2384 | 0.000115737 | 0.000140094 | 0.077351534 | 0.922392635 | 1.80E-15 | Northern |
| UG_2392 | 0.00042264 | 0.000973451 | 0.227742276 | 0.770861633 | 4.05E-15 | Northern |
| UG_2400 | 0.006420459 | 0.140474048 | 0.830634976 | 0.022470516 | 1.76E-13 | Northern |
| UG_2408 | 0.002935841 | 0.09304926 | 0.882794459 | 0.02122044 | 1.17E-15 | Northern |
| UG_2416 | 0.000170341 | 0.000314672 | 0.135756264 | 0.863758722 | 5.26E-16 | North_western |
| UG_2353 | 0.002545815 | 0.042349232 | 0.883437451 | 0.071667502 | 2.22E-15 | Northern |
| UG_2425 | 0.00011733 | 0.000174072 | 0.095564122 | 0.904144476 | 5.28E-16 | North_western |
| UG_2433 | 0.000132754 | 0.000194721 | 0.098786415 | 0.90088611 | 8.86E-16 | North_western |
| UG_2441 | 0.000100148 | 0.000131018 | 0.079409376 | 0.920359458 | 6.57E-16 | North_western |
| UG_2361 | 0.000825592 | 0.004900594 | 0.566941958 | 0.427331856 | 3.94E-16 | Northern |
| UG_2369 | 0.00177619 | 0.008688148 | 0.592018613 | 0.397517049 | 2.45E-14 | Northern |
| UG_2377 | 0.000102955 | 0.000123865 | 0.073638969 | 0.92613421 | 1.23E-15 | Northern |
| UG_2385 | 0.004400973 | 0.118109183 | 0.856535231 | 0.020954613 | 1.47E-14 | Northern |
| UG_2393 | 0.002529626 | 0.011474946 | 0.607345014 | 0.378650413 | 1.56E-13 | Northern |
| UG_2401 | 0.002953761 | 0.070719263 | 0.891274934 | 0.035052041 | 2.10E-15 | Northern |
| UG_2409 | 0.000335173 | 0.000750206 | 0.205559029 | 0.793355592 | 2.01E-15 | Northern |
| UG_2417 | 0.000173831 | 0.000296153 | 0.126281308 | 0.873248709 | 9.31E-16 | North_western |
| UG_2354 | 0.004837238 | 0.12716893 | 0.847858648 | 0.020135185 | 2.60E-14 | Northern |
| UG_2426 | 8.17E-05 | 0.000105087 | 0.0724382 | 0.927375033 | 3.51E-16 | North_western |

| | | | | | | |
|---|---|---|---|---|---|---|
| UG_2434 | 0.002259313 | 0.008914836 | 0.548918842 | 0.439907009 | 1.79E-13 | North_western |
| UG_2442 | 0.00011146 | 0.000178758 | 0.101453799 | 0.898255983 | 2.70E-16 | North_western |
| UG_2362 | 0.002262466 | 0.037101334 | 0.88175138 | 0.07888482 | 1.26E-15 | Northern |
| UG_2370 | 0.000506165 | 0.001932923 | 0.370179881 | 0.627381032 | 4.76E-16 | Northern |
| UG_2378 | 0.086456382 | 0.66905226 | 0.240755088 | 0.003597473 | 0.000138796 | Northern |
| UG_2386 | 0.003071071 | 0.074691395 | 0.889090359 | 0.033147175 | 2.50E-15 | Northern |
| UG_2394 | 0.001652227 | 0.01446812 | 0.75870027 | 0.225179383 | 1.79E-15 | Northern |
| UG_2402 | 0.000931941 | 0.006597835 | 0.638899216 | 0.353571008 | 3.08E-16 | Northern |
| UG_2410 | 0.003173624 | 0.096063075 | 0.879016198 | 0.021747103 | 1.95E-15 | Northern |
| UG_2418 | 0.000158706 | 0.000231242 | 0.104649969 | 0.894960083 | 1.76E-15 | North_western |
| UG_2355 | 0.003103556 | 0.067494574 | 0.889347156 | 0.040054714 | 3.34E-15 | Northern |
| UG_2427 | 0.000113131 | 0.000146292 | 0.082052238 | 0.91768834 | 1.09E-15 | North_western |
| UG_2435 | 0.001888772 | 0.005073493 | 0.405133255 | 0.58790448 | 5.62E-13 | North_western |
| UG_2443 | 0.004590185 | 0.041553854 | 0.825507222 | 0.128348739 | 2.13E-13 | North_western |
| UG_2363 | 0.002665594 | 0.048329807 | 0.888539611 | 0.060464987 | 2.28E-15 | Northern |
| UG_2371 | 0.001026811 | 0.007198714 | 0.646641453 | 0.345133021 | 4.90E-16 | Northern |
| UG_2379 | 0.050636517 | 0.343766202 | 0.579187034 | 0.026409912 | 3.34E-07 | Northern |
| UG_2387 | 0.000287957 | 0.000554132 | 0.169439337 | 0.829718574 | 2.82E-15 | Northern |
| UG_2395 | 0.000505964 | 0.001921032 | 0.3684597 | 0.629113304 | 4.90E-16 | Northern |
| UG_2403 | 0.001134129 | 0.008748645 | 0.687189313 | 0.302927913 | 5.17E-16 | Northern |
| UG_2411 | 0.00032192 | 0.000623906 | 0.177198088 | 0.821856085 | 4.07E-15 | Northern |
| UG_2419 | 0.000185929 | 0.000313631 | 0.128061742 | 0.871438699 | 1.26E-15 | North_western |
| UG_2356 | 0.003295672 | 0.111358305 | 0.868172693 | 0.017173331 | 1.97E-15 | Northern |
| UG_2428 | 0.010722474 | 0.074068175 | 0.800293148 | 0.114916203 | 3.21E-11 | North_western |
| UG_2436 | 0.000100359 | 0.000130801 | 0.079166188 | 0.920602652 | 6.78E-16 | North_western |
| UG_2444 | 0.000103051 | 0.000147742 | 0.08806692 | 0.911682287 | 4.10E-16 | North_western |
| UG_2364 | 0.003847187 | 0.132963653 | 0.848750291 | 0.01443887 | 4.51E-15 | Northern |
| UG_2372 | 0.001695569 | 0.016232718 | 0.782654534 | 0.199417179 | 1.49E-15 | Northern |
| UG_2380 | 0.003755049 | 0.118623873 | 0.860093113 | 0.017527964 | 4.57E-15 | Northern |

| | | | | | | |
|---|---|---|---|---|---|---|
| UG_2388 | 0.000944315 | 0.005538527 | 0.578749926 | 0.414767233 | 7.35E-16 | Northern |
| UG_2396 | 0.000443227 | 0.001083155 | 0.243951907 | 0.754521711 | 3.43E-15 | Northern |
| UG_2404 | 0.003800799 | 0.070214365 | 0.879889225 | 0.046095611 | 1.38E-14 | Northern |
| UG_2412 | 0.003271178 | 0.101891386 | 0.874702681 | 0.020134755 | 2.18E-15 | Northern |
| UG_2420 | 7.88E-05 | 9.90E-05 | 0.069731146 | 0.930091097 | 3.60E-16 | North_western |
| UG_2357 | 0.004678059 | 0.132441334 | 0.844916705 | 0.017963901 | 1.90E-14 | Northern |
| UG_2429 | 0.00696153 | 0.085640573 | 0.847092034 | 0.060305863 | 8.31E-13 | North_western |
| UG_2437 | 0.003513571 | 0.606787021 | 0.389440788 | 0.00025862 | 2.26E-15 | North_western |
| UG_2365 | 0.004521354 | 0.148230781 | 0.833360299 | 0.013887567 | 1.24E-14 | Northern |
| UG_2373 | 0.000113613 | 0.000142248 | 0.079522077 | 0.920222062 | 1.36E-15 | Northern |
| UG_2381 | 0.001812933 | 0.017361413 | 0.787701155 | 0.193124499 | 2.03E-15 | Northern |
| UG_2389 | 0.002689504 | 0.044507633 | 0.88298959 | 0.069813272 | 2.97E-15 | Northern |
| UG_2397 | 0.00375124 | 0.123672765 | 0.85641104 | 0.016164956 | 4.23E-15 | Northern |
| UG_2405 | 0.001670087 | 0.009315581 | 0.62770973 | 0.361304602 | 1.07E-14 | Northern |
| UG_2413 | 0.001698187 | 0.011784803 | 0.697037579 | 0.289479431 | 4.76E-15 | Northern |
| UG_2421 | 0.000295025 | 0.000467575 | 0.141705097 | 0.857532303 | 9.99E-15 | North_western |
| UG_2358 | 0.003251773 | 0.077399472 | 0.88631128 | 0.033037474 | 3.55E-15 | Northern |
| UG_2430 | 0.000115179 | 0.000171988 | 0.095546037 | 0.904166796 | 4.75E-16 | North_western |
| UG_2438 | 0.005752754 | 0.020980124 | 0.625766526 | 0.347500596 | 1.43E-11 | North_western |
| UG_2366 | 0.001289615 | 0.007722843 | 0.621574312 | 0.36941323 | 2.56E-15 | Northern |
| UG_2374 | 0.004278357 | 0.091384086 | 0.871635692 | 0.032701865 | 1.93E-14 | Northern |
| UG_2382 | 0.002787496 | 0.136766916 | 0.85082525 | 0.009620338 | 4.13E-16 | Northern |
| UG_2390 | 0.00134677 | 0.007612337 | 0.607703361 | 0.383337532 | 3.96E-15 | Northern |
| UG_2398 | 0.003269518 | 0.092763597 | 0.880022272 | 0.023944613 | 2.59E-15 | Northern |
| UG_2406 | 0.000110366 | 0.000131926 | 0.075046017 | 0.924711692 | 1.64E-15 | Northern |
| UG_2414 | 0.017306269 | 0.104418421 | 0.775808435 | 0.102466875 | 5.54E-10 | North_western |
| UG_2422 | 0.000621583 | 0.002301195 | 0.382643118 | 0.614434104 | 1.26E-15 | North_western |
| UG_2445 | 0.00294052 | 0.009046496 | 0.497415982 | 0.490597001 | 1.73E-12 | North_western |
| UG_2517 | 0.00019126 | 0.000307443 | 0.123346031 | 0.876155266 | 1.88E-15 | North_western |

| | | | | | | |
|---|---|---|---|---|---|---|
| UG_2525 | 0.004730804 | 0.042194332 | 0.824160228 | 0.128914635 | 2.58E-13 | North_western |
| UG_2533 | 0.000184961 | 0.000286137 | 0.117348048 | 0.882180853 | 2.11E-15 | North_western |
| UG_2453 | 0.000101418 | 0.00013397 | 0.080567106 | 0.919197505 | 6.47E-16 | North_western |
| UG_2461 | 7.69E-05 | 9.48E-05 | 0.067770546 | 0.932057755 | 3.73E-16 | North_western |
| UG_2469 | 0.000145016 | 0.00021856 | 0.104797896 | 0.894838528 | 1.03E-15 | North_western |
| UG_2477 | 0.006859545 | 0.082610452 | 0.847329799 | 0.063200204 | 8.06E-13 | North_western |
| UG_2485 | 0.000118549 | 0.000162216 | 0.08839524 | 0.911323995 | 9.09E-16 | North_western |
| UG_2493 | 0.001068677 | 0.003823213 | 0.431211349 | 0.563896761 | 1.31E-14 | North_western |
| UG_2501 | 0.000138091 | 0.000215127 | 0.106439285 | 0.893207497 | 7.02E-16 | North_western |
| UG_2509 | 0.000182574 | 0.000377024 | 0.154985675 | 0.844454727 | 3.44E-16 | North_western |
| UG_2446 | 9.50E-05 | 0.000127112 | 0.079680655 | 0.920097201 | 4.73E-16 | North_western |
| UG_2518 | 0.007965365 | 0.09622791 | 0.839443866 | 0.056362859 | 1.78E-12 | North_western |
| UG_2526 | 0.018358608 | 0.106750112 | 0.770694983 | 0.104196297 | 8.31E-10 | North_western |
| UG_2534 | 9.21E-05 | 0.000122468 | 0.078277409 | 0.921507982 | 4.41E-16 | North_western |
| UG_2454 | 0.000116321 | 0.000181461 | 0.100203084 | 0.899499134 | 3.74E-16 | North_western |
| UG_2462 | 9.55E-05 | 0.000122027 | 0.076173267 | 0.923609187 | 6.44E-16 | North_western |
| UG_2470 | 0.001357004 | 0.00330697 | 0.342574344 | 0.652761683 | 2.55E-13 | North_western |
| UG_2478 | 0.004756454 | 0.042412339 | 0.82426885 | 0.128562357 | 2.65E-13 | North_western |
| UG_2486 | 0.000119077 | 0.000177447 | 0.096505781 | 0.903197695 | 5.42E-16 | North_western |
| UG_2494 | 0.000112871 | 0.000164175 | 0.092368382 | 0.907354572 | 5.20E-16 | North_western |
| UG_2502 | 8.03E-05 | 9.63E-05 | 0.067030173 | 0.932793229 | 5.11E-16 | North_western |
| UG_2510 | 0.00011537 | 0.000165022 | 0.091542187 | 0.908177421 | 6.25E-16 | North_western |
| UG_2447 | 0.000126112 | 0.000188865 | 0.099021011 | 0.900664012 | 6.46E-16 | North_western |
| UG_2519 | 0.000114825 | 0.000175582 | 0.097751106 | 0.901958488 | 4.05E-16 | North_western |
| UG_2527 | 9.50E-05 | 0.00012418 | 0.077818932 | 0.92196188 | 5.47E-16 | North_western |
| UG_2535 | 9.05E-05 | 0.00011978 | 0.077434282 | 0.922355448 | 4.24E-16 | North_western |
| UG_2455 | 0.008562832 | 0.091763527 | 0.833970325 | 0.065703316 | 3.40E-12 | North_western |
| UG_2463 | 0.000106137 | 0.000145785 | 0.085233654 | 0.914514424 | 5.96E-16 | North_western |
| UG_2471 | 0.00013813 | 0.000224148 | 0.110875972 | 0.888761751 | 5.46E-16 | North_western |

| | | | | | | |
|---|---|---|---|---|---|---|
| UG_2479 | 8.91E-05 | 0.000109889 | 0.071609664 | 0.928191321 | 6.28E-16 | North_western |
| UG_2487 | 0.006681982 | 0.081748067 | 0.848876938 | 0.062693013 | 6.77E-13 | North_western |
| UG_2495 | 0.000134915 | 0.00022065 | 0.110805823 | 0.888838613 | 4.77E-16 | North_western |
| UG_2503 | 0.000263838 | 0.000604933 | 0.194081974 | 0.805049255 | 7.15E-16 | North_western |
| UG_2511 | 0.00011852 | 0.000174434 | 0.095141805 | 0.904565242 | 5.76E-16 | North_western |
| UG_2448 | 0.000115443 | 0.000184515 | 0.102393804 | 0.897306238 | 3.13E-16 | North_western |
| UG_2520 | 0.000135243 | 0.000178846 | 0.089581695 | 0.910104216 | 1.81E-15 | North_western |
| UG_2528 | 0.000103983 | 0.000172668 | 0.102460157 | 0.897263192 | 1.69E-16 | North_western |
| UG_2536 | 0.002127076 | 0.012971279 | 0.681352593 | 0.303549052 | 2.16E-14 | North_western |
| UG_2456 | 0.000109391 | 0.000144083 | 0.082582452 | 0.917164073 | 8.65E-16 | North_western |
| UG_2464 | 0.006785897 | 0.081809226 | 0.847830346 | 0.063574531 | 7.59E-13 | North_western |
| UG_2472 | 7.57E-05 | 0.000106789 | 0.07740501 | 0.922412509 | 1.50E-16 | North_western |
| UG_2480 | 0.006784695 | 0.079682075 | 0.847073532 | 0.066459699 | 8.05E-13 | North_western |
| UG_2488 | 7.91E-05 | 9.82E-05 | 0.069022635 | 0.930800032 | 3.92E-16 | North_western |
| UG_2496 | 0.000128944 | 0.000182717 | 0.094412204 | 0.905276134 | 9.89E-16 | North_western |
| UG_2504 | 8.69E-05 | 0.000110286 | 0.07305269 | 0.926750077 | 4.81E-16 | North_western |
| UG_2512 | 0.000138214 | 0.000209452 | 0.103569275 | 0.896083059 | 8.36E-16 | North_western |
| UG_2449 | 0.002820851 | 0.007911533 | 0.462846552 | 0.526421064 | 2.29E-12 | North_western |
| UG_2521 | 0.00010611 | 0.000161576 | 0.094597943 | 0.90513437 | 3.13E-16 | North_western |
| UG_2529 | 8.75E-05 | 0.000113174 | 0.074696532 | 0.925102789 | 4.35E-16 | North_western |
| UG_2537 | 0.000105789 | 0.000140165 | 0.082070853 | 0.917683193 | 7.39E-16 | North_western |
| UG_2457 | 8.02E-05 | 9.83E-05 | 0.068474938 | 0.931346534 | 4.47E-16 | North_western |
| UG_2465 | 0.00017467 | 0.000307646 | 0.13070476 | 0.868812924 | 7.72E-16 | North_western |
| UG_2473 | 0.000320646 | 0.000782909 | 0.219454297 | 0.779442149 | 1.02E-15 | North_western |
| UG_2481 | 9.27E-05 | 0.000150379 | 0.095994514 | 0.903762373 | 1.30E-16 | North_western |
| UG_2489 | 0.000119078 | 0.000176978 | 0.096247857 | 0.903456087 | 5.51E-16 | North_western |
| UG_2497 | 0.000401823 | 0.001322969 | 0.306636801 | 0.691638408 | 4.35E-16 | North_western |
| UG_2505 | 0.000172024 | 0.00040337 | 0.171508533 | 0.827916074 | 1.28E-16 | North_western |
| UG_2513 | 0.000112846 | 0.000167807 | 0.094445178 | 0.905274169 | 4.52E-16 | North_western |

| | | | | | | |
|---|---|---|---|---|---|---|
| UG_2450 | 0.026412378 | 0.14623201 | 0.742356514 | 0.084999091 | 6.83E-09 | North_western |
| UG_2522 | 0.026569668 | 0.151392614 | 0.741631196 | 0.080406516 | 6.65E-09 | North_western |
| UG_2530 | 0.127256525 | 0.596673681 | 0.266188854 | 0.007976995 | 0.001903945 | North_western |
| UG_2538 | 0.002860887 | 0.592902083 | 0.404009903 | 0.000227127 | 4.64E-16 | North_western |
| UG_2458 | 0.000106016 | 0.000149026 | 0.087221572 | 0.912523386 | 5.14E-16 | North_western |
| UG_2466 | 0.002838663 | 0.579415312 | 0.417498764 | 0.000247261 | 4.03E-16 | North_western |
| UG_2474 | 9.94E-05 | 0.00016479 | 0.100625306 | 0.899110474 | 1.46E-16 | North_western |
| UG_2482 | 8.92E-05 | 0.000118399 | 0.077235121 | 0.922557262 | 3.97E-16 | North_western |
| UG_2490 | 0.000104848 | 0.000171829 | 0.101419175 | 0.898304149 | 1.89E-16 | North_western |
| UG_2498 | 0.002494749 | 0.601397784 | 0.395923563 | 0.000183904 | 1.82E-16 | North_western |
| UG_2506 | 0.000203831 | 0.000476719 | 0.181259152 | 0.818060298 | 2.44E-16 | North_western |
| UG_2514 | 0.006662735 | 0.081475533 | 0.84899136 | 0.062870371 | 6.67E-13 | North_western |
| UG_2451 | 8.61E-05 | 0.00011826 | 0.078970772 | 0.92082489 | 2.81E-16 | North_western |
| UG_2523 | 0.000731198 | 0.002921922 | 0.423541197 | 0.572805682 | 1.62E-15 | North_western |
| UG_2531 | 9.83E-05 | 0.000144899 | 0.089018743 | 0.910738026 | 2.92E-16 | North_western |
| UG_2459 | 0.000123475 | 0.000188068 | 0.09995235 | 0.899736107 | 5.39E-16 | North_western |
| UG_2467 | 0.002888594 | 0.014064864 | 0.644214518 | 0.338832023 | 2.07E-13 | North_western |
| UG_2475 | 0.000111639 | 0.000191598 | 0.108637461 | 0.891059302 | 1.78E-16 | North_western |
| UG_2483 | 0.006921349 | 0.081824697 | 0.846450462 | 0.064803493 | 8.81E-13 | North_western |
| UG_2491 | 0.010239343 | 0.118917867 | 0.821090198 | 0.049752592 | 7.59E-12 | North_western |
| UG_2499 | 8.10E-05 | 0.000102349 | 0.070917716 | 0.928898978 | 3.80E-16 | North_western |
| UG_2507 | 0.003407052 | 0.576883551 | 0.419401477 | 0.000307921 | 1.49E-15 | North_western |
| UG_2515 | 0.00013007 | 0.000200608 | 0.103132201 | 0.896537121 | 6.02E-16 | North_western |
| UG_2452 | 0.000117508 | 0.000176842 | 0.097001018 | 0.902704631 | 4.86E-16 | North_western |
| UG_2524 | 0.000553099 | 0.002220158 | 0.394351897 | 0.602874846 | 5.19E-16 | North_western |
| UG_2532 | 0.000103124 | 0.000147624 | 0.087955204 | 0.911794048 | 4.15E-16 | North_western |
| UG_2460 | 0.006628448 | 0.082618786 | 0.849668852 | 0.061083914 | 6.22E-13 | North_western |
| UG_2468 | 0.006940728 | 0.082787982 | 0.846566349 | 0.063704941 | 8.76E-13 | North_western |
| UG_2476 | 0.006611498 | 0.082579534 | 0.849830439 | 0.060978529 | 6.11E-13 | North_western |

| | | | | | | |
|---|---|---|---|---|---|---|
| UG_2484 | 0.000112235 | 0.000163868 | 0.092532005 | 0.907191891 | 4.97E-16 | North_western |
| UG_2492 | 0.000105001 | 0.000151795 | 0.089420153 | 0.91032305 | 4.16E-16 | North_western |
| UG_2500 | 0.000105592 | 0.000173563 | 0.101981382 | 0.897739463 | 1.91E-16 | North_western |
| UG_2508 | 0.000130543 | 0.000178515 | 0.091487456 | 0.908203485 | 1.29E-15 | North_western |
| UG_2516 | 0.07215956 | 0.655143392 | 0.269020091 | 0.003651895 | 2.51E-05 | North_western |
| UG_2539 | 0.000134001 | 0.00020161 | 0.10168388 | 0.897980509 | 7.82E-16 | North_western |
| UG_2611 | 0.008002052 | 0.132650456 | 0.827745687 | 0.031601805 | 9.85E-13 | Northern |
| UG_2619 | 0.003178871 | 0.048452727 | 0.876642754 | 0.071725647 | 8.52E-15 | Northern |
| UG_2627 | 0.004223002 | 0.135019832 | 0.845258533 | 0.015498634 | 8.70E-15 | Northern |
| UG_2547 | 0.000119666 | 0.000186712 | 0.101250504 | 0.898443118 | 4.14E-16 | North_western |
| UG_2555 | 0.000164341 | 0.000254494 | 0.112613453 | 0.886967712 | 1.37E-15 | North_western |
| UG_2563 | 0.006879385 | 0.081535813 | 0.846780887 | 0.064803914 | 8.48E-13 | North_western |
| UG_2571 | 0.000128273 | 0.000185368 | 0.096117031 | 0.903569328 | 8.58E-16 | North_western |
| UG_2579 | 0.00011552 | 0.000147428 | 0.081580981 | 0.918156071 | 1.28E-15 | North_western |
| UG_2587 | 0.000389365 | 0.001311401 | 0.309600209 | 0.688699026 | 3.40E-16 | North_western |
| UG_2595 | 0.002545542 | 0.031233445 | 0.851360616 | 0.114860397 | 4.91E-15 | Northern |
| UG_2603 | 0.000832369 | 0.004416376 | 0.530512205 | 0.46423905 | 6.83E-16 | Northern |
| UG_2540 | 0.007132114 | 0.084228925 | 0.845094892 | 0.063544069 | 1.04E-12 | North_western |
| UG_2612 | 0.006133718 | 0.111967002 | 0.849014575 | 0.032884705 | 1.87E-13 | Northern |
| UG_2620 | 0.003531742 | 0.118426998 | 0.861582268 | 0.016458991 | 2.93E-15 | Northern |
| UG_2628 | 0.005731177 | 0.131266036 | 0.840303085 | 0.022699703 | 8.54E-14 | Northern |
| UG_2548 | 0.000125922 | 0.000187452 | 0.098372263 | 0.901314363 | 6.67E-16 | North_western |
| UG_2556 | 0.000105079 | 0.000153365 | 0.090312628 | 0.909428929 | 3.93E-16 | North_western |
| UG_2564 | 0.001135797 | 0.002920747 | 0.339898809 | 0.656044647 | 9.51E-14 | North_western |
| UG_2572 | 0.000474016 | 0.000909566 | 0.199974065 | 0.798642354 | 1.82E-14 | North_western |
| UG_2580 | 0.000229941 | 0.000467155 | 0.165206145 | 0.834096759 | 8.87E-16 | North_western |
| UG_2588 | 0.001753583 | 0.007337783 | 0.539025642 | 0.451882993 | 4.68E-14 | North_western |
| UG_2596 | 0.005003561 | 0.109117667 | 0.85801829 | 0.027860481 | 4.37E-14 | Northern |
| UG_2604 | 0.000697071 | 0.002938863 | 0.434997529 | 0.561366537 | 1.02E-15 | Northern |

| | | | | | | |
|---|---|---|---|---|---|---|
| UG_2541 | 0.000183149 | 0.000283482 | 0.11699723 | 0.882536139 | 2.03E-15 | North_western |
| UG_2613 | 0.004114226 | 0.142559107 | 0.839781755 | 0.013544913 | 6.60E-15 | Northern |
| UG_2621 | 0.004690226 | 0.128356792 | 0.847816676 | 0.019136306 | 2.04E-14 | Northern |
| UG_2629 | 0.000716189 | 0.003162084 | 0.452738777 | 0.543382949 | 8.98E-16 | Northern |
| UG_2549 | 8.36E-05 | 0.000104574 | 0.071010162 | 0.92880171 | 4.54E-16 | North_western |
| UG_2557 | 0.006766474 | 0.083123692 | 0.848410958 | 0.061698876 | 7.17E-13 | North_western |
| UG_2565 | 0.002647359 | 0.630272912 | 0.366919739 | 0.00015999 | 3.46E-16 | North_western |
| UG_2573 | 8.21E-05 | 0.000103746 | 0.071244706 | 0.928569436 | 4.01E-16 | North_western |
| UG_2581 | 0.000134308 | 0.000186979 | 0.094121008 | 0.905557706 | 1.28E-15 | North_western |
| UG_2589 | 9.87E-05 | 0.00011187 | 0.068202215 | 0.931587215 | 1.54E-15 | North_western |
| UG_2597 | 0.002188381 | 0.02565731 | 0.838706241 | 0.133448069 | 2.67E-15 | Northern |
| UG_2605 | 0.001971974 | 0.026266107 | 0.854297597 | 0.117464322 | 1.11E-15 | Northern |
| UG_2542 | 0.004630579 | 0.051504817 | 0.850383761 | 0.093480843 | 1.27E-13 | North_western |
| UG_2614 | 0.002382135 | 0.020586456 | 0.783888535 | 0.193142874 | 1.03E-14 | Northern |
| UG_2622 | 0.003371638 | 0.096913635 | 0.876888785 | 0.022825942 | 2.99E-15 | Northern |
| UG_2630 | 0.001342191 | 0.006558362 | 0.559274683 | 0.432824765 | 7.44E-15 | Northern |
| UG_2550 | 0.003230443 | 0.622651147 | 0.373907813 | 0.000210597 | 1.38E-15 | North_western |
| UG_2558 | 0.001619532 | 0.006460827 | 0.51400672 | 0.47791292 | 4.20E-14 | North_western |
| UG_2566 | 0.007055079 | 0.082062437 | 0.845172153 | 0.065710331 | 1.01E-12 | North_western |
| UG_2574 | 9.02E-05 | 0.000127414 | 0.082640956 | 0.917141434 | 2.79E-16 | North_western |
| UG_2582 | 0.003204193 | 0.571173118 | 0.425323427 | 0.000299261 | 9.22E-16 | North_western |
| UG_2590 | 8.30E-05 | 0.000105063 | 0.071686112 | 0.928125866 | 4.10E-16 | North_western |
| UG_2598 | 0.006843456 | 0.11176762 | 0.844426883 | 0.036962041 | 4.23E-13 | Northern |
| UG_2606 | 0.000396356 | 0.000981601 | 0.238168293 | 0.760453751 | 2.08E-15 | Northern |
| UG_2543 | 9.72E-05 | 0.000132648 | 0.081982866 | 0.917787256 | 4.54E-16 | North_western |
| UG_2615 | 0.00556389 | 0.101971969 | 0.857200634 | 0.035263507 | 1.09E-13 | Northern |
| UG_2623 | 0.000113418 | 0.000145081 | 0.081226648 | 0.918514853 | 1.18E-15 | Northern |
| UG_2631 | 0.004814165 | 0.132331157 | 0.844299677 | 0.018555001 | 2.34E-14 | Northern |
| UG_2551 | 9.70E-05 | 0.000127616 | 0.078922973 | 0.920852383 | 5.67E-16 | North_western |

| | | | | | | |
|---|---|---|---|---|---|---|
| UG_2559 | 0.000129829 | 0.000192222 | 0.098921978 | 0.900755971 | 7.71E-16 | North_western |
| UG_2567 | 0.00018511 | 0.000319147 | 0.130639604 | 0.86885614 | 1.09E-15 | North_western |
| UG_2575 | 0.006916521 | 0.083357669 | 0.846976028 | 0.062749782 | 8.40E-13 | North_western |
| UG_2583 | 0.000107556 | 0.000130799 | 0.07565833 | 0.924103314 | 1.34E-15 | North_western |
| UG_2591 | 0.000161864 | 0.000289456 | 0.12915436 | 0.87039432 | 5.33E-16 | North_western |
| UG_2599 | 0.007798461 | 0.127937991 | 0.83132671 | 0.032936837 | 8.68E-13 | Northern |
| UG_2607 | 0.002224808 | 0.022007835 | 0.80837554 | 0.167391817 | 4.83E-15 | Northern |
| UG_2544 | 0.000102407 | 0.0001359 | 0.08123329 | 0.918528403 | 6.51E-16 | North_western |
| UG_2616 | 0.000706003 | 0.003009405 | 0.439906427 | 0.556378165 | 1.01E-15 | Northern |
| UG_2624 | 0.00187848 | 0.019847236 | 0.811364 | 0.166910284 | 1.75E-15 | Northern |
| UG_2632 | 0.000882193 | 0.003158536 | 0.410259842 | 0.58569943 | 6.04E-15 | Northern |
| UG_2552 | 8.49E-05 | 0.000106313 | 0.071454091 | 0.928354672 | 4.80E-16 | North_western |
| UG_2560 | 8.62E-05 | 0.000115934 | 0.077312407 | 0.922485455 | 3.22E-16 | North_western |
| UG_2568 | 0.000247383 | 0.000647326 | 0.214715709 | 0.784389582 | 2.58E-16 | North_western |
| UG_2576 | 8.41E-05 | 0.00010778 | 0.072937836 | 0.926870299 | 3.99E-16 | North_western |
| UG_2584 | 0.000355966 | 0.000802929 | 0.211199274 | 0.787641831 | 2.40E-15 | North_western |
| UG_2592 | 0.000427504 | 0.001191672 | 0.270930356 | 0.727450469 | 1.41E-15 | North_western |
| UG_2600 | 0.00059362 | 0.002780886 | 0.450278058 | 0.546347436 | 3.12E-16 | Northern |
| UG_2608 | 0.00230012 | 0.027639201 | 0.844715118 | 0.125345561 | 3.17E-15 | Northern |
| UG_2545 | 9.88E-05 | 0.000138994 | 0.085075747 | 0.914686453 | 3.97E-16 | North_western |
| UG_2617 | 0.005515275 | 0.112807245 | 0.852659261 | 0.029018219 | 8.42E-14 | Northern |
| UG_2625 | 0.0045241 | 0.158030657 | 0.825216065 | 0.012229178 | 1.13E-14 | Northern |
| UG_2553 | 8.06E-05 | 9.75E-05 | 0.067658771 | 0.932163119 | 4.95E-16 | North_western |
| UG_2561 | 0.006810243 | 0.080137909 | 0.846985063 | 0.066066785 | 8.17E-13 | North_western |
| UG_2569 | 0.000115985 | 0.000164708 | 0.091052264 | 0.908667042 | 6.66E-16 | North_western |
| UG_2577 | 0.000213766 | 0.000450216 | 0.166766064 | 0.832569954 | 5.46E-16 | North_western |
| UG_2585 | 7.66E-05 | 8.71E-05 | 0.062283594 | 0.937552769 | 6.10E-16 | North_western |
| UG_2593 | 0.002749923 | 0.604373851 | 0.392675606 | 0.00020062 | 3.77E-16 | North_western |
| UG_2601 | 0.004822224 | 0.134817018 | 0.842430589 | 0.017930169 | 2.30E-14 | Northern |

| | | | | | | |
|---|---|---|---|---|---|---|
| UG_2609 | 0.003815719 | 0.115101782 | 0.862197146 | 0.018885353 | 5.42E-15 | Northern |
| UG_2546 | 0.000106131 | 0.000140158 | 0.081893773 | 0.917859938 | 7.63E-16 | North_western |
| UG_2618 | 0.006149143 | 0.117094505 | 0.846412851 | 0.030343501 | 1.76E-13 | Northern |
| UG_2626 | 0.003981699 | 0.137870079 | 0.844186654 | 0.013961568 | 5.47E-15 | Northern |
| UG_2554 | 0.000114429 | 0.000176968 | 0.098747097 | 0.900961507 | 3.72E-16 | North_western |
| UG_2562 | 0.002018135 | 0.008686093 | 0.564586974 | 0.424708799 | 7.46E-14 | North_western |
| UG_2570 | 6.70E-05 | 7.95E-05 | 0.06190339 | 0.937950099 | 2.91E-16 | North_western |
| UG_2578 | 0.007258543 | 0.0836954 | 0.843692784 | 0.065353273 | 1.20E-12 | North_western |
| UG_2586 | 9.93E-05 | 0.000125417 | 0.076353928 | 0.923421322 | 7.98E-16 | North_western |
| UG_2594 | 0.044234076 | 0.324010316 | 0.604748156 | 0.027007332 | 1.20E-07 | North_western |
| UG_2602 | 0.001949602 | 0.019708467 | 0.804583741 | 0.17375819 | 2.41E-15 | Northern |
| UG_2610 | 0.001709586 | 0.020732219 | 0.832446572 | 0.145111623 | 7.29E-16 | Northern |
| UG_2633 | 0.002730581 | 0.029408884 | 0.834304741 | 0.133555795 | 1.01E-14 | Northern |
| UG_2705 | 0.0031169 | 0.11079081 | 0.8697652 | 0.016327091 | 1.32E-15 | Northern |
| UG_2713 | 0.003385859 | 0.104587254 | 0.87212927 | 0.019897617 | 2.68E-15 | Northern |
| UG_2721 | 0.003615446 | 0.131984037 | 0.850705496 | 0.013695021 | 2.90E-15 | Northern |
| UG_2641 | 0.002277583 | 0.015454376 | 0.71893403 | 0.263334011 | 1.96E-14 | Northern |
| UG_2649 | 0.004038202 | 0.124643648 | 0.854077478 | 0.017240671 | 7.16E-15 | Northern |
| UG_2657 | 0.003507329 | 0.039779819 | 0.849361455 | 0.107351397 | 2.99E-14 | Northern |
| UG_2665 | 0.003622599 | 0.139190129 | 0.844819978 | 0.012367294 | 2.71E-15 | Northern |
| UG_2673 | 0.003741335 | 0.113002997 | 0.864109421 | 0.019146248 | 4.85E-15 | Northern |
| UG_2681 | 0.003775855 | 0.15363014 | 0.83196057 | 0.010633435 | 3.15E-15 | Northern |
| UG_2689 | 0.000699852 | 0.003106086 | 0.451772718 | 0.544421345 | 7.96E-16 | Northern |
| UG_2697 | 0.003755472 | 0.121604734 | 0.857922487 | 0.016717307 | 4.39E-15 | Northern |
| UG_2634 | 0.001223522 | 0.00564058 | 0.529124962 | 0.464010937 | 6.60E-15 | Northern |
| UG_2706 | 0.003312086 | 0.036879358 | 0.845691999 | 0.114116558 | 2.37E-14 | Northern |
| UG_2714 | 0.003998115 | 0.114054935 | 0.861745041 | 0.020201908 | 7.75E-15 | Northern |
| UG_2722 | 0.004224445 | 0.11074524 | 0.862381966 | 0.02264835 | 1.22E-14 | Northern |
| UG_2642 | 0.002811389 | 0.041952241 | 0.875070186 | 0.080166184 | 4.80E-15 | Northern |

| | | | | | | |
|---|---|---|---|---|---|---|
| UG_2650 | 0.000827953 | 0.003921013 | 0.492271405 | 0.502979629 | 1.15E-15 | Northern |
| UG_2658 | 0.002666639 | 0.042977 | 0.881102435 | 0.073253926 | 3.03E-15 | Northern |
| UG_2666 | 0.003224385 | 0.014370968 | 0.62817043 | 0.354234218 | 4.85E-13 | Northern |
| UG_2674 | 0.003969984 | 0.156378144 | 0.828814613 | 0.010837259 | 4.42E-15 | Northern |
| UG_2682 | 0.002904226 | 0.117982852 | 0.86570059 | 0.013412332 | 7.07E-16 | Northern |
| UG_2690 | 0.004473832 | 0.147022567 | 0.83454933 | 0.013954271 | 1.16E-14 | Northern |
| UG_2698 | 0.003132152 | 0.127980836 | 0.856444291 | 0.012442721 | 1.07E-15 | Northern |
| UG_2635 | 0.000135132 | 0.000191982 | 0.096279041 | 0.903393846 | 1.15E-15 | Northern |
| UG_2707 | 0.003316582 | 0.088075783 | 0.881877664 | 0.026729971 | 3.17E-15 | Northern |
| UG_2715 | 0.003979109 | 0.132128479 | 0.848732586 | 0.015159826 | 5.83E-15 | Northern |
| UG_2723 | 0.004822582 | 0.13815222 | 0.839926194 | 0.017099004 | 2.21E-14 | Northern |
| UG_2643 | 0.003244137 | 0.066134594 | 0.887178671 | 0.043442598 | 4.85E-15 | Northern |
| UG_2651 | 0.001563283 | 0.012275573 | 0.724505573 | 0.261655571 | 2.05E-15 | Northern |
| UG_2659 | 0.002642736 | 0.064767539 | 0.896180899 | 0.036408826 | 1.11E-15 | Northern |
| UG_2667 | 0.003952005 | 0.143448998 | 0.839790421 | 0.012808575 | 4.87E-15 | Northern |
| UG_2675 | 0.000162739 | 0.000224881 | 0.100133588 | 0.899478792 | 2.68E-15 | Northern |
| UG_2683 | 0.003695522 | 0.118280736 | 0.860699352 | 0.01732439 | 4.09E-15 | Northern |
| UG_2691 | 0.004470755 | 0.156253023 | 0.826925921 | 0.012350301 | 1.05E-14 | Northern |
| UG_2699 | 0.004618589 | 0.141101767 | 0.838618532 | 0.015661112 | 1.56E-14 | Northern |
| UG_2636 | 0.000701551 | 0.003046376 | 0.445077422 | 0.551174652 | 8.98E-16 | Northern |
| UG_2708 | 0.005289076 | 0.12444671 | 0.847184233 | 0.023079981 | 5.19E-14 | Northern |
| UG_2716 | 0.003768884 | 0.116912343 | 0.861227049 | 0.018091724 | 4.82E-15 | Northern |
| UG_2724 | 0.003573387 | 0.113503143 | 0.864851448 | 0.018072021 | 3.44E-15 | Northern |
| UG_2644 | 0.003866092 | 0.137797555 | 0.844799445 | 0.013536908 | 4.42E-15 | Northern |
| UG_2652 | 0.003445651 | 0.102929755 | 0.872735161 | 0.020889432 | 3.13E-15 | Northern |
| UG_2660 | 0.004122543 | 0.07575556 | 0.876211479 | 0.043910418 | 2.15E-14 | Northern |
| UG_2668 | 0.006263649 | 0.127234044 | 0.840037392 | 0.026464915 | 1.73E-13 | Northern |
| UG_2676 | 0.004422862 | 0.138467921 | 0.841600516 | 0.015508701 | 1.17E-14 | Northern |
| UG_2684 | 0.001898734 | 0.018497324 | 0.794843516 | 0.184760426 | 2.39E-15 | Northern |

| | | | | | | |
|---|---|---|---|---|---|---|
| UG_2692 | 0.003637934 | 0.12789444 | 0.853810817 | 0.014656808 | 3.20E-15 | Northern |
| UG_2700 | 0.006016431 | 0.096668544 | 0.855218816 | 0.042096209 | 2.16E-13 | Northern |
| UG_2637 | 0.004547405 | 0.111521781 | 0.859754078 | 0.024176736 | 2.08E-14 | Northern |
| UG_2709 | 0.004520946 | 0.132594686 | 0.845606573 | 0.017277795 | 1.48E-14 | Northern |
| UG_2717 | 0.00457131 | 0.137493372 | 0.841641531 | 0.016293787 | 1.51E-14 | Northern |
| UG_2725 | 0.003272772 | 0.085181649 | 0.883541577 | 0.028004002 | 3.07E-15 | Northern |
| UG_2645 | 0.000194298 | 0.000330156 | 0.131007147 | 0.868468399 | 1.42E-15 | Northern |
| UG_2653 | 0.001061413 | 0.006407761 | 0.601910759 | 0.390620067 | 1.07E-15 | Northern |
| UG_2661 | 0.00521984 | 0.112365315 | 0.854829618 | 0.027585226 | 5.65E-14 | Northern |
| UG_2669 | 0.005110269 | 0.146721042 | 0.831996347 | 0.016172342 | 3.07E-14 | Northern |
| UG_2677 | 0.004940935 | 0.580275294 | 0.414330736 | 0.000453035 | 2.25E-14 | Northern |
| UG_2685 | 0.008967673 | 0.183983181 | 0.788326386 | 0.018722761 | 1.37E-12 | Northern |
| UG_2693 | 0.001850365 | 0.016671327 | 0.774937572 | 0.206540736 | 2.74E-15 | Northern |
| UG_2701 | 0.002654642 | 0.05132677 | 0.891539691 | 0.054478898 | 1.92E-15 | Northern |
| UG_2638 | 0.001957755 | 0.019721979 | 0.804120577 | 0.174199689 | 2.48E-15 | Northern |
| UG_2710 | 0.009296467 | 0.189216061 | 0.783125601 | 0.018361871 | 1.71E-12 | Northern |
| UG_2718 | 0.001488723 | 0.014137 | 0.770438905 | 0.213935373 | 8.32E-16 | Northern |
| UG_2726 | 0.005134767 | 0.160740214 | 0.820570884 | 0.013554135 | 2.77E-14 | Northern |
| UG_2646 | 0.003105026 | 0.055626627 | 0.885478423 | 0.055789924 | 5.15E-15 | Northern |
| UG_2654 | 0.004084439 | 0.121392736 | 0.856162974 | 0.018359852 | 8.13E-15 | Northern |
| UG_2662 | 0.002593797 | 0.050456441 | 0.892191623 | 0.054758138 | 1.68E-15 | Northern |
| UG_2670 | 0.002793624 | 0.092875139 | 0.884142447 | 0.02018879 | 8.15E-16 | Northern |
| UG_2678 | 0.060008085 | 0.397068753 | 0.5219613 | 0.020960582 | 1.28E-06 | Northern |
| UG_2686 | 0.116353268 | 0.557416797 | 0.314962523 | 0.0106873 | 0.000580112 | Northern |
| UG_2694 | 0.003688296 | 0.119122616 | 0.860133788 | 0.0170553 | 3.98E-15 | Northern |
| UG_2702 | 0.004092112 | 0.125083067 | 0.85345418 | 0.017370641 | 7.84E-15 | Northern |
| UG_2639 | 0.001893029 | 0.015920457 | 0.759797033 | 0.222389481 | 3.87E-15 | Northern |
| UG_2711 | 0.004258332 | 0.132469086 | 0.84704295 | 0.016229632 | 9.54E-15 | Northern |
| UG_2719 | 0.003969695 | 0.130131037 | 0.850324319 | 0.015574949 | 5.88E-15 | Northern |

| | | | | | | |
|---|---|---|---|---|---|---|
| UG_2647 | 0.000928815 | 0.004113906 | 0.483723282 | 0.511233997 | 2.55E-15 | Northern |
| UG_2655 | 0.002034359 | 0.017113761 | 0.765699479 | 0.2151524 | 5.40E-15 | Northern |
| UG_2663 | 0.004356701 | 0.152852025 | 0.830241367 | 0.012549906 | 9.01E-15 | Northern |
| UG_2671 | 0.002127255 | 0.014384414 | 0.711556606 | 0.271931725 | 1.45E-14 | Northern |
| UG_2679 | 0.003717553 | 0.120145705 | 0.859214589 | 0.016922153 | 4.16E-15 | Northern |
| UG_2687 | 9.89E-05 | 0.000118439 | 0.07223087 | 0.92755182 | 1.09E-15 | Northern |
| UG_2695 | 0.009431821 | 0.192786787 | 0.779842787 | 0.017938605 | 1.86E-12 | Northern |
| UG_2703 | 0.003098371 | 0.600598982 | 0.396067564 | 0.000235083 | 8.70E-16 | Northern |
| UG_2640 | 0.005669982 | 0.112390903 | 0.851859977 | 0.030079138 | 1.04E-13 | Northern |
| UG_2712 | 0.004694123 | 0.160202654 | 0.822719948 | 0.012383276 | 1.45E-14 | Northern |
| UG_2720 | 0.003926191 | 0.120371873 | 0.857819341 | 0.017882594 | 6.18E-15 | Northern |
| UG_2648 | 0.006339614 | 0.130912186 | 0.837359056 | 0.025389145 | 1.80E-13 | Northern |
| UG_2656 | 0.003497834 | 0.077878832 | 0.883351725 | 0.03527161 | 6.01E-15 | Northern |
| UG_2664 | 0.000745107 | 0.003375222 | 0.465535911 | 0.53034376 | 9.28E-16 | Northern |
| UG_2672 | 0.002331769 | 0.026605737 | 0.837029537 | 0.134032958 | 3.93E-15 | Northern |
| UG_2680 | 0.003856724 | 0.12636066 | 0.853802283 | 0.015980333 | 5.00E-15 | Northern |
| UG_2688 | 0.004210923 | 0.144579594 | 0.837700027 | 0.013509455 | 7.65E-15 | Northern |
| UG_2696 | 0.003795727 | 0.14964171 | 0.835290304 | 0.011272259 | 3.40E-15 | Northern |
| UG_2704 | 0.003695241 | 0.650615226 | 0.34549019 | 0.000199343 | 4.57E-15 | Northern |
| UG_2727 | 0.004618154 | 0.134063382 | 0.844014679 | 0.017303785 | 1.69E-14 | Northern |
| UG_2799 | 0.08495657 | 0.691822323 | 0.220179958 | 0.002876994 | 0.000164155 | Northern |
| UG_2807 | 0.006969565 | 0.110882834 | 0.8439311 | 0.038216501 | 4.92E-13 | Northern |
| UG_2815 | 0.002170642 | 0.022018651 | 0.811944145 | 0.163866562 | 3.97E-15 | Northern |
| UG_2735 | 0.011761525 | 0.212716145 | 0.757091652 | 0.018430678 | 8.36E-12 | Northern |
| UG_2743 | 0.00158612 | 0.011469905 | 0.702304019 | 0.284639956 | 2.99E-15 | Northern |
| UG_2751 | 0.049496345 | 0.332145678 | 0.590169447 | 0.028188247 | 2.83E-07 | Northern |
| UG_2759 | 0.06089621 | 0.405615805 | 0.513495904 | 0.019990626 | 1.45E-06 | Northern |
| UG_2767 | 0.04246383 | 0.329079806 | 0.603435291 | 0.025020986 | 8.72E-08 | Northern |
| UG_2775 | 0.056818808 | 0.390276989 | 0.53194603 | 0.020957343 | 8.29E-07 | Northern |

| | | | | | | |
|---|---|---|---|---|---|---|
| UG_2783 | 0.000589983 | 0.002303444 | 0.392948364 | 0.604158208 | 7.75E-16 | Northern |
| UG_2791 | 0.009677414 | 0.118288202 | 0.824567867 | 0.047466517 | 5.02E-12 | Northern |
| UG_2728 | 0.003942399 | 0.151906226 | 0.832751594 | 0.011399781 | 4.39E-15 | Northern |
| UG_2800 | 0.004049239 | 0.070325544 | 0.87658141 | 0.049043808 | 2.21E-14 | Northern |
| UG_2808 | 0.003078473 | 0.122839693 | 0.860860971 | 0.013220863 | 1.01E-15 | Northern |
| UG_2816 | 0.004478315 | 0.114641758 | 0.858295248 | 0.02258468 | 1.77E-14 | Northern |
| UG_2736 | 0.01795134 | 0.24996691 | 0.711890458 | 0.020191292 | 1.61E-10 | Northern |
| UG_2744 | 0.000778221 | 0.002471036 | 0.361276591 | 0.635474152 | 6.93E-15 | Northern |
| UG_2752 | 9.15E-05 | 0.000106537 | 0.068205042 | 0.931596969 | 9.86E-16 | Northern |
| UG_2760 | 0.059578143 | 0.396717553 | 0.522822454 | 0.020880641 | 1.21E-06 | Northern |
| UG_2768 | 0.047197199 | 0.422551026 | 0.51614222 | 0.014109347 | 2.08E-07 | Northern |
| UG_2776 | 0.059438085 | 0.394158157 | 0.525184883 | 0.021217691 | 1.18E-06 | Northern |
| UG_2784 | 0.002182508 | 0.016728656 | 0.747948822 | 0.233140014 | 1.03E-14 | Northern |
| UG_2792 | 0.006180484 | 0.374170233 | 0.617251862 | 0.002397421 | 5.62E-14 | Northern |
| UG_2729 | 0.004973993 | 0.146214864 | 0.832994994 | 0.01581615 | 2.54E-14 | Northern |
| UG_2801 | 0.004478164 | 0.040309983 | 0.823957821 | 0.131254032 | 1.92E-13 | Northern |
| UG_2809 | 0.000672526 | 0.002781362 | 0.424946929 | 0.571599182 | 9.70E-16 | Northern |
| UG_2817 | 0.007669431 | 0.127232716 | 0.832388875 | 0.032708978 | 7.75E-13 | Northern |
| UG_2737 | 0.000101293 | 0.000119457 | 0.071718298 | 0.928060951 | 1.32E-15 | Northern |
| UG_2745 | 0.000953062 | 0.004654121 | 0.518899415 | 0.475493402 | 1.78E-15 | Northern |
| UG_2753 | 0.049895308 | 0.332014173 | 0.589667189 | 0.028423029 | 3.01E-07 | Northern |
| UG_2761 | 0.041360077 | 0.335881675 | 0.599561776 | 0.023196402 | 7.09E-08 | Northern |
| UG_2769 | 0.043330054 | 0.337756772 | 0.59501103 | 0.023902043 | 1.01E-07 | Northern |
| UG_2777 | 0.037174572 | 0.324988676 | 0.615103914 | 0.022732806 | 3.18E-08 | Northern |
| UG_2785 | 0.001628895 | 0.015492837 | 0.778017965 | 0.204860303 | 1.26E-15 | Northern |
| UG_2793 | 0.006706187 | 0.108092604 | 0.846724653 | 0.038476556 | 3.88E-13 | Northern |
| UG_2730 | 0.004111561 | 0.133966315 | 0.846633002 | 0.015289121 | 7.25E-15 | Northern |
| UG_2802 | 0.001186872 | 0.006654269 | 0.590480581 | 0.401678279 | 2.38E-15 | Northern |
| UG_2810 | 0.007754345 | 0.128969171 | 0.831023241 | 0.032253243 | 8.20E-13 | Northern |

| | | | | | | |
|---|---|---|---|---|---|---|
| UG_2818 | 0.001708196 | 0.018537072 | 0.810846408 | 0.168908324 | 1.02E-15 | Northern |
| UG_2738 | 0.045997262 | 0.339738283 | 0.589358831 | 0.024905465 | 1.60E-07 | Northern |
| UG_2746 | 9.90E-05 | 0.000114942 | 0.070006728 | 0.929779377 | 1.33E-15 | Northern |
| UG_2754 | 0.057155146 | 0.3386763 | 0.573609425 | 0.030558269 | 8.59E-07 | Northern |
| UG_2762 | 0.000116385 | 0.00014876 | 0.081931951 | 0.917802904 | 1.30E-15 | Northern |
| UG_2770 | 0.044429481 | 0.324328805 | 0.604188036 | 0.027053554 | 1.24E-07 | Northern |
| UG_2778 | 0.041415437 | 0.332271151 | 0.602453174 | 0.023860166 | 7.19E-08 | Northern |
| UG_2786 | 0.006460991 | 0.093908437 | 0.851966028 | 0.047664544 | 3.90E-13 | Northern |
| UG_2794 | 0.00318808 | 0.10718854 | 0.871814534 | 0.017808847 | 1.65E-15 | Northern |
| UG_2731 | 0.004875708 | 0.144201004 | 0.835014646 | 0.015908642 | 2.24E-14 | Northern |
| UG_2803 | 0.000563037 | 0.002291592 | 0.400469995 | 0.596675376 | 5.18E-16 | Northern |
| UG_2811 | 0.005789065 | 0.11016039 | 0.852148174 | 0.031902371 | 1.26E-13 | Northern |
| UG_2819 | 0.002543032 | 0.040202478 | 0.879439772 | 0.077814719 | 2.50E-15 | Northern |
| UG_2739 | 0.051337422 | 0.335792779 | 0.584508194 | 0.028361232 | 3.74E-07 | Northern |
| UG_2747 | 0.001190421 | 0.00595062 | 0.552791892 | 0.440067067 | 4.04E-15 | Northern |
| UG_2755 | 0.00650011 | 0.14999639 | 0.823472032 | 0.020031468 | 1.73E-13 | Northern |
| UG_2763 | 0.003624868 | 0.081748081 | 0.881043555 | 0.033583496 | 7.08E-15 | Northern |
| UG_2771 | 0.050786436 | 0.333563454 | 0.587096453 | 0.028553312 | 3.44E-07 | Northern |
| UG_2779 | 0.123567123 | 0.63581566 | 0.232736521 | 0.005551092 | 0.002329603 | Northern |
| UG_2787 | 0.003010376 | 0.131093328 | 0.854524935 | 0.01137136 | 7.72E-16 | Northern |
| UG_2795 | 0.087456847 | 0.668720396 | 0.240040292 | 0.003629666 | 0.000152799 | Northern |
| UG_2732 | 0.005137398 | 0.154326317 | 0.825822251 | 0.014714033 | 2.96E-14 | Northern |
| UG_2804 | 0.002606516 | 0.08519257 | 0.890233609 | 0.021967305 | 5.78E-16 | Northern |
| UG_2812 | 8.75E-05 | 0.000100949 | 0.066423797 | 0.933387731 | 8.97E-16 | Northern |
| UG_2820 | 0.003306545 | 0.083903525 | 0.883695605 | 0.029094326 | 3.41E-15 | Northern |
| UG_2740 | 0.043365977 | 0.322196911 | 0.607559903 | 0.026877105 | 1.03E-07 | Northern |
| UG_2748 | 9.25E-05 | 0.000108056 | 0.068664928 | 0.931134474 | 1.01E-15 | Northern |
| UG_2756 | 0.051807348 | 0.341605651 | 0.579191556 | 0.027395045 | 3.99E-07 | Northern |
| UG_2764 | 0.000970757 | 0.003928145 | 0.459499794 | 0.535601304 | 4.77E-15 | Northern |

| | | | | | | |
|---|---|---|---|---|---|---|
| UG_2772 | 0.002417894 | 0.020517236 | 0.780785205 | 0.196279665 | 1.18E-14 | Northern |
| UG_2780 | 0.010226394 | 0.528608944 | 0.459738674 | 0.001425989 | 3.39E-12 | Northern |
| UG_2788 | 0.004155295 | 0.149737655 | 0.833685553 | 0.012421496 | 6.58E-15 | Northern |
| UG_2796 | 0.086758449 | 0.67476773 | 0.234900746 | 0.003418504 | 0.000154571 | Northern |
| UG_2733 | 0.010427571 | 0.194946444 | 0.775134049 | 0.019491936 | 3.84E-12 | Northern |
| UG_2805 | 0.004182012 | 0.535825867 | 0.459480864 | 0.000511257 | 5.25E-15 | Northern |
| UG_2813 | 0.00116025 | 0.004644019 | 0.47645299 | 0.517742742 | 1.04E-14 | Northern |
| UG_2741 | 9.35E-05 | 0.000106627 | 0.067252138 | 0.932547687 | 1.23E-15 | Northern |
| UG_2749 | 0.002086374 | 0.016278603 | 0.748741569 | 0.232893454 | 7.89E-15 | Northern |
| UG_2757 | 0.002258337 | 0.016891243 | 0.744478254 | 0.236372166 | 1.32E-14 | Northern |
| UG_2765 | 0.003704266 | 0.09653063 | 0.874347701 | 0.025417402 | 6.00E-15 | Northern |
| UG_2773 | 0.003679337 | 0.158373907 | 0.828223263 | 0.009723493 | 2.50E-15 | Northern |
| UG_2781 | 0.141362879 | 0.601426253 | 0.243823191 | 0.007671749 | 0.005715928 | Northern |
| UG_2789 | 9.46E-05 | 0.000108113 | 0.06772866 | 0.932068654 | 1.25E-15 | Northern |
| UG_2797 | 0.069611995 | 0.565712497 | 0.35746671 | 0.007200529 | 8.27E-06 | Northern |
| UG_2734 | 0.008379116 | 0.166554923 | 0.803755833 | 0.021310128 | 9.55E-13 | Northern |
| UG_2806 | 0.003661447 | 0.128154154 | 0.853482919 | 0.01470148 | 3.34E-15 | Northern |
| UG_2814 | 0.003643335 | 0.12596168 | 0.855275801 | 0.015119183 | 3.31E-15 | Northern |
| UG_2742 | 9.64E-05 | 0.000115367 | 0.071483787 | 0.928304431 | 1.01E-15 | Northern |
| UG_2750 | 9.46E-05 | 0.000108272 | 0.067837615 | 0.931959556 | 1.24E-15 | Northern |
| UG_2758 | 9.43E-05 | 0.000107805 | 0.067663814 | 0.932134092 | 1.24E-15 | Northern |
| UG_2766 | 9.23E-05 | 0.000107195 | 0.068205576 | 0.931594897 | 1.04E-15 | Northern |
| UG_2774 | 0.052338 | 0.337712416 | 0.581495864 | 0.028453287 | 4.33E-07 | Northern |
| UG_2782 | 0.000169608 | 0.000271371 | 0.117646148 | 0.881912873 | 1.25E-15 | Northern |
| UG_2790 | 0.000365094 | 0.000952686 | 0.243093905 | 0.755588315 | 1.13E-15 | Northern |
| UG_2798 | 0.065195724 | 0.551177489 | 0.376018831 | 0.007603526 | 4.43E-06 | Northern |
| UG_2821 | 0.000774071 | 0.004025992 | 0.515322571 | 0.479877366 | 5.55E-16 | Northern |
| UG_2893 | 0.003410154 | 0.580774153 | 0.415515639 | 0.000300055 | 1.53E-15 | Eastern |
| UG_2901 | 0.020866088 | 0.602183395 | 0.375138211 | 0.001812304 | 1.01E-09 | Eastern |

| | | | | | | |
|---|---|---|---|---|---|---|
| UG_2909 | 0.008030972 | 0.529837698 | 0.461042782 | 0.001088549 | 5.85E-13 | Eastern |
| UG_2829 | 0.002696406 | 0.044869487 | 0.883357096 | 0.069077011 | 2.97E-15 | Northern |
| UG_2837 | 0.004461842 | 0.121080221 | 0.854173915 | 0.020284022 | 1.56E-14 | Northern |
| UG_2845 | 0.00418168 | 0.124465 | 0.853403466 | 0.017949855 | 9.26E-15 | Northern |
| UG_2853 | 0.004322378 | 0.157127107 | 0.826777019 | 0.011773495 | 8.16E-15 | Northern |
| UG_2861 | 0.003298639 | 0.094250225 | 0.878975729 | 0.023475407 | 2.68E-15 | Northern |
| UG_2869 | 0.002007018 | 0.020063611 | 0.804112688 | 0.173816683 | 2.86E-15 | Northern |
| UG_2877 | 0.003655177 | 0.126639851 | 0.854689245 | 0.015015728 | 3.36E-15 | Northern |
| UG_2885 | 0.002482056 | 0.011044876 | 0.599011401 | 0.387461667 | 1.56E-13 | Northern |
| UG_2822 | 0.002819955 | 0.039782151 | 0.86997096 | 0.087426933 | 5.63E-15 | Northern |
| UG_2894 | 0.01026644 | 0.672624483 | 0.316596835 | 0.000512243 | 9.60E-12 | Eastern |
| UG_2902 | 0.003111381 | 0.623731951 | 0.372956186 | 0.000200482 | 1.06E-15 | Eastern |
| UG_2910 | 0.003140599 | 0.639182031 | 0.357496184 | 0.000181186 | 1.28E-15 | Eastern |
| UG_2830 | 0.002477132 | 0.025452253 | 0.821694082 | 0.150376534 | 7.21E-15 | Northern |
| UG_2838 | 0.005275718 | 0.117645424 | 0.851479389 | 0.025599469 | 5.62E-14 | Northern |
| UG_2846 | 0.004095021 | 0.111595753 | 0.862713273 | 0.021595953 | 9.60E-15 | Northern |
| UG_2854 | 0.003383648 | 0.104860018 | 0.871969458 | 0.019786876 | 2.65E-15 | Northern |
| UG_2862 | 0.00414904 | 0.129680311 | 0.849725137 | 0.016445512 | 8.17E-15 | Northern |
| UG_2870 | 0.005380212 | 0.121877173 | 0.848291496 | 0.02445112 | 6.10E-14 | Northern |
| UG_2878 | 0.003625768 | 0.115815622 | 0.862891198 | 0.017667412 | 3.69E-15 | Northern |
| UG_2886 | 0.000956513 | 0.004863825 | 0.53276157 | 0.461418091 | 1.49E-15 | Northern |
| UG_2823 | 0.004425813 | 0.142768812 | 0.838189767 | 0.014615607 | 1.12E-14 | Northern |
| UG_2895 | 0.017989989 | 0.615987363 | 0.364614002 | 0.001408645 | 3.70E-10 | Eastern |
| UG_2903 | 0.006717843 | 0.561577138 | 0.43098354 | 0.000721479 | 1.88E-13 | Eastern |
| UG_2911 | 0.00315645 | 0.584951782 | 0.411624148 | 0.00026762 | 8.99E-16 | Eastern |
| UG_2831 | 0.004126843 | 0.142127203 | 0.840074518 | 0.013671436 | 6.78E-15 | Northern |
| UG_2839 | 0.001864968 | 0.021087065 | 0.824247099 | 0.152800868 | 1.37E-15 | Northern |
| UG_2847 | 0.000172053 | 0.000231255 | 0.09936029 | 0.900236402 | 3.89E-15 | Northern |
| UG_2855 | 0.00430787 | 0.107828681 | 0.863554879 | 0.024308571 | 1.48E-14 | Northern |

| | | | | | | |
|---|---|---|---|---|---|---|
| UG_2863 | 0.004050956 | 0.11836742 | 0.85848599 | 0.019095634 | 8.00E-15 | Northern |
| UG_2871 | 0.003153662 | 0.06077535 | 0.887276527 | 0.048794461 | 4.73E-15 | Northern |
| UG_2879 | 0.002923744 | 0.10544087 | 0.874904249 | 0.016731137 | 9.02E-16 | Northern |
| UG_2887 | 0.001655715 | 0.015391815 | 0.77378089 | 0.209171579 | 1.47E-15 | Northern |
| UG_2824 | 0.003243831 | 0.048778333 | 0.875602767 | 0.072375069 | 9.76E-15 | Northern |
| UG_2896 | 0.003179315 | 0.617815035 | 0.378791466 | 0.000214183 | 1.18E-15 | Eastern |
| UG_2904 | 0.024537692 | 0.669305056 | 0.304868333 | 0.001288913 | 6.04E-09 | Eastern |
| UG_2912 | 0.003649771 | 0.606785175 | 0.389295346 | 0.000269708 | 2.98E-15 | Eastern |
| UG_2832 | 0.002835325 | 0.093719106 | 0.883273295 | 0.020172274 | 8.93E-16 | Northern |
| UG_2840 | 0.005648737 | 0.117277745 | 0.849398041 | 0.027675477 | 9.36E-14 | Northern |
| UG_2848 | 0.003582403 | 0.118657852 | 0.861107934 | 0.016651811 | 3.24E-15 | Northern |
| UG_2856 | 0.004987066 | 0.055068271 | 0.849694997 | 0.090249666 | 1.89E-13 | Northern |
| UG_2864 | 0.003257545 | 0.110391094 | 0.869109703 | 0.017241659 | 1.83E-15 | Northern |
| UG_2872 | 0.005414173 | 0.121843119 | 0.848115031 | 0.024627677 | 6.39E-14 | Northern |
| UG_2880 | 0.001856971 | 0.022320527 | 0.835155798 | 0.140666704 | 1.12E-15 | Northern |
| UG_2888 | 0.004399438 | 0.119215829 | 0.855805449 | 0.020579283 | 1.45E-14 | Northern |
| UG_2825 | 0.003771425 | 0.110321326 | 0.865700115 | 0.020207133 | 5.36E-15 | Northern |
| UG_2897 | 0.004951823 | 0.559585662 | 0.434938834 | 0.000523682 | 2.02E-14 | Eastern |
| UG_2905 | 0.003170498 | 0.622289106 | 0.374333577 | 0.000206819 | 1.20E-15 | Eastern |
| UG_2913 | 0.003027744 | 0.629528338 | 0.367257314 | 0.000186604 | 9.08E-16 | Eastern |
| UG_2833 | 0.001867866 | 0.016053177 | 0.76413934 | 0.217939617 | 3.37E-15 | Northern |
| UG_2841 | 0.000101648 | 0.000120057 | 0.071922548 | 0.927855747 | 1.32E-15 | Northern |
| UG_2849 | 0.003442788 | 0.117991366 | 0.862440835 | 0.016125011 | 2.45E-15 | Northern |
| UG_2857 | 0.003365038 | 0.09194552 | 0.87959438 | 0.025095062 | 3.25E-15 | Northern |
| UG_2865 | 0.00347881 | 0.099777443 | 0.87438127 | 0.022362478 | 3.56E-15 | Northern |
| UG_2873 | 0.001366092 | 0.008251847 | 0.630858388 | 0.359523673 | 3.17E-15 | Northern |
| UG_2881 | 0.000422742 | 0.001163071 | 0.266907902 | 0.731506285 | 1.45E-15 | Northern |
| UG_2889 | 0.006541343 | 0.633156741 | 0.35987849 | 0.000423426 | 2.51E-13 | Eastern |
| UG_2826 | 0.001606554 | 0.004845999 | 0.423146003 | 0.570401445 | 1.62E-13 | Northern |

| | | | | | | |
|---|---|---|---|---|---|---|
| UG_2898 | 0.003288645 | 0.621940064 | 0.374555404 | 0.000215887 | 1.56E-15 | Eastern |
| UG_2906 | 0.003244525 | 0.626439007 | 0.370110524 | 0.000205944 | 1.46E-15 | Eastern |
| UG_2914 | 0.001543422 | 0.006512326 | 0.526927428 | 0.465016823 | 2.64E-14 | Eastern |
| UG_2834 | 0.001780465 | 0.017843809 | 0.796629071 | 0.183746655 | 1.61E-15 | Northern |
| UG_2842 | 0.004429801 | 0.104027495 | 0.864777781 | 0.026764922 | 1.94E-14 | Northern |
| UG_2850 | 0.003737541 | 0.103572249 | 0.870165514 | 0.022524696 | 5.62E-15 | Northern |
| UG_2858 | 0.000755257 | 0.00385523 | 0.50622103 | 0.489168483 | 5.48E-16 | Northern |
| UG_2866 | 0.001281666 | 0.006970932 | 0.589371433 | 0.40237597 | 3.79E-15 | Northern |
| UG_2874 | 0.004327438 | 0.132608572 | 0.846584089 | 0.016479901 | 1.07E-14 | Northern |
| UG_2882 | 0.003470683 | 0.130527613 | 0.852614259 | 0.013387445 | 2.19E-15 | Northern |
| UG_2890 | 0.002982975 | 0.617074616 | 0.379741744 | 0.000200665 | 7.42E-16 | Eastern |
| UG_2827 | 0.0033261 | 0.072381239 | 0.886205333 | 0.038087329 | 4.82E-15 | Northern |
| UG_2899 | 0.059554495 | 0.70094021 | 0.237361296 | 0.002134921 | 9.08E-06 | Eastern |
| UG_2907 | 0.009827269 | 0.427317848 | 0.560127129 | 0.002727755 | 1.78E-12 | Eastern |
| UG_2835 | 0.005408217 | 0.112283203 | 0.853635037 | 0.028673543 | 7.35E-14 | Northern |
| UG_2843 | 0.004473746 | 0.108715131 | 0.861890763 | 0.02492036 | 1.93E-14 | Northern |
| UG_2851 | 0.002352455 | 0.031316866 | 0.859945361 | 0.106385318 | 2.66E-15 | Northern |
| UG_2859 | 0.005563851 | 0.120999233 | 0.847753306 | 0.025683611 | 7.91E-14 | Northern |
| UG_2867 | 0.00459047 | 0.127016049 | 0.84931172 | 0.019081761 | 1.77E-14 | Northern |
| UG_2875 | 0.003323389 | 0.124085697 | 0.858501794 | 0.014089119 | 1.74E-15 | Northern |
| UG_2883 | 0.003765669 | 0.133104126 | 0.849051302 | 0.014078903 | 3.85E-15 | Northern |
| UG_2891 | 0.008036579 | 0.540274216 | 0.45067458 | 0.001014625 | 6.19E-13 | Eastern |
| UG_2828 | 0.00229099 | 0.030189828 | 0.857847296 | 0.109671886 | 2.40E-15 | Northern |
| UG_2900 | 0.00795428 | 0.523013381 | 0.46790394 | 0.001128399 | 5.28E-13 | Eastern |
| UG_2908 | 0.003392434 | 0.624533579 | 0.371854676 | 0.000219311 | 1.99E-15 | Eastern |
| UG_2836 | 0.001411973 | 0.0101925 | 0.689975639 | 0.298419887 | 1.78E-15 | Northern |
| UG_2844 | 0.001444348 | 0.009480723 | 0.663301788 | 0.325773141 | 2.88E-15 | Northern |
| UG_2852 | 0.003354578 | 0.036834845 | 0.844152566 | 0.115658012 | 2.62E-14 | Northern |
| UG_2860 | 0.000462862 | 0.001736377 | 0.355846589 | 0.641954172 | 3.68E-16 | Northern |

| | | | | | | |
|---|---|---|---|---|---|---|
| UG_2868 | 0.003459363 | 0.110891524 | 0.867410506 | 0.018238608 | 2.82E-15 | Northern |
| UG_2876 | 0.00498135 | 0.119517731 | 0.852119912 | 0.023381007 | 3.58E-14 | Northern |
| UG_2884 | 0.003923484 | 0.096377547 | 0.87260446 | 0.027094509 | 9.19E-15 | Northern |
| UG_2892 | 0.012523032 | 0.662431648 | 0.324361718 | 0.000683603 | 3.76E-11 | Eastern |
| UG_2915 | 0.003694002 | 0.609872587 | 0.386165991 | 0.00026742 | 3.32E-15 | Eastern |
| UG_2987 | 0.02134425 | 0.657404181 | 0.32002036 | 0.001231207 | 1.88E-09 | Eastern |
| UG_2995 | 0.003551687 | 0.61560882 | 0.38059362 | 0.000245873 | 2.60E-15 | Eastern |
| UG_3003 | 0.010055244 | 0.665775948 | 0.323640739 | 0.000528069 | 7.72E-12 | Eastern |
| UG_2923 | 0.089631513 | 0.531223873 | 0.367914473 | 0.011178204 | 5.19E-05 | Eastern |
| UG_2931 | 0.003328683 | 0.600303792 | 0.396112527 | 0.000254999 | 1.46E-15 | Eastern |
| UG_2939 | 0.00291472 | 0.643702933 | 0.35322092 | 0.000161427 | 7.72E-16 | Eastern |
| UG_2947 | 0.009740444 | 0.696575095 | 0.293283337 | 0.000401124 | 8.34E-12 | Eastern |
| UG_2955 | 0.092136565 | 0.701192131 | 0.203568388 | 0.002708489 | 0.000394427 | Eastern |
| UG_2963 | 0.004429586 | 0.617696079 | 0.377565378 | 0.000308957 | 1.31E-14 | Eastern |
| UG_2971 | 0.010107484 | 0.684097852 | 0.305333935 | 0.00046073 | 9.60E-12 | Eastern |
| UG_2979 | 0.003559244 | 0.550241835 | 0.445811005 | 0.000387916 | 1.76E-15 | Eastern |
| UG_2916 | 0.015332799 | 0.628513564 | 0.355064578 | 0.001089059 | 1.25E-10 | Eastern |
| UG_2988 | 0.00375487 | 0.577130705 | 0.418772195 | 0.00034223 | 3.01E-15 | Eastern |
| UG_2996 | 0.032327796 | 0.624058961 | 0.341230557 | 0.002382653 | 3.23E-08 | Eastern |
| UG_3004 | 0.003615043 | 0.579175996 | 0.416885376 | 0.000323585 | 2.32E-15 | Eastern |
| UG_2924 | 0.009516698 | 0.66565988 | 0.324325053 | 0.000498368 | 5.15E-12 | Eastern |
| UG_2932 | 0.050163273 | 0.594722688 | 0.350681812 | 0.004431469 | 7.58E-07 | Eastern |
| UG_2940 | 0.008310716 | 0.516211211 | 0.474238654 | 0.001239419 | 7.05E-13 | Eastern |
| UG_2948 | 0.059861118 | 0.697521604 | 0.240397318 | 0.002210924 | 9.04E-06 | Eastern |
| UG_2956 | 0.079075739 | 0.688190557 | 0.229778336 | 0.002872215 | 8.32E-05 | Eastern |
| UG_2964 | 0.003643147 | 0.603960696 | 0.39212158 | 0.000274577 | 2.88E-15 | Eastern |
| UG_2972 | 0.014487093 | 0.652215485 | 0.332436781 | 0.000860641 | 1.00E-10 | Eastern |
| UG_2980 | 0.003453601 | 0.118240806 | 0.862191071 | 0.016114522 | 2.49E-15 | Eastern |
| UG_2917 | 0.003325551 | 0.586889627 | 0.409505094 | 0.000279728 | 1.33E-15 | Eastern |

| | | | | | | |
|---|---|---|---|---|---|---|
| UG_2989 | 0.009032653 | 0.603426563 | 0.386797457 | 0.000743328 | 2.12E-12 | Eastern |
| UG_2997 | 0.028788708 | 0.649522547 | 0.319931722 | 0.001757005 | 1.67E-08 | Eastern |
| UG_3005 | 0.011677285 | 0.580161336 | 0.407008219 | 0.00115316 | 1.19E-11 | Eastern |
| UG_2925 | 0.003956123 | 0.59621994 | 0.399506456 | 0.000317481 | 4.97E-15 | Eastern |
| UG_2933 | 0.014375331 | 0.668856243 | 0.316016581 | 0.000751845 | 1.11E-10 | Eastern |
| UG_2941 | 0.011099504 | 0.559493424 | 0.428146594 | 0.001260479 | 7.26E-12 | Eastern |
| UG_2949 | 0.003437544 | 0.581456271 | 0.414804892 | 0.000301293 | 1.63E-15 | Eastern |
| UG_2957 | 0.010099123 | 0.670280284 | 0.319108062 | 0.000512531 | 8.32E-12 | Eastern |
| UG_2965 | 0.003961331 | 0.610650692 | 0.385100736 | 0.000287241 | 5.54E-15 | Eastern |
| UG_2973 | 0.01066018 | 0.670758266 | 0.318040675 | 0.00054088 | 1.24E-11 | Eastern |
| UG_2981 | 0.132646705 | 0.59430604 | 0.262084053 | 0.008204309 | 0.002758894 | Eastern |
| UG_2918 | 0.003062573 | 0.618297306 | 0.378435318 | 0.000204804 | 9.07E-16 | Eastern |
| UG_2990 | 0.003228612 | 0.630727781 | 0.365845004 | 0.000198603 | 1.46E-15 | Eastern |
| UG_2998 | 0.002998607 | 0.590372266 | 0.406385616 | 0.000243512 | 6.42E-16 | Eastern |
| UG_3006 | 0.006191994 | 0.534037333 | 0.458974091 | 0.000796582 | 8.99E-14 | Eastern |
| UG_2926 | 0.008471338 | 0.641022904 | 0.349976823 | 0.000528934 | 1.77E-12 | Eastern |
| UG_2934 | 0.049839811 | 0.601258732 | 0.344703567 | 0.004197131 | 7.59E-07 | Eastern |
| UG_2942 | 0.015999935 | 0.650347187 | 0.332684993 | 0.000967884 | 2.06E-10 | Eastern |
| UG_2950 | 0.00673825 | 0.594582739 | 0.398103228 | 0.000575784 | 2.35E-13 | Eastern |
| UG_2958 | 0.003799923 | 0.570560555 | 0.425276693 | 0.000362829 | 3.16E-15 | Eastern |
| UG_2966 | 0.006533048 | 0.602637602 | 0.390303306 | 0.000526044 | 1.98E-13 | Eastern |
| UG_2974 | 0.002671545 | 0.607514455 | 0.38962395 | 0.000190049 | 3.12E-16 | Eastern |
| UG_2982 | 0.003054118 | 0.599527116 | 0.397185648 | 0.000233117 | 7.79E-16 | Eastern |
| UG_2919 | 0.003666987 | 0.618326144 | 0.377757057 | 0.000249813 | 3.34E-15 | Eastern |
| UG_2991 | 0.003284605 | 0.625610369 | 0.370895025 | 0.000210001 | 1.59E-15 | Eastern |
| UG_2999 | 0.005924741 | 0.675039647 | 0.318757478 | 0.000278134 | 1.76E-13 | Eastern |
| UG_3007 | 0.032459617 | 0.666496269 | 0.299320307 | 0.001723759 | 4.94E-08 | Eastern |
| UG_2927 | 0.007226204 | 0.631766214 | 0.360530994 | 0.000476588 | 5.13E-13 | Eastern |
| UG_2935 | 0.050750732 | 0.609526002 | 0.335716243 | 0.004006087 | 9.35E-07 | Eastern |

| | | | | | | |
|---|---|---|---|---|---|---|
| UG_2943 | 0.031447665 | 0.714432226 | 0.252999269 | 0.001120773 | 6.74E-08 | Eastern |
| UG_2951 | 0.02989579 | 0.678491869 | 0.290163644 | 0.001448668 | 2.98E-08 | Eastern |
| UG_2959 | 0.003354187 | 0.641440164 | 0.355013996 | 0.000191653 | 2.10E-15 | Eastern |
| UG_2967 | 0.013252727 | 0.687247087 | 0.298902259 | 0.000597926 | 7.31E-11 | Eastern |
| UG_2975 | 0.002991092 | 0.609603527 | 0.387193167 | 0.000212214 | 7.18E-16 | Eastern |
| UG_2983 | 0.007751699 | 0.594345952 | 0.397230706 | 0.000671644 | 6.52E-13 | Eastern |
| UG_2920 | 0.008686599 | 0.450661683 | 0.538620429 | 0.002031289 | 7.69E-13 | Eastern |
| UG_2992 | 0.003301072 | 0.585883875 | 0.410535656 | 0.000279397 | 1.25E-15 | Eastern |
| UG_3000 | 0.009395255 | 0.669170819 | 0.320955373 | 0.000478553 | 4.85E-12 | Eastern |
| UG_3008 | 0.002836539 | 0.595187099 | 0.401754932 | 0.00022143 | 4.43E-16 | Eastern |
| UG_2928 | 0.0033713 | 0.634451081 | 0.361974837 | 0.000202782 | 2.06E-15 | Eastern |
| UG_2936 | 0.008047707 | 0.522724797 | 0.468082435 | 0.001145062 | 5.74E-13 | Eastern |
| UG_2944 | 0.011154293 | 0.540984309 | 0.446422571 | 0.001438827 | 6.81E-12 | Eastern |
| UG_2952 | 0.003343594 | 0.627729789 | 0.368715685 | 0.000210932 | 1.84E-15 | Eastern |
| UG_2960 | 0.054684618 | 0.409801571 | 0.517864591 | 0.017648587 | 6.33E-07 | Eastern |
| UG_2968 | 0.003250994 | 0.635117456 | 0.361437678 | 0.000193872 | 1.59E-15 | Eastern |
| UG_2976 | 0.00357604 | 0.635337231 | 0.360871708 | 0.000215021 | 3.18E-15 | Eastern |
| UG_2984 | 0.003951541 | 0.586603731 | 0.409105666 | 0.000339061 | 4.63E-15 | Eastern |
| UG_2921 | 0.002863265 | 0.627171951 | 0.369786357 | 0.000178426 | 5.95E-16 | Eastern |
| UG_2993 | 0.003425155 | 0.614418581 | 0.381918034 | 0.00023823 | 1.98E-15 | Eastern |
| UG_3001 | 0.003385712 | 0.618652654 | 0.377733409 | 0.000228226 | 1.88E-15 | Eastern |
| UG_2929 | 0.007328775 | 0.622211586 | 0.369941107 | 0.000518531 | 5.28E-13 | Eastern |
| UG_2937 | 0.005966355 | 0.484250078 | 0.508711604 | 0.001071962 | 5.56E-14 | Eastern |
| UG_2945 | 0.056638323 | 0.647015787 | 0.293071132 | 0.003271591 | 3.17E-06 | Eastern |
| UG_2953 | 0.031510507 | 0.708833012 | 0.258477178 | 0.00117924 | 6.37E-08 | Eastern |
| UG_2961 | 0.076472726 | 0.650524631 | 0.269018398 | 0.003945585 | 3.87E-05 | Eastern |
| UG_2969 | 0.122510538 | 0.581212277 | 0.286135264 | 0.00900249 | 0.001139431 | Eastern |
| UG_2977 | 0.003444228 | 0.545733038 | 0.450436982 | 0.000385751 | 1.35E-15 | Eastern |
| UG_2985 | 0.016934185 | 0.576720494 | 0.404595744 | 0.001749576 | 1.80E-10 | Eastern |

| | | | | | | |
|---|---|---|---|---|---|---|
| UG_2922 | 0.007869013 | 0.588309135 | 0.403109695 | 0.000712157 | 7.00E-13 | Eastern |
| UG_2994 | 0.002834444 | 0.612272131 | 0.384697211 | 0.000196214 | 4.95E-16 | Eastern |
| UG_3002 | 0.010234215 | 0.650597531 | 0.338564807 | 0.000603447 | 7.66E-12 | Eastern |
| UG_2930 | 0.00362496 | 0.600519146 | 0.395576145 | 0.000279749 | 2.71E-15 | Eastern |
| UG_2938 | 0.003279127 | 0.614591575 | 0.381902537 | 0.000226761 | 1.45E-15 | Eastern |
| UG_2946 | 0.051468133 | 0.654386803 | 0.291290424 | 0.002853051 | 1.59E-06 | Eastern |
| UG_2954 | 0.012039177 | 0.606899754 | 0.380074477 | 0.000986592 | 1.78E-11 | Eastern |
| UG_2962 | 0.00366367 | 0.603616964 | 0.392442414 | 0.000276953 | 2.99E-15 | Eastern |
| UG_2970 | 0.133815866 | 0.606344788 | 0.249007248 | 0.007360514 | 0.003471583 | Eastern |
| UG_2978 | 0.003924652 | 0.634785061 | 0.361051109 | 0.000239179 | 6.21E-15 | Eastern |
| UG_2986 | 0.00455019 | 0.13912428 | 0.84048217 | 0.01584336 | 1.43E-14 | Eastern |
| UG_3009 | 0.003329589 | 0.580632221 | 0.415745673 | 0.000292517 | 1.29E-15 | Eastern |
| UG_3081 | 0.006078026 | 0.594480086 | 0.398927003 | 0.000514885 | 1.11E-13 | Eastern |
| UG_3089 | 5.99E-13 | 1.83E-13 | 1.89E-15 | 1.14E-16 | 1 | South_western |
| UG_3097 | 1.73E-12 | 5.88E-13 | 6.65E-15 | 3.74E-16 | 1 | South_western |
| UG_3017 | 0.010080338 | 0.667516387 | 0.321880798 | 0.000522476 | 7.99E-12 | Eastern |
| UG_3025 | 0.003376694 | 0.597714109 | 0.398645389 | 0.000263808 | 1.59E-15 | Eastern |
| UG_3033 | 0.011737087 | 0.638591487 | 0.348908169 | 0.000763257 | 1.89E-11 | Eastern |
| UG_3041 | 0.025857241 | 0.50260542 | 0.467023364 | 0.004513971 | 2.83E-09 | Eastern |
| UG_3049 | 0.006903033 | 0.600166002 | 0.392362574 | 0.00056839 | 2.91E-13 | Eastern |
| UG_3057 | 1.40E-12 | 4.41E-13 | 5.30E-15 | 3.37E-16 | 1 | Eastern |
| UG_3065 | 0.004894486 | 0.564463678 | 0.430141827 | 0.000500009 | 1.91E-14 | Eastern |
| UG_3073 | 0.008117311 | 0.542813284 | 0.448061321 | 0.001008084 | 6.74E-13 | Eastern |
| UG_3010 | 0.006717432 | 0.600472192 | 0.392259794 | 0.000550582 | 2.39E-13 | Eastern |
| UG_3082 | 0.003737012 | 0.12140872 | 0.858174052 | 0.016680216 | 4.24E-15 | Eastern |
| UG_3090 | 8.14E-13 | 2.54E-13 | 2.73E-15 | 1.63E-16 | 1 | South_western |
| UG_3098 | 4.50E-13 | 1.34E-13 | 1.34E-15 | 8.05E-17 | 1 | South_western |
| UG_3018 | 0.011215195 | 0.536462194 | 0.450829916 | 0.001492695 | 6.92E-12 | Eastern |
| UG_3026 | 0.002943211 | 0.580078801 | 0.416721801 | 0.000256186 | 5.26E-16 | Eastern |

| | | | | | | |
|---|---|---|---|---|---|---|
| UG_3034 | 0.00983475 | 0.651176051 | 0.338413377 | 0.000575822 | 5.75E-12 | Eastern |
| UG_3042 | 0.00319975 | 0.591831134 | 0.404710118 | 0.000258998 | 1.04E-15 | Eastern |
| UG_3050 | 0.010524271 | 0.634323799 | 0.354450691 | 0.000701239 | 8.20E-12 | Eastern |
| UG_3058 | 0.007241416 | 0.643870037 | 0.348451355 | 0.000437192 | 5.75E-13 | Eastern |
| UG_3066 | 0.007433925 | 0.607780686 | 0.384201612 | 0.000583778 | 5.27E-13 | Eastern |
| UG_3074 | 0.010001533 | 0.628862602 | 0.360444813 | 0.000691052 | 5.40E-12 | Eastern |
| UG_3011 | 0.014187972 | 0.661943253 | 0.323086654 | 0.000782122 | 9.41E-11 | Eastern |
| UG_3083 | 8.55E-12 | 3.57E-12 | 4.39E-14 | 2.08E-15 | 1 | South_western |
| UG_3091 | 3.79E-13 | 1.08E-13 | 1.10E-15 | 7.14E-17 | 1 | South_western |
| UG_3099 | 1.96E-11 | 8.62E-12 | 1.20E-13 | 5.74E-15 | 1 | South_western |
| UG_3019 | 0.010187868 | 0.669911954 | 0.319381382 | 0.000518796 | 8.84E-12 | Eastern |
| UG_3027 | 0.009373857 | 0.667283035 | 0.322858775 | 0.000484334 | 4.68E-12 | Eastern |
| UG_3035 | 0.015552173 | 0.668256678 | 0.315371504 | 0.000819644 | 1.97E-10 | Eastern |
| UG_3043 | 0.003509268 | 0.507378782 | 0.488601214 | 0.000510736 | 1.30E-15 | Eastern |
| UG_3051 | 0.007346151 | 0.561764293 | 0.430095251 | 0.000794305 | 3.61E-13 | Eastern |
| UG_3059 | 0.00625306 | 0.642929113 | 0.350442461 | 0.000375367 | 1.96E-13 | Eastern |
| UG_3067 | 0.002842719 | 0.600898506 | 0.396045506 | 0.00021327 | 4.68E-16 | Eastern |
| UG_3075 | 0.008226133 | 0.539431608 | 0.451295615 | 0.001046644 | 7.30E-13 | Eastern |
| UG_3012 | 0.005640889 | 0.624882088 | 0.369094531 | 0.000382492 | 8.01E-14 | Eastern |
| UG_3084 | 5.09E-13 | 1.48E-13 | 1.58E-15 | 1.02E-16 | 1 | South_western |
| UG_3092 | 1.30E-12 | 4.00E-13 | 4.92E-15 | 3.27E-16 | 1 | South_western |
| UG_3100 | 3.55E-10 | 1.93E-10 | 3.96E-12 | 1.92E-13 | 0.999999999 | South_western |
| UG_3020 | 0.004548082 | 0.499419974 | 0.495314276 | 0.000717668 | 8.21E-15 | Eastern |
| UG_3028 | 0.008986441 | 0.597316176 | 0.392925553 | 0.000771831 | 1.96E-12 | Eastern |
| UG_3036 | 0.00318199 | 0.618991898 | 0.377613519 | 0.000212593 | 1.20E-15 | Eastern |
| UG_3044 | 0.006281901 | 0.620069919 | 0.373202883 | 0.000445297 | 1.69E-13 | Eastern |
| UG_3052 | 0.00585548 | 0.600440654 | 0.393229807 | 0.000474059 | 8.81E-14 | Eastern |
| UG_3060 | 0.007205585 | 0.554883496 | 0.437095541 | 0.000815377 | 3.02E-13 | Eastern |
| UG_3068 | 0.007520045 | 0.621665697 | 0.370278937 | 0.00053532 | 6.35E-13 | Eastern |

| | | | | | | |
|---|---|---|---|---|---|---|
| UG_3076 | 0.002815365 | 0.592636165 | 0.404324927 | 0.000223543 | 4.13E-16 | Eastern |
| UG_3013 | 0.003444574 | 0.611401765 | 0.384908758 | 0.000244903 | 2.02E-15 | Eastern |
| UG_3085 | 2.71E-08 | 1.92E-08 | 7.59E-10 | 4.10E-11 | 0.999999953 | South_western |
| UG_3093 | 8.26E-13 | 2.67E-13 | 2.73E-15 | 1.53E-16 | 1 | South_western |
| UG_3101 | 1.78E-10 | 8.55E-11 | 1.78E-12 | 9.97E-14 | 1 | South_western |
| UG_3021 | 0.008484041 | 0.686078992 | 0.305060711 | 0.000376255 | 2.71E-12 | Eastern |
| UG_3029 | 0.003172427 | 0.610260076 | 0.386342063 | 0.000225435 | 1.10E-15 | Eastern |
| UG_3037 | 0.003172713 | 0.622214923 | 0.374405275 | 0.000207089 | 1.21E-15 | Eastern |
| UG_3045 | 0.004861157 | 0.594983435 | 0.399753707 | 0.000401701 | 2.20E-14 | Eastern |
| UG_3053 | 0.015226986 | 0.671040459 | 0.312947785 | 0.000784769 | 1.73E-10 | Eastern |
| UG_3061 | 0.009227487 | 0.505892354 | 0.48339102 | 0.001489139 | 1.45E-12 | Eastern |
| UG_3069 | 0.012141761 | 0.495935023 | 0.489781501 | 0.002141714 | 1.03E-11 | Eastern |
| UG_3077 | 0.002745162 | 0.617359148 | 0.379713019 | 0.000182671 | 4.08E-16 | Eastern |
| UG_3014 | 0.010119961 | 0.670033537 | 0.319331869 | 0.000514633 | 8.43E-12 | Eastern |
| UG_3086 | 8.19E-11 | 4.17E-11 | 6.58E-13 | 2.90E-14 | 1 | South_western |
| UG_3094 | 3.58E-13 | 1.01E-13 | 1.04E-15 | 6.76E-17 | 1 | South_western |
| UG_3102 | 1.60E-12 | 5.43E-13 | 6.09E-15 | 3.43E-16 | 1 | South_western |
| UG_3022 | 0.013949718 | 0.623804514 | 0.361224944 | 0.001020825 | 5.99E-11 | Eastern |
| UG_3030 | 0.038676843 | 0.659615611 | 0.299568671 | 0.002138697 | 1.78E-07 | Eastern |
| UG_3038 | 0.002883043 | 0.617198074 | 0.37972581 | 0.000193073 | 5.81E-16 | Eastern |
| UG_3046 | 0.007036657 | 0.596063385 | 0.396302596 | 0.000597362 | 3.26E-13 | Eastern |
| UG_3054 | 0.006003398 | 0.571689042 | 0.421712399 | 0.000595161 | 8.79E-14 | Eastern |
| UG_3062 | 0.008443627 | 0.63976316 | 0.351261237 | 0.000531975 | 1.71E-12 | Eastern |
| UG_3070 | 0.00570925 | 0.603603924 | 0.390235858 | 0.000450967 | 7.49E-14 | Eastern |
| UG_3078 | 0.003058198 | 0.620085848 | 0.376654064 | 0.00020189 | 9.09E-16 | Eastern |
| UG_3015 | 0.02138983 | 0.715377311 | 0.26246183 | 0.000771026 | 3.60E-09 | Eastern |
| UG_3087 | 8.34E-13 | 2.44E-13 | 2.91E-15 | 2.01E-16 | 1 | South_western |
| UG_3095 | 1.33E-12 | 4.46E-13 | 4.83E-15 | 2.68E-16 | 1 | South_western |
| UG_3023 | 0.011597707 | 0.64263243 | 0.345038403 | 0.000731461 | 1.79E-11 | Eastern |

| | | | | | | |
|---|---|---|---|---|---|---|
| UG_3031 | 0.033146292 | 0.672244571 | 0.292930717 | 0.001678358 | 6.16E-08 | Eastern |
| UG_3039 | 0.002720942 | 0.630794856 | 0.366319901 | 0.0001643 | 4.23E-16 | Eastern |
| UG_3047 | 0.008546637 | 0.652726092 | 0.338237655 | 0.000489617 | 2.08E-12 | Eastern |
| UG_3055 | 0.002909466 | 0.589745702 | 0.407108291 | 0.000236542 | 5.14E-16 | Eastern |
| UG_3063 | 0.005697956 | 0.596010367 | 0.397817008 | 0.000474669 | 7.01E-14 | Eastern |
| UG_3071 | 0.007463132 | 0.630572502 | 0.361466518 | 0.000497848 | 6.44E-13 | Eastern |
| UG_3079 | 0.004765088 | 0.607128544 | 0.387745538 | 0.000360829 | 2.06E-14 | Eastern |
| UG_3016 | 0.008509525 | 0.494056366 | 0.495956174 | 0.001477935 | 7.64E-13 | Eastern |
| UG_3088 | 1.94E-12 | 6.66E-13 | 7.66E-15 | 4.31E-16 | 1 | South_western |
| UG_3096 | 5.86E-13 | 1.76E-13 | 1.85E-15 | 1.15E-16 | 1 | South_western |
| UG_3024 | 0.011502546 | 0.682237965 | 0.305723346 | 0.000536143 | 2.44E-11 | Eastern |
| UG_3032 | 0.003560967 | 0.572153557 | 0.423951436 | 0.00033404 | 1.99E-15 | Eastern |
| UG_3040 | 0.035102913 | 0.69141994 | 0.271967167 | 0.001509862 | 1.19E-07 | Eastern |
| UG_3048 | 0.00528477 | 0.592813749 | 0.401454485 | 0.000446995 | 3.97E-14 | Eastern |
| UG_3056 | 0.002827504 | 0.574516234 | 0.422401616 | 0.000254646 | 3.80E-16 | Eastern |
| UG_3064 | 0.004901577 | 0.611937985 | 0.382800711 | 0.000359727 | 2.62E-14 | Eastern |
| UG_3072 | 0.005338324 | 0.505559002 | 0.488281487 | 0.000821187 | 2.69E-14 | Eastern |
| UG_3080 | 0.003603024 | 0.084051946 | 0.880595569 | 0.031749462 | 6.41E-15 | Eastern |
| UG_3103 | 2.51E-12 | 8.69E-13 | 1.05E-14 | 6.08E-16 | 1 | South_western |
| UG_3175 | 2.71E-07 | 2.34E-07 | 1.21E-08 | 6.22E-10 | 0.999999482 | South_western |
| UG_3183 | 1.74E-06 | 1.69E-06 | 1.14E-07 | 6.11E-09 | 0.999996456 | South_western |
| UG_3191 | 1.01E-12 | 3.20E-13 | 3.52E-15 | 2.11E-16 | 1 | South_western |
| UG_3111 | 2.30E-12 | 7.62E-13 | 9.67E-15 | 6.03E-16 | 1 | South_western |
| UG_3119 | 6.55E-13 | 1.96E-13 | 2.13E-15 | 1.35E-16 | 1 | South_western |
| UG_3127 | 9.83E-13 | 3.32E-13 | 3.31E-15 | 1.74E-16 | 1 | South_western |
| UG_3135 | 3.74E-12 | 1.29E-12 | 1.74E-14 | 1.08E-15 | 1 | South_western |
| UG_3143 | 0.000711402 | 0.002206833 | 0.345601608 | 0.651480157 | 5.53E-15 | South_western |
| UG_3151 | 1.95E-12 | 6.62E-13 | 7.79E-15 | 4.52E-16 | 1 | South_western |
| UG_3159 | 1.61E-12 | 5.20E-13 | 6.29E-15 | 3.92E-16 | 1 | South_western |

| | | | | | | |
|---|---|---|---|---|---|---|
| UG_3167 | 0.02560524 | 0.531217905 | 0.439505299 | 0.003671553 | 3.01E-09 | South_western |
| UG_3104 | 2.51E-12 | 8.02E-13 | 1.10E-14 | 7.49E-16 | 1 | South_western |
| UG_3176 | 6.19E-12 | 2.32E-12 | 3.10E-14 | 1.74E-15 | 1 | South_western |
| UG_3184 | 7.56E-12 | 2.78E-12 | 4.02E-14 | 2.41E-15 | 1 | South_western |
| UG_3192 | 4.04E-13 | 1.15E-13 | 1.20E-15 | 7.80E-17 | 1 | South_western |
| UG_3112 | 1.42E-12 | 4.55E-13 | 5.37E-15 | 3.32E-16 | 1 | South_western |
| UG_3120 | 2.60E-12 | 9.41E-13 | 1.07E-14 | 5.65E-16 | 1 | South_western |
| UG_3128 | 1.26E-12 | 4.20E-13 | 4.53E-15 | 2.52E-16 | 1 | South_western |
| UG_3136 | 7.46E-13 | 2.23E-13 | 2.50E-15 | 1.61E-16 | 1 | South_western |
| UG_3144 | 3.92E-07 | 5.02E-07 | 1.55E-08 | 3.72E-10 | 0.999999091 | South_western |
| UG_3152 | 0.001429938 | 0.007761241 | 0.601294847 | 0.389513974 | 6.11E-15 | South_western |
| UG_3160 | 1.11E-12 | 3.49E-13 | 3.99E-15 | 2.47E-16 | 1 | South_western |
| UG_3168 | 2.06E-08 | 1.76E-08 | 4.90E-10 | 1.73E-11 | 0.999999961 | South_western |
| UG_3105 | 1.02E-12 | 3.17E-13 | 3.60E-15 | 2.23E-16 | 1 | South_western |
| UG_3177 | 6.38E-13 | 2.00E-13 | 2.01E-15 | 1.15E-16 | 1 | South_western |
| UG_3185 | 0.106770444 | 0.569037086 | 0.059868129 | 0.000627649 | 0.263696693 | South_western |
| UG_3193 | 8.41E-13 | 2.52E-13 | 2.91E-15 | 1.91E-16 | 1 | South_western |
| UG_3113 | 3.77E-12 | 1.43E-12 | 1.66E-14 | 8.36E-16 | 1 | South_western |
| UG_3121 | 4.52E-12 | 1.62E-12 | 2.14E-14 | 1.26E-15 | 1 | South_western |
| UG_3129 | 2.67E-07 | 2.24E-07 | 1.20E-08 | 6.46E-10 | 0.999999496 | South_western |
| UG_3137 | 0.000917162 | 0.005083228 | 0.556489303 | 0.437510307 | 8.39E-16 | South_western |
| UG_3145 | 2.28E-12 | 7.69E-13 | 9.49E-15 | 5.71E-16 | 1 | South_western |
| UG_3153 | 0.002770315 | 0.090750334 | 0.88559784 | 0.020881511 | 8.00E-16 | South_western |
| UG_3161 | 0.000534141 | 0.002240214 | 0.403874607 | 0.593351037 | 3.59E-16 | South_western |
| UG_3169 | 1.54E-13 | 4.01E-14 | 3.79E-16 | 2.58E-17 | 1 | South_western |
| UG_3106 | 5.22E-13 | 1.58E-13 | 1.60E-15 | 9.50E-17 | 1 | South_western |
| UG_3178 | 6.60E-13 | 2.05E-13 | 2.11E-15 | 1.24E-16 | 1 | South_western |
| UG_3186 | 5.37E-13 | 1.58E-13 | 1.68E-15 | 1.07E-16 | 1 | South_western |
| UG_3194 | 3.04E-12 | 1.05E-12 | 1.34E-14 | 7.93E-16 | 1 | South_western |

| | | | | | | |
|---|---|---|---|---|---|---|
| UG_3114 | 0.09095437 | 0.52586396 | 0.371380784 | 0.011744126 | 5.68E-05 | South_western |
| UG_3122 | 2.53E-12 | 8.95E-13 | 1.05E-14 | 5.83E-16 | 1 | South_western |
| UG_3130 | 2.91E-12 | 1.06E-12 | 1.23E-14 | 6.49E-16 | 1 | South_western |
| UG_3138 | 2.52E-12 | 8.56E-13 | 1.06E-14 | 6.37E-16 | 1 | South_western |
| UG_3146 | 5.93E-13 | 1.78E-13 | 1.88E-15 | 1.17E-16 | 1 | South_western |
| UG_3154 | 0.151233252 | 0.606802193 | 0.221490095 | 0.006967053 | 0.013507407 | South_western |
| UG_3162 | 0.126195565 | 0.593315754 | 0.270595254 | 0.008197048 | 0.001696378 | South_western |
| UG_3170 | 3.22E-13 | 9.64E-14 | 8.78E-16 | 4.97E-17 | 1 | South_western |
| UG_3107 | 7.06E-12 | 2.95E-12 | 3.45E-14 | 1.57E-15 | 1 | South_western |
| UG_3179 | 9.56E-11 | 4.87E-11 | 7.99E-13 | 3.61E-14 | 1 | South_western |
| UG_3187 | 1.16E-12 | 3.80E-13 | 4.15E-15 | 2.38E-16 | 1 | South_western |
| UG_3195 | 3.03E-12 | 1.02E-12 | 1.35E-14 | 8.54E-16 | 1 | South_western |
| UG_3115 | 1.23E-12 | 4.12E-13 | 4.39E-15 | 2.42E-16 | 1 | South_western |
| UG_3123 | 7.51E-13 | 2.44E-13 | 2.42E-15 | 1.32E-16 | 1 | South_western |
| UG_3131 | 1.74E-12 | 5.90E-13 | 6.72E-15 | 3.80E-16 | 1 | South_western |
| UG_3139 | 8.02E-13 | 2.46E-13 | 2.70E-15 | 1.68E-16 | 1 | South_western |
| UG_3147 | 1.01E-07 | 8.41E-08 | 3.60E-09 | 1.71E-10 | 0.999999811 | South_western |
| UG_3155 | 1.04E-12 | 3.21E-13 | 3.74E-15 | 2.40E-16 | 1 | South_western |
| UG_3163 | 4.75E-12 | 1.78E-12 | 2.24E-14 | 1.21E-15 | 1 | South_western |
| UG_3171 | 6.81E-13 | 2.12E-13 | 2.19E-15 | 1.28E-16 | 1 | South_western |
| UG_3108 | 1.43E-12 | 4.65E-13 | 5.42E-15 | 3.29E-16 | 1 | South_western |
| UG_3180 | 6.91E-13 | 2.07E-13 | 2.28E-15 | 1.46E-16 | 1 | South_western |
| UG_3188 | 1.93E-12 | 6.07E-13 | 7.99E-15 | 5.40E-16 | 1 | South_western |
| UG_3196 | 6.89E-13 | 2.15E-13 | 2.22E-15 | 1.31E-16 | 1 | South_western |
| UG_3116 | 1.96E-08 | 1.38E-08 | 5.11E-10 | 2.68E-11 | 0.999999966 | South_western |
| UG_3124 | 2.69E-12 | 8.88E-13 | 1.17E-14 | 7.52E-16 | 1 | South_western |
| UG_3132 | 3.79E-12 | 1.31E-12 | 1.76E-14 | 1.08E-15 | 1 | South_western |
| UG_3140 | 1.03E-12 | 3.22E-13 | 3.63E-15 | 2.23E-16 | 1 | South_western |
| UG_3148 | 3.73E-12 | 1.27E-12 | 1.74E-14 | 1.10E-15 | 1 | South_western |

| | | | | | | |
|---|---|---|---|---|---|---|
| UG_3156 | 1.86E-12 | 6.18E-13 | 7.41E-15 | 4.44E-16 | 1 | South_western |
| UG_3164 | 6.36E-08 | 7.37E-08 | 1.70E-09 | 3.80E-11 | 0.999999861 | South_western |
| UG_3172 | 2.78E-08 | 2.01E-08 | 7.77E-10 | 4.08E-11 | 0.999999951 | South_western |
| UG_3109 | 8.99E-11 | 4.32E-11 | 7.62E-13 | 3.85E-14 | 1 | South_western |
| UG_3181 | 4.89E-12 | 1.83E-12 | 2.31E-14 | 1.25E-15 | 1 | South_western |
| UG_3189 | 8.59E-13 | 2.58E-13 | 2.98E-15 | 1.95E-16 | 1 | South_western |
| UG_3117 | 1.32E-10 | 6.86E-11 | 1.18E-12 | 5.41E-14 | 1 | South_western |
| UG_3125 | 1.89E-12 | 6.14E-13 | 7.64E-15 | 4.81E-16 | 1 | South_western |
| UG_3133 | 3.47E-07 | 3.65E-07 | 1.48E-08 | 5.27E-10 | 0.999999272 | South_western |
| UG_3141 | 6.52E-13 | 1.98E-13 | 2.10E-15 | 1.29E-16 | 1 | South_western |
| UG_3149 | 0.113994251 | 0.589817611 | 0.287390565 | 0.008152118 | 0.000645454 | South_western |
| UG_3157 | 0.003824325 | 0.112119096 | 0.864161157 | 0.019895422 | 5.77E-15 | South_western |
| UG_3165 | 1.81E-11 | 6.93E-12 | 1.17E-13 | 7.42E-15 | 1 | South_western |
| UG_3173 | 1.27E-10 | 6.38E-11 | 1.15E-12 | 5.60E-14 | 1 | South_western |
| UG_3110 | 2.63E-12 | 9.04E-13 | 1.12E-14 | 6.54E-16 | 1 | South_western |
| UG_3182 | 6.74E-13 | 1.90E-13 | 2.27E-15 | 1.63E-16 | 1 | South_western |
| UG_3190 | 2.99E-12 | 9.45E-13 | 1.37E-14 | 9.80E-16 | 1 | South_western |
| UG_3118 | 3.82E-12 | 1.37E-12 | 1.74E-14 | 9.88E-16 | 1 | South_western |
| UG_3126 | 7.80E-13 | 2.39E-13 | 2.62E-15 | 1.62E-16 | 1 | South_western |
| UG_3134 | 0.000351711 | 0.000563647 | 6.56E-05 | 2.81E-06 | 0.999016251 | South_western |
| UG_3142 | 4.93E-05 | 7.35E-05 | 5.90E-06 | 2.17E-07 | 0.999871057 | South_western |
| UG_3150 | 5.44E-12 | 1.99E-12 | 2.68E-14 | 1.55E-15 | 1 | South_western |
| UG_3158 | 4.62E-13 | 1.35E-13 | 1.39E-15 | 8.70E-17 | 1 | South_western |
| UG_3166 | 2.53E-12 | 8.69E-13 | 1.07E-14 | 6.29E-16 | 1 | South_western |
| UG_3174 | 4.37E-12 | 1.45E-12 | 2.14E-14 | 1.46E-15 | 1 | South_western |
| UG_3197 | 4.80E-12 | 1.76E-12 | 2.29E-14 | 1.30E-15 | 1 | South_western |
| UG_3269 | 0.006137164 | 0.460544776 | 0.532018903 | 0.001299157 | 6.31E-14 | Eastern |
| UG_3277 | 0.005922842 | 0.469176445 | 0.523722631 | 0.001178082 | 5.01E-14 | Northern |
| UG_3285 | 0.003238232 | 0.55409698 | 0.442324436 | 0.000340353 | 9.04E-16 | Northern |

| | | | | | | |
|---|---|---|---|---|---|---|
| UG_3205 | 8.65E-12 | 3.27E-12 | 4.68E-14 | 2.71E-15 | 1 | South_western |
| UG_3213 | 0.003655077 | 0.018914305 | 0.686195769 | 0.291234849 | 4.66E-13 | Northern |
| UG_3221 | 0.011275714 | 0.516966245 | 0.470043361 | 0.00171468 | 6.57E-12 | Northern |
| UG_3229 | 9.57E-13 | 3.08E-13 | 3.29E-15 | 1.90E-16 | 1 | South_western |
| UG_3237 | 0.005041812 | 0.040068432 | 0.808129866 | 0.14675989 | 4.94E-13 | Northern |
| UG_3245 | 0.002259831 | 0.019132889 | 0.775340333 | 0.203266946 | 8.64E-15 | Northern |
| UG_3253 | 0.002726663 | 0.014488487 | 0.665111766 | 0.317673085 | 1.13E-13 | Central |
| UG_3261 | 0.079098121 | 0.649589881 | 0.267198428 | 0.004062721 | 5.08E-05 | Central |
| UG_3198 | 9.70E-13 | 2.85E-13 | 3.50E-15 | 2.44E-16 | 1 | South_western |
| UG_3270 | 0.090777354 | 0.645416699 | 0.25908902 | 0.004559181 | 0.000157747 | Eastern |
| UG_3278 | 0.006087441 | 0.301808367 | 0.688009869 | 0.004094324 | 5.10E-14 | Northern |
| UG_3286 | 6.83E-12 | 2.45E-12 | 3.59E-14 | 2.25E-15 | 1 | South_western |
| UG_3206 | 1.38E-12 | 4.28E-13 | 5.27E-15 | 3.46E-16 | 1 | South_western |
| UG_3214 | 0.003308701 | 0.033898415 | 0.833744292 | 0.129048592 | 2.98E-14 | Northern |
| UG_3222 | 6.71E-13 | 1.99E-13 | 2.21E-15 | 1.43E-16 | 1 | South_western |
| UG_3230 | 2.48E-12 | 8.51E-13 | 1.04E-14 | 6.12E-16 | 1 | South_western |
| UG_3238 | 0.003226792 | 0.629125411 | 0.367447013 | 0.000200785 | 1.44E-15 | Northern |
| UG_3246 | 0.00088334 | 0.006096479 | 0.624572978 | 0.368447203 | 2.72E-16 | Northern |
| UG_3254 | 0.113944997 | 0.583797808 | 0.293085458 | 0.008564785 | 0.000606952 | Central |
| UG_3262 | 0.004117475 | 0.143347607 | 0.839125146 | 0.013409772 | 6.58E-15 | Central |
| UG_3199 | 1.16E-12 | 3.54E-13 | 4.32E-15 | 2.90E-16 | 1 | South_western |
| UG_3271 | 0.01160799 | 0.629842093 | 0.357745567 | 0.000804349 | 1.62E-11 | Eastern |
| UG_3279 | 0.011205071 | 0.097190369 | 0.814350081 | 0.077254479 | 2.31E-11 | Northern |
| UG_3287 | 2.92E-12 | 9.32E-13 | 1.32E-14 | 9.22E-16 | 1 | South_western |
| UG_3207 | 3.34E-12 | 1.08E-12 | 1.56E-14 | 1.08E-15 | 1 | South_western |
| UG_3215 | 0.086378246 | 0.665597669 | 0.244181611 | 0.003710878 | 0.000131597 | Northern |
| UG_3223 | 1.33E-12 | 4.39E-13 | 4.87E-15 | 2.80E-16 | 1 | South_western |
| UG_3231 | 1.55E-12 | 4.96E-13 | 6.04E-15 | 3.82E-16 | 1 | South_western |
| UG_3239 | 0.003672064 | 0.119064574 | 0.860273137 | 0.016990224 | 3.86E-15 | Northern |

| | | | | | | |
|---|---|---|---|---|---|---|
| UG_3247 | 0.131530788 | 0.632121364 | 0.226458901 | 0.005752279 | 0.004136667 | Central |
| UG_3255 | 0.004817216 | 0.129396428 | 0.846397928 | 0.019388428 | 2.45E-14 | Central |
| UG_3263 | 0.007449212 | 0.643532145 | 0.348566722 | 0.00045192 | 7.05E-13 | Central |
| UG_3200 | 1.07E-10 | 4.91E-11 | 9.62E-13 | 5.45E-14 | 1 | South_western |
| UG_3272 | 0.097070564 | 0.596845199 | 0.298808751 | 0.007106905 | 0.000168581 | Eastern |
| UG_3280 | 0.017149658 | 0.186832275 | 0.760491439 | 0.035526628 | 1.64E-10 | Northern |
| UG_3288 | 1.88E-12 | 6.18E-13 | 7.55E-15 | 4.65E-16 | 1 | South_western |
| UG_3208 | 3.72E-12 | 1.33E-12 | 1.69E-14 | 9.70E-16 | 1 | South_western |
| UG_3216 | 0.002837345 | 0.04955681 | 0.885847577 | 0.061758267 | 3.43E-15 | Northern |
| UG_3224 | 4.87E-12 | 1.69E-12 | 2.39E-14 | 1.52E-15 | 1 | South_western |
| UG_3232 | 2.71E-12 | 9.49E-13 | 1.15E-14 | 6.52E-16 | 1 | South_western |
| UG_3240 | 0.000123066 | 0.000200651 | 0.106884044 | 0.89279224 | 3.49E-16 | Northern |
| UG_3248 | 0.000744379 | 0.003279745 | 0.456487303 | 0.539488573 | 1.06E-15 | Central |
| UG_3256 | 0.125485178 | 0.626548273 | 0.239506616 | 0.006071284 | 0.002388649 | Central |
| UG_3264 | 0.002034059 | 0.020419305 | 0.805842016 | 0.171704621 | 3.01E-15 | Eastern |
| UG_3201 | 2.59E-11 | 1.12E-11 | 1.72E-13 | 8.94E-15 | 1 | South_western |
| UG_3273 | 0.004487876 | 0.097361369 | 0.867504244 | 0.030646511 | 2.43E-14 | Central |
| UG_3281 | 0.010564041 | 0.589762068 | 0.398705283 | 0.000968607 | 6.08E-12 | Northern |
| UG_3289 | 4.28E-12 | 1.50E-12 | 2.03E-14 | 1.25E-15 | 1 | South_western |
| UG_3209 | 1.18E-10 | 5.27E-11 | 1.11E-12 | 6.87E-14 | 1 | South_western |
| UG_3217 | 0.028107293 | 0.352976593 | 0.604913763 | 0.014002347 | 3.80E-09 | Northern |
| UG_3225 | 3.54E-12 | 1.20E-12 | 1.63E-14 | 1.03E-15 | 1 | South_western |
| UG_3233 | 0.003302272 | 0.584286115 | 0.412128989 | 0.000282624 | 1.24E-15 | Northern |
| UG_3241 | 0.001374505 | 0.008631444 | 0.643962315 | 0.346031736 | 2.77E-15 | Northern |
| UG_3249 | 0.141934218 | 0.571970952 | 0.271854367 | 0.010043653 | 0.00419681 | Central |
| UG_3257 | 0.004369568 | 0.158202614 | 0.825677014 | 0.011750804 | 8.75E-15 | Central |
| UG_3265 | 0.003327855 | 0.113179094 | 0.866664643 | 0.016828409 | 2.05E-15 | Eastern |
| UG_3202 | 2.72E-12 | 8.67E-13 | 1.21E-14 | 8.41E-16 | 1 | South_western |
| UG_3274 | 0.008219084 | 0.466816841 | 0.523251382 | 0.001712693 | 5.39E-13 | Central |

| | | | | | | |
|---|---|---|---|---|---|---|
| UG_3290 | 1.93E-12 | 6.01E-13 | 7.98E-15 | 5.47E-16 | 1 | South_western |
| UG_3210 | 8.40E-13 | 2.55E-13 | 2.88E-15 | 1.84E-16 | 1 | South_western |
| UG_3218 | 0.006942425 | 0.341182104 | 0.648399206 | 0.003476264 | 1.29E-13 | Northern |
| UG_3226 | 9.23E-13 | 2.86E-13 | 3.21E-15 | 2.00E-16 | 1 | South_western |
| UG_3234 | 0.0044841 | 0.090301261 | 0.870121105 | 0.035093533 | 2.80E-14 | Northern |
| UG_3242 | 0.004506862 | 0.108244675 | 0.861928539 | 0.025319925 | 2.05E-14 | Northern |
| UG_3250 | 0.003159661 | 0.582132348 | 0.414434785 | 0.000273206 | 8.90E-16 | Central |
| UG_3258 | 0.00613207 | 0.636973328 | 0.356510758 | 0.000383844 | 1.62E-13 | Central |
| UG_3266 | 3.52E-12 | 1.19E-12 | 1.62E-14 | 1.03E-15 | 1 | Eastern |
| UG_3203 | 1.44E-12 | 4.59E-13 | 5.50E-15 | 3.46E-16 | 1 | South_western |
| UG_3275 | 0.004379724 | 0.116084182 | 0.857995665 | 0.021540428 | 1.47E-14 | Central |
| UG_3283 | 0.144197122 | 0.582678194 | 0.258449925 | 0.00911514 | 0.005559619 | Central |
| UG_3211 | 1.01E-12 | 3.28E-13 | 3.52E-15 | 2.02E-16 | 1 | South_western |
| UG_3219 | 0.003136057 | 0.038364021 | 0.856259682 | 0.10224024 | 1.40E-14 | Northern |
| UG_3227 | 1.63E-12 | 5.49E-13 | 6.20E-15 | 3.52E-16 | 1 | South_western |
| UG_3235 | 0.004879969 | 0.137765534 | 0.83994192 | 0.017412577 | 2.42E-14 | Northern |
| UG_3243 | 0.004284229 | 0.038765605 | 0.823373728 | 0.133576438 | 1.52E-13 | Northern |
| UG_3251 | 0.007162249 | 0.129831583 | 0.833686717 | 0.02931945 | 4.50E-13 | Central |
| UG_3259 | 0.121455332 | 0.573597342 | 0.294410476 | 0.009556945 | 0.000979906 | Central |
| UG_3267 | 0.030815876 | 0.462475126 | 0.499618041 | 0.007090949 | 9.09E-09 | Eastern |
| UG_3204 | 6.50E-13 | 1.89E-13 | 2.14E-15 | 1.44E-16 | 1 | South_western |
| UG_3276 | 0.002799654 | 0.607202794 | 0.389796945 | 0.000200606 | 4.37E-16 | Northern |
| UG_3284 | 0.013477706 | 0.569983303 | 0.415097024 | 0.001441968 | 3.21E-11 | Northern |
| UG_3212 | 0.007425637 | 0.122863933 | 0.835943848 | 0.033766582 | 6.50E-13 | Northern |
| UG_3220 | 0.002406659 | 0.025078897 | 0.822787384 | 0.14972706 | 6.01E-15 | Northern |
| UG_3228 | 1.18E-12 | 3.76E-13 | 4.26E-15 | 2.57E-16 | 1 | South_western |
| UG_3236 | 0.002452188 | 0.034081097 | 0.866192337 | 0.097274377 | 2.91E-15 | Northern |
| UG_3244 | 0.004862019 | 0.112886465 | 0.856880975 | 0.025370541 | 3.32E-14 | Northern |
| UG_3252 | 0.004447663 | 0.140381127 | 0.839982512 | 0.015188698 | 1.19E-14 | Central |

| | | | | | | |
|---|---|---|---|---|---|---|
| UG_3260 | 0.003716813 | 0.113510692 | 0.863921647 | 0.018850847 | 4.58E-15 | Central |
| UG_3268 | 0.019394465 | 0.643008583 | 0.336350326 | 0.001246625 | 8.08E-10 | Eastern |
| UG_3291 | 1.45E-13 | 3.65E-14 | 3.57E-16 | 2.57E-17 | 1 | South_western |
| UG_3299 | 0.003898006 | 0.575801697 | 0.419940359 | 0.000359938 | 3.92E-15 | Eastern |
| UG_3307 | 0.004775848 | 0.1036504 | 0.862402739 | 0.029171012 | 3.41E-14 | Northern |
| UG_3315 | 0.000116831 | 0.000148868 | 0.081787685 | 0.917946616 | 1.35E-15 | Northern |
| UG_3323 | 0.004195685 | 0.137616824 | 0.843359235 | 0.014828257 | 8.05E-15 | Northern |
| UG_3331 | 0.01087788 | 0.511900727 | 0.475513858 | 0.001707535 | 4.94E-12 | Northern |
| UG_3292 | 9.19E-05 | 0.00011649 | 0.074500822 | 0.925290784 | 5.89E-16 | North_western |
| UG_3300 | 0.141668379 | 0.576547457 | 0.267810282 | 0.009645779 | 0.004328103 | Central |
| UG_3308 | 0.004544787 | 0.067606613 | 0.868906343 | 0.058942258 | 5.73E-14 | Northern |
| UG_3316 | 0.000695127 | 0.002716718 | 0.411070395 | 0.585517759 | 1.48E-15 | Northern |
| UG_3324 | 0.000577648 | 0.002311321 | 0.398076339 | 0.599034691 | 6.27E-16 | Northern |
| UG_3332 | 0.011938858 | 0.185513042 | 0.777681045 | 0.024867055 | 1.12E-11 | Northern |
| UG_3293 | 0.003462411 | 0.019358926 | 0.703251209 | 0.273927454 | 2.72E-13 | North_western |
| UG_3301 | 0.002405581 | 0.040160829 | 0.883613154 | 0.073820436 | 1.65E-15 | Northern |
| UG_3309 | 0.005917781 | 0.121522845 | 0.845371363 | 0.02718801 | 1.24E-13 | Northern |
| UG_3317 | 0.000631698 | 0.002553968 | 0.410994663 | 0.585819671 | 8.48E-16 | Northern |
| UG_3325 | 0.000372243 | 0.000945933 | 0.238865469 | 0.759816355 | 1.42E-15 | Northern |
| UG_3333 | 0.002603009 | 0.027321202 | 0.827899325 | 0.142176464 | 8.59E-15 | Northern |
| UG_3294 | 1.39E-12 | 4.33E-13 | 5.35E-15 | 3.52E-16 | 1 | South_western |
| UG_3302 | 0.001852152 | 0.01204742 | 0.686588169 | 0.299512259 | 9.03E-15 | Northern |
| UG_3310 | 0.003522497 | 0.099717739 | 0.874071968 | 0.022687796 | 3.90E-15 | Northern |
| UG_3318 | 0.000399884 | 0.000933165 | 0.226347545 | 0.772319405 | 3.04E-15 | Northern |
| UG_3326 | 0.00063189 | 0.002791699 | 0.438721444 | 0.557854967 | 5.39E-16 | Northern |
| UG_3334 | 0.014673891 | 0.22752996 | 0.737674669 | 0.02012148 | 3.97E-11 | Northern |
| UG_3295 | 9.23E-05 | 0.000107256 | 0.068238808 | 0.931561591 | 1.04E-15 | Northern |
| UG_3303 | 0.006084142 | 0.060863511 | 0.840116574 | 0.092935772 | 6.72E-13 | Northern |
| UG_3311 | 0.001573937 | 0.014257415 | 0.763232215 | 0.220936433 | 1.27E-15 | Northern |

| | | | | | | |
|---|---|---|---|---|---|---|
| UG_3319 | 8.01E-13 | 2.47E-13 | 2.69E-15 | 1.65E-16 | 1 | Northern |
| UG_3327 | 0.000805346 | 0.003789743 | 0.486952599 | 0.508452312 | 1.06E-15 | Northern |
| UG_3335 | 0.004470725 | 0.144442164 | 0.836646655 | 0.014440455 | 1.19E-14 | Northern |
| UG_3296 | 1.48E-12 | 4.69E-13 | 5.73E-15 | 3.68E-16 | 1 | South_western |
| UG_3304 | 0.004528804 | 0.042152701 | 0.829129649 | 0.124188846 | 1.84E-13 | Northern |
| UG_3312 | 0.037339611 | 0.317097023 | 0.621321174 | 0.024242158 | 3.32E-08 | Northern |
| UG_3320 | 0.004439061 | 0.125309291 | 0.851359605 | 0.018892042 | 1.42E-14 | Northern |
| UG_3328 | 0.015782663 | 0.207072686 | 0.750664909 | 0.026479742 | 7.63E-11 | Northern |
| UG_3336 | 2.02E-12 | 6.84E-13 | 8.09E-15 | 4.70E-16 | 1 | South_western |
| UG_3297 | 0.00217346 | 0.031770618 | 0.869402702 | 0.09665322 | 1.39E-15 | Northern |
| UG_3305 | 0.005853904 | 0.140270945 | 0.833449195 | 0.020425955 | 8.94E-14 | Northern |
| UG_3313 | 0.004202746 | 0.148827261 | 0.834241006 | 0.012728986 | 7.21E-15 | Northern |
| UG_3321 | 0.005896393 | 0.141627485 | 0.832273961 | 0.020202161 | 9.28E-14 | Northern |
| UG_3329 | 0.013279919 | 0.193409939 | 0.767799989 | 0.025510153 | 2.32E-11 | Northern |
| UG_3298 | 7.53E-12 | 2.77E-12 | 3.99E-14 | 2.40E-15 | 1 | South_western |
| UG_3306 | 0.005771142 | 0.116005768 | 0.849333519 | 0.02888957 | 1.12E-13 | Northern |
| UG_3314 | 0.000297266 | 0.000589188 | 0.176166118 | 0.822947429 | 2.65E-15 | Northern |
| UG_3322 | 0.001008321 | 0.005177569 | 0.542221203 | 0.451592908 | 1.78E-15 | Northern |
| UG_3330 | 0.014188409 | 0.200257979 | 0.760121669 | 0.025431942 | 3.61E-11 | Northern |
