## Supplementary material for "Genetic diversity analysis and characterization of Ugandan sorghum": Table_S2

Table S2: Conditions of the field experiments conducted in Gross-Gerau, Germany (49°55' N, 8°29' E) in 2019 and 2020

| Year | Duration | Mean soil temperature | Mean maximum temperature | Mean minimum temperature | Absolute minimum temperature |
|---|---|---|---|---|---|
| 2019 | 08/04-22/05 | 13.8 °C | 17.7 °C | 5.6 °C | -1.5 °C |
| 2020 | 22/04-26/05 | 16.3 °C | 20.6 °C | 6.1 °C | -0.7 °C |
