## Supplementary material for "Genetic diversity analysis and characterization of Ugandan sorghum": Table_S3

Table S3: Conditions of the climate chamber experiment

| Total duration | Temperature (day/night) from sowing until emergence (12 days) | Temperature (day/night) from emergence until end of experiment (60 days) | Day/night cycle |
|---|---|---|---|
| 72 days | 25/18 °C | 13/10 °C | 14h day/10h night |
