## Supplementary material for "Genetic diversity analysis and characterization of Ugandan sorghum": Table_S4

Table S4: Genetic diversity Indices for the sorghum collection in the Uganda National GeneBank

| District | MAF | He | Ho | PIC | $F_{st}$ |
|---|---|---|---|---|---|
| **Northern region** | | | | | |
| Agago | 0.190 ± 0.0017 | 0.260 ± 0.0020 | 0.033 ± 0.0007 | 0.210 ± 0.0014 | 0.001 |
| Amuru | 0.188 ± 0.0018 | 0.258 ± 0.0020 | 0.035 ± 0.0008 | 0.210 ± 0.0015 | 0.006 |
| Gulu | 0.158 ± 0.0015 | 0.226 ± 0.0018 | 0.056 ± 0.0008 | 0.186 ± 0.0013 | 0.132 |
| Kitgum | 0.217 ± 0.0017 | 0.301 ± 0.0018 | 0.055 ± 0.0009 | 0.244 ± 0.0013 | -0.158 |
| Omoro | 0.135 ± 0.0015 | 0.198 ± 0.0017 | 0.037 ± 0.0008 | 0.166 ± 0.0013 | 0.238 |
| Nwoya | 0.177 ± 0.0016 | 0.246 ± 0.0018 | 0.041 ± 0.0007 | 0.202 ± 0.0013 | 0.053 |
| Pader | 0.170 ± 0.0016 | 0.244 ± 0.0018 | 0.061 ± 0.0009 | 0.201 ± 0.0014 | 0.062 |
| Lira | 0.201 ± 0.0017 | 0.277 ± 0.0019 | 0.030 ± 0.0007 | 0.224 ± 0.0014 | -0.064 |
| Kole | 0.185 ± 0.0017 | 0.264 ± 0.0019 | 0.044 ± 0.0009 | 0.217 ± 0.0014 | -0.016 |
| Otuke | 0.206 ± 0.0017 | 0.286 ± 0.0019 | 0.032 ± 0.0008 | 0.232 ± 0.0014 | -0.101 |
| Alebtong | 0.314 ± 0.0013 | 0.418 ± 0.0011 | 0.078 ± 0.0028 | 0.334 ± 0.0007 | -0.608 |
| Otuke | 0.206 ± 0.0017 | 0.286 ± 0.0019 | 0.032 ± 0.0008 | 0.232 ± 0.0014 | -0.101 |
| **Eastern region** | | | | | |
| Mbale | 0.188 ± 0.0017 | 0.261 ± 0.0019 | 0.048 ± 0.0008 | 0.213 ± 0.0014 | -0.004 |
| Serere | 0.189 ± 0.0017 | 0.261 ± 0.0019 | 0.031 ± 0.0007 | 0.213 ± 0.0014 | -0.004 |
| Soroti | 0.203 ± 0.0017 | 0.283 ± 0.0019 | 0.049 ± 0.0009 | 0.230 ± 0.0014 | -0.089 |
| Moroto | 0.195 ± 0.0020 | 0.271 ± 0.0022 | 0.017 ± 0.0000 | 0.221 ± 0.0016 | -0.041 |
| Ngora | 0.169 ± 0.0016 | 0.239 ± 0.0019 | 0.076 ± 0.0001 | 0.197 ± 0.0014 | 0.081 |
| Iganga | 0.197 ± 0.0017 | 0.273 ± 0.0019 | 0.027 ± 0.0007 | 0.222 ± 0.0014 | -0.05 |
| Katakwi | 0.197 ± 0.0019 | 0.273 ± 0.0021 | 0.065 ± 0.0001 | 0.222 ± 0.0015 | -0.05 |
| Kumi | 0.175 ± 0.0019 | 0.245 ± 0.0021 | 0.037 ± 0.0000 | 0.200 ± 0.0016 | 0.058 |
| **West Nile region** | | | | | |
| Arua | 0.189 ± 0.0017 | 0.265 ± 0.0020 | 0.047 ± 0.0009 | 0.217 ± 0.0015 | -0.02 |
| Yumbe | 0.171 ± 0.0016 | 0.244 ± 0.0019 | 0.031 ± 0.0009 | 0.201 ± 0.0014 | 0.06 |
| Zombo | 0.137 ± 0.0017 | 0.196 ± 0.0020 | 0.019 ± 0.0000 | 0.163 ± 0.0015 | 0.245 |

| | | | | | |
|---|---|---|---|---|---|
| **Kigezi region** | | | | | |
| Kabale | 0.185 ± 0.0022 | 0.266 ± 0.0023 | 0.059 ± 0.0009 | 0.220 ± 0.0017 | -0.024 |
| Rubanda | 0.080 ± 0.0013 | 0.124 ± 0.0016 | 0.027 ± 0.0001 | 0.106 ± 0.0012 | 0.525 |
| Rukiga | 0.147 ± 0.0017 | 0.219 ± 0.0020 | 0.045 ± 0.0014 | 0.185 ± 0.0015 | 0.157 |
