## Supplementary material for "Genetic diversity analysis and characterization of Ugandan sorghum": Table_S5

Table S5: Statistical data of the field experiments and the climate chamber experiment

| **Experiment** | **d.f.** | **genotypic variance (mean squares)** | **CV genotypes** | **residual variance** |
|---|---|---|---|---|
| field experiment Gross-Gerau 2019 (GG19) | 443 | 90.859*** | 1.08 | 30.858 |
| field experiment Gross-Gerau 2020 (GG20) | 443 | 134.408*** | 0.76 | 83.878 |
| climate chamber experiment | 254 | 50.653*** | 0.97 | 19.605 |
