## Supplementary material for "Genetic diversity analysis and characterization of Ugandan sorghum": Table_S6

Table S6: Summary of different cold tolerance traits QTL overlapping the haploblock regions

| Chromosome | QTL Id | Population | Trait Description | Start-End (bp; v3.0) | Publication |
|---|---|---|---|---|---|
| Sb02 | QCHLF2.29 | Pop1 | Chlorophyll fluorescence | 2:60411376-71013958 | Fiedler et al 2016 |
| Sb02 | QCHLF2.30 | Pop1 | Chlorophyll fluorescence | 2:61660713-69899062 | Fiedler et al 2016 |
| Sb02 | QCHLF2.28 | Pop1 | Chlorophyll fluorescence | 2:61909731-69785972 | Fiedler et al 2016 |
| Sb02 | QDMGR2.15 | Pop1 | Dry matter growth rate | 2:62050122-71145714 | Fiedler et al 2016 |
| Sb02 | QLFAR2.11 | Pop2 | Leaf appearance rate | 2:62721350-69834382 | Fiedler et al 2016 |
| Sb02 | QCHLF2.26 | Diversity set (biomass) (n=194) | Chlorophyll fluorescence | 2:68082309-68502429 | Fiedler et al 2014 |
| Sb02 | QLFAR2.2 | Diversity set (biomass) (n=194) | Leaf appearance rate | 2:68082309-68502429 | Fiedler et al 2014 |
| Sb06 | QSURV6.1 | SS79/M71 | Survival | 6:47812728-52065118 | Bekele et al 2014 |
| Sb06 | QDTEM6.5 | SS79/M71 | Time to emergence | 6:47987146-52006841 | Bekele et al 2014 |
| Sb06 | QRTLN6.1 | SS79/M71 | Root length | 6:48603544-51584852 | Bekele et al 2014 |
| Sb06 | QLFTE6.30 | Sorghum Association Panel (SAP) | Transpiration rate | 6:51474125-51656848 | Ortiz et al 2017 |
| Sb06 | QLFTE6.31 | Sorghum Association Panel (SAP) | Transpiration rate | 6:51474158-51656881 | Ortiz et al 2017 |
| Sb06 | QLFTE6.32 | Sorghum Association Panel (SAP) | Transpiration rate | 6:51474220-51656943 | Ortiz et al 2017 |
| Sb06 | QLFTE6.33 | Sorghum Association Panel (SAP) | Transpiration rate | 6:51474222-51656945 | Ortiz et al 2017 |
| Sb06 | QRTSR6.1 | SS79/M71 | Root to shoot ratio | 6:54398329-59964018 | Bekele et al 2014 |
| Sb06 | QCHLF6.17 | Pop2 | Chlorophyll fluorescence | 6:54671423-61260478 | Fiedler et al 2016 |
| Sb06 | QGERM6.8 | Sorghum Association Panel (SAP) | Germination rate (%) | 6:58270809-59806933 | Moghimi et al 2019 |

| | | | | | |
|---|---|---|---|---|---|
| Sb06 | QGERM6.9 | Sorghum Association Panel (SAP) | Germination rate (%) | 6:58270809-59806933 | Moghimi et al 2019 |
| Sb06 | QGERM6.10 | Sorghum Association Panel (SAP) | Germination rate (%) | 6:58270809-59806933 | Moghimi et al 2019 |
