## Supplementary material for "Genetic diversity analysis and characterization of Ugandan sorghum": Table_S7

Table S7: Summary table of genes identified in the haploblock regions involved in cold stress tolerance

| **Gene name** | **Chromosome** | **Start (bp; v3.0)** | **End (bp; v3.0)** | **Gene Description** | **Studied in** | **Reference** |
|---|---|---|---|---|---|---|
| Sobic.006G155700 | Sb06 | 51486353 | 51493123 | ABC-2 type transporter family protein | arabidopsis; barley | (Kang et al. 2010; Zhang et al. 2020) |
| Sobic.006G157100 | Sb06 | 51601722 | 51604971 | fatty acid desaturase family protein | rice; maize | (Wang et al 2019; Zhao et al. 2019) |
| Sobic.006G245200 | Sb06 | 58481412 | 58483311 | F-box and associated interaction domains-containing protein | rice | (Jain et al. 2007) |
| Sobic.006G245800 | Sb06 | 58535274 | 58539822 | Pentatricopeptide repeat (PPR) superfamily protein | arabidopsis | (Jiang et al. 2015) |
| Sobic.009G257900 | Sb09 | 59188770 | 59194161 | F-box and associated interaction domains-containing protein | rice | (Jain et al. 2007) |
| Sobic.009G260500 | Sb09 | 59350969 | 59367238 | Tetratricopeptide repeat (TPR)-like superfamily protein | tomato | (Zhou et al. 2021) |
